# Environmental DNA Reveals Reykjavík’s Human and Ecological History

**DOI:** 10.1101/2025.10.08.681091

**Authors:** Kurt H. Kjær, Anthony H. Ruter, Mateu Menendez-Serra, Nicola A. Vogel, Abigail D. Ramsøe, Wesley R. Farnsworth, Marie-Louise Siggaard-Andersen, Zihao Huang, Thorfinn S. Korneliussen, Karina K. Sand, Ana Prohaska, Lasse Vinner, Jesper Stenderup, Martin Sikora, Ólafur Ingólfsson, Bjarni F. Einarsson, Egill Erlendsson, Jesper Petersen, Peter C. Ilsøe, Esther R. Guðmundsdóttir, Arni Einarsson, Jón Eríksson, AEGIS Consortium, Frederikke M. Sønderborg, Ladislav Hamerlik, Scott J. Riddell, Orri Vésteinsson, Lars Wörmer, Katherine Richardson, Nicolaj K. Larsen, Ainara Sistiaga, Christoph Dockter, Morten E. Jørgensen, Robbie Waugh, Miriam Schreiber, Joanne R. Russell, Pete E. Hedley, Micha Bayer, Malcolm Macaulay, Sidsel B. Schmidt, Ronja Wonneberger, Yu Guo, Marina P. Marone, Erwang Chen, Axel Himmelbach, Martin Mascher, Nils Stein, Haoran Dong, Yuanyang Cai, Ruairidh Macleod, Lucas P. P. Braga, Chai Hao Chiu, Astrid K. N. Iversen, Michael K. Borregaard, Guðrún Þ. Larsen, Skafti Brynjolfsson, Árni D. Júlíusson, Ralph Fyfe, Laura Scoble, Max Ramsøe, Richard Durbin, Rasmus Nielsen, Yucheng Wang, Mikkel W. Pedersen, Antonio Fernandez-Guerra, David J. Meltzer, Eske Willerslev

## Abstract

Iceland was among the last large islands settled by humans, with colonization (Landnám) in the late 9th century CE (Common Era) and is often portrayed as an ecological disaster driven by the Norse settlers. Here, we revisit this narrative through environmental DNA (eDNA) and multiproxy analyses of sediment cores from Lake Tjörnin in central Reykjavík, one of Iceland’s earliest and longest-occupied settlements. Originally a marine embayment, Tjörnin became a freshwater lake around 660 CE. Our record reveals a human presence decades before the long-accepted arrival date of 877 CE, marked by the Landnám volcanic tephra. Early settlement brought livestock, barley cultivation, and other introduced taxa that enhanced nutrient cycling and unexpectedly increased local biodiversity. Contrary to the conventional view of rapid deforestation, eDNA shows that birch and willow expanded during the settlement period, likely supported by deliberate management. Pronounced ecological and land use shifts occurred after 1200 CE, but these were coeval with the Little Ice Age cooling, compounded by volcanic eruptions, storm surges, and plague, rather than anthropogenic degradation. Crop cultivation ceased, arboreal taxa retracted, and grazing pressure maintained open landscapes. Even more profound ecological changes came after c. 1750 CE with urbanization and industrialization, as wastewater discharge, heavy-metal pollution, and fossil fuel use reshaped Tjörnin’s ecosystem. These findings challenge the prevailing model of Norse-induced environmental collapse, revealing instead a dynamic human–environment relationship shaped by both cultural practices and external stressors. By applying eDNA to a long-occupied urban catchment, we demonstrate the power of genomic methods to refine settlement chronologies, reassess ecological baselines and changes, and integrate natural and cultural histories. This approach offers a model for revisiting human–environment interactions in urban centers worldwide.

## Environmental DNA and human settlement history

The early history of cities is often poorly documented, particularly regarding the environmental factors that shaped their trajectories from initial settlement to urbanization, and how human activities in turn changed the landscape^1^. Environmental DNA (eDNA)^2^ retrieved directly from sediments, can provide high-resolution records of such processes and changes, though its application has so far been confined to natural environments^3–5^. We propose that eDNA can reveal the effect of natural stressors on settlements and activities, and how humans impact their setting through, e.g. species introductions and losses, landscape modifications, and the spread of pathogens and pollutants – and thus enhance and perhaps help explain conventional social and political histories. To this end, we analysed sediment cores from Tjörnin in central Reykjavík, one of Iceland’s earliest and longest occupied settlements^6^ (Fig. 1; Fig. S1). Tjörnin played a central role in the lives of its inhabitants, and its deposits record the evolution of the city^7^.

**Figure 1.**
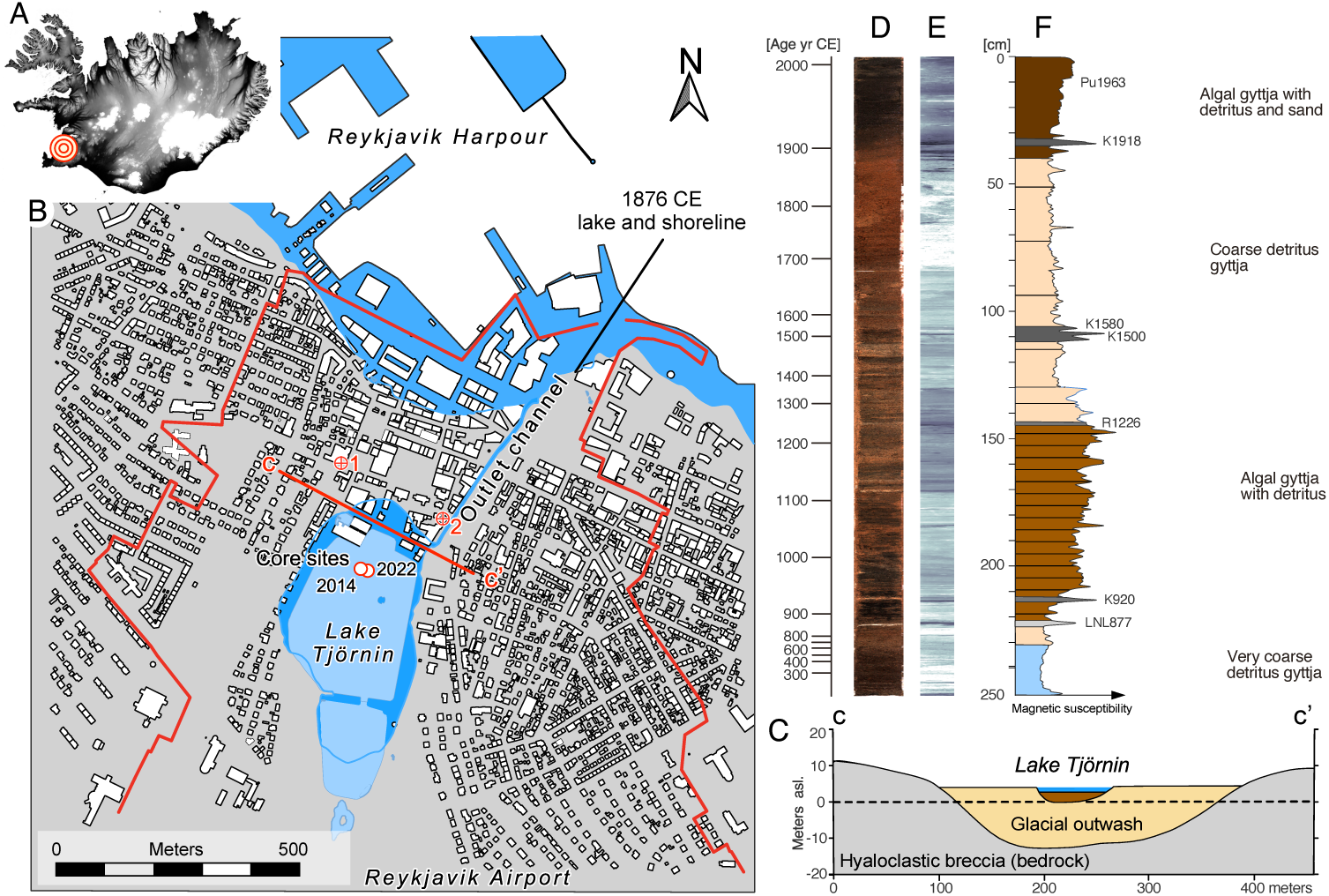
Geographic location and depositional sequence of Tjörnin, Reykjavík, Iceland. A. Location of Reykjavik on Iceland. B. Present-day cityscape of Reykjavík. Between the harbour and the Tjörnin, the old town area of Reykjavík is positioned on gravel deposits that constitute the coastal ridge separating the marine foreshore from the inland wetland area. The red polygon outlines the catchment area of Tjörnin. See supplementary information for full catchment outline. Dark blue colour reflects lake and coast shoreline in 1876 CE^108^ before major landfill deposits altered and extended the coastline several hundred meters into the fjord. Starting in the 1860s, Tjörnin was gradually filled with deposits to generate more space for housing and infrastructure in Reykjavík. This process continued into the late 19^th^ century, with various organizations and individuals filling sections of the lake for construction purposes. As a result, Tjörnin became surrounded by new buildings and roads. Red line is the geological cross section B-B’. Red crosshairs indicate location of the early settlement halls at Aðalstræti 16 (1) and Lækjargata 10-12 (2) archaeological sites mentioned in the text. C. Geological cross section (B-B’) of the north-south trending bedrock channel that probably acted as a meltwater conduit during the retreat of the Icelandic ice-sheet c. 12 cal. ka BP, Modified from Magnússson and colleagues^65^. The bedrock channel is filled with glacial outwash sediment with Tjörnin situated in the central part and bottom of the sediments beginning at 0 m asl altitude. D. Optical imagery of the 250 cm long sediment sequence at core site T22. E. X-ray image of the Tjörnin core site T22. Light areas indicate organic-rich sediment intervals, while darker areas show more minerogenic intervals. F. Stratigraphic log showing the grain-size variation based on magnetic susceptibility measurements. Blue is coastal lagoon deposits; sand colour is terrestrial lake deposits consisting of coarse detritus gyttja with faint lamination. Brown colour is a laminated algal gyttja with minerogenic detritus. Dark brown colour is very fine algal gyttja with detritus and sand.

Settlement began in the 9^th^ century with the arrival of Norse farmers who brought crops and animals from Scandinavia and the British Isles. The locality became increasingly commercial, industrial and urbanized from the mid-18^th^ century onwards and is today the most densely populated region on this ecologically sensitive North Atlantic Island^8^. Although there are written records of Iceland’s history, the earliest date to more than two centuries after human arrival. The sagas allude to earlier environmental conditions and contexts, but their reliability requires careful assessment^9^, particularly on the key issue of whether Iceland was draped by woodlands at the outset of human occupation. Thus, the idea of a blanketing forest may be more an “imaginative reconstruction” than a reliable ecological assessment^10^. This limits its use in estimating the ‘baseline’ vegetation at human arrival, and in turn its utility for calculating woodlands lost since. Moreover, palynological studies show forest distribution before settlement was more variable than assumed^11^, with some regions already treeless^12^ (Supplementary Information p.63, Figs. S42, 43). To fully understand Iceland’s environmental history including biodiversity change and resource use, native and non-native^13–15^, it must be reconstructed directly with eDNA and other paleoenvironmental proxies, as presented here.

## The Tjörnin record

Two stratigraphically overlapping sediment cores were collected from Tjörnin (64.14496° N, 21.94245° W) in 2014 and 2022 (Figs. 1, 2; Figs. S2, S3). A chronological framework for the composite core was established using dated volcanic ash layers (tephra), radiocarbon dating (^14^C), and plutonium isotope analysis (Pu) (Figs. S4-5; Table S1-S3). These yielded an age-depth model that spans from 200 CE to the present (Extended Data Fig. 1), its accuracy attested by its alignment with known historic events (Fig. 2; Supplementary Information p. 88).

**Figure 2.**
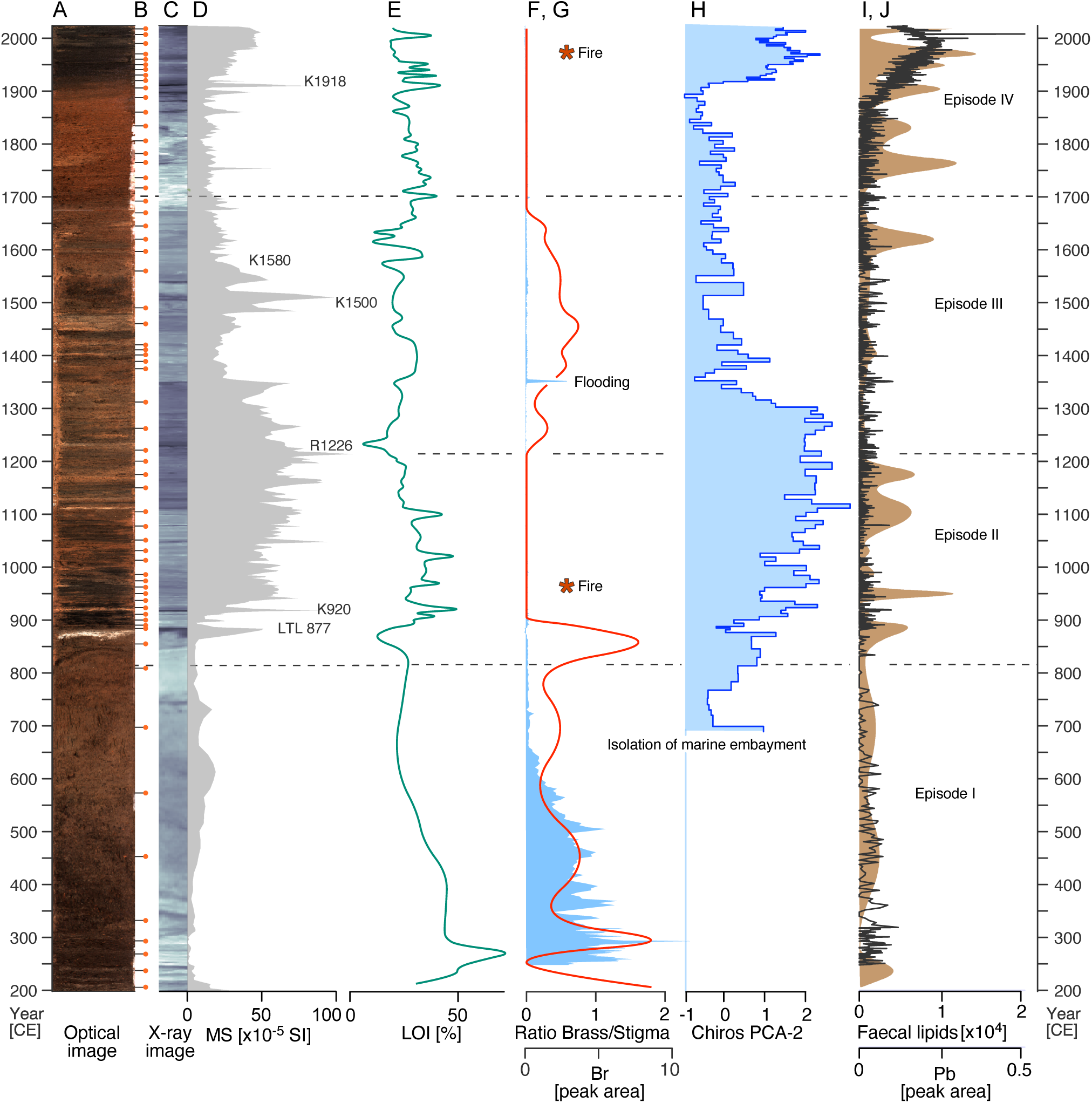
Proxy data from Tjörnin between Episodes I-IV. A. Optical image of the sediment core T22. B. Age/stratigraphical position of the 62 discrete samples i.e. eDNA, pollen and lipids. C. X-ray image of the core as proxy for density and internal structures. D. Magnetic Susceptibility (MS) as proxy for minerogenic input from the catchment. E. Loss on ignition (LOI) as a proxy for the organic content. F. G. Blue colour shows the concentration of Bromine (Br) while red coloured line indicates marine-brackish water presence derived from the ratio between brassicasterol/stigmasterol lipids. Asterix indicate burning of woody material derived from the concentration of Naphtaline (ng/g) lipids. H. The 2nd axis of the PCA (15 % of the total variation) of the chironomid assemblage as a proxy for eutrophication related to nutrient inputs and saltwater conditions. Chironomid analyses are unavailable below 700 CE, as these were conducted on core T14, which has a shorter stratigraphy. Faecal lipid signals suggest minimal input from animals in the catchment, and heavy metal concentrations are negligible. I, J. Brown is the sum of faecal biomarkers (Coprostanol, 5𝞫-stigmastanol, lithocolic acid and deoxycholic acid) reflecting herbivore and human faecal input, while black line shows the concentration (peak area) of lead (Pb) in the lake sediments.

From 62 sampled layers, we compiled a total of 21.5 billion (10⁹) DNA sequence reads (hereafter referred to as “reads”) between 16-208 million per library (median 90.6 million), which after quality control, deduplication, and low complexity filtering (Methods) resulted in 14.7 billion sequences (between 1.9-168 million, median 61.3 million) distributed across the 225 libraries prepared from double stranded DNA (Figs. S6-25). We mapped all quality-controlled and filtered sequences against comprehensive reference databases and then parsed them for taxonomic classification and DNA authentication (Methods). We used variance in the abundance and relative proportions of matched nucleotide sequences as a proxy for vegetation biomass within the lake (Extended Data Fig. 3) and its catchment (Extended Data Fig. 4). In addition to eDNA, we collected numerous other proxies that together provide a longitudinal record of shifts in catchment activity and environmental conditions. This record shows changes on varying spatial and temporal scales, the latter from abrupt events on a scale of months to years, to trends over decades and centuries. We use key inflection points in the environmental records to structure the results and discussion into episodes of environmental history (Fig. 2).

## Reykjavik before people – Episode I

The Tjörnin basin is inset into glacial outwash deposits resting unconformably atop hyaloclastic breccia. Our eDNA record shows the earliest deposits were dominated by marine animals, including echinoderms, molluscs, bryozoans, priapulids, nemerteans, hemichordates, tunicates, crustaceans, and several fish (Fig. 3, Extended Data Fig. 2, Table S5). This fauna indicates water depths sufficient to allow biomolecules from deeper-dwelling organisms to reach the basin. The marine signal is supported by vegetation and microbial records, which reveal a rich marine microbiome including fish pathogens (Fig. 3, Extended Data Fig. 9). Some ∼20 plant taxa are present, including macroalgae typical of saline coasts and eelgrass, which declined and was ultimately replaced by *Myriophyllum*. Terrestrial vegetation in the surrounding basin included salt-tolerant genera typical of coastal environments, with ryegrass, sedges, and willows dominant (Fig. 3, Extended Data Fig. 4). Thus, in its earliest phase Tjörnin was a high-salinity marine embayment.

**Figure 3.**
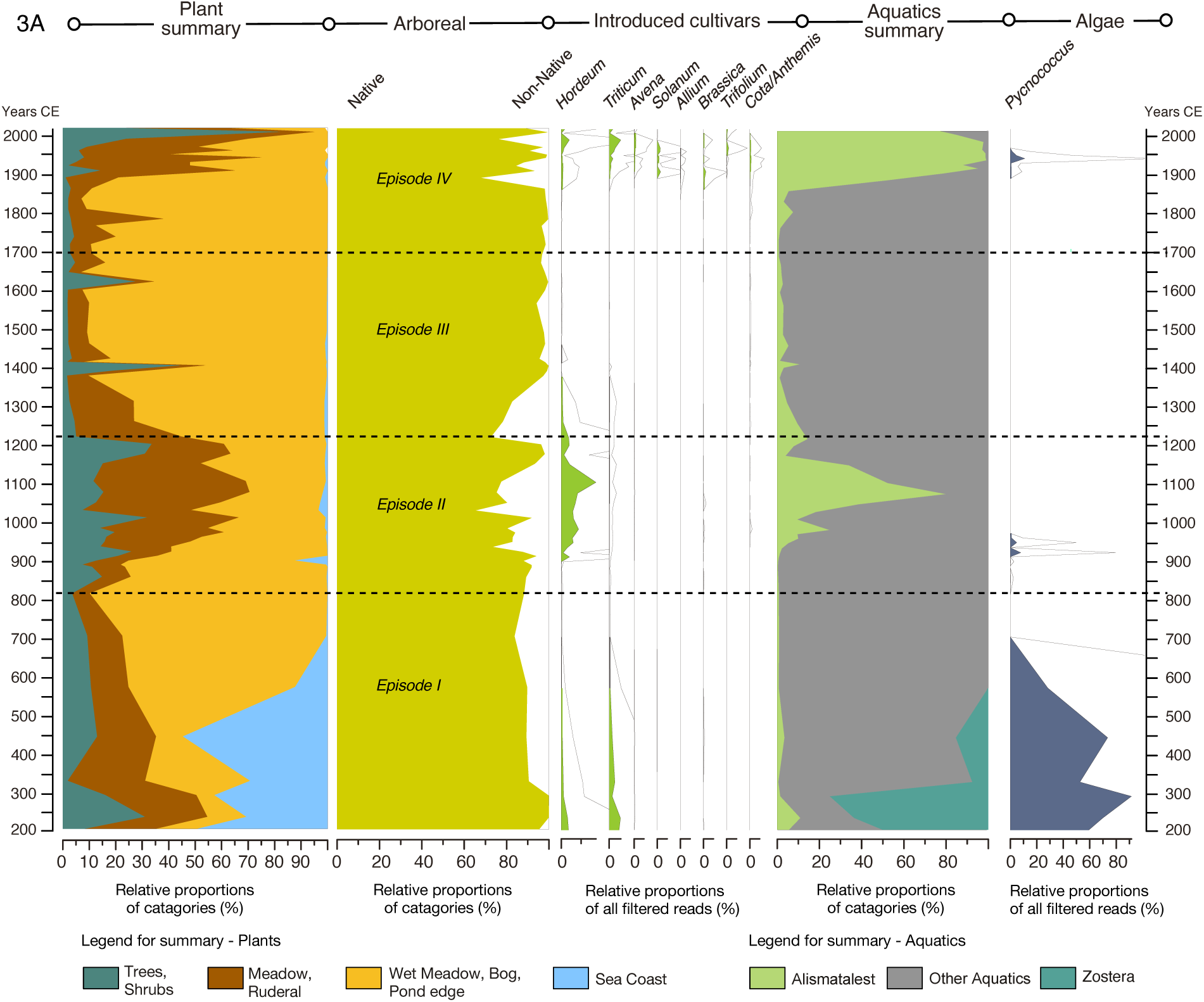
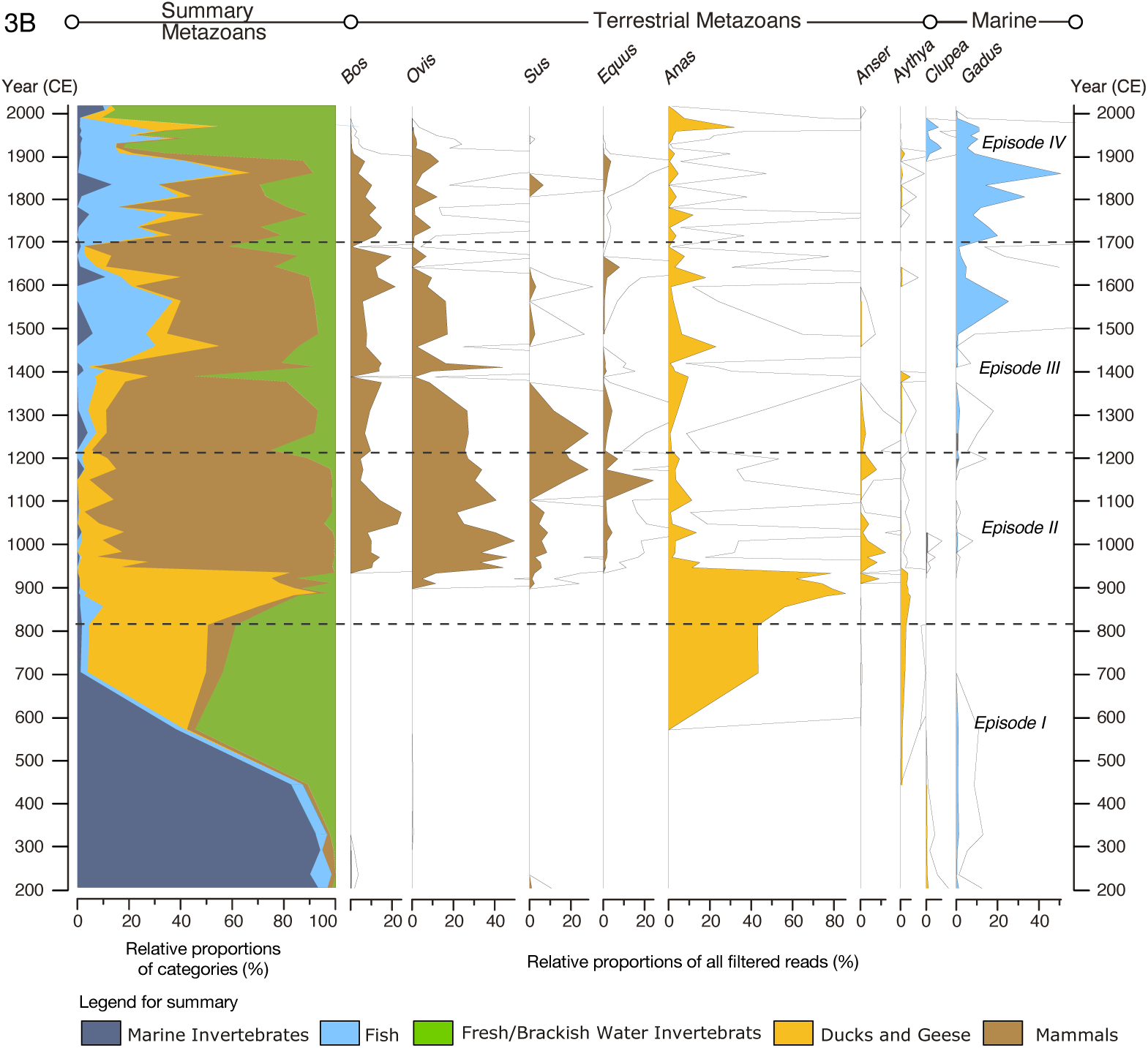
Environmental DNA data. A. Terrestrial and telmatic vascular plant genera as well as aquatic angiosperms and algae in the Tjörnin sequence (core T22), against the modelled age in calendar years. From left relative proportion of all terrestrial plants, proportion of arboreal genera; relative proportions of all filtered reads (mean read length ca. 45 base pairs) matched to introduced cultivars; relative proportion of aquatics and far right relative proportions of all filtered reads matched to algae. B. Key genera of Metazoan genera in the Tjörnin sequence, against the modelled age in calendar years. From left relative proportion of all metazoans in five categories; the relative proportions of all filtered reads (mean read length ca. 45 base pairs) matched to terrestrial metazoans; far right two commercially important fish genera, herring and cod. The hollow curve indicates a 10X exaggeration to amplify variation in minor taxa. See extended data figures 2, 3, 4 and 5 for complete data.

The basin was gradually isolated from the sea by a growing coastal barrier^16^. Proxies indicate a transition from marine to brackish to freshwater between ∼585–660 CE: organic content declined from ∼70% to ∼20% since ∼300 CE, bromine concentrations decreased, and sterol ratios shifted from marine to terrestrial sources^17^ (Fig. 2). Variations in sterol ratios also imply occasional marine incursions until the 9th century, when human arrival permanently altered the system (Fig. 2G). Finally, long-chain *n*-alkanoic acids document the rise of submerged freshwater plants, completed by ∼676 CE (Fig. S16).

During the freshwater phase, eDNA shows a shift to brackish and freshwater taxa such as chironomids (non-biting midges) and rotifers, while microbial profiles indicate changing salinity and temperature (Fig. 4A-B; Extended Data Fig. 6; Figs. S28, S29; Table S6). Terrestrial plant genera increased, freshwater algae became dominant, and ferns expanded to comprise much of the record (Extended Data Figs. 2–4).

**Figure 4.**
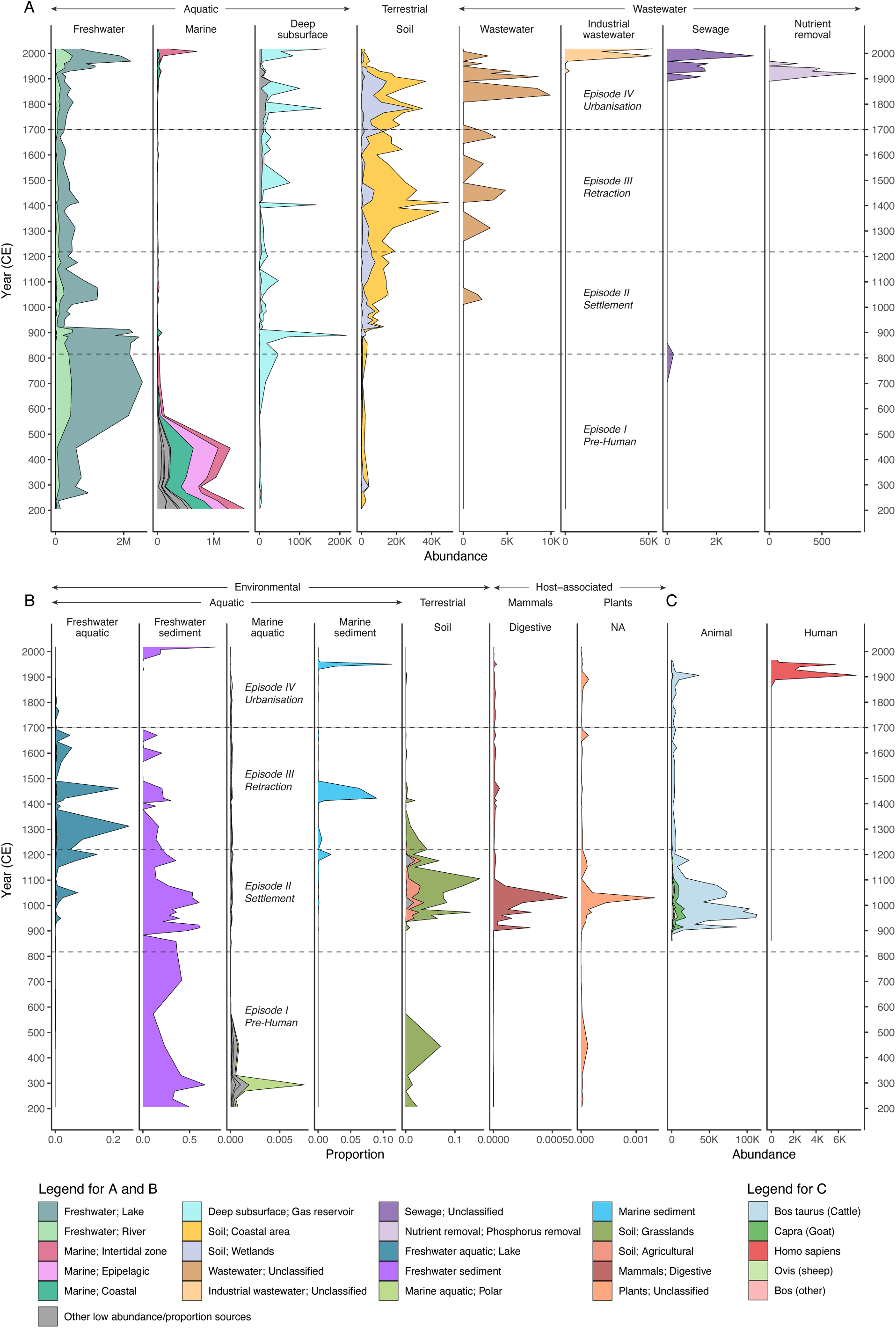
Microbial community analyses of the Tjörnin lake record. A. Viral fraction by isolation source, inferred from ecological metadata associated with reference genomes. B. Source attribution of bacterial and archaeal fractions using meta-Sourcetracker. C. Abundance of reads mapping to selected reference genomes isolated from the gut microbiome of domesticated animals and humans. Note the scale of the x-axis changes in each column.

## Human arrival and settlement of Reykjavik – Episode II

Human settlement in Iceland involved the introduction of new plant and animal taxa and the loss of native ones; a reduction in soil stability and a significant increase in erosion; and in places the replacement of forest biomes by Gramineae (grasses) and hardier, low sward vegetation better suited to an agro-pastoralist economy in this Subarctic environment^13,14,18–21^. Although the broad outlines of these changes can be seen in pollen, stratigraphic, historical and archaeological records, these are often spatially and temporally discontinuous. eDNA makes it possible to gain a more continuous, fine-grained record of the changing ecosystem to which humans adapted, and the scope and consequences of human settlement in a focal area.

Apart from a possible brief visit to Iceland by Irish monks c. 800 CE^9,22^, the beginning of a sustained human presence is traditionally associated with the Landnám tephra layer (LTL) deposited 877±1 CE^23,24^. Most early archaeological sites are situated immediately atop the LTL. However, there is archaeological evidence from two sites (including Aðalstræti in Reykjavík^25^) of turf walls, and pollen evidence at a third site of cereal cultivation, found immediately below the LTL ^7,19,26–31^

Yet, a more pronounced and earlier human presence is visible in the Tjörnin record, seen in an increase c. 810 CE in levoglucosan (an indicator of biomass burning), a rise in the early to mid-9^th^ century of wastewater-associated viruses in the microbial record (Fig. 4; Extended Data Fig. 7), an increase in faecal lipids (Coprostanol:5 β -Stigmastanol ratios) indicative of a human source (Figs. S9-15), and increase in the ambient pathogen load in the mid to late 800s CE, the latter including a component of the human gut microbiome; *Debaryomyces hansenii*, *Staphylococcus sp. AntiMn-1*, associated with the nose and skin microbiome; and *Bacillus* sp*. 1s-1* which could originate from the gastrointestinal tract of humans or their ruminant animals (Extended Data Fig. 9; Fig. S51). Sharply increasing at this time as well are certain plant-associated microbial species including *Botrytis cinerea*, and *Pseudomonas syringae* and a variety of plant symbionts, all likely linked in one manner or another to crops, farming, and the soil disturbance that follows (Extended Data Fig. 9) (Supplementary information p.84).

Previous pollen studies in the region e.g. Hallsdóttir 1987, suggested the area was a *Betula* (birch) woodland when humans arrived, which was then rapidly deforested. Yet, that apparent decline may be a statistical artefact of increases in other taxa driving down the relative representation of birch, i.e. the Fagerlind effect^32,32^. More recent studies using absolute pollen influx, do not support the reconstruction of a sudden island-wide decline in birch forest, or even a significant pre-Landnám woodland^12^ (Supplementary Information p.63, Figs. S42, S43; Table S6). Regarding the latter, the eDNA record suggests that neither *Betula* nor *Salix* (willow) were dominant components of the local pre-Landnám flora. Reads and microfossils from these taxa were matched to the genus level or higher so could match both *B. pubescens* and *B. nana*, and *S. herbacea* and *S. arctica* as well as *S. lanata* and *S. phylicifolia*. Most are shrubs or low-sward vegetation rather than trees. Nor is there evidence of a decline in forests: the Tjörnin high-resolution record of n-alkanoic acids shows no change in their relative abundance before/after Landnám, indicating there was no major reduction of woodlands in favour of grass or sedges (Fig. S16).

Instead, and counter to the conventional narrative of deforestation^33^, we see a post-Landnám expansion of woodlands in matched nucleotide sequences and pollen from birch and willow. For example, the accumulation of birch pollen grains cm^-2^ year^-1^ increases by a factor of 5.3 in the centuries from 900 to 1200 CE (Extended Data Fig. 5; Fig. S42). Given the scarcity of trees before arrival, it is reasonable to attribute this increase to human activities. Excluding livestock from disturbed substrates colonized by trees and shrubs may have been a means of nurturing sustainable wood production, providing settlers easier access to wood for framing timber and charcoal for fuel. In fact, the only native taxon in the eDNA record that experienced an adverse impact postdating human arrival were *Anas* (dabbling ducks). Their populations had risen for several centuries prior to Landnám as the onetime saltwater re-entrant ‘freshened.’ However, the duck population declined precipitously in the first few decades of the 10^th^ century, likely owing to hunting, possibly combined with the loss of their nesting habitats. Perhaps not coincidentally, *Bacillus licheniformis*, which resides in bird feathers, vanishes within150 years of human arrival, not returning until c. 1900 CE. While the duck population was declining, the introduced domestic *Anser* (grey goose) population was almost simultaneously increasing, becoming a primary avian food source alongside seabirds as seen in the zooarchaeological record (Table S11).

Genomic evidence identifies Norway, the Scottish Isles and Ireland as the principal source regions of Viking Age migrants^34^. These migrants would have sought to transplant their culture and economic strategies into this new setting, perhaps at first without fully recognizing the ecological differences^9^. Iceland was settled during the Medieval Warm Period when conditions, though never optimal, were nonetheless suitable for some cereal cultivation, especially in the south. Even so, of the cereals cultivated by the Norse in Britian and Scandinavia, only sequences matched to *Hordeum* (barley) were detected in substantial numbers in the Tjörnin record. Barley reads increased by an order of magnitude in the two to three centuries after arrival (Figs. 3; S44; Table S8), and there was a corresponding sharp increase by the late 10^th^ century in C33 alkanes, the major lipid component in barley (Fig. S9 – 15). Although there is debate whether early settlements depended on locally grown or imported barley^35^, the eDNA and lipid evidence supports the former, as does some archaeological evidence, such as the abundant evidence of charred barley from the Aðalstræti and Lækjargata sites within the catchment^35,36^ (Fig. 1).

Although *Avena* (oats) are a very minor component of these archaeological samples, there is no evidence that this cereal was cultivated intentionally. The presence of oats may be explained as seeds inadvertently mixed with barley, and possible their survival as a weed in barley fields^35^, but reads matched to oats are not detected, confirming that they were unlikely to have been cultivated. Ancient barley eDNA reads from Tjörnin were analysed against the barley pangenome, globally representative barley diversity panels, and two new Scandinavian barley landrace reference genome assemblies^37,38^ (Supplementary information: p.67). In summary, this analysis suggests that the Tjörnin ancient barley shares close common ancestry with Scottish Bere barleys, northern Europe’s oldest known landraces^39,40^. This Icelandic settlement period barley probably shared key traits with Bere including its ∼90 day growing season, making it an advantageous variety for Iceland’s brief summers.

There is a significant increase in reads mapping to introduced taxa in the Amygdaloideae subfamily which contains many fruit cultivars, including the native genus *Sorbus*, as well as imported *Prunus* (plums)^35^. The genus *Rubus* which includes blackberries, raspberries, and stone bramble, appears for the first time along with a trace amount of *Brassica* (cabbage), as well as *Cota* (chamomile) which includes an important dye-yielding cultivar (Extended Data Fig. 4).

Grasses, including *Festuca* and *Poa,* both important hay cultivar increase as well, along with several disturbance-tolerant plants. Accompanying this vegetation turnover were increases in over two dozen host-associated microbial species (Fig. 4; Extended Data Fig. 4), including *Hyaloscypha bicolor*, which primarily forms ericoid mycorrhizal symbiosis with Ericaceae plants like *Calluna* (heather) and *Vaccinium* (bilberries) (Extended data Fig. 9). Overall, terrestrial plant taxa detected by eDNA expanded after human arrival (Extended Data Fig. 4), resulting in an increase in the richness and diversity of the vegetation^41^ (Fig. 5, Extended Data Fig. 4). Given the taxa involved, the vegetation record is consistent with human use of the Tjörnin catchment for grazing, hay meadows and small-scale barley cultivation for beer brewing.

**Figure 5.**
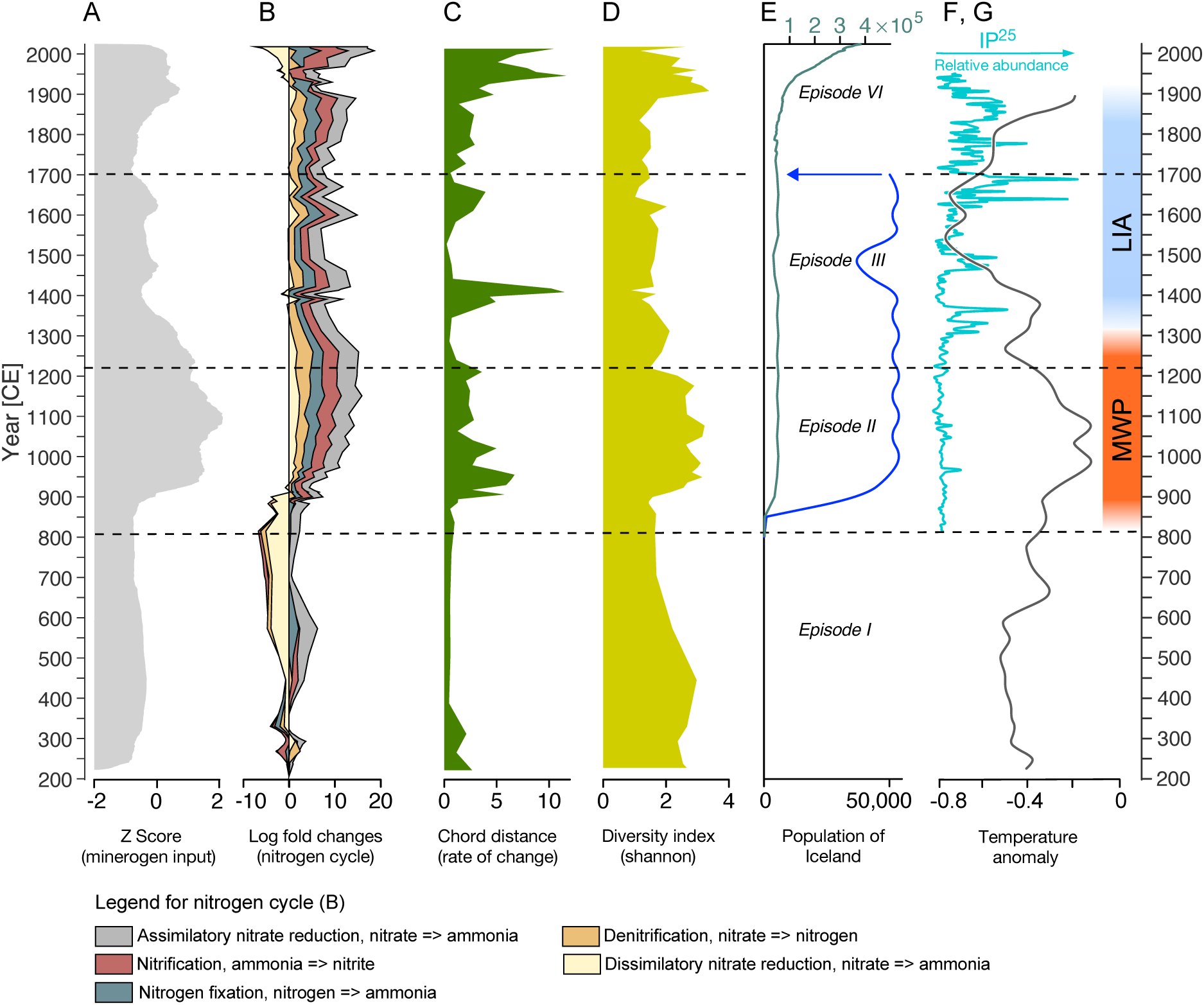
Summary diagram illustrating environmental, ecological, and demographic change. A. Smoothed minerogenic input into the lake, reflecting variations in catchment erosion and hydrological activity. Tephra layers - indicating volcanic events - have been filtered to isolate background sedimentary signals. B. Schematic representation of the complete nitrogen cycle as reconstructed microbial eDNA data, showing all five major components (nitrogen fixation, mineralization, nitrification, denitrification, and assimilation; annamox and complete nitrification were also targeted but not detected), providing insight into nutrient dynamics within the lake ecosystem. C. Rate of ecological change derived from time-series analysis of the Viridiplantae environmental DNA (eDNA) record, capturing fluctuations in terrestrial plant community composition and turnover. D. Shannon diversity index calculated from the Viridiplantae eDNA record, used as a proxy for plant biodiversity over time and indicating periods of ecological stability or disruption. E. Estimated population size of Iceland, reconstructed from historical and archaeological sources (see Supplementary Information) reflecting demographic trends and their potential correlation with environmental changes. F. Abundance of sea ice inferred from the IP25 biomarker proxy^54^, indicating past sea-ice extent and its variability as a critical component of North Atlantic climate dynamics. G. Northern Hemisphere temperature anomalies reconstructed from multi-proxy records^51^, providing a regional climate context against which local Icelandic environmental and human patterns can be interpreted. MWP: Medieval Warm Period. LIA: Little Ice Age.

The Reykjavík settlers’ livestock, *Ovis* (sheep), *Bos*, (cattle) *Sus* (pigs), and *Equus* (horses) are seen in the Tjörnin eDNA record, although reads matched to their reference genomes and those matched to associated taxon-specific bacterial microbes, archaeal species and gut microbiome records (Figs. 4; Figs. S49-S52). These appear several decades after the initial settlement.

Nevertheless, there is reason to expect the animals were an essential part of the ‘settlers kit’^9,42,43^ colonists brought to establish their traditional economy in this new setting ^20,44^. The lag in their appearance in the Tjörnin record is likely a function of low numbers: it would have taken about 20 years to build a viable herd for farms within the catchment, assuming each started with one of each sex of sheep and cattle^45^, after which there would be sufficient biomass in the basin to be detectable in the eDNA record. The increase of animals later might also, in part, reflect human demographics. The major pulse of migration to Iceland occurred after the LTL, and it was a rapid, island-wide radiation, except in the highest elevation areas^24^. Within a few decades of the LTL, there were an estimated 20,000-30,000 people^7,13,22^ (Fig. 5E; Supplementary Information p.88). Along with the expanding human population, newly imported livestock numbers likely increased as well.

The Iceland horses are related to lineages in mainland Europe (Extended Data Fig. 10), while the sheep belong to mitochondrial haplogroup B1, linked to Northern European short-tailed breeds, with one branch ancestral to the Icelandic Leader sheep, prized for their ability to cope in harsh conditions (Extended Data Fig. 10). One perhaps inevitable consequence of the expanding settlement was the accumulation of refuse which led to an increase in pathogenic microbial species. Of note is the appearance of several *Aspergillus* species, including *A. fumigatus*, a fungus found in soil and compost heaps (Extended Data Fig. 9), which thrives in damp environments, can cause indoor mould, and lead to Aspergillosis, a respiratory infection.

DNA derived from terrestrial microbes rose following Landnám. The reconstructed microbial functional profiles reflect changes in nutrient availability and cycling in the surrounding landscape driven by growing flocks and herds both fertilizing and disturbing the catchment substrates (Figs. 2, 5B; Figs. S43-44). Increased nutrient input enhanced overall nitrogen and sulphur turnover, but certain metabolic pathways became especially relevant in what was an originally nutrient-poorer system. In particular, the remarkable increase in dissimilatory nitrate reduction probably played a key role in retaining bioavailable nitrogen – otherwise lost via denitrification – thereby potentially sustaining higher productivity over extended periods (Fig. 5; Figs. S47-52).

In the decades prior to human arrival, Tjörnin was a eutrophic freshwater lake, but was still subject to occasional influxes of saltwater. These declined sharply in the settlement period, probably owing to a partial impoundment of the Lækurinn outlet (Fig. 2), attesting to water management within the Tjörnin catchment. Instead, *Potamogeton* (pondweed) became well established, consistent with somewhat less eutrophic conditions within the lake and a more turbid photic zone probably due to eroded sediment in suspension.

By the end of the Viking Age, around 1100 CE, the one-time flood of colonists had long since ceased^46,47^, and settlement patterns had stabilised after an initial period of experimentation and relocation^48^. In Reykjavík, several farmhouses lining the path to the sea had been abandoned, and the farm at Lækjargata was on its last legs. It is not known whether this reflects a decline in the population or a reorganization of the settlement pattern, e.g., if the subsidiary farms later recorded in the hills around the farm core were established at this time.

## Retraction – Episode III

After 1200 CE there are shifts in several of the Tjörnin record proxies: minerogenic content starts to decline; brackish water again re-enters; and terrestrial plant diversity declined (Figs. 3, 5). Yet, most striking in the eDNA record is the nearly complete disappearance of birch, a sharp reduction in willow, and the loss of crops and trees introduced by Norse settlers (Figs 3; Extended Data Fig. 4). These taxa were replaced in the catchment by *Carex* (sedges), *Poa* (grasses) and an increase in pond edge vegetation. We also find that microbes associated with fruit trees disappear *(Pseudomonas cerasi)* or are only occasionally detected *(Pseudomonas syringae)* (Extended Data Fig. 9). The proximate cause(s) of these changes likely included external stressors, as well as changes in human activity within the lake basin^14,49^. In terms of the latter, the several centuries’ effort to prevent marine incursions were no longer practised or at least no longer fully effective, and saltwater entered the lake (Fig. 2). A decline in minerogenic content likely signalled the expansion of nonarboreal vegetation, perhaps due to changes in cultivation or grazing patterns. Such changes reflect alterations in land management, although of uncertain cause(s).

That both native and non-native trees and introduced crops largely vanished, suggests a breaching of their ecological tolerances. As this happened to multiple genera at roughly the same time, climate change is the most likely cause. There were strong cold snaps around the end of the 13^th^ century^50–52^. These mark the onset of more pronounced and longer-lasting cooling of the Little Ice Age that followed^51,53^ (Fig. 5). This was a time of lowered air and sea surface temperatures, which led to episodic increases in sea ice buildup and duration, at times extending up to half the year^54,55^ (Fig. 5). The Little Ice Age cooling impacted people in several ways: for one, sea ice would have severely delayed spring plant growth and reduced grazing and fodder production^12^, making it more challenging to maintain livestock through the winter^14,53^. Cattle and sheep decline in the Tjörnin eDNA record after 1200 CE was similarly evident in the drop-off of faecal biomarkers (Fig. 2). In contemporary Greenland, the percentages of cattle remained stable on larger manor farms, but on small to medium sized holdings they become far less common after ca. 1300^14^. That could have been the case in Iceland as well. The Iceland zooarchaeological record shows that at some farms the sheep:cattle ratio increased, perhaps reflecting a ‘risk-spreading’ strategy that favoured less expensive animals that could nonetheless produce valuable wool^14^.

Relaxed grazing pressure and the decline of hay-meadow management and cultivation are reflected in the reestablishment of ferns as a more dominant component of the plant community. However, the impact of grazing was still sufficient to prevent tree growth^14^ (Hartman et al. 2017). The pig population declined during this period, and eventually largely faded from the eDNA and the archaeological record, except for occasional minor recurrences over the following centuries. Similarly, horses, never abundant, largely disappear from the eDNA record by the early 1300s. Microbial taxa associated with domesticated animals also showed a marked decline in both abundance and species richness after ∼1200 CE, although they remained detectable at lower levels in subsequent periods.

Barley declined sharply in the Tjörnin eDNA record in the 1200s and had altogether vanished by the 1300s (Fig. 3, Fig. Extended Data Fig. 4). Cooler temperatures during the Little Ice Age, reduced both average warmth which slowed growth, and the length of growing season. In these conditions, even ∼90-day varieties like Bere barley may fail many years to produce a crop. Furthermore, the ancient Icelandic barley is phylogenetically closest to a group of seven barleys (Figs. S44, S45), three of which have a distinctive MKK3 locus haplotype^56^. Varieties with this haplotype can convert starch better into fermentable sugars but they are also subject to pre-harvest sprouting in damp weather, leading to crop failure^56^. Repeated crop failures from cold and rain are recorded in sagas of the time^57^. Although the Little Ice Age was not a period of constant cold and rain, it need not have been, to make barley cultivation uneconomical, for even under present-day climatic conditions crop failure might be expected every 2-3 years^35,43,58,59^. With Little Ice Age temperatures in the North Atlantic more than 1° C cooler than present^60^, the failure rate would not have been less, leaving little incentive to pursue barley cultivation.

Accompanying the decline of cultivation was a decline in *Rhizophagus irregularis* (Extended Data Fig. 9), which forms a symbiotic relationship with barley and other plants, and serves to improve nutrient uptake. *Myriophyllum* recovered its pre-Landnám dominance in the lake. Most ruderal terrestrial plants declined or disappeared, as dominance was reclaimed by genera adapted to poor drainage, or shallow water (Fig. 3). With these changes in plant DNA, we see an increase in wet meadow and near-coastal terrestrial microbes in the soil (Fig. 4). Few aquatic metazoans and very few algal reads were registered during these centuries of cooling, apart from two detections c. 1550 and 1600 CE.

Significant and more abrupt environmental changes also occur in these centuries, owing to a variety of external stressors, although their effects vary. For example, while there were repeated volcanic eruptions, only a few produced ash falls in the catchment area. The nearest, a 1226 CE eruption that occurred off the coast ∼60 km southwest of Reykjavík is very pronounced in the Tjörnin magnetic susceptibility curve but left only a 3 mm thick deposit in the lake. Nonetheless, historical sources describe a very hard ‘sand winter’^9^. Ash can ruin a harvest and contaminate the soil over the short term, ash rich in fluorine poses a severe risk to grazing livestock, but over the longer term it can be beneficial in terms of increased soil nutrients.

Roughly a century later, c. 1350 CE, amidst an upward trend of brackish water seeping into the lake, there was a prominent, isolated peak in bromine (Br) and an abrupt decline in minerogenic input (Fig. 2). The spike represents a major storm surge and flood that breached the coastal barrier, introducing a massive amount of saltwater into Tjörnin. That also spike corresponds with a documented historically storm surge in 1345 CE, which damaged coastal islands and headlands, and flooded houses^61^, including the Aðalstræti farmstead, where sand and gravel were deposited above archaeological levels^36^. Around 1400 CE, the Tjörnin record reveals two significant changes: one is a marked excursion in vegetation composition (Fig. 3, Extended Data Fig. 4), which includes a brief reappearance of willow and ferns and a reduction in sedges in the catchment, and the other is a transient disruption in the microbial community that was also reflected in the nitrogen cycle. These are likely linked processes and, in turn, both a consequence of changes in human activity in the catchment.

One intriguing possibility to explain the change in human activity is the arrival of plague episodes, the first between 1402 and 1404 CE, the second in 1494-1495^49,62,63^. The Tjörnin record does not provide evidence on the impact of the plague, but it sheds light on the longstanding debate over how plague came to Iceland. The most common plague vector across Europe were rats and fleas carrying *Yersinia pestis*^64^. Yet, rats are not mentioned in Iceland’s historical annals, nor are they found in its archaeological sites^49,63^. Because rats may have gone unnoticed amidst the grain and cloth on trading vessels, or in the cod waste of fish processing stations^62^, we cannot eliminate the possibility that rats played a role in bringing plague to Iceland. However, there is no evidence of rats (*Rattus* sp.) or other potential rodent vectors in the Tjörnin eDNA record (Table S10). Nor do we detect any of the pathogenic bacteria known to be carried by rats, e.g., *Leptospira borgpetersensii* and *L. interrogans* ^64^. Furthermore, there is no evidence of *Y. pestis* in the microbial record - admittedly this could be a function of its low biomass, in absolute terms and relative to other lake-dwelling microorganisms. This negative evidence suggests the that the plague in Iceland was pneumatic rather than bubonic. In the 15^th^ and 16^th^ centuries there were few significant excursions in the eDNA record, except for an increase in cod. Their presence likely represents some form of lakeside storage and processing, possibly indicative of the growing importance of marine fishing. There was also a brief spike in willow around the mid-17^th^ century, which roughly coincides with very cold temperatures and the most massive spike in sea ice of the last millennium^54^.

## Urbanization and industrialization – Episode IV

In 1703 CE, Iceland’s first national census recorded Reykjavík as a dense settlement^8^ (Fig. 1) and with the establishment of wool factories in the 1750s, a trading licence, and the arrival of government institutions and residences, it grew into a small town. These developments marked the beginning of urbanisation, driving demand for energy and increasing activity in the Tjörnin catchment^8^. Historic records show that tenant farmers continued offshore fishing. That practice is clearly visible in the eDNA record (Fig. 3; Extended data Fig. 2), suggesting wastewater from fish processing was discharged directly into the lake, both well before and after the first cooling houses were built on the banks of Tjörnin in 1905 CE.

An Iceland-wide reintroduction of barley may have occurred in this period, as the genomes of herbarium specimens collected from northern and eastern Iceland in 1897 and 1932 CE, differ from those detected in the Tjörnin sediments. At the genome level, these barleys still show large proportions of Scottish landraces of the “Bere” type but also a proportion of Scandinavian landrace “Asplund” suggesting import from Scandinavia (Figs. S47, S48). The most striking changes occurred in the late 19^th^ to early 20^th^ centuries, when native birch, willow, and aspen are again visible in the eDNA record. The first appearance of maple and rising proportions of trees in the Maleae tribe, which includes apples and pears, likely reflect intentional arboriculture in the growing town. Several new species also appear, including potato and wheat in the late 19^th^ century, the latter increasing in the 20^th^ century along with oats and exotic imports like cacao and tea. These signals are best explained as food and beverage residues incorporated into lake sediments via urban effluent.

The eDNA record of the final two centuries is marked by a steady decline in domestic animal signals and a reduction in fish-processing residues, patterns probably linked to the expansion of urban green spaces and a shift to small-scale industrial activity. Wetland-adapted taxa such as sedges, and sundew disappeared as natural surfaces were drained, reclaimed, and paved. The lake itself became increasingly eutrophic from domestic and industrial effluent: chironomid diversity declined as aquatic macrophytes and algae reached their highest recorded diversity, (Extended Data Figs. 2 and 3; Fig. S17, S18). The collapse of *Myriophyllum* and its replacement by various genera in the Alismatales marked a major ecological shift. Planktonic and macroalgae also diversified, with earlier dominant taxa reappearing alongside the first occurrence of new groups. This period of industrialisation and urban development produced the most diverse aquatic plant assemblage in Tjörnin’s history (Extended Data Fig. 3). Microbial data likewise register urban growth, with human-associated taxa first appearing around 1890 and peaking during the early 20^th^ century (Fig. 4, Fig. S48-53).

By the late 19th century, Tjörnin had effectively become a sink for waste generated by Reykjavík’s population. Proxies show increases in faecal lipids and wastewater-associated microbes’ decades before the city’s physician declared the waters of Tjörnin unsafe for health c. 1920 CE^65^. Peat cutting and drainage of the Vatnsmýrin wetlands facilitated urban expansion, altering water flow into the lake, while new roads, buildings, and sewers reduced natural runoff^65^. The shift to fossil fuels further transformed Reykjavík. Imported coal rose sharply after 1860^66^, and its combustion is reflected in rising zinc (Zn) and polyaromatic hydrocarbons (PAHs) in sediments (Extended Data Fig. 8, Fig. S17 – 18). Concentrations of lead (Pb) also rose from the 1880s, along with sulphur, linking early industrialisation to polluted wastewater. Pb peaked around 1990 before declining after the phase-out of leaded gasoline in 2000. These chemical records mirror Iceland’s energy transitions from peat to coal and petroleum. The biological record captures the ecological consequences of this industrial growth. Increased pollutant loads coincide with rising deformities in chironomid head capsules (Fig. 2; Extended Data Fig. 8; Fig. S17 – 18), signalling stress in aquatic invertebrate populations. Together, these lines of evidence demonstrate how 18th–20th century urbanisation and industrialisation radically reshaped the ecology of Tjörnin, creating a record of pollution, eutrophication, and biodiversity turnover that reflects the environmental costs of Reykjavík’s transformation into a modern city.

## Discussion

Our data shed new light on the Viking Age environmental history of Iceland. The conventional narrative of “foolish” behaviour on the part of settlers who did not use the soil and vegetation sustainably^33^ is not supported by the eDNA and multiproxy records. Nor is this a case comparable to that of many islands in the Pacific, in which human hunting, introduced predators, land clearance, and habitat destruction resulted in rapid decimation of the native fauna^67–71^.

Instead, the evidence points to agropastoralists who adapted to their environment and created a landscape that enabled several centuries of success, rather than a Norse-induced ecological disaster^72^. Settlers cultivated crops, created pastures, and unintentionally enhanced biodiversity through increased nutrient cycling, soil formation, and woodland cover, especially birch. Increases in magnetic susceptibility and minerogenic input after Landnám reflect small-scale mobilization of sediments, distinct from the coarse-grained erosion typical of degraded rangelands near volcanic zones and glacial margins.

Iceland’s settlers introduced many non-native plants and animals, but these had limited negative impacts on native species, although they altered ecosystem dynamics. A key driver of these transformations was the introduction of large herbivores. On nutrient-poor volcanic landscapes, ruminants accelerated decomposition through their gut microbiota and redistributed nutrients inland^73^. Cultural practices such as feeding sheep with seaweed, known since the Neolithic in Orkney^74^, may have further integrated marine nitrogen into terrestrial ecosystems.

This adaptive system was not disrupted by human mismanagement but by external forcing. From the late 1200s, colder summers soon followed by the Little Ice Age resulted in the loss of native and non-native plants and likely led to the abandonment of barley cultivation, given that the crop was already on the edge of its climate envelope. Environmental stress was compounded by specific catastrophic events such as a major storm surge in 1350 CE and episodes of plague in the 15^th^ century. Given the labour demands of harvesting barley, population losses from plague may have dealt a lasting blow to farms already challenged by climatic downturns. Human impacts rose most steeply in the 18^th^ century, with population growth, peat cutting, and intensified lakeside industries such as fish processing^65^. These activities led to increasing industrial water and air pollution, which worsened over the next two centuries as coal and then petroleum became the dominant fuels.

Although Reykjavík’s history may be locally distinctive, it has implications for Iceland’s wider settlement history, that without an accurate ecosystem ‘baseline,’ would be impossible to measure the degree of deforestation and soil loss attributable to human actions or gauge other possible human impacts on the environment. Comparable multiproxy studies across Iceland are likely to reveal similarly complex and dynamic environmental histories. Such reconstructions must also integrate the role of climate, which can amplify human pressures and determine the long-term resilience of ecosystems. Environmental genomics, combined with other proxies, offers a powerful means of reconstructing the deep history of cities, from pre-human landscapes through settlement and industrialisation. Extending this approach to other historic centres such as Rome, Athens, Angkor, promises new insights into the coupled trajectories of human societies and their environments.

## Methods

### Core collection and sampling procedure for robot workflow

Sediment cores were collected from the northwest region of Tjörnin (64.14496° N; 21.94245° W) in Spring 2014 (T14-7) and again in March 2022 (T22). The cores were collected with a lightweight piston-coring system with 64 mm diameter coring tubes that allow for the collection of 200 cm long overlapping sediment cores. Coring was performed during ice cover. Coring in 2014 and 2022 yielded a 2.0 and 2.5 m long sediment sequence, respectively. The two core sites were primarily correlated through the different tephra layers and distinct patterns in the XRF element data (Supplementary information p.5, Fig. S3). Following sediment collection, core samples were stored cool (4-6 °C), packed and flown to Denmark for laboratory analysis at Globe institute, University of Copenhagen.

After the core splitting, visible stratigraphical layers, such as visible volcanic layers, were identified. Both core halves, a working half and an archive half, were marked up accordingly using adhesive tape with cm marks and then placed in a cold room. Prior to archive, samples sediment cores and other bulk samples were photographed to provide documentation of source material. The surface of the collected substrate was scraped clean using bleached metal plate and the newly exposed surface flipped with a sterile disposable spatula before the archive sample was collected by inserting a 5 ml sterile syringe with bottom cut off into the substrate, recovering 2-4 ml substrate. Cores were sampled continuously with a 1.2 cm resolution. The archival sample is collected in 4-8 ml barcoded LVL tubes. The archive tube barcode is scanned and registered along with sampling depth and organic content of the substrate. All archive samples are stored at -18°C in barcoded 24 format SBS format racks. See figure 2 for sampling resolution on absolute age scale.

The archive racks were collected in the freezer and thawed on a thawing station using RT air flow and subsequently spun down at low speed to separate potential substrate from the lids. On a bleached lab bench (not LAF) ∼180mg material was transferred by sterile disposable spatula from 4 ml archive tubes to 2 ml barcoded LVL tubes in 48 format for subsequent treatment using an automated workflow.

### Absolute age dating

#### Tephra dating

Tephra samples were extracted from specific horizons identified within Tjörnin lake cores (visually or via x-ray/XRF data). Tephra samples were cleaned for humic material, sieved (63 μm), dried, mounted in epoxy, polished and carbon-coated for geochemical analysis. Tephra geochemistry was analysed at the University of Iceland using a JEOL JXA-8230 electron probe micro-analyser (EPMA). The acceleration voltage was 15 kV, the beam current 10 nA, with a beam diameter of 5-10 μm. The standards A99 (for basaltic tephra), ATHO and Lipari Obsidian (both for silicic and intermediate tephra), were measured prior to, and after, the analyses to verify consistency in analytical conditions. Data were then inspected for, and cleaned of, anomalies and analyses with sums <95% and >101%.

#### ^14^C dating

Macrofossils for ^14^C analysis were retrieved from residues derived from 0.25 mm sieving. Plant material was identified and isolated using tweezers and a binocular microscope. Radiocarbon ages were obtained through accelerator mass spectrometry (AMS) at the Ångström Laboratory, Uppsala University, Sweden and The Aarhus AMS Center, University of Aarhus, (AAR) Aarhus, Denmark.

#### Pu dating

To support the collection and preservation of modern sediments at the sediment-water interface, bulk sediment subsamples (*c.* 2 cm^3^ each) were collected from the uppermost 20 cm of the ISL22-T1A core for plutonium radiometric dating. The peak in Pu^239+240^ is associated with nuclear weapons testing in 1963/1964 while the onset of global fall-out begins 1952^75,76^. Samples were run on an ICP-MS using a Thermo X2 quadrupole at the Northern Arizona University, following procedures adapted from Ketterer et al.^77^.

### Element composition

We used the archive core half for XRF core scanning to obtain the variation in element composition. The Itrax µ-XRF core-scanner is equipped with a Rhodium X-ray tube and a Silicon drift X-ray detector with a resolution of 70 eV, and further it is equipped with a high-resolution optical line-scan camera, a high-resolution X-ray camera for radiographic line-scan imaging, and a Bartington MS2E magnetic susceptibility sensor. For measurement, the instrument set-up was 60 kV, 30 mA, and 1000 msec for radiography imaging, and 30 kV, 50 mA and 35 sec for XRF analyses. The resolution was 200 µm for radiography, 1 mm for XRF, and 4 mm for magnetic susceptibility. The XRF data were evaluated using the Q-spec software, and core correlations and depth assignments were performed using Matlab for post processing of the scanning data.

### Pollen data

Samples of 2 cm^3^ were used for pollen analysis. Following the addition of an exotic marker grain, sediment was disaggregated, with the 10–180-micron fraction separated through sieving. Silicates were removed using the density separation method (using sodium polytungstate), followed by acetolysis to remove cellulose. Taxon identification was made using published keys^78,79^ and a modern reference collection. Pollen accumulation rates were calculated based on taxon concentrations (calculated using the exotic marker count) and the sediment accumulation rate from the core age-depth model.

### Conventional and high-resolution molecular biomarkers

Sediment samples were freeze-dried prior to extraction. Lipids were extracted using an Ethos Ex microwave extraction system, with 15 mL of dichloromethane (DCM) and methanol (MeOH) (9:1, v/v). The microwave was operated with a program consisting of a 10 min ramp to 70 °C (1000 W), 10 min hold at 70 °C (1000 W), and 20 min cool down. The supernatant was collected after 24 h and the extraction was repeated 3 times. The total lipid extract (TLE) was then concentrated using a turboVap (Biotage). After measuring the TLE, an aliquot (20%) of the TLE was derivatized via acid methanolysis (0.5 M HCl in methanol) diluted in H_2_O and extracted with hexane:DCM (4:1 vol/vol). The resulting extract was then silylated using N,O-bis(trimethylsilyl) trifluoroacetamide (BSTFA_Merck).

Biomarkers were characterised by gas chromatography–mass spectrometry (GC-MS) with an Agilent 7890B series gas chromatograph interfaced with a flame Ionization Detector (FID) and an Agilent 5977B mass selective detector. A 1-μL aliquot of derivatized extracts was injected in splitless mode onto a 60 m DB-5MS fused-silica column (60 m × 0.25 mm internal diameter and film thickness of 0.25 μm). The GC oven temperature was programmed as follows: 60 °C injection and hold for 2 min, ramp at 6 °C min to 300 °C, followed by isothermal hold of 20 min. The transfer line, source, and quadrupole temperatures were set at 320 °C, 270 °C, and 150 °C, respectively, and the source was operated at 70 eV. Internal injection and response factors were calculated for faecal biomarkers using representative standards (epiandrosterone, 1-nonadecanol). Procedural blanks were run to monitor background interferences. Each biomarker was identified using the software MassHunter (Agilent) by comparison with a spectral reference database (NIST) and published spectral patterns.

In addition, high-resolution records of molecular biomarkers were obtained for the period of human arrival and settlement. Therefore, a 16 cm interval spanning across the LTL (615-917 CE) was subsampled and divided into four sections, which were freeze-dried and embedded following the procedure described by Alfken et al.^80^. Mass spectrometry imaging (MSI) can be used for the µm scale mapping of target molecules on intact sample surfaces. Recently we demonstrated that MSI of non-disturbed sediments can be used for paleoenvironmental studies; using matrix-assisted laser desorption/ionization coupled to Fourier transform-ion cyclotron resonance-mass spectrometry we visualized the spatial distributions of archaeal glycerol dibiphytanyl glycerol tetraether (GDGT) lipids. There is a pressing need for implementing sample preparation procedures that allow exploiting the full potential of sediment MSI. Here we present a suite of sample preparation steps, optimized for the analysis of GDGTs in marine sediments. It considers the crucial requirements for successful MSI and optional combination with elemental imaging via micro X-Ray Fluorescence Spectroscopy (µ-XRF). Preservation of the sediment’s spatial distribution is achieved with freeze-drying and subsequent embedding in a mixture of gelatin and carboxymethyl cellulose. This enables sectioning the sample into sequential slices from 20 to 500 µm in thickness. Thinner sections showed enhanced signal intensity in MSI, but elemental mapping by µ-XRF is more accurate for thicker sections; 100 µm thick slices provide satisfactory results for both analyses and are recommended for congruent elemental and biomarker imaging. When applied to the uppermost ∼5 cm of marine sediment from a Santa Barbara Basin box core, the optimized sample preparation yields reproducible ultra-high-resolution GDGT records from sequential slices, thus demonstrating the robustness of the method. Congruent µ-XRF results aid the establishment of a contextual framework regarding supply of terrigenous and marine detritus as well as the assignment of molecular data to annual layers ^78^. Thin slices (100 µm) from these samples were obtained using a Medite Cryostat M630 and placed on indium-tin-oxide-coated glass slides. These slices were used for elemental mapping on a Bruker M4 Tornado 300 and for mass spectrometry imaging (MSI) of molecular biomarkers on a 7T Bruker solariXR mass spectrometer. MSI was performed with 200 µm raster resolution and around 40 spots per horizon, resulting in ∼ 2000 spots per cm depth. Our targets were levoglucosan and its isomers, which can be used as a tracer of fire^81^, as well as long-chain *n*-alkanoic acids (Supplementary information p.16). For levoglucosan detection, spectra were recorded in continuous accumulation of selected ions mode, with a *m*/*z* range of 150-250. X-y-coordinates, intensity and signal-to-noise ratio of the target molecule (*m*/*z* 185.042) were exported for each spot by DataAnalysis. Using MSIAlign, these abundance maps were aligned with an image of the core to place them in the correct depth scale. Subsequently the same software was used to generate a downcore profile of average abundance of levoglucosan. Therefore, individual spots were binned into 200 µm horizons and their intensities were averaged. Horizons were only considered when they contained at least ten spots in which the target compound had been successfully detected.

### Chironomid analysis

Subsamples were wet sieved through a 90 µm sieve. The material was sorted with forceps under a stereo microscope at 20-40x magnification. Head capsules from chironomids were dehydrated in 99% ethanol and mounted on microscope slides using Euparal. The identification was carried out following Brooks *et al.*^82^ and Andersen at al.^81^.

### DNA recovery

Sampling, DNA extraction and library preparation were performed in dedicated clean rooms at the Globe Institute, University of Copenhagen, Denmark. Sediment subsamples (≤0.2 grams) were transferred to SAFE® 2D barcoded 2 ml tubes (LVL technologies) previously loaded with 200 μL Omni 0.1 mm, 200 μL Omni 0.5 mm Ceramic Bulk Beads and the lysate buffer. All samples were then bead-beated on a FastPrep-96™ running two rounds at 1600 rpm for 30 seconds, interspaced by a 30 seconds break, samples were next incubated shaking overnight at 37°C. DNA was extracted on a Tecan Fluent DreamPrep 780 using the Qiagen® MagAttract® Power Soil Pro kit following the manufacturer’s protocol with the following modifications: The DNA lysate volume was reduced to 240 μl for downstream purification, and adjusting the binding buffer ratio to 1:3 lysate:binding buffer. All extracts were lastly converted to unique double-stranded (N=225) dual-indexed Illumina sequencing libraries following the protocol as described previously (Mccoll et al: Steppe Ancestry in western Eurasia and the spread of the Germanic Languages, accepted Nature 2025 and lastly sequenced on the NovaSeq 6000 S4 flowcells running 2×101 bp paired-end.

### Metagenomes

We hereafter parsed all QC reads through the Holi pipeline for taxonomic profiling and DNA damage estimation (for details about version of tools and options set see SI). To increase resolution and sensitivity of our taxonomic assignments, we supplemented the RefSeq (92 excluding bacteria) and the nucleotide database (NCBI) with a recently published Arctic-boreal plant database (PhyloNorway). All alignments were hereafter merged using samtools and sorted by coordinate and parsed through filterBAM reassign and filter functions to refine alignments and generate reference-wide statistics. Cytosine deamination frequencies were then estimated using metaDMG, by first finding the lowest common ancestor across all possible alignments for each read and then calculating the nucleotide misincorporations (deamination frequencies) due to DNA damage at each taxonomic level (See SI for details on damage filtering). In parallel, we computed the mean read length as well as number of reads per taxonomic nodes. Using the DNA damage for plant species with >500 reads allowed us to create a DNA damage model which we used to filter all eukaryotic taxa as described in the SI. In addition, using biological and technical replicates we investigated inter and intra sample variability, in which we found the species compositions to be highly similar. We therefore merged all biological and technical data by layer to increase the statistical power of the genome wide statistics (see SI).

Microbial taxonomic profiles were reconstructed following the approach described in Fernandez-Guerra et al. (2023). Briefly, quality-controlled and dereplicated reads were mapped against a curated, pseudo-de-replicated database encompassing Bacteria, Archaea, Viruses and Eukaryotic chloroplasts and mitochondria (see SI for details). Alignments were then refined and filtered using filterBAM (reassign and filter functions), which was also used to estimate Lowest Common Ancestor (LCA) genome-normalize abundances (lca function, see SI for details and specific parameters). Post-mortem DNA damage was estimated with metaDMG in LCA mode. Functional profiles were reconstructed by estimating the metabolic potential of archaeal and bacterial reference genomes using Anvi’o^83^. Potential source environments were inferred separately for Bacteria and Archaea versus Viruses. Bacterial and archaeal sources were inferred using meta-Sourcetracker^84^ with a curated set of modern metagenomes representing multiple biomes. Viral sources were inferred based on ecological metadata available in IMG/VR v4.

### Ancient microbial species screening

We performed ancient microbial DNA screening for specific pathogen and host-associated taxa using a previously described computational workflow. Briefly, the workflow performs initial taxonomic classification using *KrakenUniq*^61^ against a database of complete bacterial, archaeal, viral, protozoan genomes in the RefSeq database, followed by genus-level read mapping and authentication stages. The targeted genera for these stages included the complete set of fungal genera in the database (n=165 genera), as well as selected bacterial (n=40) and protozoan (n=41) genera including known human pathogens.

Each targeted genus with n≥150 unique *k*-mers assigned was included in the read mapping stage. An initial list of putative microbial “hits” (sample/species combinations) was obtained by using the following criteria to filter the resulting alignments:

- Number of reads mapped with MQ20 or above ≥ 50
- Average read ANI ≥ 0.965
- Ratio of observed / expected breadth of coverage ≥ 0.8
- Relative entropy of read start positions ≥ 0.9
- 5’ C>T deamination rate ≥ 0.01

This initial list was subjected to Bayesian inference of ancient DNA damage rates using *metaDMG,* and the species with the highest read ANI showing evidence for aDNA damage (5’ damage rate estimate ≥ 0.03, Z ≥ 2) was selected as the putative origin of the ancient microbial DNA reads for each genus and sample. This approach therefore results in a single, best-matching species for each genus/sample combination. While this has been shown to guard against false-positives in the context of screening for human pathogen DNA with closely related non-pathogenic species within the same genus, it will result in false negatives if ancient microbial DNA deriving from more than one species within the same genus is present in the sample, as could be expected for environmental samples. To account for this and for a possible lack of aDNA damage evidence due to lower read numbers, we extended our putative species list by also reporting the presence of authenticated species in samples where the respective species had lower read ANI rank and/or non-significant aDNA damage profile.

## Statistical analyses of virus data

To test if virus and host community diversity were changing across the different episodes, we performed non-metric multidimensional scaling (NMDS) analyses with PERMANOVA using the R package vegan followed by a pairwise comparison between groups using the R package rvaidememoire. In both cases 999 permutations were considered, and for the pairwise comparison Pillai test with False Discovery Rate for *p*-value correction was used. Next, to identify the individual viruses significantly contributing to the changes across episodes, an indicator species analysis was performed using the multipatt function from the R package indicspecies with 999 permutations.

## Phylogenetic placement of animal mitochondrial DNA

Reference phylogenetic trees for the 4 target animal taxa were first constructed. This was done by first aligning the collected mitochondrial genomes of each taxon using MAFFT^85^. A consensus sequence for each taxon was then generated using EMBOSS^86^. Informative SNPs were identified by simulating pair-ended reads from the collected genomes using ART^87^, mapping the simulated reads against the consensus using BWA^88^, calling variants using GATK^89^ and filtering the SNPs with BCFtools^90^. The SNPs set was then converted to multi–FASTA alignments and used for inferring ultrametric trees using BEAST^91^, to work as the reference tree. Ancient eDNA reads were thereafter mapped against the consensus genome of each taxon using BWA, with taxon-specific mitochondrial reads identified through the eDNA competitive mapping pipeline. The identified reads were then trimmed to remove the effects of deamination damage, remapped with BWA against the consensus genome, and filtered with SAMtools^88^. PathPhynder^92^ was applied to place these eDNA reads on branches of the reference phylogenetic tree based on SNPs supports/conflicts of each read. More details can be found in the Supplementary Information.

## Bere genome assembly

A genome assembly of the barley variety Bere Unst was conducted by combining PacBio HiFi reads with Omni-C data as described in Jayakodi et al.^37^. All sequence data have been deposited at the European Nucleotide Archive^93^ under the BioProject: PRJEB91373 and the genome assembly under the accession number: GCA_965637285.

## Asplund genome assembly

We searched for European barley varieties derived or closely related to landraces described before 1950 in the Directory of Barley Cultivars^94^. We selected accessions that had available sequencing data^95–97^. to compare to previous data we had and to try to maximize capturing haplotypes widely dispersed during breeding history. The selected accession (Asplund, BCC1451, DOI: 10.25642/IPK/GBIS/638469) was grown under controlled greenhouse conditions with a 16-hour light/8-hour dark photoperiod at 25 °C during the day and 18 °C at night. Humidity was maintained at 60%, and plants were irrigated weekly with a nutrient solution. Leaf tissue samples were collected from young plants (2–3 weeks old) for DNA extraction. High molecular weight (HMW) DNA was extracted from a single Asplund plant using the MACHEREY-NAGEL NucleoBond HMW DNA kit, following the manufacturer’s protocol. DNA concentration was assessed using the Qubit 2.0® dsDNA HS assay (Invitrogen, Waltham, USA), and fragment size distribution was evaluated with the Agilent Femto Pulse System (Agilent Technologies, Santa Clara, USA).

For HiFi sequencing, the HMW DNA was sheared into ∼20 kb fragments using a Megaruptor device (Diagenode) at speed setting 30. HiFi SMRTbell libraries were prepared using the Pacific Biosciences SMRTbell Express Template Prep Kit. Final libraries were size-selected to a narrow 17–20 kb range using the SageELF system with a 0.75% agarose gel cassette (Sage Science), in accordance with the manufacturer’s instructions. HiFi circular consensus sequencing (CCS) reads were generated on the PacBio Revio platform (Pacific Biosciences) following standard operating procedures (https://github.com/pacificbiosciences).

PacBio HiFi reads were assembled using hifiasm (v.0.19.9)^98^. The resulting raw unitigs p_utg (N50, 413 kb) were scaffolded using the RagTag pipeline (v2.1.0)^99^, with Morex V3^100^ as the reference genome. Genome completeness and consensus accuracy were evaluated using Merqury (v.1.3)^101^. Additionally, BUSCO (Benchmarking Universal Single-Copy Orthologs; v5.8.2)^102^ was employed with the embryophyta_odb10 database to evaluate gene space completeness of the assembly.

## Ancient barley variant calling and core1000 SNP analysis

Variant calling was done in a two-step process using BCFtools v1.15^103^ on .bam files from the CGG3016635 sample libraries (x and y). Initially, per-position genotype likelihoods were computed using bcftools and the command *bcftools mpileup --min-MQ 20 --min-BQ 20 --count-orphans --skip-indels --adjust-MQ 50 -d 10000*, outputting results in uncompressed BCF format. Subsequently, variant calling was performed using bcftools call, identifying single nucleotide polymorphisms (SNPs). The resulting SNP matrix was merged with a pre-existing matrix comprising 1000 diverse domesticated barley samples^97^. A total of 4,901 SNPs, for which the ancient sample had genotype data, were retained for phylogenetic analysis. Following filtration, ancient SNP variants and the Core1000 lines were combined into a single SNP diversity matrix. This matrix exclusively included SNP positions derived from the ancient samples. Lastly, a phylogenetic tree was computed from the merged SNP diversity matrix using TASSEL v5.2.94^104^. The Neighbour-Joining method was applied to reconstruct the phylogenetic relationships.

## Ancient barley variant calling and domesticated barley sample analysis

Reads from the ancient sample (sample x) were mapped to the MorexV3 genome assembly (Mascher et al., 2021) using Minimap2(Li et al., 2021). The resulting BAM files were sorted, and duplicate reads were marked using Novosort (version 3.06.05) (https://www.novocraft.com/products/novosort/). Variant calling was performed with BCFtools ^103^ (version 1.15.1) using the command mpileup -a DP,AD -q 20 -Q 20 --ns 3332. The --variants-only option was omitted in bcftools call to retain genotypes at all sites. The resulting SNP matrix was merged with a pre-existing matrix comprising 116 high-coverage domesticated samples^38^. Prior to merging, the SNP matrix had been filtered to exclude sites with a missing rate greater than 20%. A total of 19,338 SNPs, for which the ancient sample had genotype data, were retained for phylogenetic analysis. An identity-by-state (IBS) genetic distance matrix was calculated using PLINK^105^(version 1.9). This matrix was used to construct a neighbour-joining (NJ) tree with Fneighbor (http://emboss.toulouse.inra.fr/cgi-bin/emboss/fneighbor), part of the EMBOSS package^86^. The resulting tree was visualized using the Interactive Tree Of Life (iTOL) platform^106^.

## Ancient barley coverage analysis

Coverage analysis was conducted on ancient samples: CGG3016635, CGG3016637, CGG3016639, and CGG3016641. Primary alignments were extracted using SAMtools (v1.15)^103^ by excluding secondary alignments (SAMtools option -F 256). Genome size files, indicating chromosome names and lengths, were generated from the indexed primary alignment BAM files using samtools idxstats. These were converted into BED files specifying genomic intervals for subsequent depth analysis. Coverage depth and breadth were computed using samtools depth with the BED intervals file to ensure comprehensive analysis across each chromosome. Depth values were processed with AWK scripts to calculate the coverage breadth, represented as a percentage of bases covered (Coverage Breadth %).

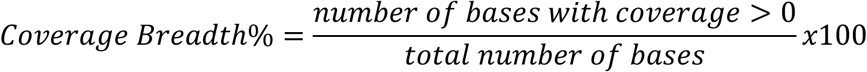

## Likelihood estimates of the proportions of reference genomes

We here describe the method used to identify the origin of the barley variety based on the short read data. This method is based on a slight modification of that described by Pipes and colleagues^107^. This method provides maximum likelihood estimates of the proportions of reference genomes represented in a pool of short reads:

Notation

For each read *i* = 1, …, *N* and reference genome *j* = 1, …, *K*:

- *n_ij_*: number of sites compared (read length)
- *d_ij_*: number of mismatches observed
- *ð_j_*: true proportion of reads from genome *j* (to be estimated)

The goal is to estimate the mixture proportions *π* = (*π*_1_, …, *π_K_*) subject to Σ_*j*=1_*^K^* π*_j_* = 1 given the observation of alignments of reads to all genomes.

Likelihood Model

This method assumes a single error rate *ε* for all comparisons and symmetric errors among all nucleotides. For read *i* and genome *j*, the conditional likelihood is then:

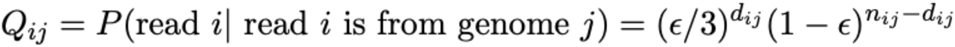

The full log likelihood is obtained as

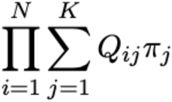

We optimize this likelihood function using an EM algorithm with SQUAREM optimization.

EM Algorithm

E-step: Calculate posterior probabilities

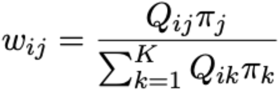

M-step: Update proportions

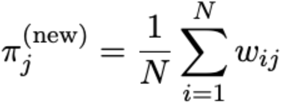

The error rate is similarly estimated using the EM algorithm using the following M-step:

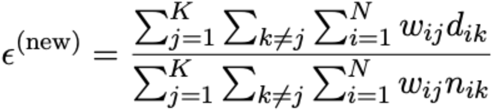

## Code availability

All codes used will be available at https://github.com

## Data availability

The datasets generated and analysed in this study are included in the paper, in the Extended Data Figures, its Supplementary Information, and the Source Data files.

## Acknowledgements

EW, KHK, DJM, MWP, ADR, AHR, MMS, MLSA, TSK, KKS, AP, AFG, LV, JS, MS, PCI, NKL, NAV, LPB, CHC, YW and the AEGIS Consortium were supported by the Novo Nordisk Foundation (AEGIS project), the Danish National Research Foundation – Center for Ancient Environmental Genomics (CAEG), and The Carlsberg Foundation (grant CF20-0398 to WRF and grant CF18-0024 to YW) within HM Queen Margrethe II’s and Vigdís Finnbogadóttir’s Interdisciplinary Research Centre on Ocean, Climate and Society (ROCS). WRF and KR were supported by The Carlsberg Foundation (CF20-0398, ROCS). JP received support from the Danish National Research Foundation, and LW was funded by Germany’s Excellence Strategy (EXC-2077, Project 390741603) and the AEGIS project. AS was funded by the Carlsberg Foundation – ROCS project. CD and MEJ received support from Carlsberg Foundation grants CF15-0236, CF15-0476, and CF15-0672. RW, MSch, JRR, PEH, MB, and MMac were funded by the Scottish Government RESAS project KJHI-B1-2. MM and NS were supported by the German Federal Ministry of Education and Research (SHAPE-P3, FKZ 031B1302A). MKB received funding from Carlsberg Foundation grant CF19-0695. A.K.N.I. was supported by the OAK foundation (OFIL-20-095). YW acknowledges funding from the Excellent Research Group Program for Tibetan Plateau Earth System (No. 42588201), and the CAS Youth Interdisciplinary Team Fund. DJM is supported by the Quest Archaeological Research Fund and the Potts and Sibley Foundation. Data analysis was supported by the National Key Scientific and Technological Infrastructure project “Earth System Numerical Simulation Facility” (EarthLab, 2023-EL-ZD-000111), the National Supercomputer Center in Wuxi utilising the computational resources of the Sunway TaihuLight Supercomputer, and the High-Performance Computing (HPC) clusters at Southern Methodist University. Hrefna Dögg Gunnarsdóttir is thanked for her support in field-data collection. We thank Miles Langner and Hassan Abdullah Aamir for generation of the MSI data. We acknowledge Research Computing at the James Hutton Institute for providing computational resources and technical support for the “UK’s Crop Diversity Bioinformatics HPC” (BBSRC grants BB/S019669/1 and BB/X019683/1), the use of which contributed to the results reported herein (please cite: https://doi.org/10.1002/ppp3.10607). We also thank Sylvia Swetik and Manuela Knauft for technical assistance with DNA work, the HiFi Lab assistants Frida Wolter Mastek and Victoria Schiller, and technicians Maria Madrona and Suzan Rabia Coskun from the GeoGenetics Sequencing Core staff for their invaluable support.

## Author Contributions

KHK, JE, and EW conceived the idea. KHK, DJM, and EW conceptualized, designed and supervised the study and wrote the manuscript with input from co-authors. EW, KR and KHK funding acquisition. JE also contributed to logistics, selection of all core sites, and collection of the 2014 cores. MWP: supervision, ancient DNA analysis, data interpretation, writing methods and supplementary information on eDNA. AHR: Data interpretation, figure generation, writing, review and editing. ADR: Bioinformatics analyses, data curation, review and editing. WRF: Stratigraphic interpretation, radiocarbon dating coordination, chronological analysis, review and editing as well as collection of the 2022 cores. MMS: Microbial community analysis, including replicate assessment, taxonomic profiling, ordination-based analysis, source tracking and functional profiling. MLSA: XRF analyses and interpretation, supplementary information writing, review and editing. TSK: Bioinformatic pipeline development, supervision, review and editing. KKS: geochemical data interpretation, writing – original draft and review. AP: Data interpretation, review and editing. AFG: Metagenomic analysis, review and editing. LV: Ancient DNA authentication, lab supervision. JS: Sampling development and sample preparation. MS: Genomic data supervision, interpretation, and writing – review and editing. OI: Regional geological context, review and editing. BFE: Archaeological context, review and editing. EE: Data interpretation of palaeoecological data, review and editing. JP: Entomological analysis, figure preparation, review and editing. PCI: field sampling. ERG:Tephrochronology, data interpretation. AE: Paleoecology, interpretation of lake sediment data. FMS: Ancient DNA lab work, review and editing. LH: Entomological analysis, multivariate statistical analysis, figure preparation, review and editing. SJR: Summary of archaeological and historical records for Reykjavík, data interpretation, review and editing. OV: Archaeological synthesis, writing – review and editing. LW: High-resolution molecular biomarkers: investigation, formal analysis, visualization. Writing – review and editing. KR: Climate context synthesis, review and editing. NKL: Geochronological data supervision, review and editing. AS, FMS: Lipid biomarker analysis, interpretation, and review. NAV: Ancient cereal DNA authentication, review and editing. CD: Conceived and supervised the cereal data analysis. Wrote cereal paragraph, reviewed and edited text. Funding. MEJ: Conceived and designed cereal data analysis executed and interpreted the analysis of ancient barley reads in a contemporary context. Wrote cereal paragraph, reviewed and edited text. RW, MSch, JRR, PEH, MB, MMac, SBS: Contributed with the Bere barley genome assembly. RWon: Sample analysis support. YG: NJ tree with ancient samples. MPM: Selection of Asplund. EC: Asplund draft assembly. AH: PacBio sequencing. MM: Conception, supervision of data analysis, funding. NS: Conception, supervision of data generation, funding. YW, ZH: Phylogenetic placement for domestic animals. HD, YC, RM: Bioinformatics support and analysis. LPPB: Virus community diversity data analysis, figure generation, results interpretation and writing. CHC: Data interpretation and writing (arbuscular mycorrhiza/ericoid mycorrhiza section). AKNI: Analysed and interpreted microbial species data, writing and editing. MKB: Contributed writing on island ecology, discussion section. GTL Tephra chronology and supervision. SB: Regional environmental context. ADJ: Historical sources and context data and supplementary writing. RF, LS: Data analysis and interpretation (pollen), review and editing. RD: Supervision of phylogenetic placement for domestic animals, reviewed and edited text. RN: Population genomics interpretation, review and editing. AEGIS Consortium Contributed comparative genomic and reference data.

**Correspondence and requests for materials should be addressed to Kurt H. Kjær** and Eske Willerslev

## Additional Information

All reads to mapped human DNA sequences from the Tjörnin sediments has been omitted from the bioinformatic analyses. Supplementary Information is available online for this paper.

## Extended data figures and captions

**Extended Data Figure 1.**
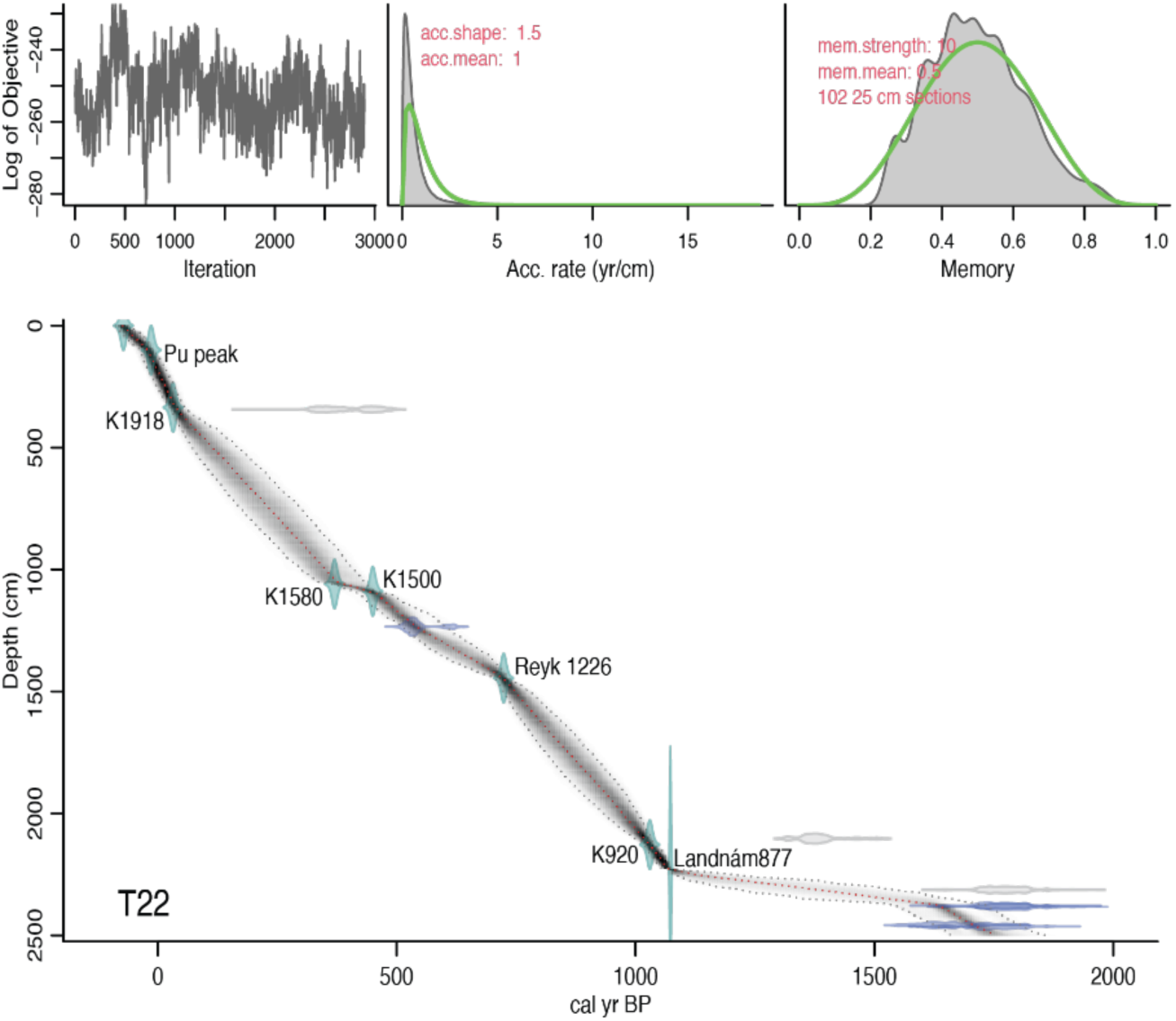
Age depth model for T22 stratigraphy. Tephra and radiocarbon age details are given in Table S1 and S2.

**Extended Data Figure 2.**
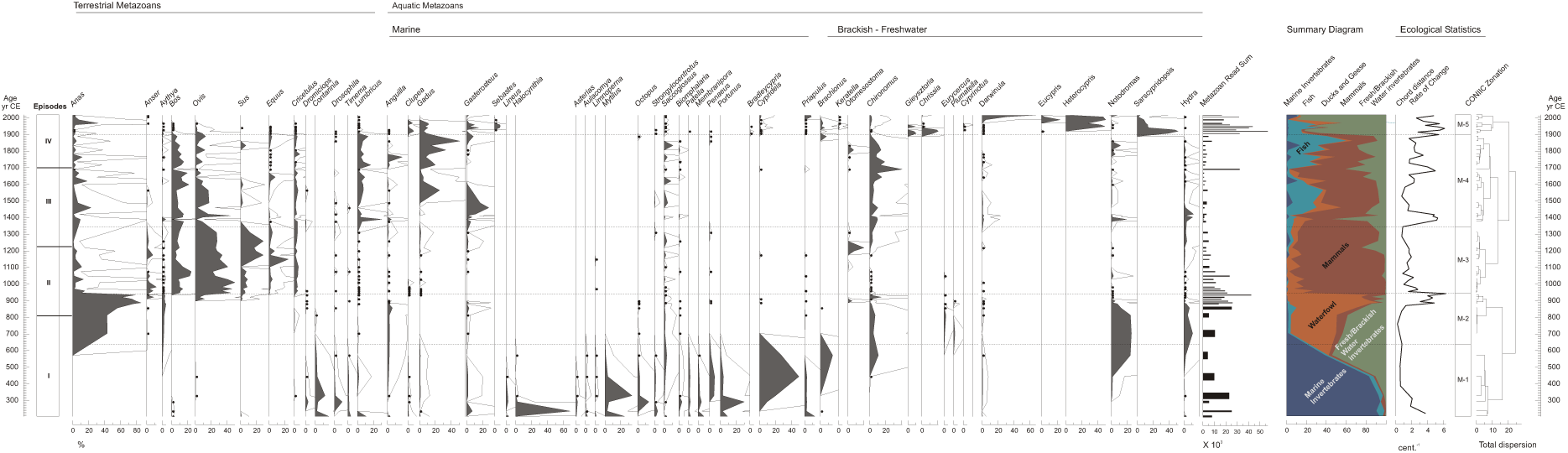
Metazoan genera. The relative proportions of all filtered reads (mean read length ca. 45 base pairs) matched to Metazoan genera in the Tjörnin sequence, against the modelled age in calendar years. The episodes used to describe the sequence in the main text are on the left next to the age scale. The hollow curve indicates a 10X exaggeration to amplify variation in minor taxa. Bullets indicate trace values ≤ 0.5%. To facilitate interpretation, the genera are ordered primarily based on habitat (e.g. terrestrial, aquatic, fully marine or freshwater) and secondarily by taxonomy (e.g. osteichthyes/tetrapods to the left followed by echinoderms, molluscs, arthropods etc… for each habitat). Non-native taxa are not segregated. The read sums of all metazoans are plotted as histograms with widths adjusted to reflect the deposition interval of the sample. The sequence has been divided into five bio-zones (M-1 to M-5) produced from a stratigraphically constrained cluster analysis using the information statistic as a distance measure. All metazoans were included in the zonation. These zones and the cladogram drawn from the dispersion distance are presented in the final column. The summary diagram on the right-hand margin plots variation in the summed read proportions of composite categories (i.e. marine Invertebrates, fish, ducks and geese, all mammals, and fresh/brackish water invertebrates).

**Extended Data Figure 3.**
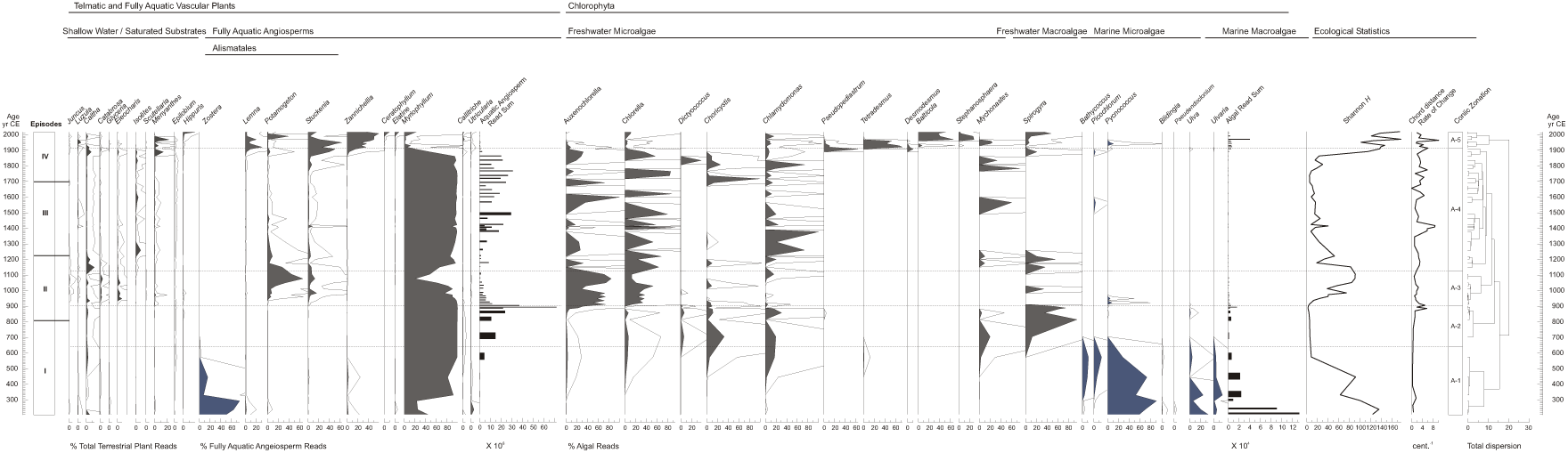
Aquatic Angiosperms and Algae. The relative proportions of all filtered reads (mean read length ca. 45 base pairs) matched to aquatic viridiplantae (fully aquatic vascular plants and chlorophyte) in the Tjörnin sequence, against the modelled age in calendar years. The episodes used to describe the sequence in the main text are on the left next to the age scale Several genera that grow in shallow water or on permanently saturated substrates are included on this diagram as well, to the left of the fully aquatic angiosperms for comparison. The hollow curve indicates a 10X exaggeration to amplify variation in minor taxa. The genera are ordered primarily by taxonomy and subdivided by habitat (e.g. flowing plants or algae and freshwater or marine, respectively). All marine taxa are shaded blue. The read sums of aquatic angiosperms and algae are plotted as histograms with widths adjusted to reflect the deposition interval of the sample. The sequence has been divided into five bio-zones (A-1 to A-5) produced from a stratigraphically constrained cluster analysis using the information statistic as a distance measure. All fully aquatic taxa were included in the zonation. These zones and the cladogram drawn from the dispersion distance are presented in the final column.

**Extended Data Figure 4.**
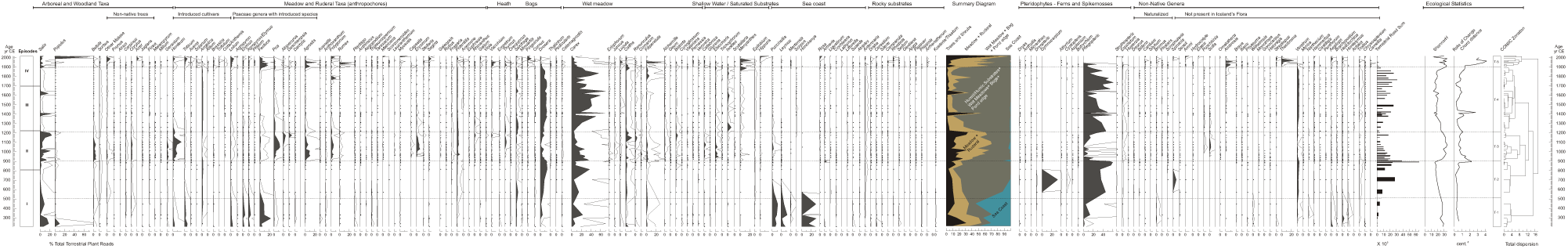
Terrestrial Viridiplantae. Relative proportions of all filtered reads (mean read length ca. 45 base pairs) matched to the 275 terrestrial and telmatic vascular plant genera in the Tjörnin sequence, against the modelled age in calendar years. The episodes used to describe the sequence in the main text are on the left next to the age scale. The hollow curve indicates a 10X exaggeration to amplify variation in minor taxa. Bullets indicate trace values ≤ 0.1%. The genera are ordered by habitat (e.g., arboreal and woodland taxa, wet meadow, heath, bogs) except for pteridophytes and non-native genera. The read sums of are plotted as histograms with widths adjusted to reflect the deposition interval of the sample. The sequence has been divided into five bio-zones (T-1 to T-5) produced from a stratigraphically constrained cluster analysis using the information statistic as a distance measure. All taxa were included in the zonation. These zones and the cladogram drawn from the dispersion distance are presented in the final column to the right of the Shannon diversity index and a rate-of change statistic.

**Extended Data Figure 5.**
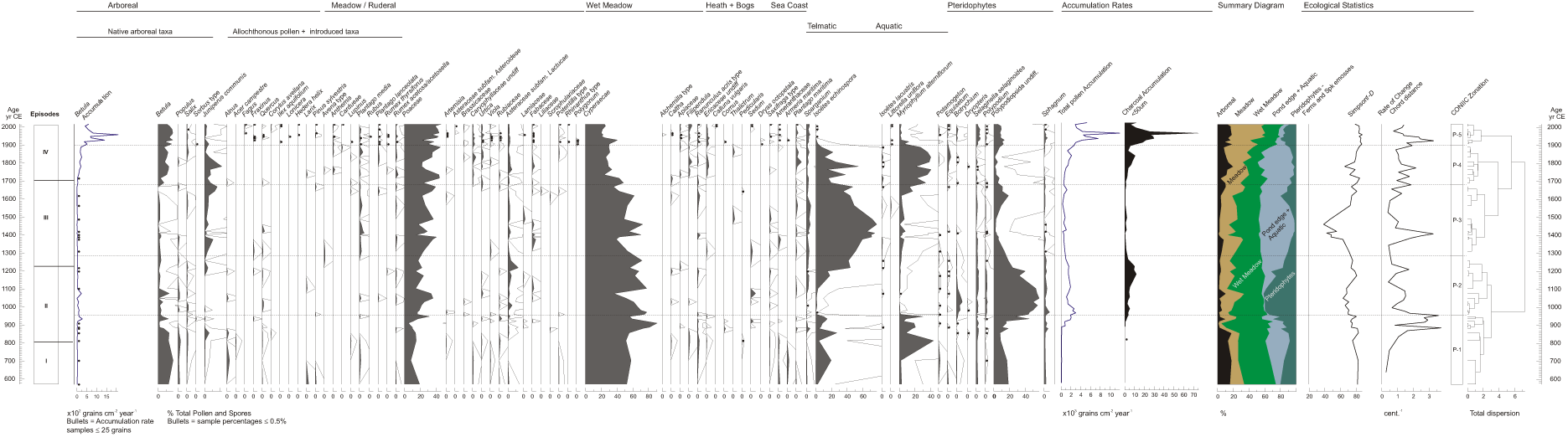
Pollen and Spores. The percentages of pollen and spores in the Tjörnin sequence, against the modelled age in calendar years. The episodes used to describe the sequence in the main text are on the left next to the age scale. The hollow curve on the percentage plot indicates a 10X exaggeration to amplify variation in minor taxa. Bullets indicate values ≤ 0.5%. The genera are ordered by habit / habitat and general taxonomy (e.g. arboreal, heath and bog, pteridophytes). Bullets indicate trace values ≤ 0.5%. The modelled accumulation for *Betula* total pollen and total charcoal are included for comparison with the percentages. The sequence has been divided into five bio-zones (P-1 to P-5) produced from a stratigraphically constrained cluster analysis using the information statistic as a distance measure. All pollen and spores taxa were included in the zonation. These zones and the cladogram drawn from the dispersion distance are presented in the final column. The summary diagram plots variation in the summed read proportions of composite categories (i.e. Arboreal, Meadow, Aquatic and Pteridophytes).

**Extended Data Figure 6.**
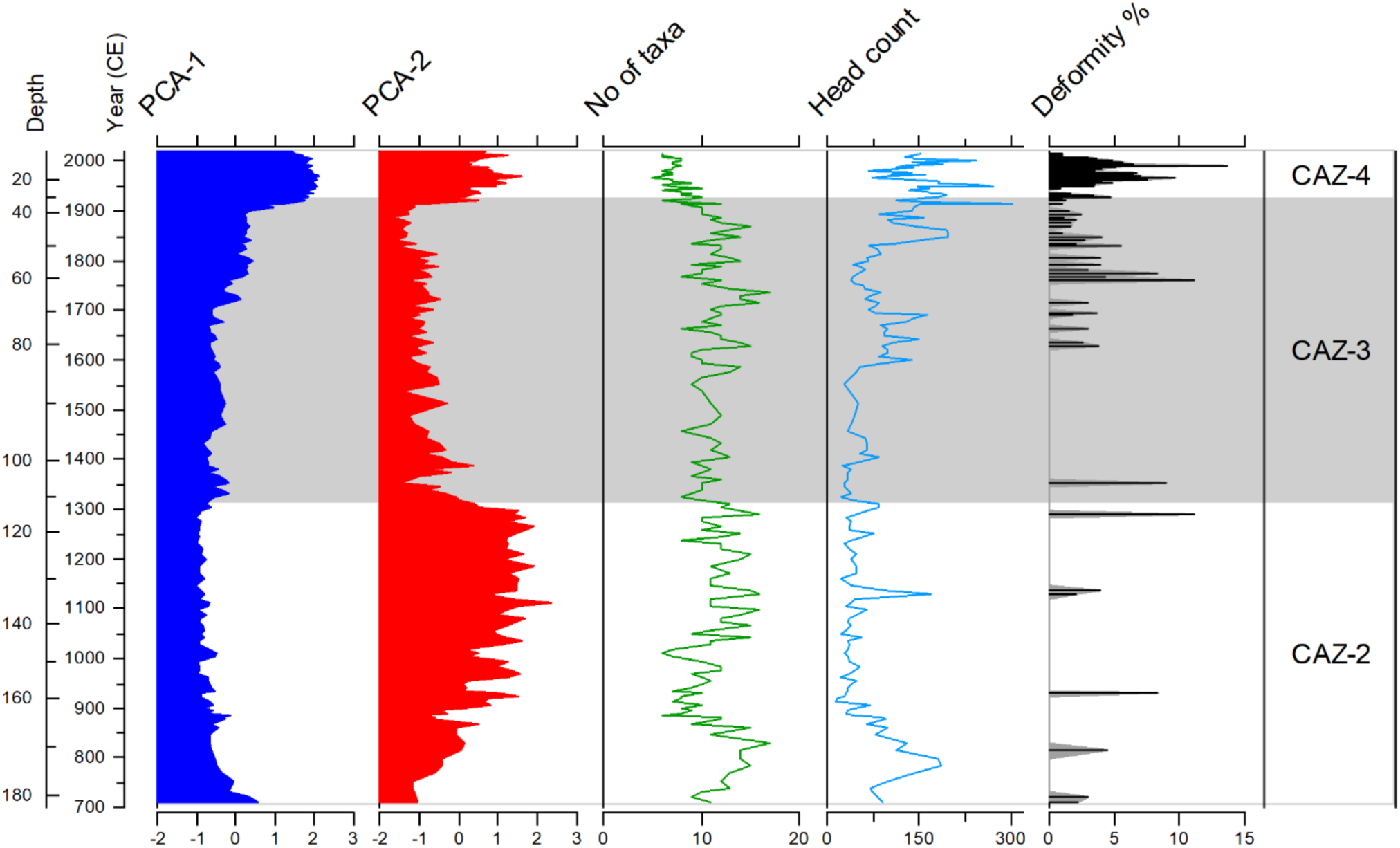
Chironomid analyses (core T22). Summary table of PCA axes scores, number of taxa, chironomid head count, and percentage of morphological deformities in the chironomid assemblages. CAZ 2 – 4 refer to Chironomid assemblage zones; Zone 1 with extremely low amount or no chironomid head capsules is not included.

**Extended Data Figure 7.**
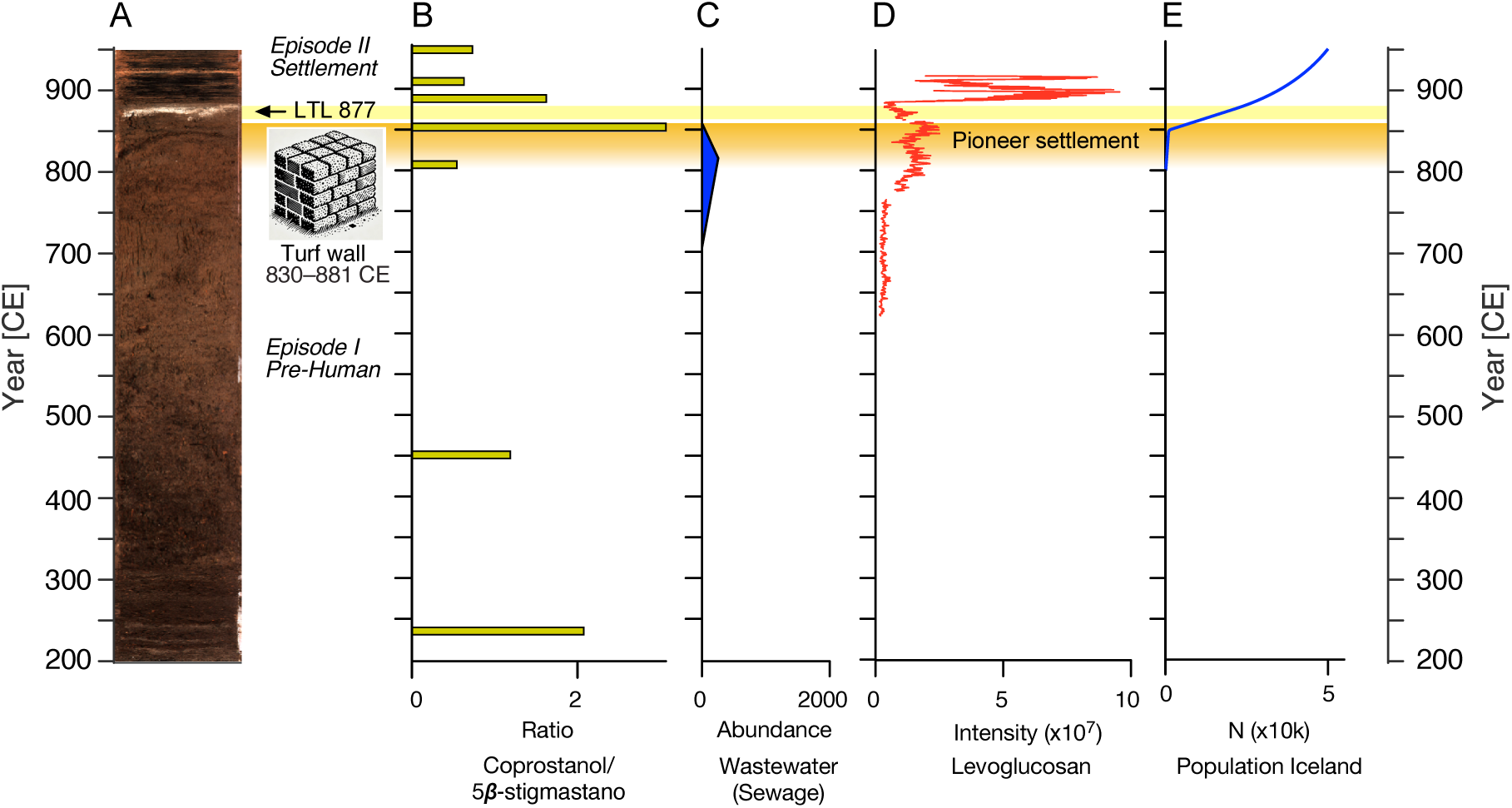
Pioneer settlement indicators. A. Optical image of the T22 core with placement of episodes and archaeological evidence of pre-Landnám turf wall construction^7^. B. Coprostanol/5𝞫-stigmastanol lipid ratio indicative of human waste. C. Abundance of sewage wastewater from microbial sources. D. Reconstruction of fire use based on mass spectrometry imaging of levoglucosan intensity. E. Estimate of the Icelandic population (Supplementary information p.88).

**Extended Data Figure 8.**
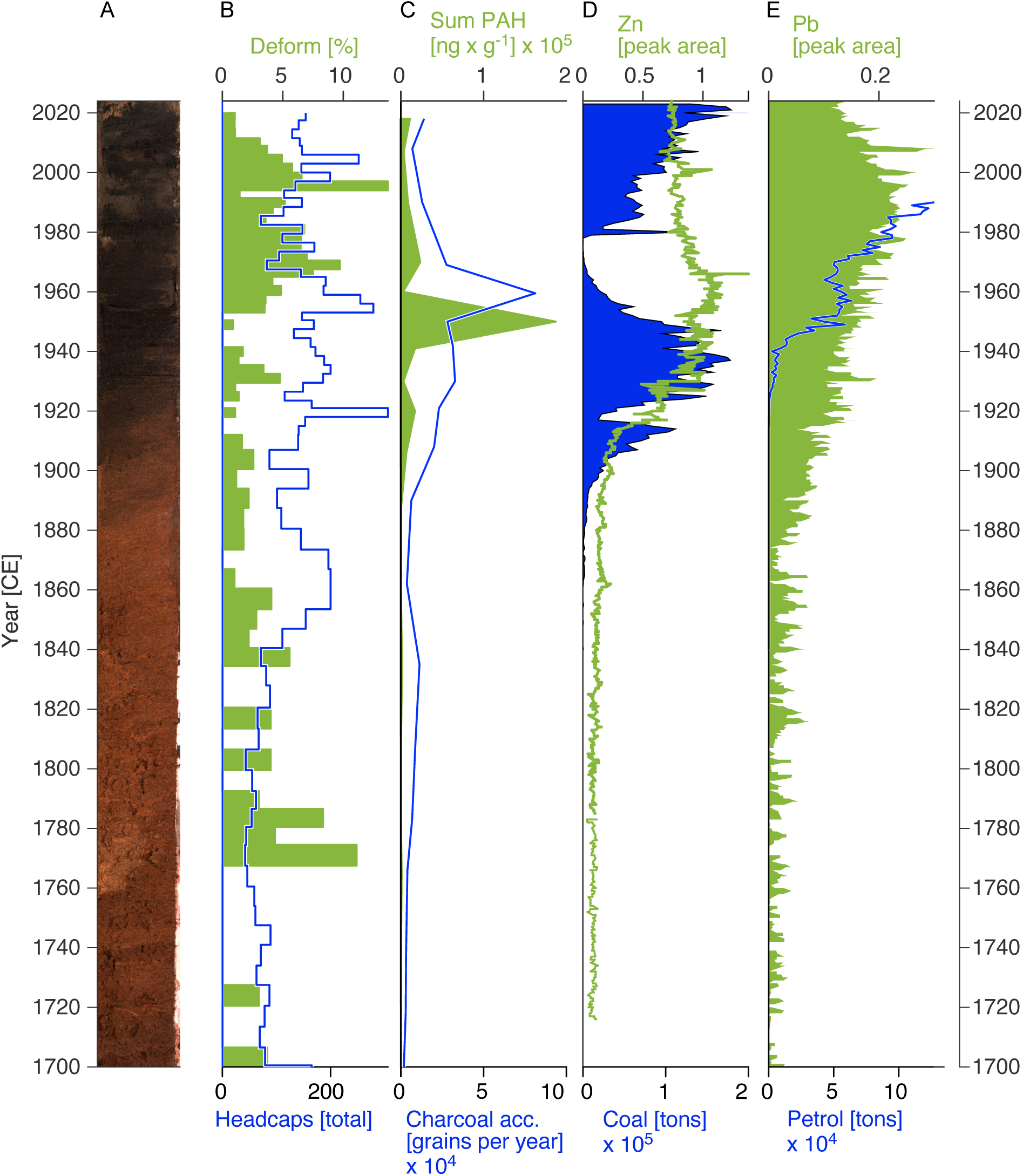
Pollution indicators from Reykjavík since 1700 CE – Episode IV. A. optical image from core T14. B. Blue: counts of total head capsules from chironomid taxa. Green: percentage of head capsules that are identified as deformed indicative of environmental stress. C. Green: sum of Polyaromatic Hydrocarbons (PAHs). Blue: Charcoal counts from pollen analyses. D. Blue: Zink (Zn) content compared to (Green) import of coal to Iceland. E. Green: Lead (Pb) content compared to (Blue) import of gasoline (petrol) to Iceland.

**Extended Data Figure 9.**
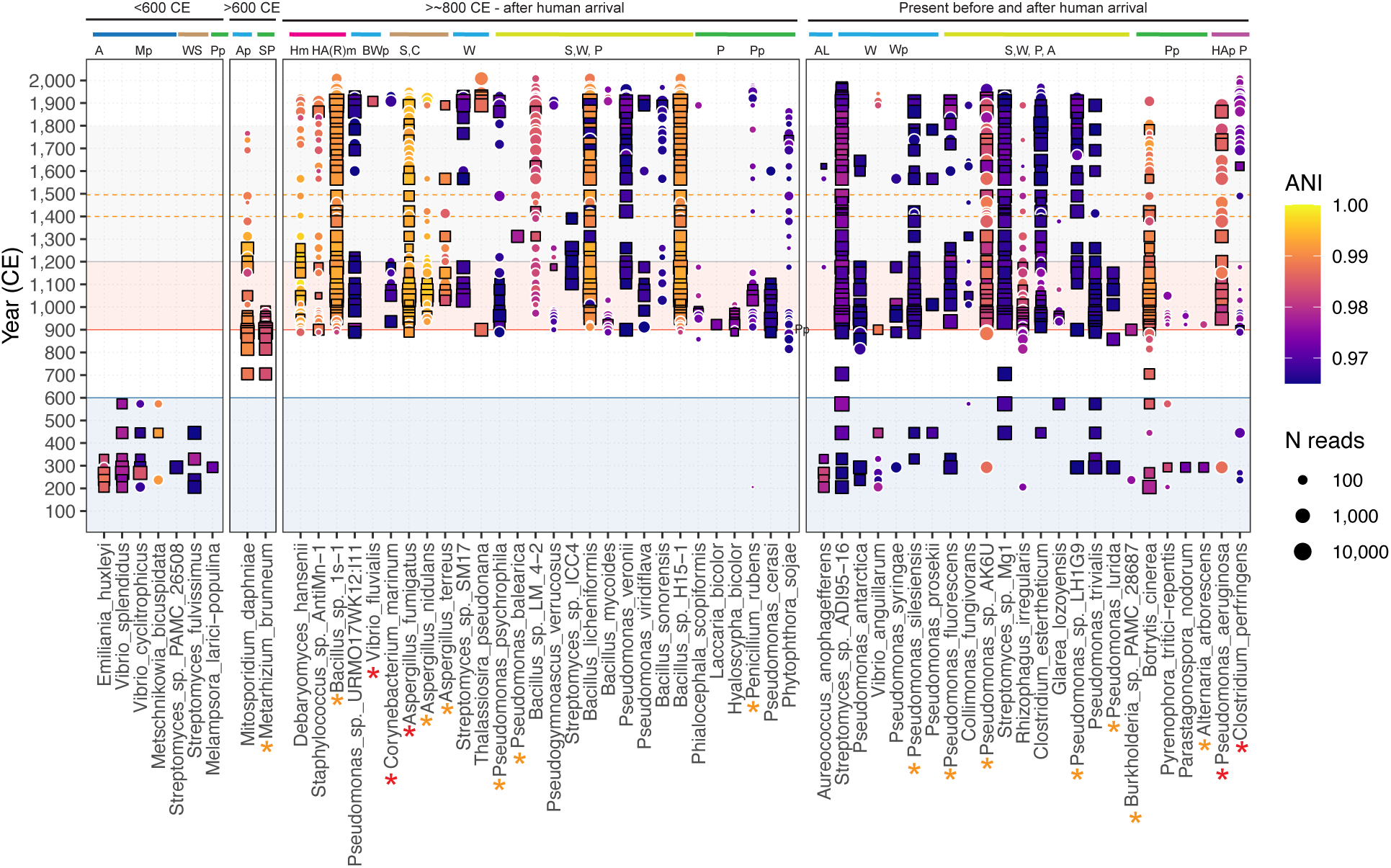
Ancient microbial species screening. The observed microbial species are categorized by the time periods in which they were first identified. Plot symbols show individual species identified in the respective samples and time periods. The number of reads aligned passing all filters is indicated by symbol size, whereas read ANI is indicated by symbol fill colour. Species hits passing all authentication criteria are indicated by black squares; samples showing species hits without significant aDNA damage but authenticated with aDNA damage in at least one other sample are indicated with white circles. Uppercase A signifies animals; M signifies marine; W signifies water; S signifies soil; P signifies plants; H signifies human; R signifies ruminants; C signifies compost; and AL signifies algae. Lowercase p signifies pathogens in some animals or plants, while lowercase m stands for microbiome. Red stars highlight microbial species that can cause disease in previously healthy individuals, whereas orange stars signify species that may cause disease in humans, but primarily in those who are immunocompromised.

**Extended Data Figure 10.**
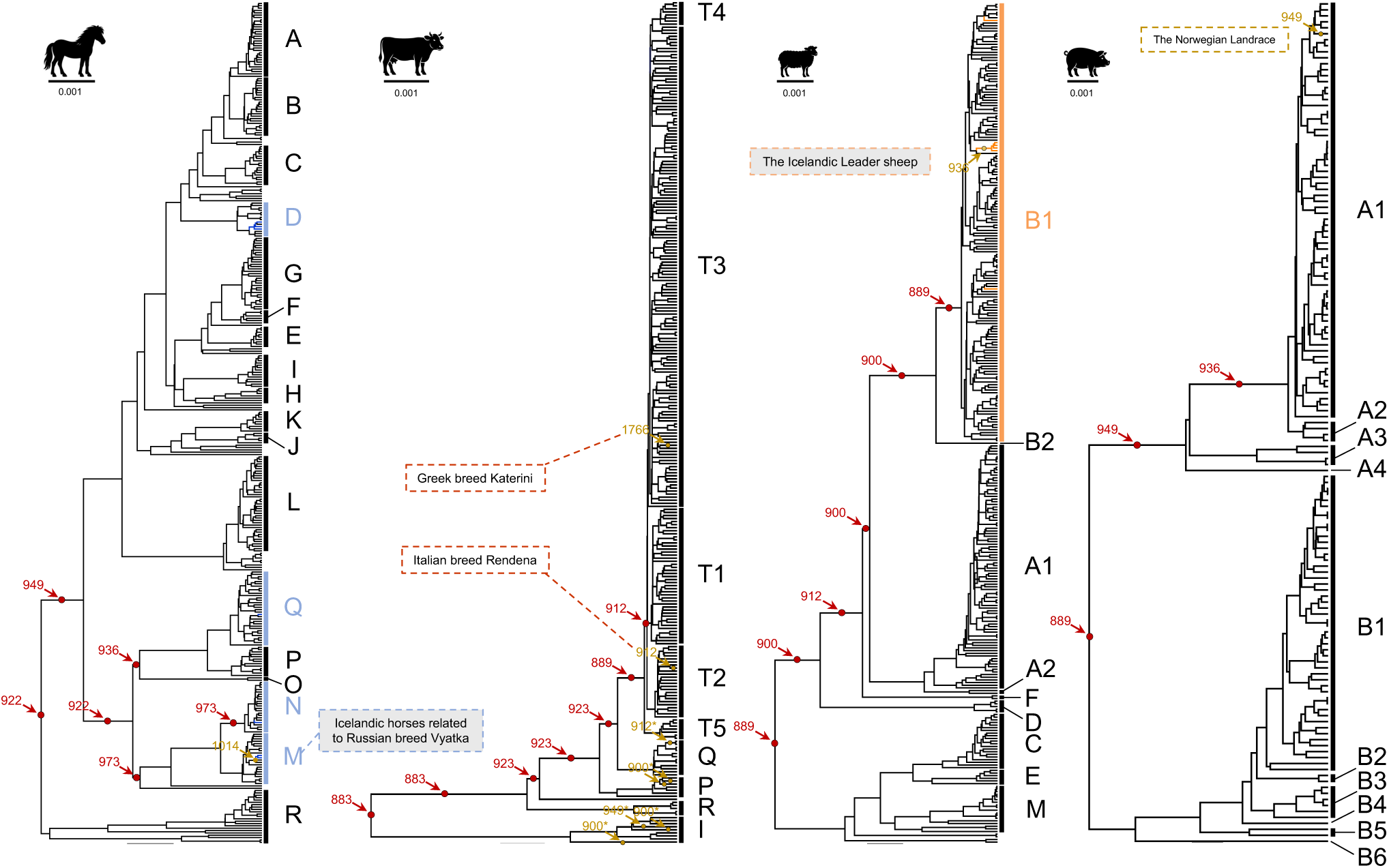
Phylogenetic placement of domesticated animals. From left to right, mitochondrial phylogenetic placements of eDNA for horse, cattle, sheep, and pig. Branch lengths are proportional to nucleotide substitutions per site. Red dots indicate backbone branches supported by eDNA, along with the earliest age of eDNA signal occurrence on the given lineage (in CE). Brown dots indicate eDNA-supported lineages closer to the tips of the tree; age marked by an asterisk (*) indicates cases where eDNA supports the tip-level branch only but not its corresponding backbone lineage. Haplogroups annotated with grey background labels indicate lineages currently present in Iceland. Coloured dashed boxes highlight haplogroups or individual samples supported by eDNA.

---

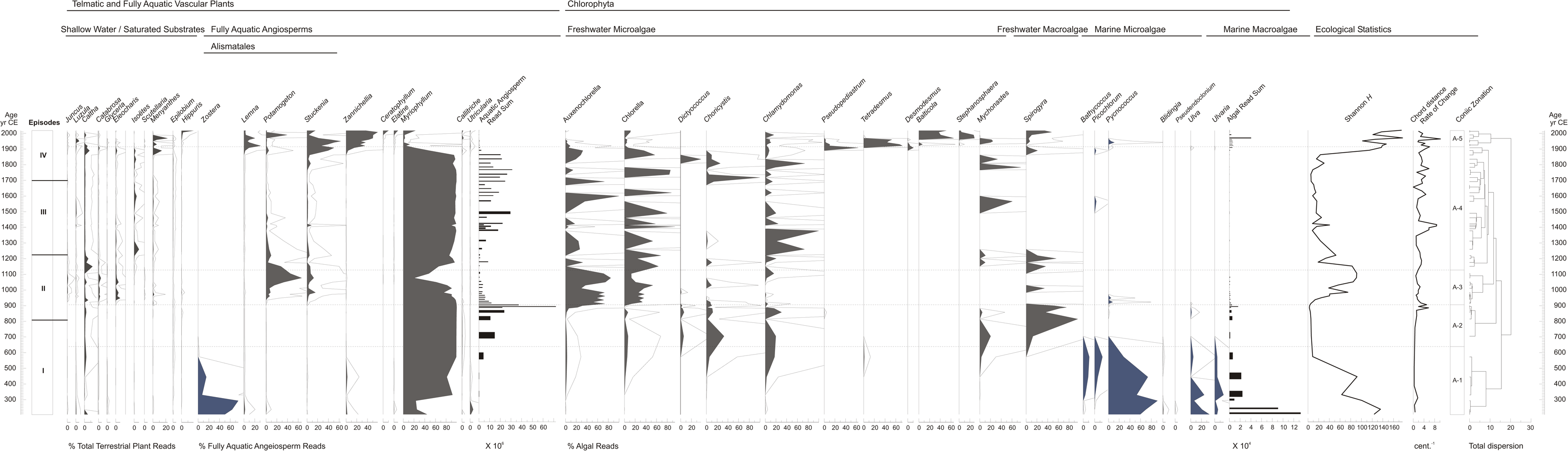

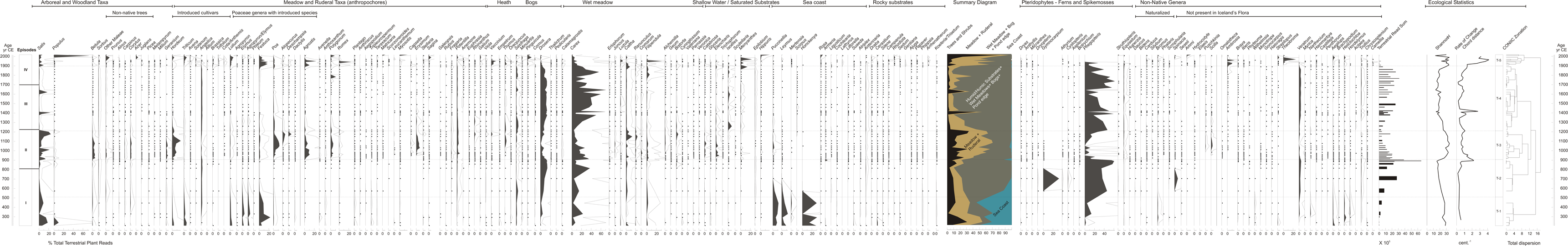

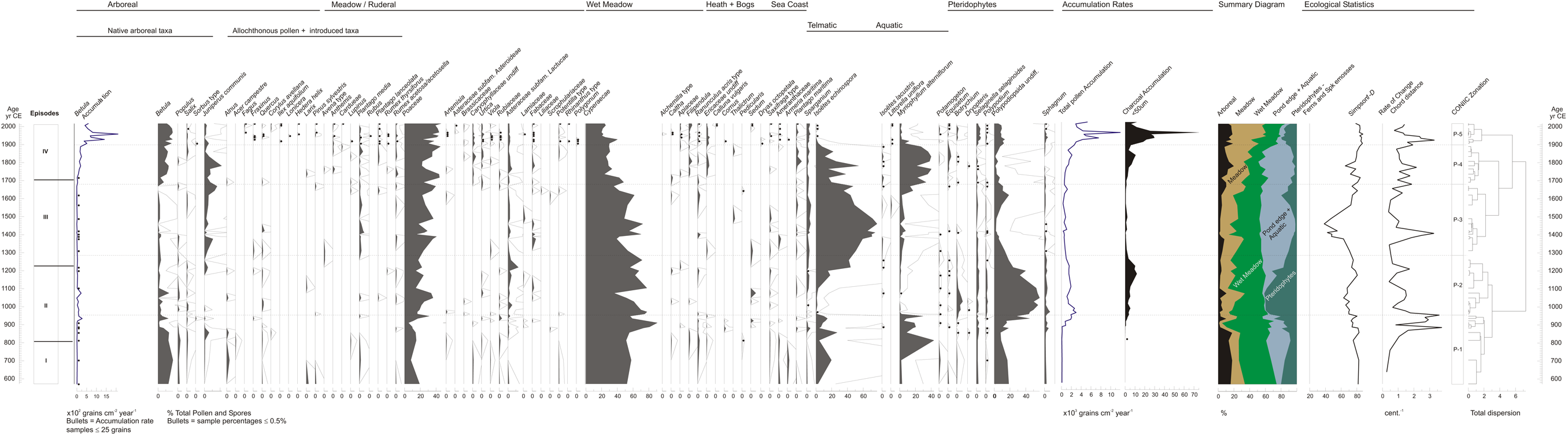

