## Supplementary information for "Environmental DNA Reveals Reykjavík’s Human and Ecological History"

### Table of Contents

*The Tjörnin Lake Record* ... 2
**Sediment description** ... 3
**Chronology** ... 5
*Geochemical and Macrofossil Proxies* ... 10
**XRF Elemental Analyses** ... 10
**Lipid Analyses** ... 13
**High-resolution molecular biomarker analysis *via* Mass Spectrometry Imaging** ... 15
**Chironomid Analysis** ... 17
Results and description of zones ... 18
Principal Component Analysis ... 19
Canonical Correspondence Analysis ... 19
*The Eukaryotic Fraction of the Metagenomes* ... 20
**Details about versions and tools used in the Holi pipeline** ... 20
Reproducibility for ancient environmental metagenomic samples ... 21
Defining DNA Damage and Model Fit Quality to filter data ... 24

Chloroplast DNA and TrnL barcodes ... 36
**Description and Interpretation of the Eukaryote Metagenomic Records ... 41**
Metagenomic record of aquatic plants ... 42
Metagenomic record of terrestrial plants ... 45
Comparison of palynological and metagenomic records ... 52
Metazoan metagenomic record ... 55
**Metazoan eDNA phylogeny ... 61**
Horse ... 61
Sheep ... 61
Cattle ... 62
Pigs ... 62
**Impact of the Landnám on *Betula* populations ... 62**
**The ancestry of Icelandic Viking Age barley ... 66**
EM Method ... 69
Comparative analysis of historical herbarium barley ... 71
DNA recovery ... 72
Basic bioinformatics ... 73
Ancestry estimation ... 73
***Prokaryotic Components of the Metagenome ... 74***
**Microbial taxonomic profiling ... 74**
Dataset filtering ... 76
Ordination-based analyses ... 76
Reference based metabolic reconstruction ... 79
**Microbial source tracking ... 81**
**The ancient microbial record ... 82**
**Viral source ecosystems ... 84**
Viral community diversity ... 84
***Historical, Archaeological, and Demographic Context ... 86***
**Estimate of the Icelandic population ... 86**
**Summary of the archaeological evidence from Reykjavik area ... 86**
The archaeology of Reykjavík ... 87
Reykjavík and Jarðabók, 1703 CE ... 92
Reykjavík, 1750–1800 CE ... 95
Reykjavík, ca. 1810 CE ... 96
Tables listing taxa reported from archaeological excavations and historical sources ... 97

### The Tjörnin Lake Record

The present-day Tjörnin is a shallow lake with maximum depth of 0.8 m and a surface area of ca. 8.94 hectares situated in Miðborg, Reykjavik's city centre (64.14496°N 21.94255°W). The lake basin is today separated from Faxaflói bay by a coastal lagoon barrier which, before landfilling and recontouring, was approximately 300 meters wide. Tjörnin drains from its northeast corner through an extended culvert beneath Lækjargata, reaching sea at Faxaflói, about 700 meters to the north (Fig. 1). Tjörnin, located south of the coastal barrier, receives freshwater and sediment from a catchment area of 2.7 km$^2$ (Fig. 1; S1)[1]. Its surface lies 3 m above sea-level. The lake's outlet, the

Lækurinn, was confined to a culvert and paved over in 1911. In the following years, a lock was installed on the canal to impede storm surges, but marine influx was not completely blocked until 1989. Historically, surges and storm tides made the lake periodically brackish and probably stratified the water column when the lake was deeper. However, influx from its extensive catchment including the Vatnsmýri marsh ensured that the lake's photic zone was fresh enough to support *Myriophyllum* for most of the sequence. The lake was larger prior to the $20^{th}$ century, when the city began systematically filling its southern margin with domestic trash. By the 1920s, ca. 30% of the lake's former area had been lost to land reclamation. Later the lake was divided by a bridge and its banks paved. The roads and buildings adjacent to the lake today are mostly built on historic fill[1].

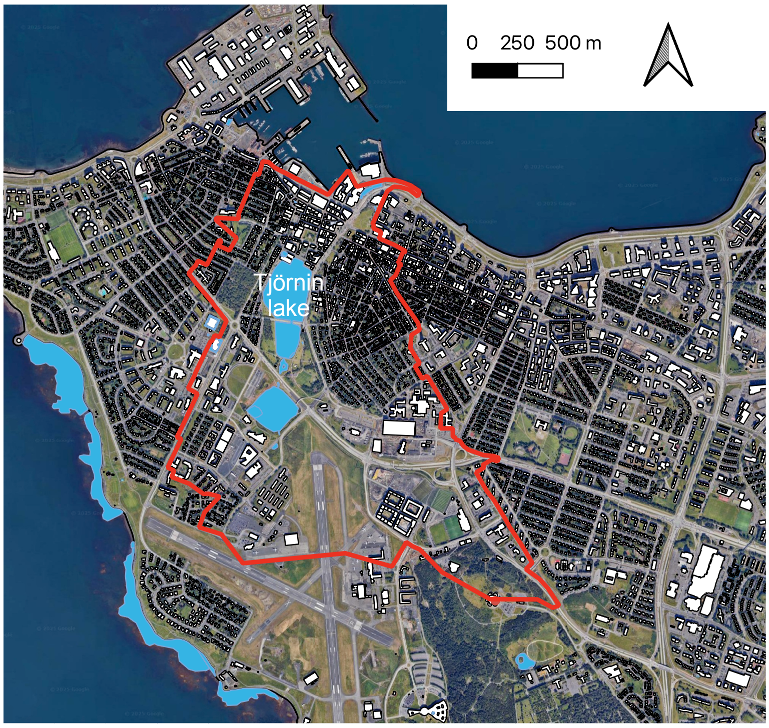

**Figure S1. Present-day outline of the Tjörnin lake catchment (red polygon) equal to 2.74 $km^2$.** Blue is lake areas and white is building infrastructure. Tjörnin is bounded to the north by the Miðbær, with urban development extending onto the surrounding hills of Skólavörðuholt to the east (39 m a.s.l.), and Landakotshæð to the west (22 m.a.s.l.). Via a culvert beneath a major road (Hringbraut, route 49), a ditch feeds into Tjörnin from the heavily modified ponds and wetlands of Vatnsmýri, a nature conservation area to its south. Vatnsmýri once covered a much greater area, now occupied by the University of Iceland campus, and predominantly by Reykjavík Airport. The airport is bordered by the high ground of Skólavörðuholt to the north, Öskjuhlið to the east (61 m.a.s.l.), the sea to the south (Skerjafjörður), while the neighbourhoods of Melar and Skerjafjörður sit upon very low hills to the west and southwest respectively.

**Sediment description**

Two stratigraphically overlapping sediment cores were collected in 2014 (T14_7) and 2022 (T22) (Fig. 1). In core T22, the uppermost 38 cm of the sequence is a gelatinous algal gyttja with fine mostly terrestrial plant detritus (mean organic matter ca. 25%). Its large minerogenic component is silty clay, with inclusions of coarse basalt sand and particles resembling bituminous coal. The ephippia (resting eggs) of *Daphnia*, chironomid head capsules and bryozoan statoblasts are present throughout the upper 195 cm of the sequence, consistent with a eutrophic, fresh to brackish lake (Fig. S2).

The segment between 38 and 164 cm is comprised of detrital gyttja like that of the overlying strata (mean organic matter ca. 25%). The detritus is comprised of indeterminate fragments of aquatic plants and terrestrial matter including oribatid mites, uncarbonized bark and wood as well as charcoal. Plant macrofossils identified included deciduous leaf fragments (cf. *Betula*), and seeds of *Betula* cf. *pubescens*, *Cerastium* sp., *Potomogeton* sp. and *Juncus* sp., along with the oospores of *Nitella* and *Isoëtes echinospora* megaspores. The recovery of the *Nitella* oospores suggests that some of the indeterminate aquatic detritus could be derived from these macroalgae. From a depth of 164 to 234 cm, the sequence is comprised primarily of algal gyttja (organic matter mean ca. 30%). Oospores of *Nitella*, stem fragments were present in between 222 and 234 cm.

The base of the sequence (ca. 234 – 250 cm) is comprised of detritus gyttja (mean organic matter ca. 46%). *Betula* leaf fragments, indeterminate bark and leaf fragments and seed of a Caryophyllid (cf. *Cerastium* sp.). Roots in the layer may be derived from *Zostera marina*, which could have colonized the detrital surface. Coleoptera fragments and Diptera larvae were also noted. Gastropods were the only molluscs and charcoal was not present. Fig. S2 shows the sediment lithology against depth in centimetres below the lake's mud-water interface.

The variable interval of the modelled age scale, in the left margin attests to both slower deposition and greater compression at the base of the sequence. The deposition rate increased after 900 CE, slowed beginning in the 13$^{th}$ century and increased again after 1600 CE. Very fine gyttja with aquatic detritus began to accumulate after ca. 600 CE and the minerogenic component increased to comprise ca. 50 to 80 % of the sediment by weight. By 1100 CE coarser detritus gyttja had begun to accumulate again, probably derived from decomposed *Myriophyllum*. A final depositional shift, to a fine algal gyttja occurred at about 1900 CE and continues to the present.

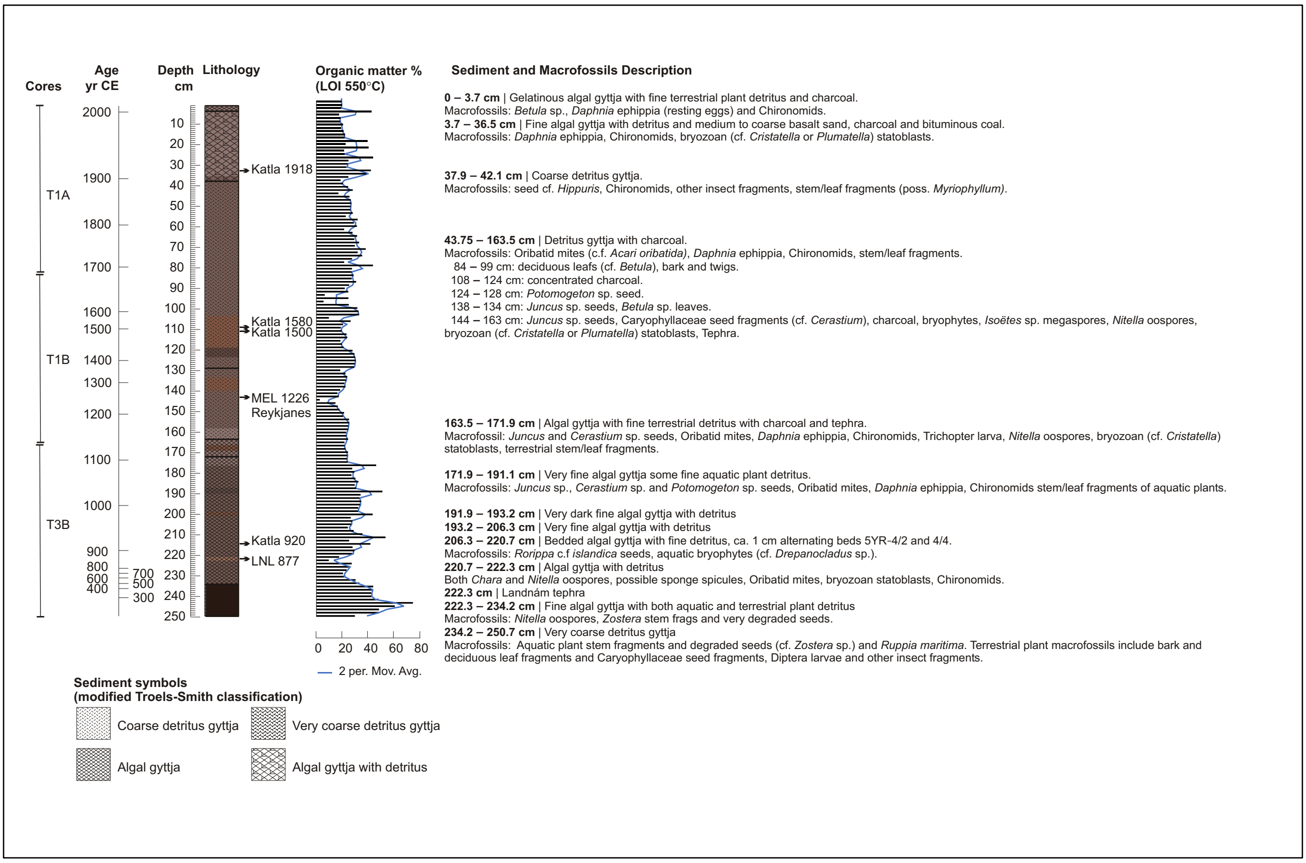

***Figure S2.* The Tjörnin record (Composite Core T22).** Sediment lithology and the proportion of organic matter determined by loss on ignition are shown against depth in centimetres below the modern lake's mud-water interface. The positions of identified tephra layers are indicated on the lithology column. Sediment symbols are modified from Troels-Smith's classification, and the colours match the Munsell values of the sediment units. The sub-sampled cores are indicated in the left margin next to the modelled age scale.

#### Chronology

A chronological framework for the Tjörnin stratigraphy was established using historical volcanic ash layers (tephra), radiocarbon dating of macrofossils ($^{14}C$), and plutonium isotope analysis (Pu) (Figs. S2-5; Tables S1-S3). This approached was done independently for T22 and T14. The age-depth model for the T22 composite core spans from 200 CE to the present and T14 from 600 CE the present (Extended Data Fig. 1, Fig. S3). To enhance chronological control, a stratigraphic alignment was performed based on diagnostic features shared between the two records. In total, 31 tie points were identified along the sequences, indicating that core T14_7 is characterized by reduced temporal resolution and potential short hiatuses, particularly around ca. 900 CE and 1750–1900 CE. The figure illustrates optical and X-radiographic line-scan images of the two cores (T22 to the left, T14_7 to the right), each displayed on its independent age model. Tie lines denote depth–depth correlations between corresponding stratigraphic horizons (Fig. S3).

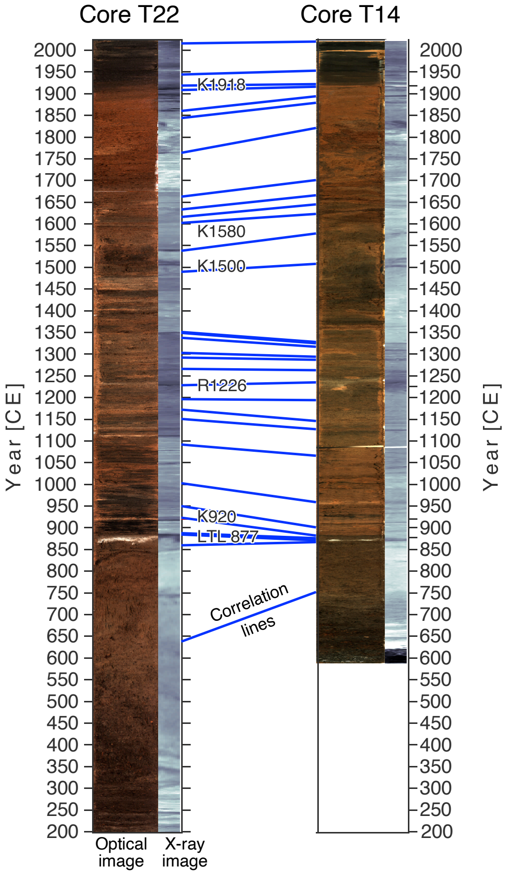

**Figure S3. Correlation between T22 and T14_7.**

The Tjörnin lake record contains key regional marker layers that one would expect to find within the stratigraphic record including K1918, K1500, R1226, K920 and LTL877. These tephra marker layers are all historic with known age and previous descriptions from West Iceland[3–5]. An additional tephra layer was also identified, sampled and analysed roughly 3 cm above K1500 in the T22 record (c. 22 cm in the T1B core – 1115 mm composite depth). Although this tephra layer had trace visibility (compared to K1500) it was also visible in the radiograph and scan data (Fig. 2). Probe results (Tjörnin Tephra Source File) prove 18 points consistent with Katla basaltic glass like that of the K1500 tephra[3,4].

The high concentration (with trace visibility) homogenous tephra from Katla several centimetres above K1500 could be the product of reworking. However, the sharp peak in MS, the clear horizon in the radiograph and a similar couplet visible in the T14 core suggests the tephra horizon is the result of a separate eruption from Katla which deposited relatively less tephra in Tjörnin than the K1500 eruption. The next documented eruption from Katla occurred 1580 (K1580). The K1580 tephra layer is only detected in localities due south of the volcanic province and is unknown to have been deposited across other regions of Iceland. While it seems unlikely to find the K1580 in Tjörnin given this dispersal pattern, it may not be impossible given the occasional discordance of local isopach orientation verse distal deposition highlighted by Kalliokoski and colleagues[6]. The allocation of this tephra to K1580 seems a simpler solution in comparison to creating a new historic Katla eruption.

While tephra K1721 was deposited westward[7] and presumably may have been deposited within the lake catchment, this tephra layer was not identified, nor have other studies presented it[4].

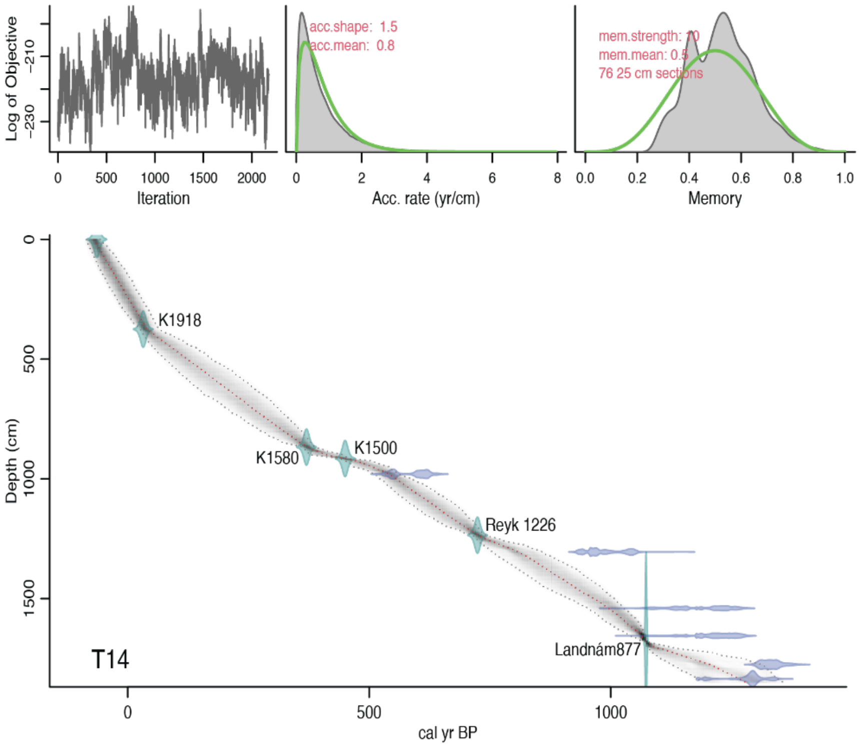

**Figure S4. Age depth model for T14-7 composite stratigraphy.** Tephra and radiocarbon age details are given in Table S1 and S2.

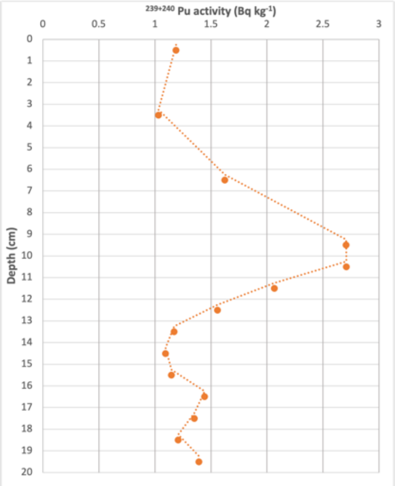

**Figure S5. Plot of Pu239+240 concentrations in the upper most 20 cm of the Tjörnin T22-1A core.** Peak fall out (10 ± 1 cm) is associated with nuclear weapons testing from 1963-1964.

**Table S1. Radiocarbon ages of macrofossils from Tjörnin.** Modelled ages derived from Bacon age depth model using Intcal20 (Reimer et al. 2020). Median calibrated ages in **bold.** * Unreliable result – excluded from age depth model

| **Lab ID** | **Sample ID** | **Sample material** | **Core** | **Depth (cm)** | **Comp. Depth (mm)** | **Mass (mg)** | δ13C‰ V-PDB | **$^{14}$C age (yr BP)** | **Error** | **Modeled median (cal. yr BP)** | **Modeled range (cal. yr BP)** |
|---|---|---|---|---|---|---|---|---|---|---|---|
| Ua-75012 | T1 | cf. *Eriophorum* sp. | ISL22-T1A | 69-70 | 343 | 0.83 | -19.5 | 352* | 27 | * | * |
| Ua-75013 | T2 | cf. *Andromeda* sp.; *Vaccinium* sp. | ISL22-T1B | 39-40 | 1234 | 0.85 | -19.3 | 522 | 28 | **538** | 513-602 |
| Ua-75014 | T3 | cf. *Eriophorum* sp. *Andromeda* sp. aquatic bryophyte | ISL22-T3B | 59.5-60 | 2103 | 0.61 | -22.1 | 1505* | 29 | * | * |
| Ua-75015 | T4 | cf. *Andromeda* sp.; *Vaccinium* sp.; *Betula* | ISL22-T3B | 85-86 | 2380 | 1.21 | -21.5 | 1877 | 29 | **1641** | 1552-1739 |
| Ua-75016 | T5 | cf. *Andromeda* sp.; *Vaccinium* sp.; *Betula* | ISL22-T3B | 93-94 | 2460 | 1.44 | -20.7 | 1755 | 29 | **1708** | 1626-1805 |
| Ua-79107 | ISL-C58 | Frag. terrestrial detritus; poss. *Salix* & *Empetrum* leaf & bark | ISL22-T3B | 78-79 | 2313 | 0.53 | -18.5 | 1863* | 29 | * | * |
| Ua-77313 | ISL-C21 | Plant frag. not IDed | ISL22-T3B | 85-86 | 2380 | 0.84 | -17.6 | 1855 | 30 | **1636** | 1546-1735 |
| Ua-77314 | ISL-C22 | Stem frags. (poss. *Nitella*) | ISL22-T3B | 93-94 | 2460 | 1.87 | -19.3 | 1846 | 29 | **1714** | 1631-1810 |
| AAR-22625 | GeoG_6 | *Poaceae* sp. | T14-7 | 71-74 | 725 | 0.7 | -22.06 | 411 | 23 | **270** | 15-323 |
| AAR-22626 | GeoG_7 | *Poaceae* sp., *Eleocharis* sp. | T14-7 | 77-80 | 775 | 0.6 | -20.56 | 391 | 24 | **307** | 255-348 |
| AAR-22627 | GeoG_8 | *Poaceae* sp. | T14-7 | 97.5-98.5 | 980 | 0.4 | -10.9* | 566 | 25 | **541** | 510-562 |
| AAR-22628 | GeoG_9 | *Poaceae* sp., *Eleocharis* sp. | T14-7 | 130-131 | 1305 | 0.6 | -17.41 | 1078 | 22 | **807** | 755-858 |
| AAR-22629 | GeoG_10 | *Poaceae* sp., *Eleocharis* sp., char | T14-7 | 154 | 1540 | 0.5 | -16.24 | 1248 | 27 | **998** | 954-1027 |
| AAR-22630 | GeoG_11 | *Poaceae* sp. | T14-7 | 165-166 | 1655 | 1.1 | -17.96 | 1269 | 25 | **1065** | 1042-1072 |
| AAR-22631 | GeoG_12 | *Betula nana* | T14-7 | 177-178 | 1775 | 1.1 | -27.43 | 1441 | 25 | **1195** | 1125-1314 |
| AAR-22632 | GeoG_13 | *Betula nana* | T14-7 | 183-184 | 1835 | 2.6 | -23.55 | 1369 | 26 | **1280** | 1180-1346 |

**Table S2. Tephrochronological constraints from Tjörnin.** All electron microprobe analysis conducted at University of Iceland. *Unsampled / unanalysed horizon correlated from ISL22-T1B (XRF) scan data and probe analysis.

| **Tephra** | **Probe points** | **Province** | **Core** | **Sample Depth (mm)** | **Comp. Depth (mm)** | **Age (BP)** | **Name** | **Reference** |
|---|---|---|---|---|---|---|---|---|
| Katla 1918 | 12 | Katla | ISL22-T1A | 350-360 | 335 | 32 | **K1918** | Thorarinsson 1958; Oladóttir et al. 2008, 2011 |
| Katla 1580 | 18 | Katla | ISL22-T1B | 215-225 | 1059 | 370 | **K1580** | Larsen 1993; Haflidason et al. 2000 |
| Katla 1500 | 14 | Katla | ISL22-T1B | 245-255 | 1089 | 450 | **K1500** | Haflidason et al. 1992; Eiríksson et al. 2000; Larsen 2010 |
| Reykanes 1226 | 18 | Reykanes | ISL22-T1B | 615-625 | 1445 | 724 | **MEL1226** | Jóhannesson & Eirnarsson 1988 |
| Katla 920 | 18 | Katla | ISL22-T3B | 615-625 | 2128 | 1030 | **K920** | Haflidason et al. 1992; |
| Bardabunga-Veidivötn | 14 | Bardabunga-Veidivötn | ISL22-T3B | 715-725 | 2228 | 1073 | **Landnám** | Grönvold et al. 1995; Larsen et al. 1999; Schmid et al. 2017 |

| | | | | | | | | |
|---|---|---|---|---|---|---|---|---|
| Torfajökull | 24 | Torfajökull | ISL22-T3B | 715-725 | 2228 | 1073 | **Landnám** | Grönvold et al. 1995; Larsen et al. 1999; Schmid et al. 2017 |
| Katla 1918 | 12 | Katla | T14-7 | 370-380 | 375 | 32 | **K1918** | Thorarinsson 1958; Oladóttir et al. 2008, 2011 |
| Katla 1580 | * | Katla | T14-7 | * | | 370 | **K1580** | XRF tie point |
| Katla 1500 | 10 | Katla | T14-7 | 910-920 | 915 | 450 | **K1500** | Haflidason et al. 1992; Eiríksson et al. 2000; Larsen 2010 |
| Reykanes 1226 | 15 | Reykanes | T14-7 | 1230-1240 | 1235 | 724 | **MEL1226** | Jóhannesson & Eirnarsson 1988 |
| Bardabunga-Veidivötn | 2 | Bardabunga-Veidivötn | T14-7 | 1675-1685 | 1680 | 1073 | **Landnám** | Grönvold et al. 1995; Larsen et al. 1999; Schmid et al. 2017 |
| Torfajökull | 11 | Torfajökull | T14-7 | 1675-1685 | 1680 | 1073 | **Landnám** | Grönvold et al. 1995; Larsen et al. 1999; Schmid et al. 2017 |

#### Table S3. Plutonium data for the core T22 lake core from Tjörnin, Iceland.

| ISL ID | Lake ID | Top (cm) | Bottom (cm) | NAU Lab ID | # meas | Bqkg 239+240 | Bq/kg sd | $^{240}Pu/^{239}Pu$ | 240239 sd |
|---|---|---|---|---|---|---|---|---|---|
| ISL-P43 | ISL22-T1A | 0 | 1 | 43 | 3 | **1.189** | 0.051 | 0.188 | 0.023 |
| ISL-P44 | ISL22-T1A | 3 | 4 | 44 | 3 | **1.031** | 0.079 | 0.181 | 0.026 |
| ISL-P45 | ISL22-T1A | 6 | 7 | 45 | 3 | **1.624** | 0.146 | 0.168 | 0.024 |
| ISL-P46 | ISL22-T1A | 9 | 10 | 46 | 2 | **2.706** | 0.025 | 0.186 | 0.016 |
| ISL-P47 | ISL22-T1A | 10 | 11 | 47 | 3 | **2.708** | 0.154 | 0.178 | 0.006 |
| ISL-P48 | ISL22-T1A | 11 | 12 | 48 | 3 | **2.066** | 0.09 | 0.182 | 0.009 |
| ISL-P49 | ISL22-T1A | 12 | 13 | 49 | 3 | **1.559** | 0.098 | 0.182 | 0.022 |
| ISL-P50 | ISL22-T1A | 13 | 14 | 50 | 3 | **1.171** | 0.094 | 0.158 | 0.013 |
| ISL-P51 | ISL22-T1A | 14 | 15 | 51 | 3 | **1.095** | 0.08 | 0.213 | 0.005 |
| ISL-P52 | ISL22-T1A | 15 | 16 | 52 | 3 | **1.147** | 0.066 | 0.183 | 0.024 |
| ISL-P53 | ISL22-T1A | 16 | 17 | 53 | 3 | **1.442** | 0.076 | 0.171 | 0.029 |
| ISL-P54 | ISL22-T1A | 17 | 18 | 54 | 3 | **1.352** | 0.1 | 0.189 | 0.035 |
| ISL-P55 | ISL22-T1A | 18 | 19 | 55 | 3 | **1.207** | 0.064 | 0.15 | 0.008 |
| ISL-P56 | ISL22-T1A | 19 | 20 | 56 | 3 | **1.394** | 0.077 | 0.188 | 0.023 |

The accuracy of the age-depth model for the uppermost part of the Tjörnin core is attested by a major fire detected as an isolated peak in naphthalene lipid concentrations (Fig. 2; Figs. S9-15). The age-depth model dates the peak to the late 1960s CE which aligns closely to a large fire in 1967 that destroyed or severely damaged three houses including a large bank (Iðnaðarbankinn)[8]. As an additional anchor on the age-depth model, there is a historically documented massive storm surge that brought saltwater into the basin in 1345 CE, which corresponds with an isolated, prominent bromine spike (Fig. 2) indicative of saltwater flooding the basin that by the model dates to 1350 CE. The sediment accumulation rate in the Tjörnin basin before the arrival of humans was relatively slow, ~0.40 mm/yr. After human arrival, the rate increased to ~1.96 mm/yr[5]. Also, the distinct variations in detrital input, microfossil content, and elemental composition were observed throughout the core. These patterns, together with the preservation of intact, unbroken tephra layers, indicate that bioturbation was primarily confined to the mud–water interface, and therefore, is estimated to impact only the ~10 cm within the mud–water interface. Thus, we infer that the Tjörnin record yields a decadal temporal resolution.

It should also be noted that although the bottom of the basin today aligns with present-day sea level (Figure 1), regional sea-level studies in western Iceland indicate that ca. 100 CE the sea level was approximately 1.3 m lower. This was previously thought to imply that the lowermost deposits in the basin, which date to 200 CE, could have a terrestrial origin. That inference is consistent with the sedimentary sequence of Tjörnin which consists of organic gyttja overlaying layers of gravel, sand, and volcanic tephra[9]. In this scenario, the terrestrial material was eroded or buried during the subsequent formation of the lagoon.

#### Geochemical and Macrofossil Proxies

**XRF Elemental Analyses**

X-Ray-Fluorescence (XRF) measurements was performed using an Itrax μ-XRF core-scanner from COX Analytical Systems obtaining high-resolution optical and radiography line-scan images, and variations in the element composition and magnetic susceptibility (see method section for details). Changes in chemical contents in Iceland lake sediment is largely influenced by depositions of debris from volcanic eruptions. These short-term changes appear as spikes in the chemical record and overly changes on a longer timescale that reflect environmental changes in and around the lakes. While the volcanic signals are good for core correlations and age markers, they obstruct data analyses that have the purpose of investigating environmental changes, therefore, to analyse environmental changes using XRF data, the volcanic spikes must be filtered away. We filtered the XRF data stepwise in an empirical way. First the data was median filtered using a broad window of 15 cm whereby not only volcanic spikes were effectively filtered away, but also short-term changes in the environmental signal. To compensate for this, the remanence (difference between raw data and median filtered data) was then divided into a positive and a negative contribution, where the positive values will mainly consist of volcanic spikes, and the negative values will mainly contain short term variations in the environmental signal. In case of positive values in the remanence, the negative series value was set to zero and vice versa. The negative series were then smoothed using running means with a window of 3 cm and added to the median filtered data. The XRF raw data including volcanic spikes is shown together with the filtered data in the Figure S6.

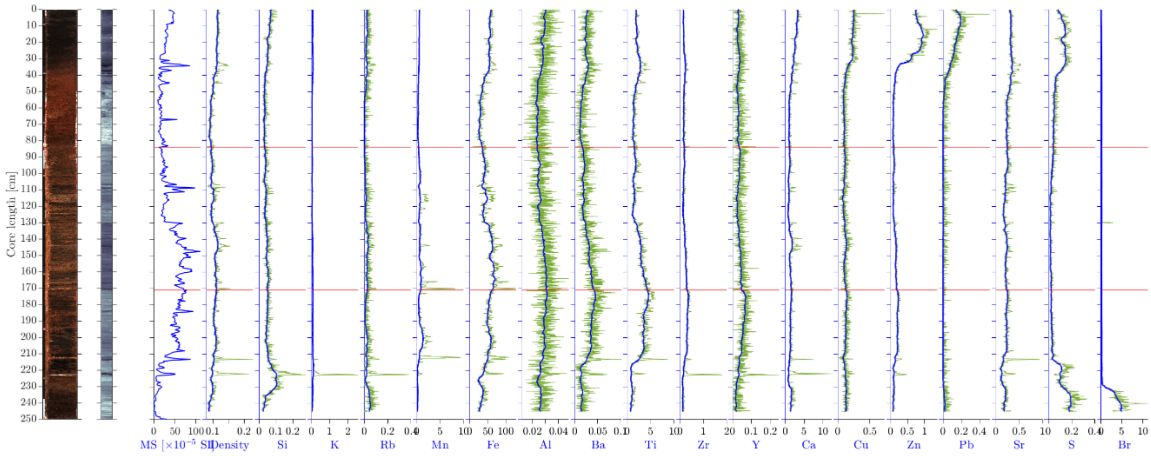

**Figure S6. Itrax data.** From the left: Optical image, radiography image, magnetic susceptibility, density proxy (obtained from the Coherent/Incoherent Rhodium scatter peak ratio), and elements (in peak area normalized to the coherent Rhodium scatter peak). Density proxy and elements were filtered. The unfiltered data are shown in green thin lines while the filtered data are shown in thick blue.

Figure S7 shows correlation maps for the filtered data. The map to the left show correlations of the whole data series, while the map to the right show correlations from when the basin was a freshwater lake at 660 C.E. (229 cm depth). The correlations suggest three major contributions to elements in the lake sediments: Br indicates marine influx to the lake, while Ca, Cu, Zn, Pb, Sr, and S indicate wastewater inflow and air-pollution depositions in the lake, and the remaining elements indicate minerogenic material from soil erosion.

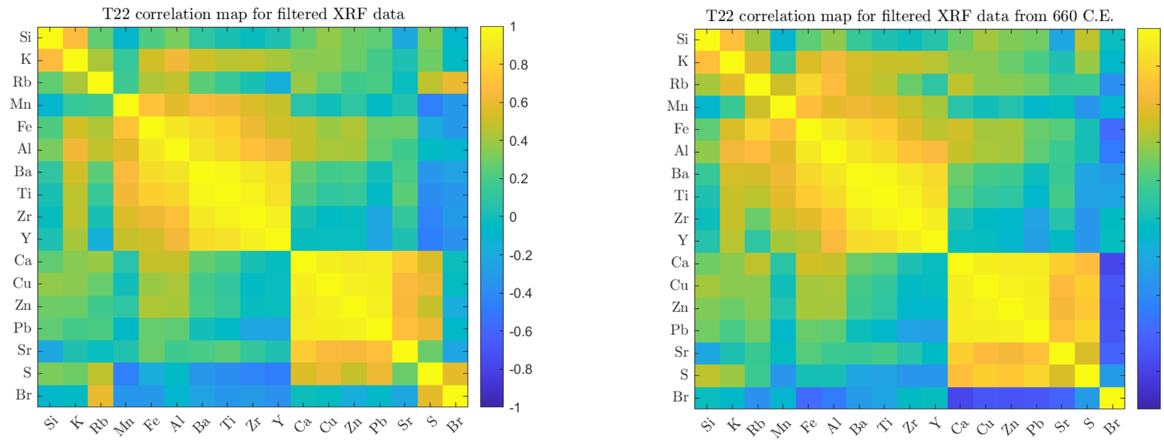

**Figure S7.** ***Correlations between filtered element series.*** *Left, the whole series. Right, from 660 C.E (229 cm) and up.*

To obtain robust proxies for minerogenic influx, as well as depositions from wastewater/air-pollution, we use based on the correlation map Al, Fe, Ba, Ti, Zr, and Y combined as an indicator for soil erosion, and Ca, Cu, Zn, Pb, Sr, and S combined as an indicator of wastewater / air pollution. We use the means of z-scores of filtered data where the z-scores are evaluated based on mean and variance from 660 C.E. and up (Fig. S8).

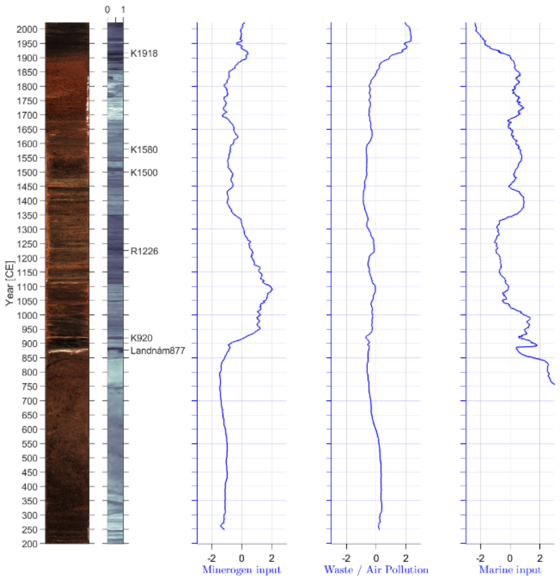

**Figure S8. Proxy for minerogenic input, waste/air pollution and marine input to lake Tjörnin**

#### Lipid Analyses

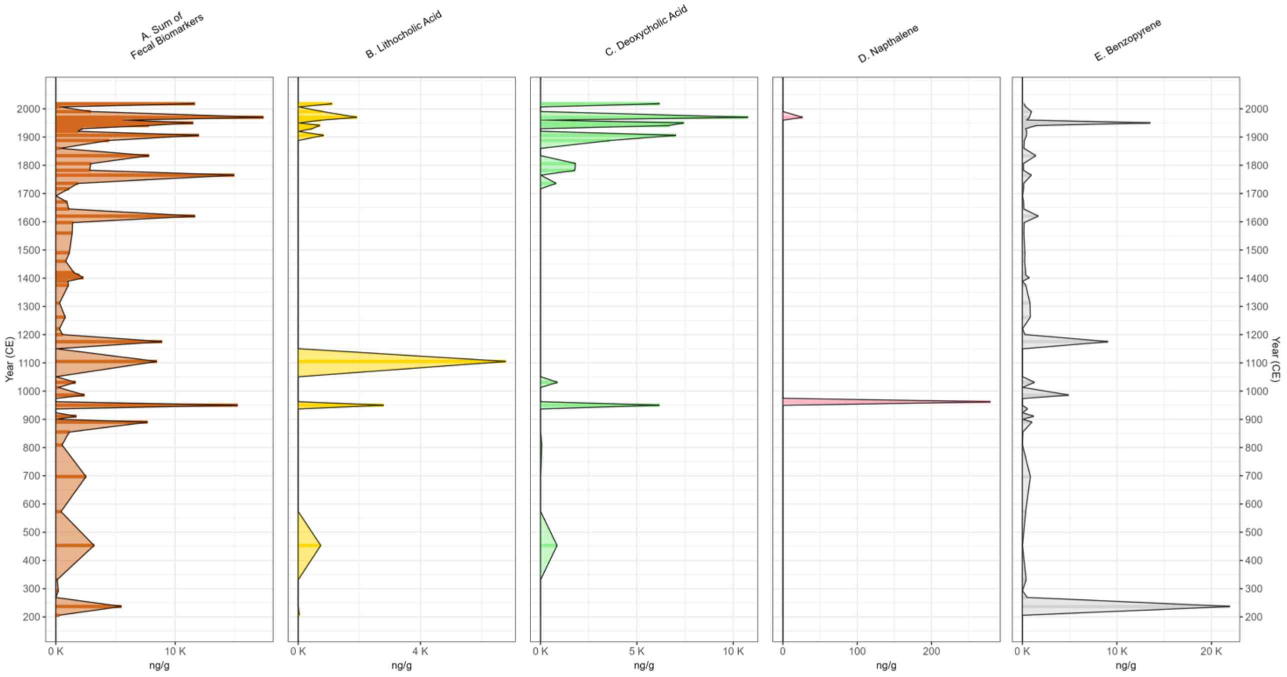

**Figure S9. Downcore profiles showing the ng/g concentration**. A) the sum of faecal biomarkers (Coprostanol, 5β--stigmastanol, lithocolic acid and deoxycholic acid) reflecting herbivore and human faecal input, B,C) Lithocolic acid and Deoxycholic acid ( bile acids associated with human faecal waste[10], D,E) Naphtalene and Benzopyrene, both polycyclic aromatic hydrocarbons produced during the incomplete combustion of organic matter, often used as proxies for vegetation burning[11].

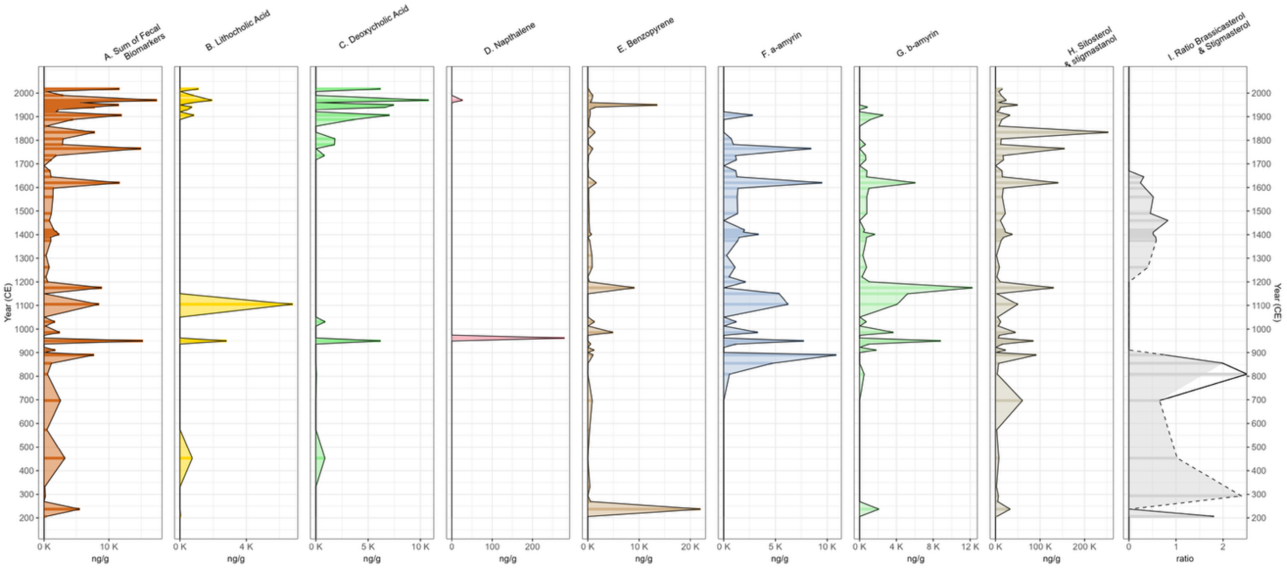

**Figure S10. Downcore profile from Tjörnin displaying several geochemical proxies.** A) the sum of faecal biomarkers (coprostanol, 5-β--stigmastanol, lithocolic acid and deoxycholic acid) reflecting herbivore and human faecal input, B,C) lithocolic acid and deoxycholic acid ( bile acids associated with human faecal waste[10], D,E) naphtalene and benzopyrene, both polycyclic aromatic hydrocarbons produced during the incomplete combustion of organic matter, often used as proxies for wood burning[11], F,G) α -amyrin and β -amyrin, plant triterpenoids signalling changes in the plant terrestrial input, H) sum of sitosterol and stigmastanol, plant sterols also showing changes in terrestrial vegetation[12], and I) ratio of brassicasterol/stigmasterol (higher values indicate an increase in lipids produced by marine diatoms in contrast to lipids produced by terrestrial plants)[13].

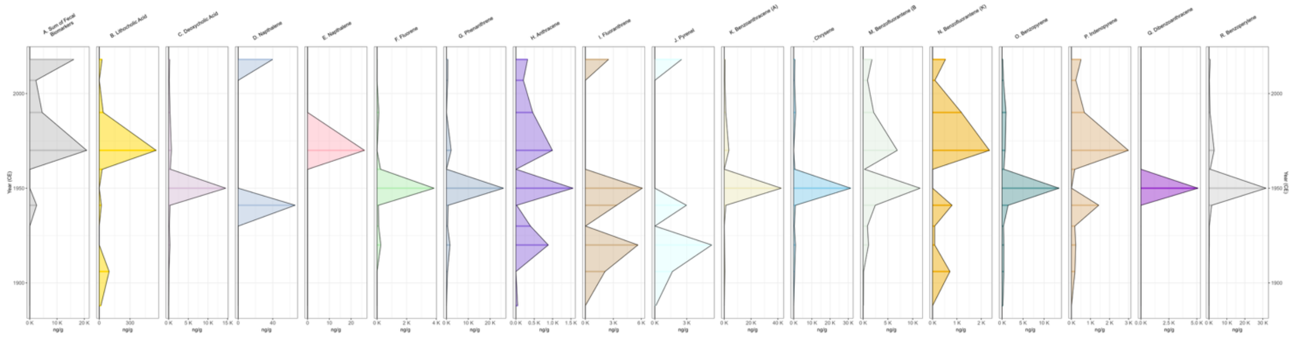

**Figure S11. Top of the profile (1888-2018) reflecting concentrations (ng/g)**. A) sum of lipids indicating the presence of mature organic matter often associated with the presence of oil (28-nor-17¨-a-hopane, bis-nor-hopane, homohopane, A'-Neogammacer-22(29)-ene)[12] and B-R) and most common polycyclic aromatic hydrocarbons[11].

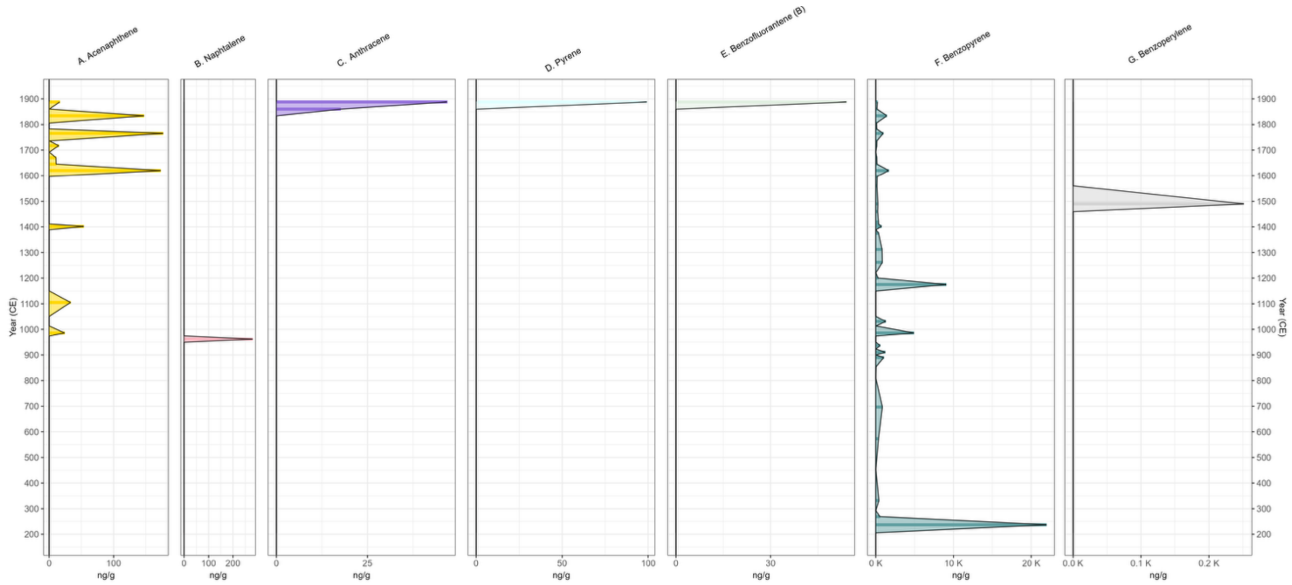

**Figure S12. Concentrations (ng/g) of most common polycyclic aromatic hydrocarbons[11] in the downcore profile from 206 to 1888 CE.**

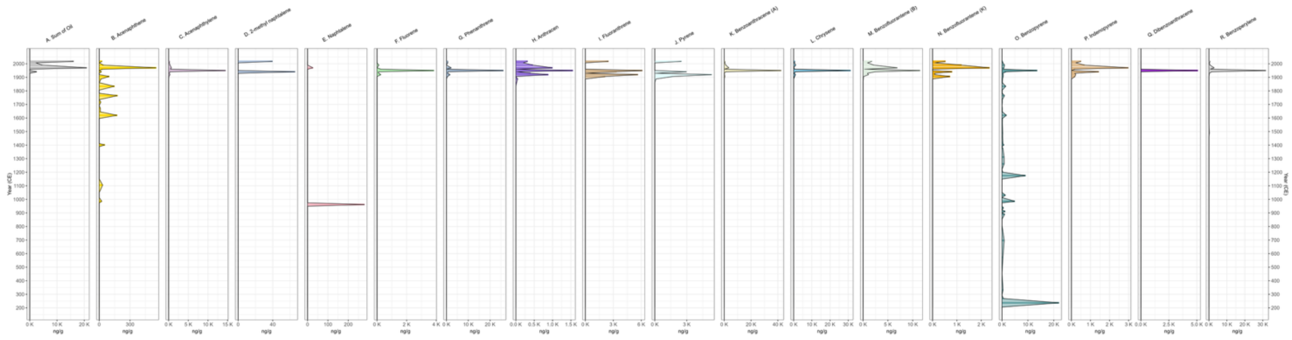

**Figure S 13. Downcore profile reflecting concentrations (ng/g).** A) sum of some lipids indicating the presence of mature organic matter often associated with the presence of oil and combustion (28-nor-17-a-hopane, bis-nor-hopane, homohopane, A'-Neogammacer-22(29)-ene)[12] and B-R) and most common polycyclic aromatic hydrocarbons[11].

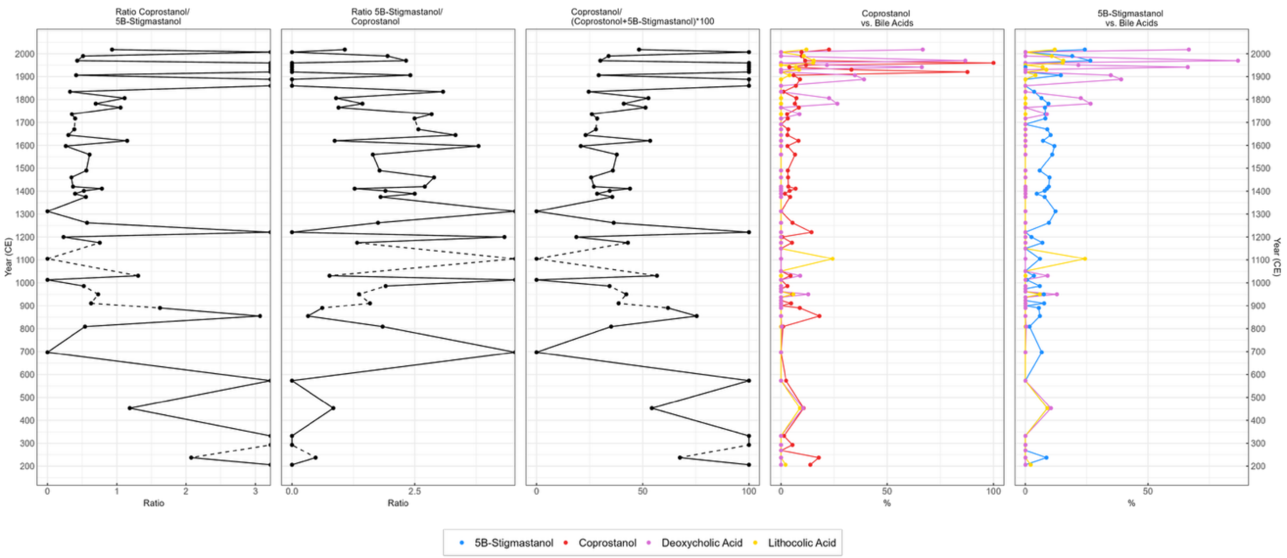

**Figure S14. Downcore profile showing ~~the~~ concentrations (ng/g)**. A) Coprostanol and bile acids, B) 5-β -stigmastanol and bile acids, C) ratio coprostanol/5-β -stigmastanol (higher values suggest human faecal input), D) ratio of 5 β -stigmastanol/coprostanol (higher values suggest hervibore faecal input), and E) ratio of coprostanol/(coprostanol+5-β -stigmastanol)*100 (values higher than 60 indicative of human faecal input)[10].

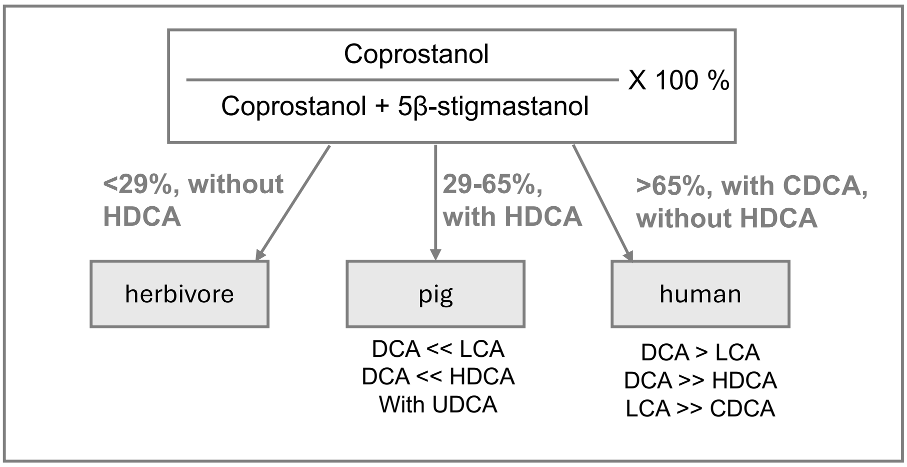

**Figure S15. Criteria for identifying faeces based on their steroid signatures.** Key distinguishing parameters are highlighted in bold. Differentiation is made between faeces from herbivores, pigs, and humans. CDCA = chenodeoxycholic acid, DCA = deoxycholic acid, HDCA = hyodeoxycholic acid, LCA = lithocholic acid, UDCA = ursodeoxycholic acid. Modified from Prost and colleagues[10].

**High-resolution molecular biomarker analysis *via* Mass Spectrometry Imaging**

A high-resolution record of molecular biomarkers indicative of fire activity (levoglucosan) and terrestrial vegetation (long chain *n*-alkanoic acids) were obtained for the period of human arrival and settlement. Therefore, a ~16-cm interval (~214-230 cm, ~615-917 CE) including the LTL was subsampled and divided into four sections, which were freeze-dried and embedded following the procedure described by Alfken and coworkers[14]. Thin slices (100µm) from these samples were obtained using a Medite Cryostat M630 and placed on indium-tin-oxide-coated glass slides. These slices were used for elemental mapping on a Bruker M4 Tornado 300 and for mass spectrometry imaging (MSI) of molecular biomarkers on a 7T Bruker solariXR mass spectrometer. MSI was performed with 200 µm raster resolution and around 40 spots per horizon, resulting in ~ 2000 spots per cm depth. To improve sensitivity, spectra were recorded in continuous accumulation of selected ions mode, with a *m*/*z* range of 150-250 for levoglucosan and 350-540 for the *n*-alkanoic acids. X-y-coordinates, intensity and signal-to-noise ratio of the target molecules were exported for each spot *via* the DataAnalysis software (Bruker Daltonics). Using MSIAlign, these abundance maps were aligned with an image of the core in order to place them in the correct depth scale. Subsequently the same software was used to generate downcore profiles in which individual spots were binned into 200 µm horizons.

Levoglucosan and its isomers (mannosan, galactosan) are major biomarkers in smoke originated during the burning of cellulose, hemicellulose, and lignin[15]. In the sedimentary record they can be used as a tracer of relatively low temperature and smouldering regional fires, including fire associated with human presence[16–18]. In the present record (Extended Data Figure 7) an initial increase in abundance is observed immediately below the LTL, while a much more pronounced maximum is recorded between ~890 and 900 CE

Because natural fire does occur in Iceland[19], we cannot be certain that the observed trend is entirely anthropogenic. However, the sudden increase post-LTL is concurrent with Norse settlement in the Tjörnien catchment. We suggest that its source is primarily domestic fires in houses and forges, possible combined with limited prescribed burning to rejuvenate grazed meadows. However, there is no historic evidence for the latter at any time in Iceland's history. Alternatively, increased erosion due to grazing and cultivation may have mobilized levoglucosan previously accumulated on catchment substrates, redepositing it in the lake.

Long-chain *n*-alkanoic acids (>C24) found in sediments mostly originate from epicuticular waxes of terrestrial higher plants and are transported through soil erosion and riverine input. The average chain length (ACL) of leaf wax components in sedimentary records, including *n*-alkanoic acid and the more commonly employed *n*-alkanes can be indicative of altered contribution of different vegetation types. This is, for example, based on the observation that these lipids in grasslands have longer chains than those originating from forests[20]. Although this general assumption has been challenged[21], ACL is still widely employed in paleovegetation research. In our record (Figure S16), a surprisingly stable ACL is observed, also post-LTL, suggesting that human presence did not significantly alter the composition of terrestrial vegetation. A massive clearance of birch forests and increase of grasslands would be expected to result in an increased ACL, as C26 and C28 alkanoic acids are the dominant species in *Betula*[22].

The only major change in ACL, i.e. a strong and fast decrease, takes place at the oldest part of the studied interval (~ 615-670 CE) and mostly results from an increase of C24 and C26 species. Such mid-chain length species are especially abundant in non-emergent aquatic macrophytes[23], including *Myriophyllum* [24] (Liu and Liu 2017), and are therefore consistent with the suggested replacement of marine taxa by a freshwater hydrophyte community strongly dominated by *Myriophyllum*.. Breakpoint detection in our high-resolution dataset suggests that this transition was completed around 676 CE.

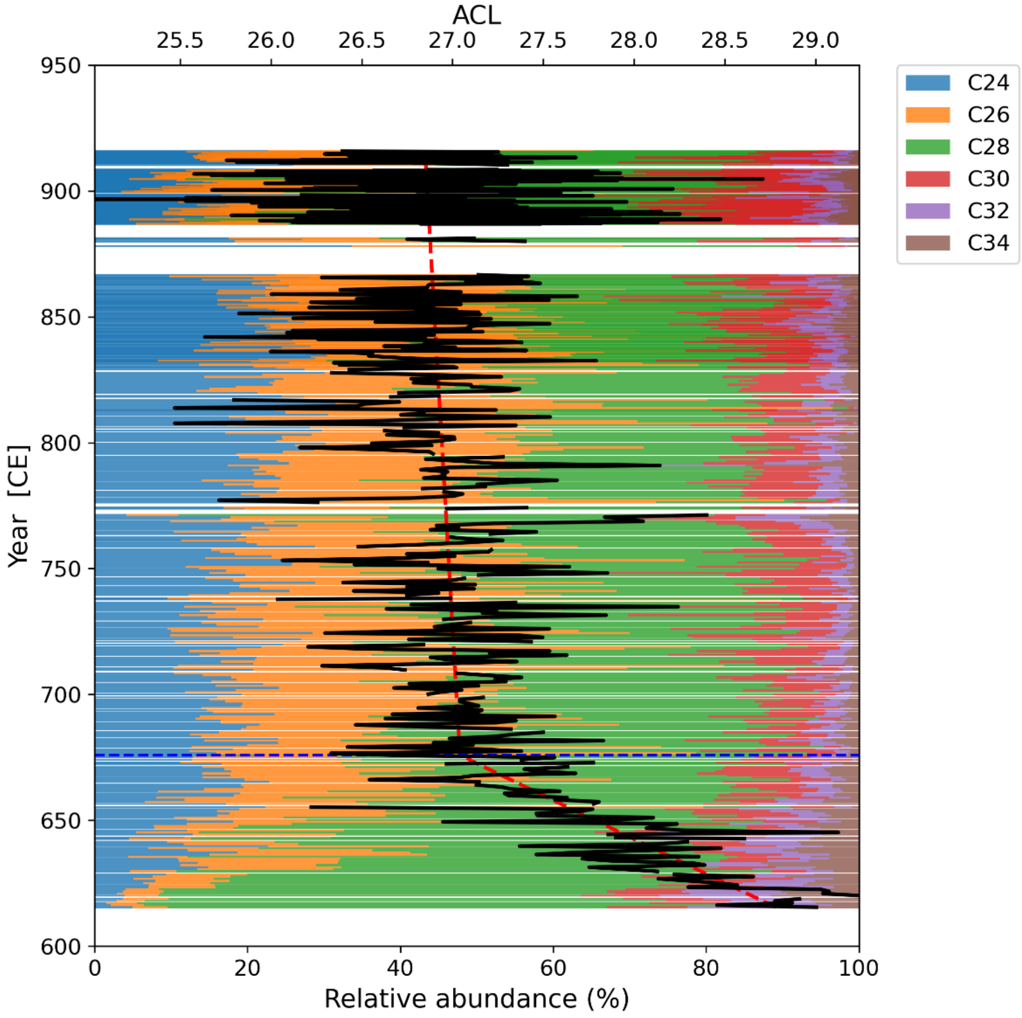

**Figure S16. Relative abundance of even numbered long-chain n-alkanoic acids and their average chain length (ACL, black solid line).** Piece-wise linear regression of ACL values (red dashed lines) identifies a breakpoint at 676 CE (blue dashed line).

**Chironomid Analysis**

We used Detrended Correspondence Analysis (DCA) to visualize the variance of chironomid assemblages in the whole sediment sequence along the main axes; gradient length was 3.4 standard deviation (SD) turnover units. After removing 19 bottom samples with no or only few chironomid remains found, we repeated the analysis and the gradient lengths was 2.6 SD, and consequently Principal Component Analysis (PCA) was used to visualize the assemblage variance in that part of the sediment sequence. Relative abundance (percentage) data were square root transformed prior the analysis and only taxa with minimal 20% fit on both displayed axes were shown. To examine the relationship of the elemental composition of the sediment to the assemblage composition Canonical Correspondence Analysis (CCA) was employed. Forward selection was used to select variables with significant influence on the taxonomic composition and $p < 0.05$ was considered significant. Only taxa with 20% fit on both displayed axes are shown. CANOCO 5.0 package [24] was employed for ordination analysis of assemblage data. Hierarchical classification using Bray-Curtis similarity of square root-transformed relative abundance data using stratigraphically constrained order of samples was used to distinguish zones of higher internal assemblage similarities.

**Results and description of zones**

Hierarchical classification distinguished four main Chironomid Assemblage Zones (CAZ) (Figs. S17 and S18).

**CAZ 1 (Marine phase; 202 – 182 cm, – ~700 CE).** The characteristic feature of this zone is the extremely low amount of chironomid remains, and thus low taxonomic richness combined with unstable assemblage structure. In most samples no, or only one remain was found, so there was no real freshwater community at that time. The remains found in this zone belong to taxa with rich representation in younger sediment layers, so it is likely that surface sediment from the lake somehow ended up in the bottom of the core as it was retrieved. Since the recorded taxa do not reflect an actual community, this zone was excluded from further statistical analyses.

**CAZ 2 (Isolation of the lagoon - Freshwater phase; 182 – 112 cm, ~700 – 1300 CE).** Head capsule count and taxon richness steeply increases at the beginning of the zone. The assemblage composition is stable along the whole zone with the dominance of *Chironomus anthracinus*-type and the cold-stenothermal *Orthocladius consobrinus*-type. *Arctopelopia* sp., *Heterotrissocladius grimshawi*-type, *Corynocera oliveri*-type/*Tanytarsus lugens*-type, *Microseptra insignilobus*/*contracta*-type, *Tanytarsus mendax*-type and *Micropsectra* type A occur in considerable amounts and frequencies only in this zone. The high representation of taxa that can thrive also in brackish waters, such as *Ablabesmyia*, *Procladius* and especially *Psectrocladius barbimanus*-type (which proportion is very high at the beginning of the zone but drops shortly after), indicates that brackish environment could persist for a shorter period (or saltwater intrusion could have happened). Head capsule deformities occur only sporadically.

**CAZ 3 (Little Ice Age and partly Saltwater Intrusion; 112 – 35 cm, ~1300 – 1900 CE).** Head capsule count slightly increases, and taxonomic richness is roughly equal to the previous zone. The dominant taxa from the previous zone remain but the relative abundance of *Chironomus anthracinus*-type slightly increases while that of *Orthocladius consobrinus*-type slightly decreases. *Arctopelopia* sp., *Heterotrissocladius grimshawi*-type, *Corynocera oliveri*-type/*Tanytarsus lugens*-type, *Microseptra insignilobus*/*contracta*-type, *Tanytarsus mendax*-type and *Micropsectra* type A gradually decrease and disappear from the sediment sequence. *Psectrocladius barbimanus*-type reappears in this zone and *Dicrotendipes* types become abundant and frequent together with *Cricotopus intersectus*-type indicating presence of macrophyte stands. In the middle of the zone morphological deformities become more frequent. PCA 1$^{st}$ axis scores slightly increase while axis 2 scores considerably decrease indicating changes along the second axis.

**CAZ 4 (Industrialization – pollution of lake water; 35 – 0 cm, ~1900 CE – 2014 CE)**. This zone saw the biggest compositional change in the sediment, as seen in both PCA axes scores. While head capsule counts increased considerably, taxonomic richness decreased due to the synchronized disappearance of many previously abundant taxa. Proportion of morphological deformities markedly increased. While the previously dominating taxa virtually disappear, *Chironomus plumosus*-type highly dominates together with *Tanytarsus gracilentus*-type and in the youngest samples *Procladius*; proportion of *Cricotopus intersectus*-type remains unchanged. The high chironomid abundance combined with low diversity indicates eutrophication while frequent deformities point to pollution with heavy metals.

**Principal Component Analysis**

PCA explained 63% of the total variability in the assemblage composition (1st axis: 48%, 2nd axis 15%). *Chironomus plumosus*-type correlated positively and *Orthocladius consobrinus*-type negatively with the 1st axis, while *Psectrocladius barbimanus*-type (negatively) and *Procladius* (positively) correlated with the 2nd axis. *Chironomus plumosus*-type remains are often numerous in subfossil records of warm, productive conditions. Increase of *Chironomus* usually indicates a transition of a lake into higher trophic state (eutrophication) due to increased nutrient inputs connected to oxygen depletion at the lake bottom[25]. Larvae of the genus, together with those of *Procladius* and *Psectrocladius*, can also thrive in brackish waters. *Psectrocladius* larvae are typically limnobionts, characteristic for the littoral zone[26,27]. Other species are associated with macrophytes[28]. Species of the genus show different response to pH; while some species are associated with calcareous waters (e.g., *P. barbimanus*), some are acidotolerat to acidophilic[29]. The genus covers a wide range of temperature conditions with *P. barbatipes*-type being closer to the warm end of the gradient, and *P. sordidellus*-type being restricted to colder lakes[30,31]. In case of the Lake Tjörnin sediment sequence *P. barbimanus*-type seems to correlate with saltwater conditions (see results of the CCA, Fig. S18), which may have increased the pH level of the lake.

**Canonical Correspondence Analysis (CCA, Fig. S18)**

Samples of the first zone are scattered along the biplot randomly, indicating instable assemblage composition (as a result of low head capsule counts). Assemblages characteristic of the second zone (dominated by *Tanytarsus mendax*-type, *Arctopelopia*, *Micropsectra insignilobus*-type) were mainly determined by high concentration of P, Al, Mn, Nb and Pr, which can indicate higher nutrient content and increased lake productivity. Samples of Zone 3 are scattered significantly along the 2nd axis, which correlated significantly with Cr, Ni and Br with *Psectrocladius barbimanus*-type and *Chironomus anthracinus*-type as characteristic taxa. Higher concentration of Br together with the presence of *Psectrocladius barbimanus*-type and partly of *Chironomus anthracinus*-type indicate saltwater intrusion in the lake. The assemblage composition of the last Zone 4 is driven by the high concentrations of Pb and Zn and is dominated by *Chironomus plumosus*-type. At the same time, morphological mouthpart deformities that may indicate high heavy metal concentrations increased significantly in this zone.

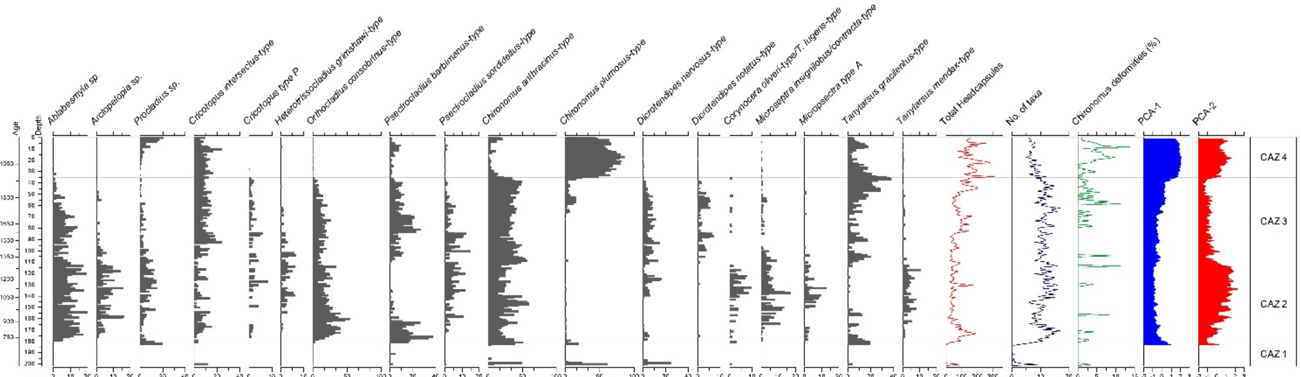

**Figure S17. Chironomid stratigraphy showing relative abundance changes along the whole sediment sequence**. CAZ 1 – 4: Chironomid Assemblage Zones. Only taxa with > 5% relative abundance in at least one sample are shown.

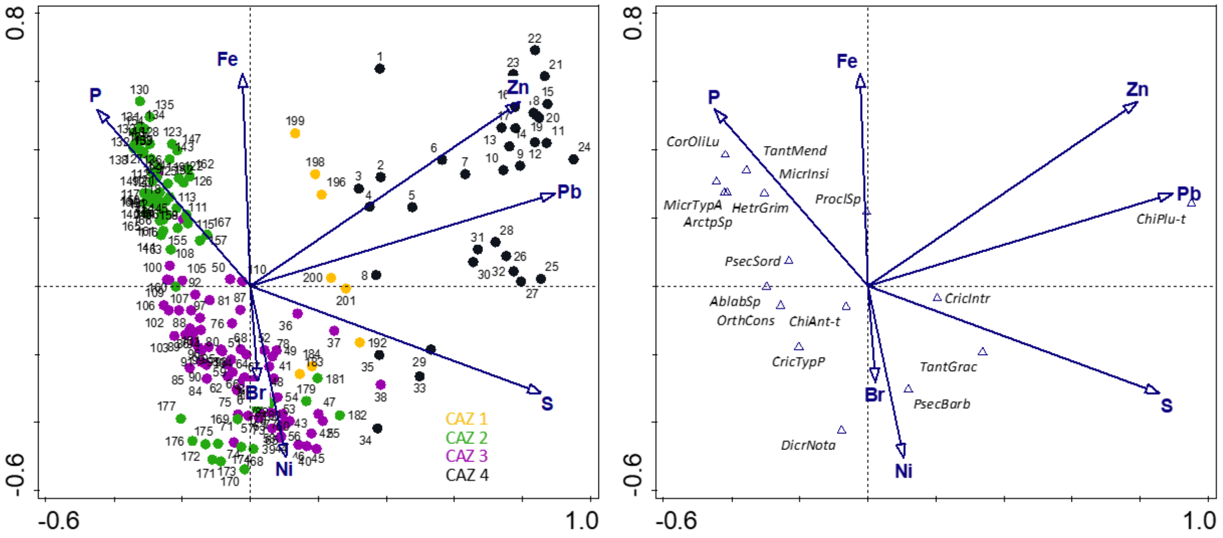

**Figure S18. CCA biplots visualizing the** Distribution of samples and variables (left), and chironomid taxa and variables (right). Only significant variables ($p < 0.05$) and taxa with minimum 20% fit on both displayed axes are shown. The bottom Zone 1 was not included in the analysis.

#### The Eukaryotic Fraction of the Metagenomes

**Details about versions and tools used in the Holi pipeline**

All raw reads were trimmed using fastp (v0.23.4) with options "--verbose --detect_adapter_for_pe --dont_eval_duplication --trim_poly_g --trim_poly_x --qualified_quality_phred 30 --average_qual 25 --length_required 30 --low_complexity_filter --complexity_threshold 30 --overrepresentation_analysis --correction --overlap_len_require 30 --overlap_diff_limit 5 --overlap_diff_percent_limit 20 --merge". Trimmed and merged reads were hereafter dereplicated with seqkit (v2.8.1) with default parameters and filtered for low complexity reads using bbduk (v39.06) with options "maxns=25 minlen=30 entropy=0.7 entropywindow=30 entropyk=4"). We next parsed all QC reads through HOLI for taxonomic assignment. To increase resolution and sensitivity of our taxonomic assignment, we supplemented the RefSeq (92 excluding bacteria) and the nucleotide database NCBI[32] with a recently published Arctic-boreal plant database PhyloNorway[33]. All alignments were hereafter merged and sorted by coordinate using samtools and parsed through filterBAM reassign and filter functions (v1.5.1) to refine taxonomic alignments and generate genome-wide statistics, with options "--iters 0 --min-read-ani 94 --min-read-count 3" and "--min-read-ani 94 --min-read-count 3 --min-expected-breadth-ratio 0.5 --min-normalized-entropy auto --min-normalized-gini auto --min-breadth 0 --min-avg-read-ani 90 --min-coverage-evenness 0.4 --min-coverage-mean 0 --include-low-detection", respectively. Cytosine deamination frequencies were then estimated using metaDMG (v0.4-93), by first finding the lowest common ancestor across all possible alignments for each read and then calculating damage patterns for each taxonomic level.

**Reproducibility for ancient environmental metagenomic samples**

We first tested the similarity between technical and biological replicates in ancient environmental metagenomics by analysing sediment samples from a lake core collected at Tjörnin, Reykjavík. Although replicates are essential for assessing the reproducibility and representativeness of small subsamples, their use in the field has been limited - likely due to the field's relative infancy and the high costs involved. To directly evaluate intra- and inter-sample variability, we analysed two subsamples of the same sediment layers (biological replicates) and split each of these extracts into two libraries (technical replicates). This setup allowed us to more systematically assess how well individual subsamples reflect the broader genetic signal and to better understand the reliability of ancient environmental DNA results.

Overall, 62 sediment layers were extracted for DNA spanning the entire core. 40 of these samples were selected and processed according to the description above, where 2 biological replicates were extracted and build into two separate dual indexed double stranded DNA libraries. A total of 225 libraries were built and subsequently sequenced on NovaSeq6000 S4 flowcells, running 100Bp paired end. Experimental setup, workflow and sequences generated and proportions of classified reads to eukaryotes can be found in Figures S19–S22.

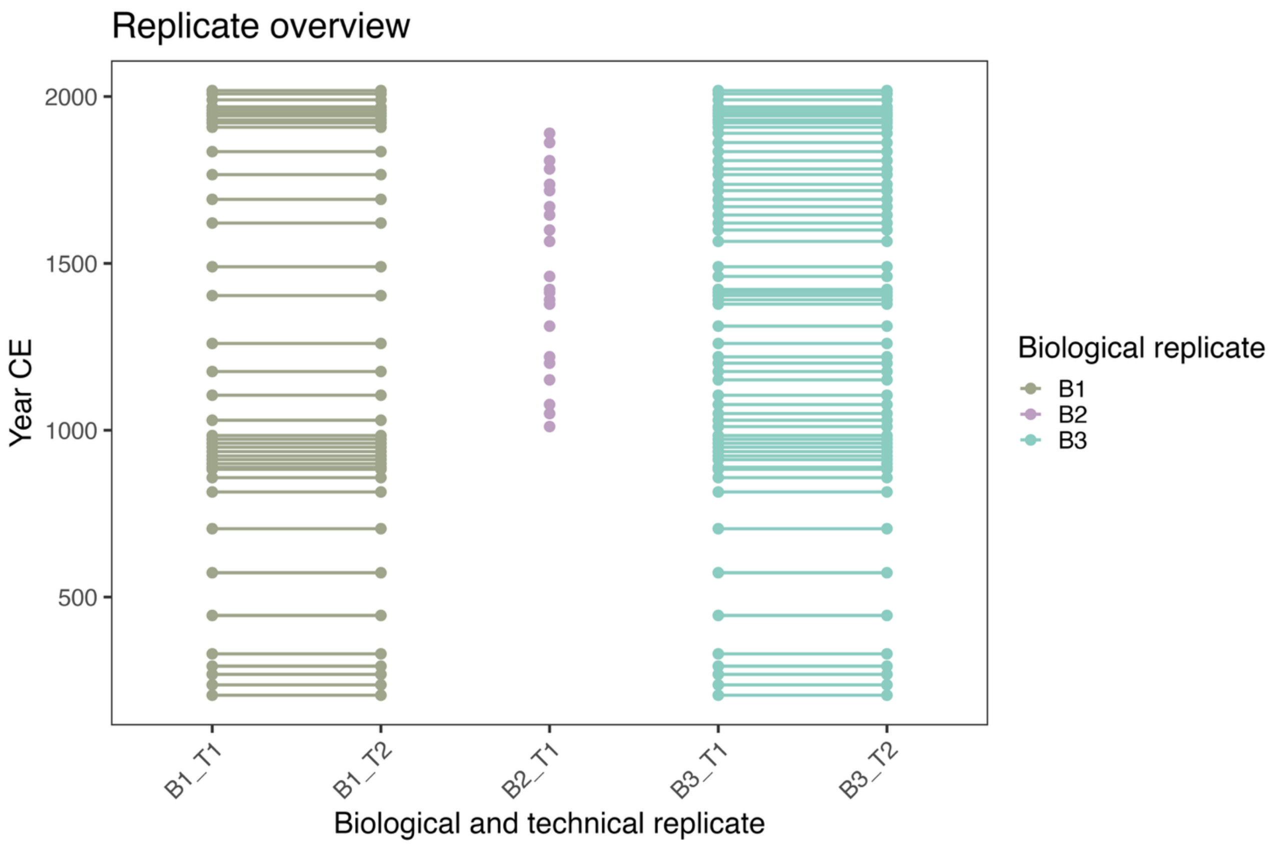

**Figure S19. Sample overview.** Technical replicates are connected by a line, and the corresponding biological replicates are the samples from the same depth but not connected by a line. The horizontal axis delineated tick marks are the individual NovaSeq 6000 flow cell runs.

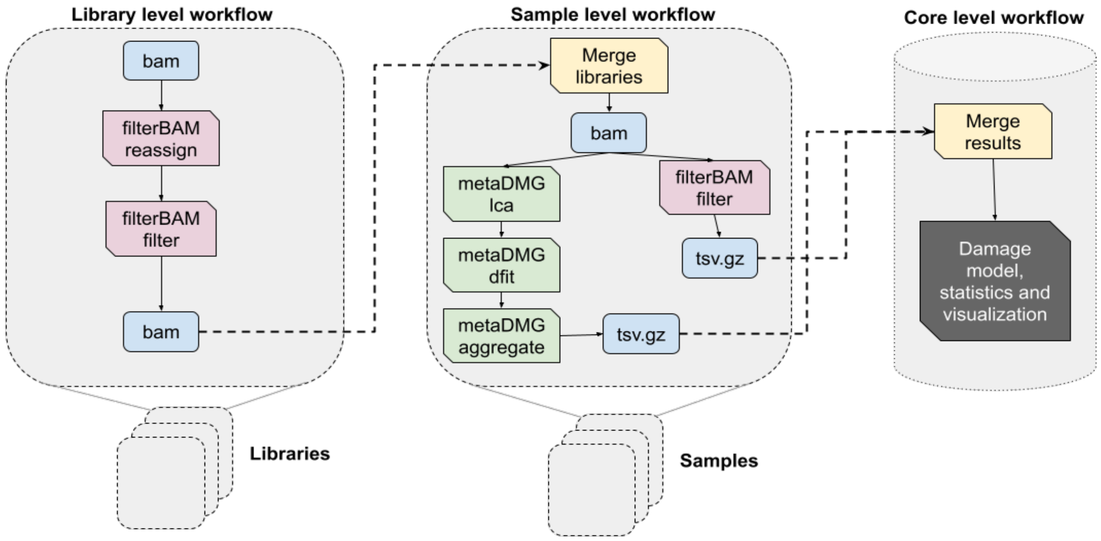

**Figure S20. Overview of the downstream analysis.** Each library is processed independently through filterBAM. Then, libraries from the same sample are merged, and processed through filterBAM and metaDMG. The results from these are then merged and processed in R for each layer throughout the core.

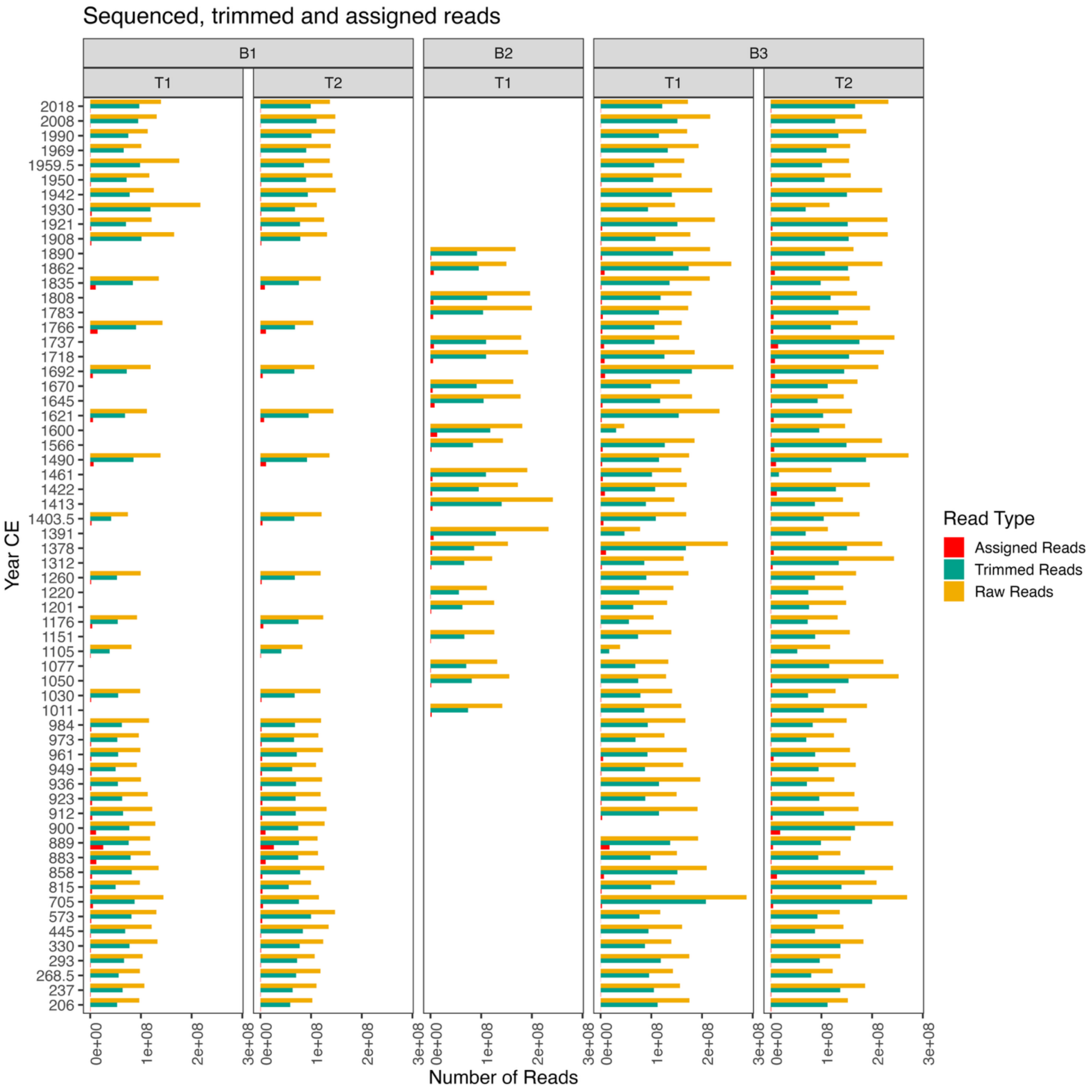

**Figure S21. Read counts by step in the pipeline.** Total raw reads sequenced (yellow); reads that collapsed after trimming adapters (green); and reads assigned to a species by metaDMG (red).

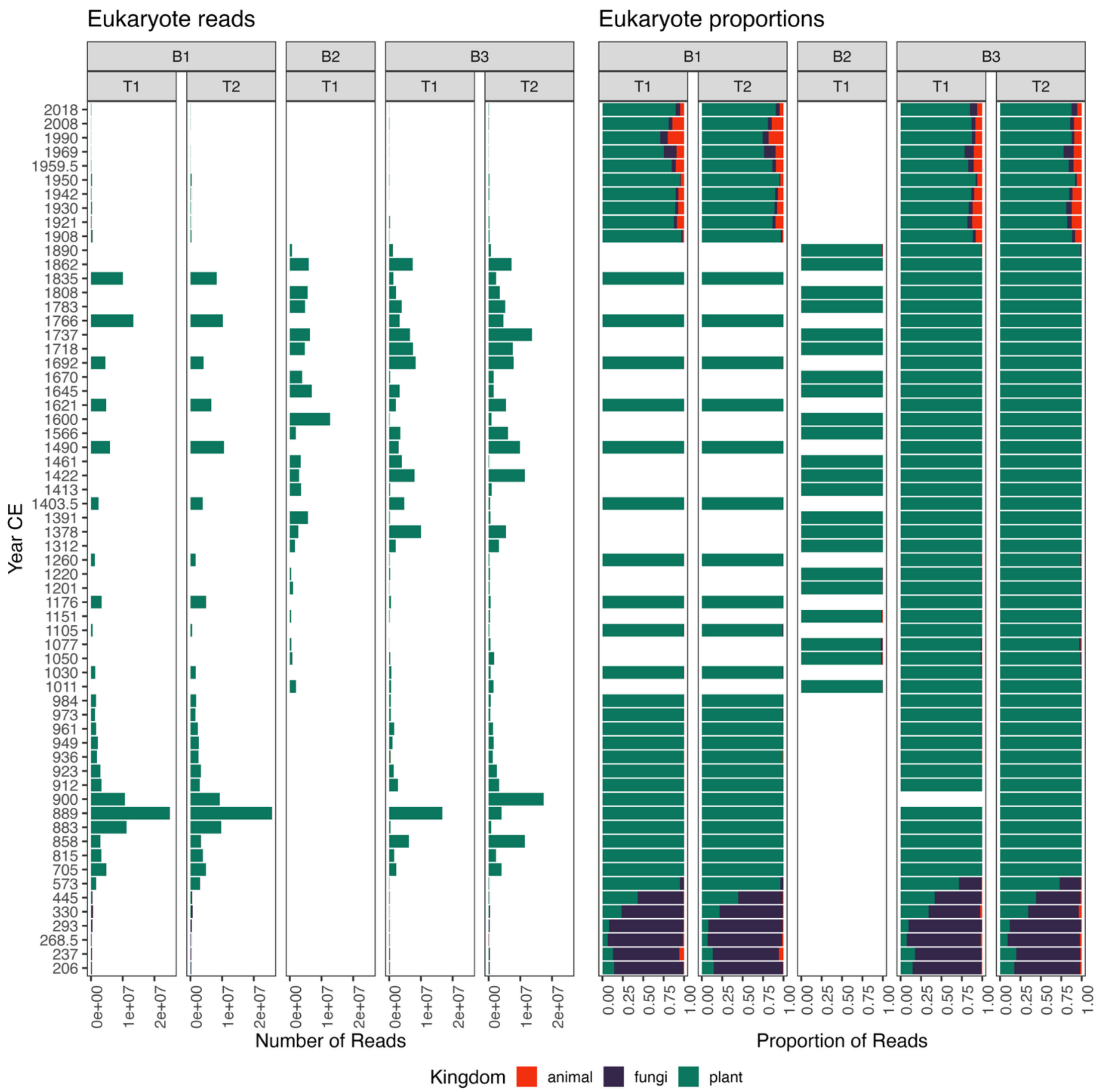

**Figure S22. Read counts by kingdom within Eukaryote**. On the left, the total number of reads assigned to each kingdom is shown, and on the right, this is represented as a proportion of all eukaryotic reads at each date within each replicate.

**Defining DNA Damage and Model Fit Quality to filter data**

For each taxonomic level, we estimated DNA damage using metaDMG[34], which provides both predicted damage values and model-derived noise baselines. To assess the quality of the estimation, we compared observed and predicted damage using Lin's Concordance Correlation Coefficient (CCC), a statistical measure of agreement.

A model fit was classified as good if all of the following conditions were met: the CCC ($\rho_c$) was greater than or equal to 0.85, the bias correction factor ($C_b$) was greater than 0.9, the p-value associated with $\rho_c$ was less than 0.1, and the confidence interval for the noise baseline parameter (c), as estimated, lay entirely between 0 and 1. If any of these criteria were not satisfied, the fit was classified as bad (Fig. S23).

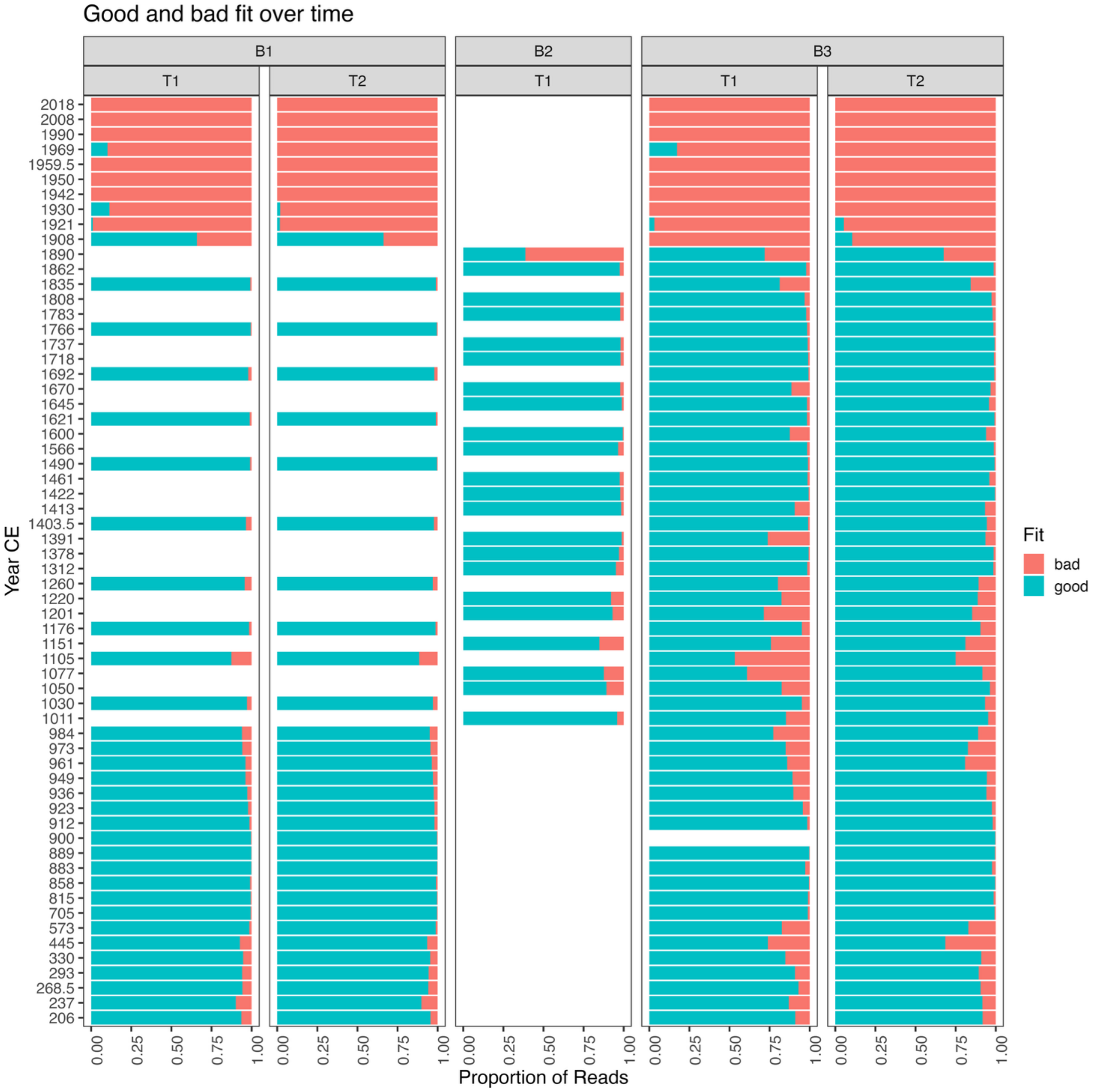

**Figure S23. Proportion of reads classified as good or bad fit to the damage model per replicate at each time point.**

**DNA Damage - age models and filter**

We next plotted all plant species with 500 reads or more, categorising them by their fit as mentioned above. We find overall similar trends exhibited by the damage, but we have a lower rate of deamination observed in the marine sediments compared to the lacustrine layers of similar age. Another key characteristic, that has been found by earlier studies, and which should be expected, is that the DNA damage decreases with age from the oldest lacustrine metagenomes to the youngest. In addition, we found that the youngest samples >1900 AD, had little to no DNA damage. Lastly,

we found that taxa from samples dating to between 1200-1900 CE, showed higher variation in the fit and hence the accumulated damage. In some cases, we found taxa with damage in older layers that showed poor fit in younger layers, even though the species have been historically documented during the entire period such as sheep and cattle (Fig. S24).
For the plant taxa with 500 reads or more and good fit to the damage model (for the dates between 660 CE and 1900 CE), we find the 5th percentile of DNA damage (A_b) across all taxa (Fig. S25). These quantile thresholds were smoothed with a loess regression, and taxa were retained only if their A_b exceeded the lower 95% confidence boundary for their corresponding date range. However, as noted above, this strict filter removed many cases of taxa in younger layers. Matched reads from sediment accumulated after the Landnám but before 1900 CE, did not always show clear signs of DNA damage possible because there was too little time for it to accumulate in these conditions.
As Pedersen and colleagues demonstrated[35], a thousand years at northern latitudes is often insufficient to accumulate significant DNA damage in lake sediments. Therefore, we setup a criterion in which we accepted taxa as genuine if they had been observed in older layers with a good fit and DNA damage.

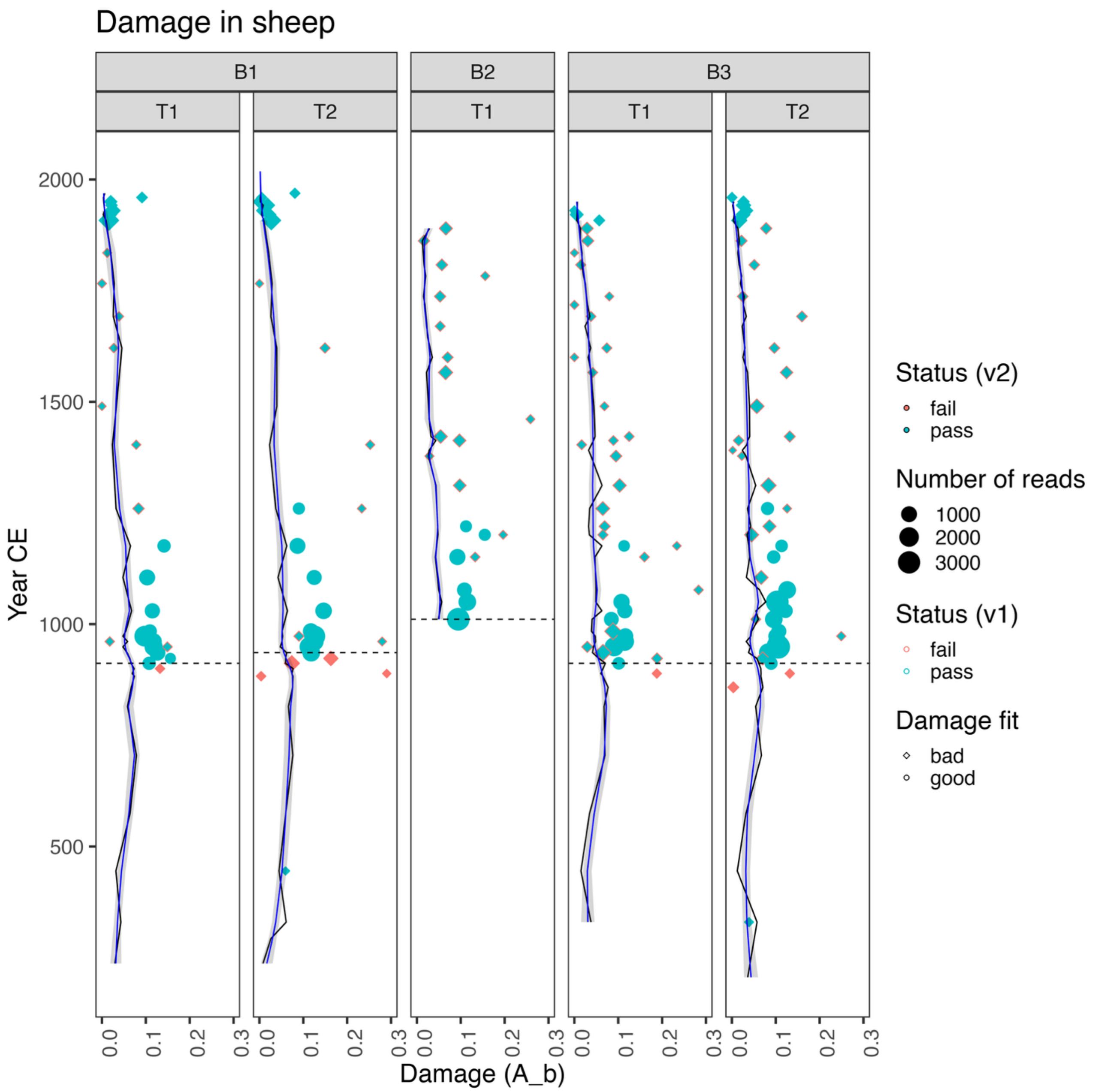

**Figure S24. DNA damage (median A_b) over time for Ovis (sheep) sequences, shown separately for each biological and technical replicate.** Point shape indicates model fit (circle = good, diamond = bad). The lower damage threshold is shown by black and blue curves with shaded uncertainty. Outline colour denotes the original status filter (v1), while fill colour shows the updated filter (v2) applied after the "oldest pass date" (horizontal dashed line). This highlights sequences failing v1 but passing v2, and vice versa.

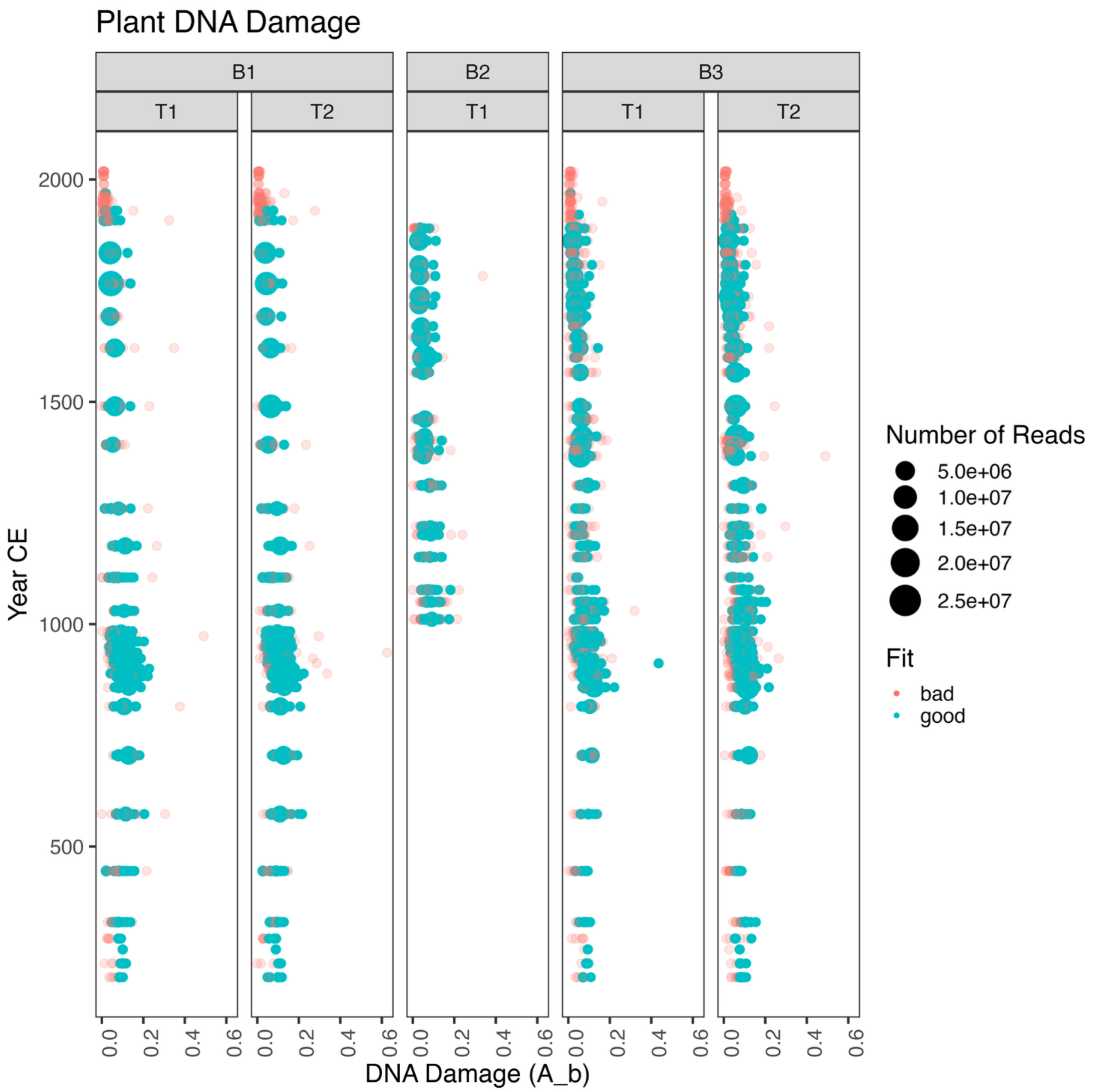

**Figure S25. Plant taxa with >=500 reads classified split by technical and biological replicates.** Blue dots indicate a good fit found using the CCC and thresholds mentioned above. Red dots indicate a bad fit.

**Comparing biological and technical replicates**

We sought to assess the structure of eukaryotic communities across both biological and technical replicates by comparing their individual metagenomic profiles using Principal Coordinate Analysis (PCoA). This allowed us to reduce the complex, high-dimensional data into principal coordinate axes, and visualize the similarities or dissimilarities between both types of replicates across the entire data (Fig. S26) and between the age of the samples (Fig. S27). Through this approach, we evaluated the extent to which replicate type contributed to variation in eukaryotic community composition.

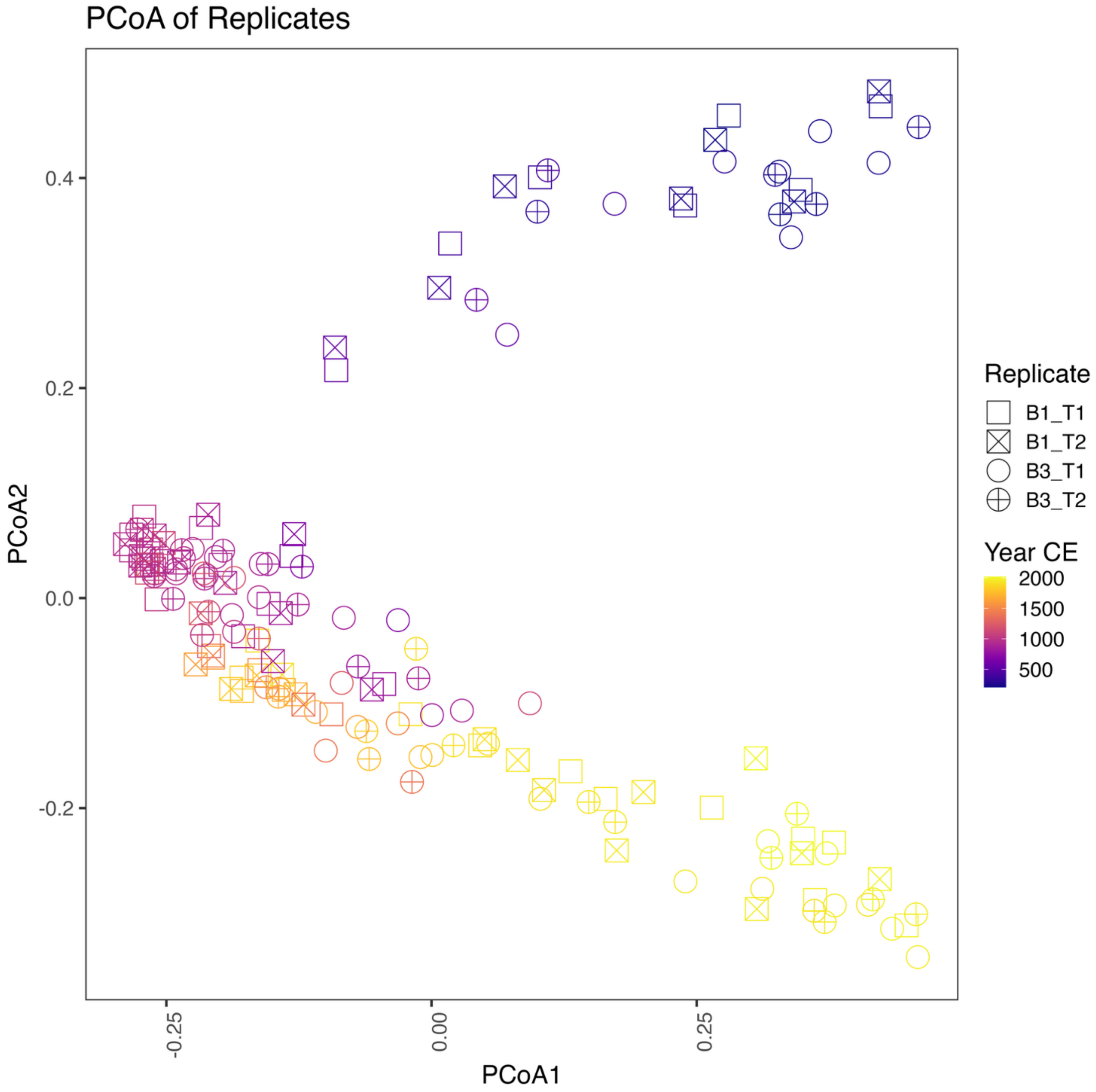

**Figure S26. PCoA of eukaryotic community profiles across biological and technical replicates.** The PCoA compares only samples from layers that have both a biological and a technical replicate.

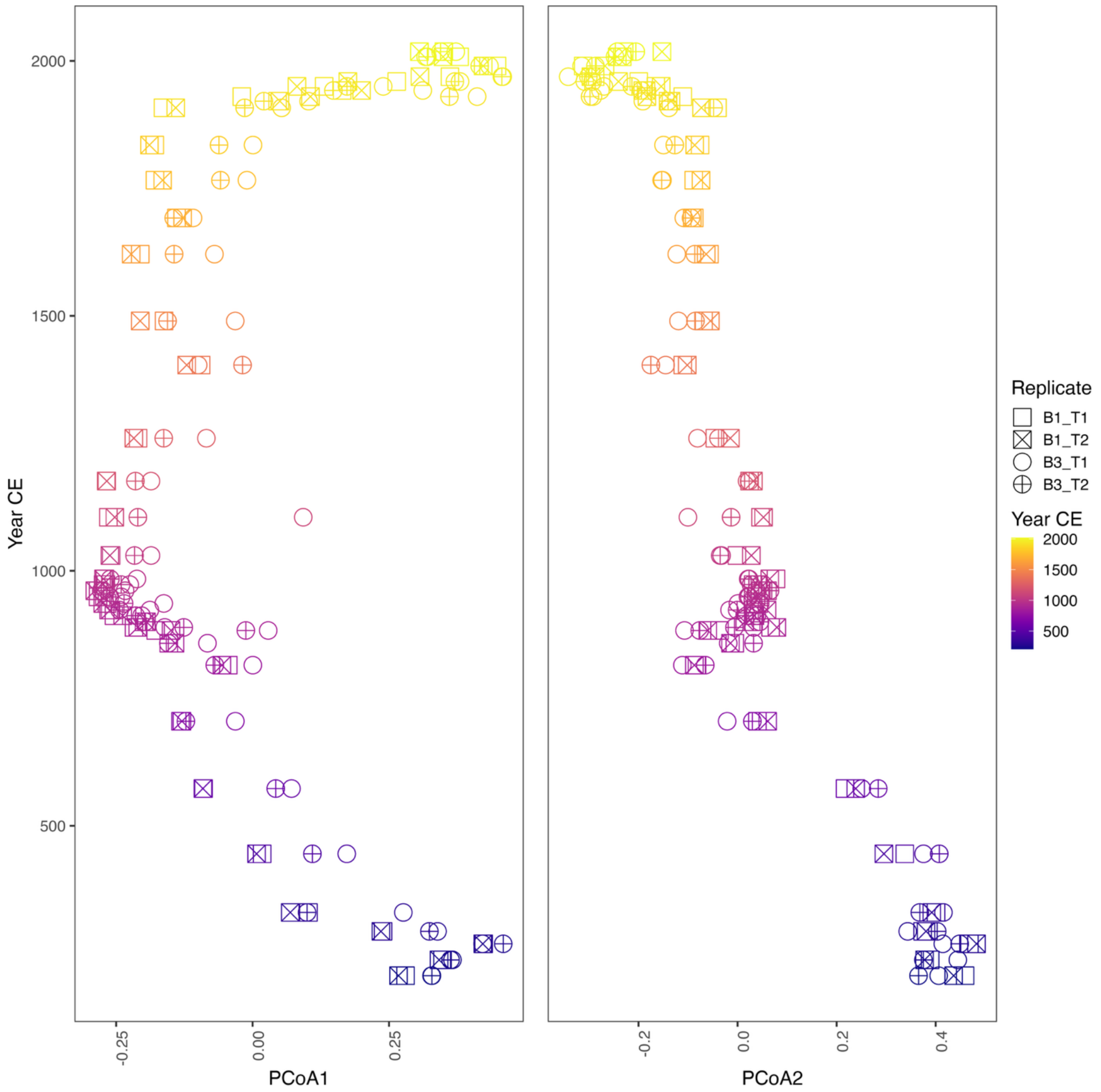

**Figure S27. PCoA of eukaryotic community profiles across biological and technical replicates, splitting the data by principal coordinate axes (1 & 2) and plotting these along the age.**

From Figure S26–27, we conclude that the variability within both technical and biological replicates are neglectable small, and that the main driver of the differences is that of time which relates to the environmental changes rather than the subsampling or technical nature of the data. It was therefore decided to merge all alignments with a post FilterBAM reassign step, to increase the number of sequences per sample and hence all downstream statistical power.

**Combined dataset with all replicates**

We then created a combined DNA damage - age model keeping the thresholds and criterions from the individual models split by replicates (Fig. S28). And plotted the 20 most abundant plant taxa and animals and their respective DNA damage by age as examples of the dataset (Figs S29 and S30).

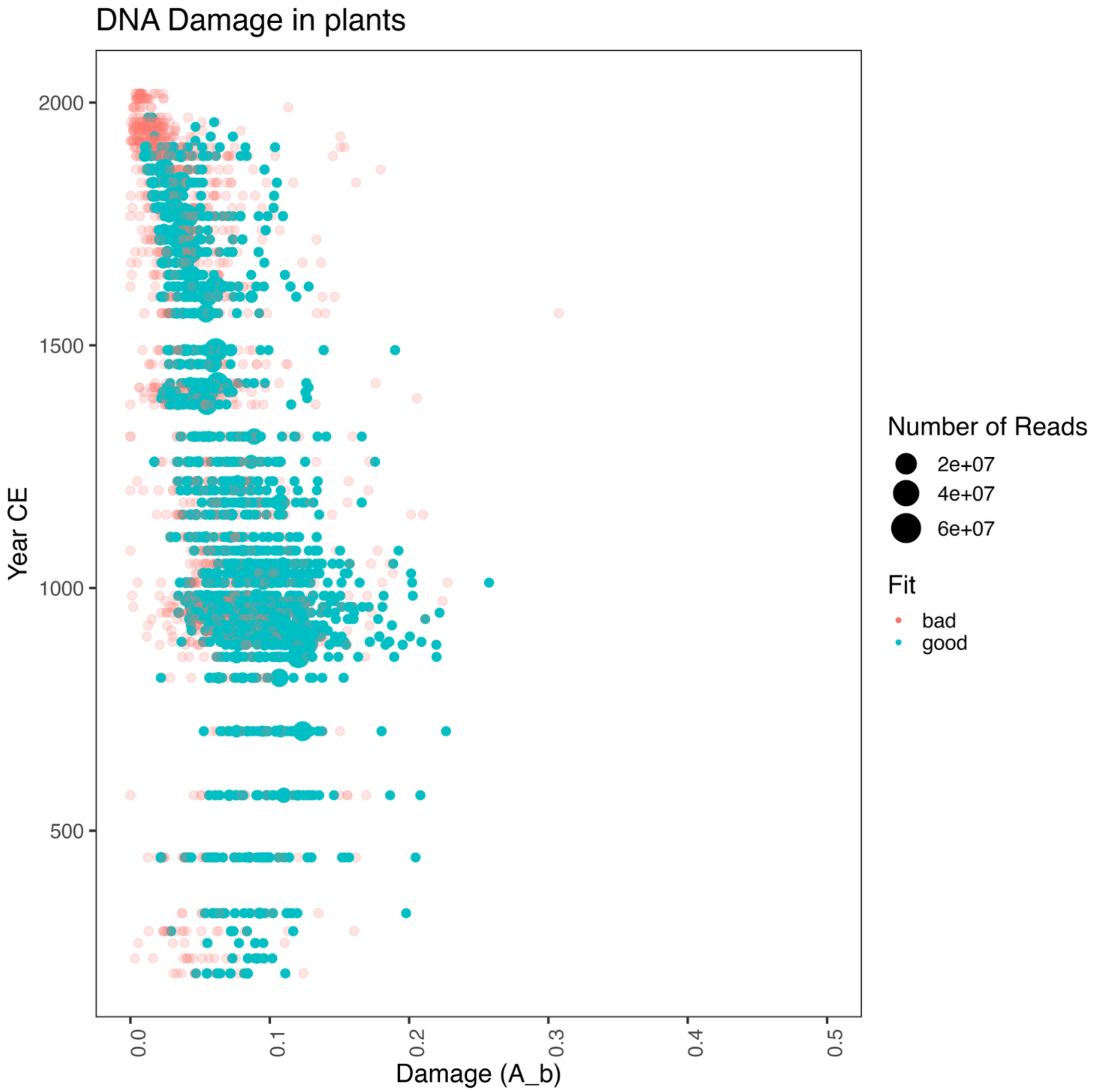

**Figure S28. Combined DNA damage - Age model for all plant taxa >=500 reads.** The black line shows the 5th percentile of the damage at each year, and the blue line is a smoothed fit (with confidence intervals) of this.

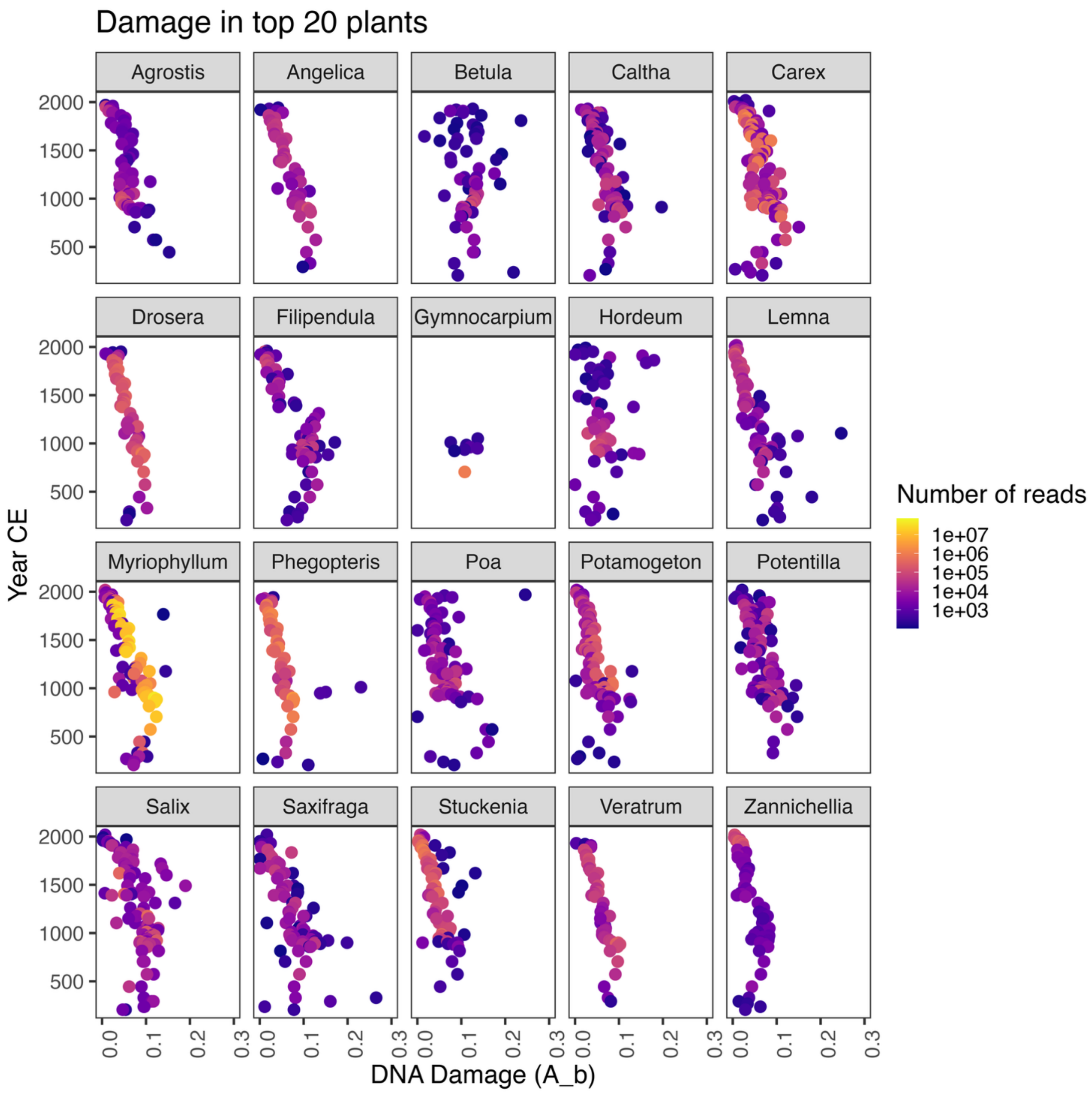

**Figure S29. Twenty most abundant plant taxa and their respective DNA damage by age.**

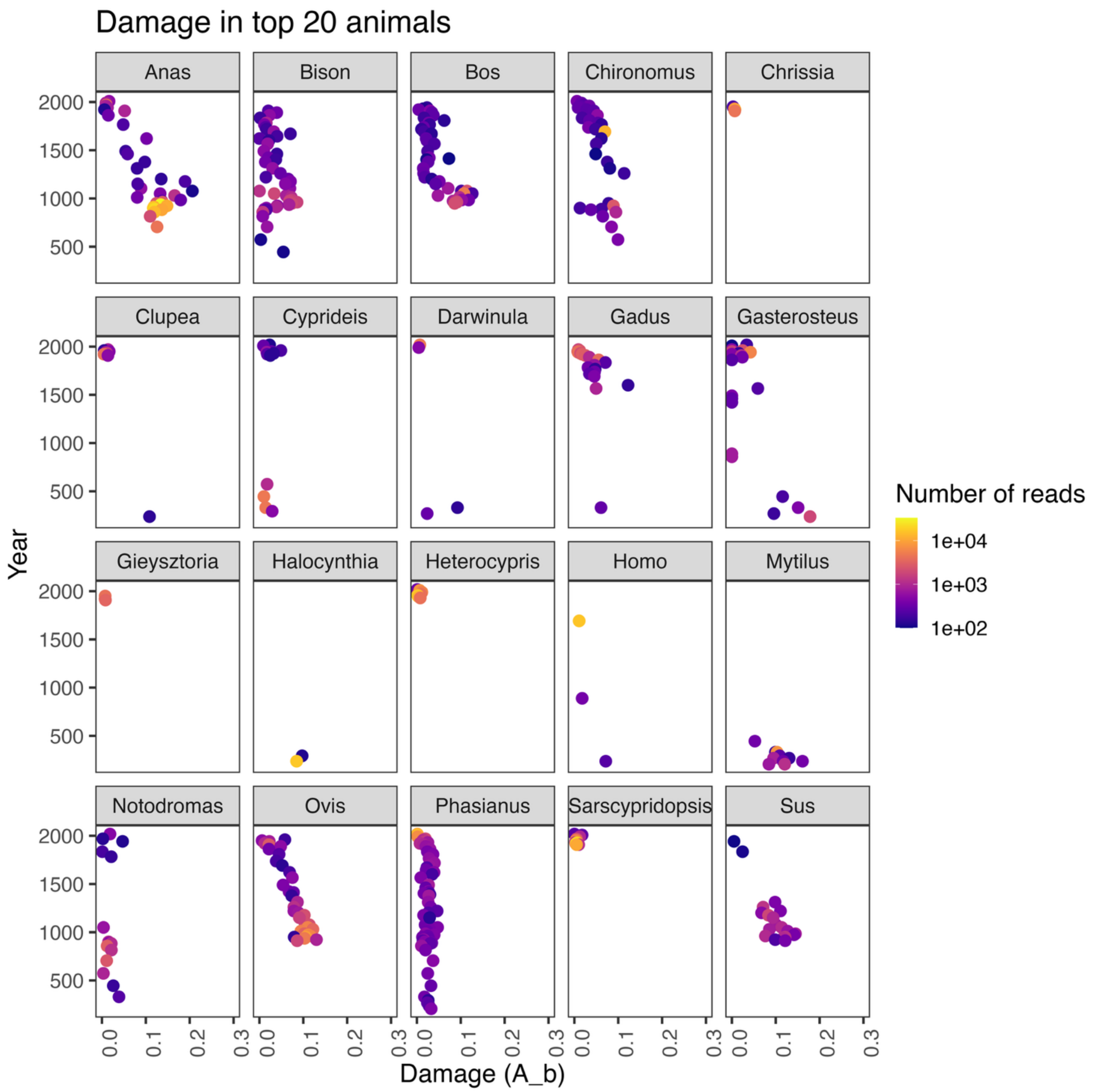

**Figure S30. Twenty most abundant animal taxa and their respective DNA damage by age.**

Rarefaction analysis was performed to evaluate whether sequencing depth was sufficient to capture the underlying taxonomic diversity. To assess whether further sequencing would identify more taxa, we generated rarefaction curves for each sample separately for plant and animal genera parsing the filtering criteria above but before the minimum read number filter. The R package vegan (v2.7)[36] was used to compute rarefaction curves based on repeated subsampling without replacement. For each sample, expected genus richness was plotted against the number of subsampled reads, with separate curves shown for plants and animals. Plateaus in the curves were used to evaluate whether additional sequencing would be expected to yield further genera. Visual inspection indicated that more than 95% of samples had reached saturation in taxa discovery (see Fig. S31).

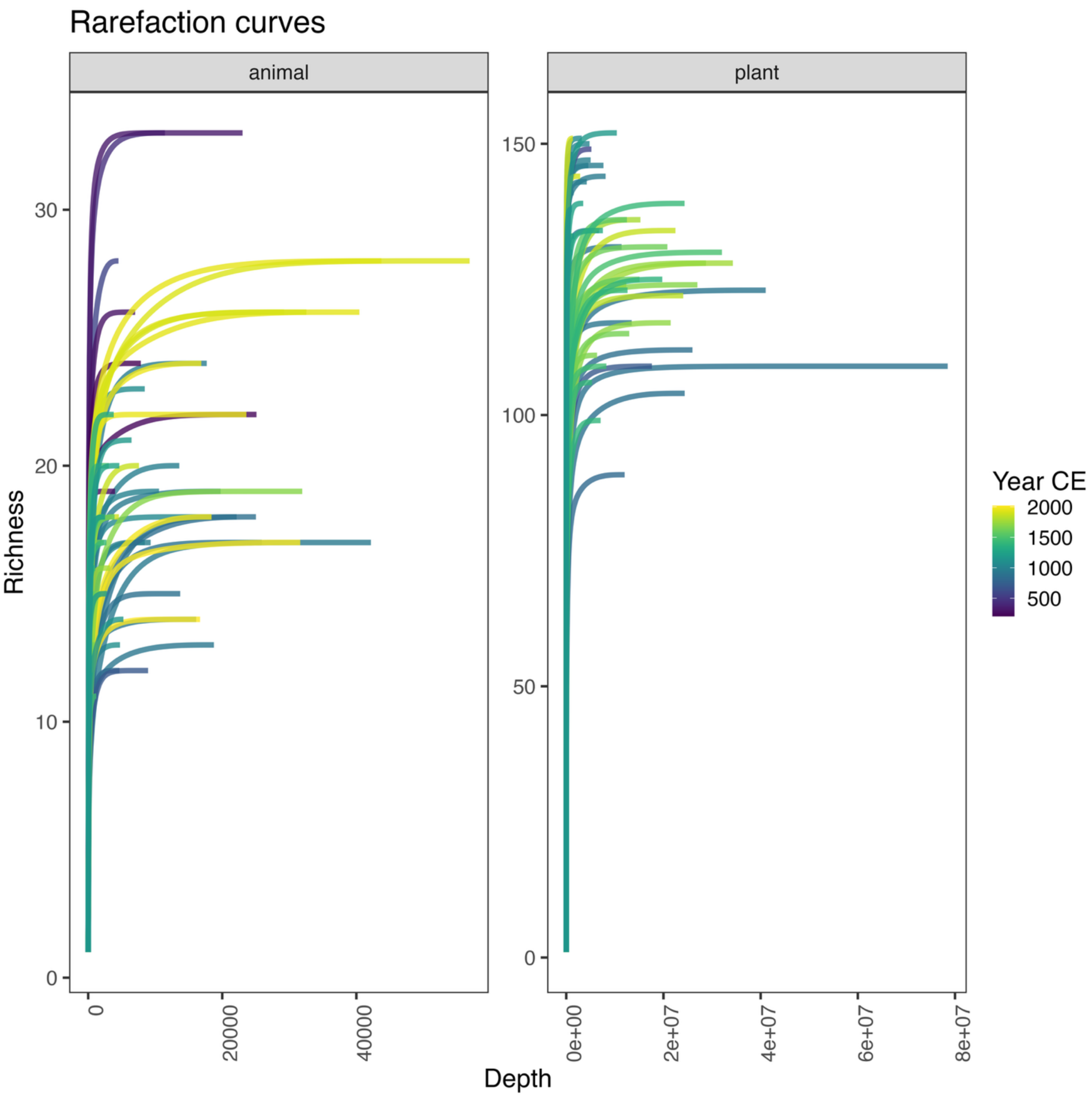

**Figure S31. Rarefaction curves for each metagenome merged by layer.** The two panels show animals (left) and plants (right), with curves coloured according to the sample age.

**Negative controls**

Following the extractions and library preparation, ten controls were included to test for reagent contamination and potential cross-contamination (Figs. S32–S33). All sequence data from the controls were processed through Holi and assessed for DNA damage to evaluate whether ancient DNA or other sources of contamination were present, like the other data and samples. Although we observed a small overlap between some taxa in the samples and those in the controls, DNA damage results indicated that these sequences do not derive from ancient sources. Only five taxa exhibited any degree of DNA damage, which may reflect cross-contamination or sequencing spillover, although the exact source remains uncertain. In any case, the contribution of these signals is negligibly small and does not affect the ecological reconstructions or the conclusions drawn in this study.

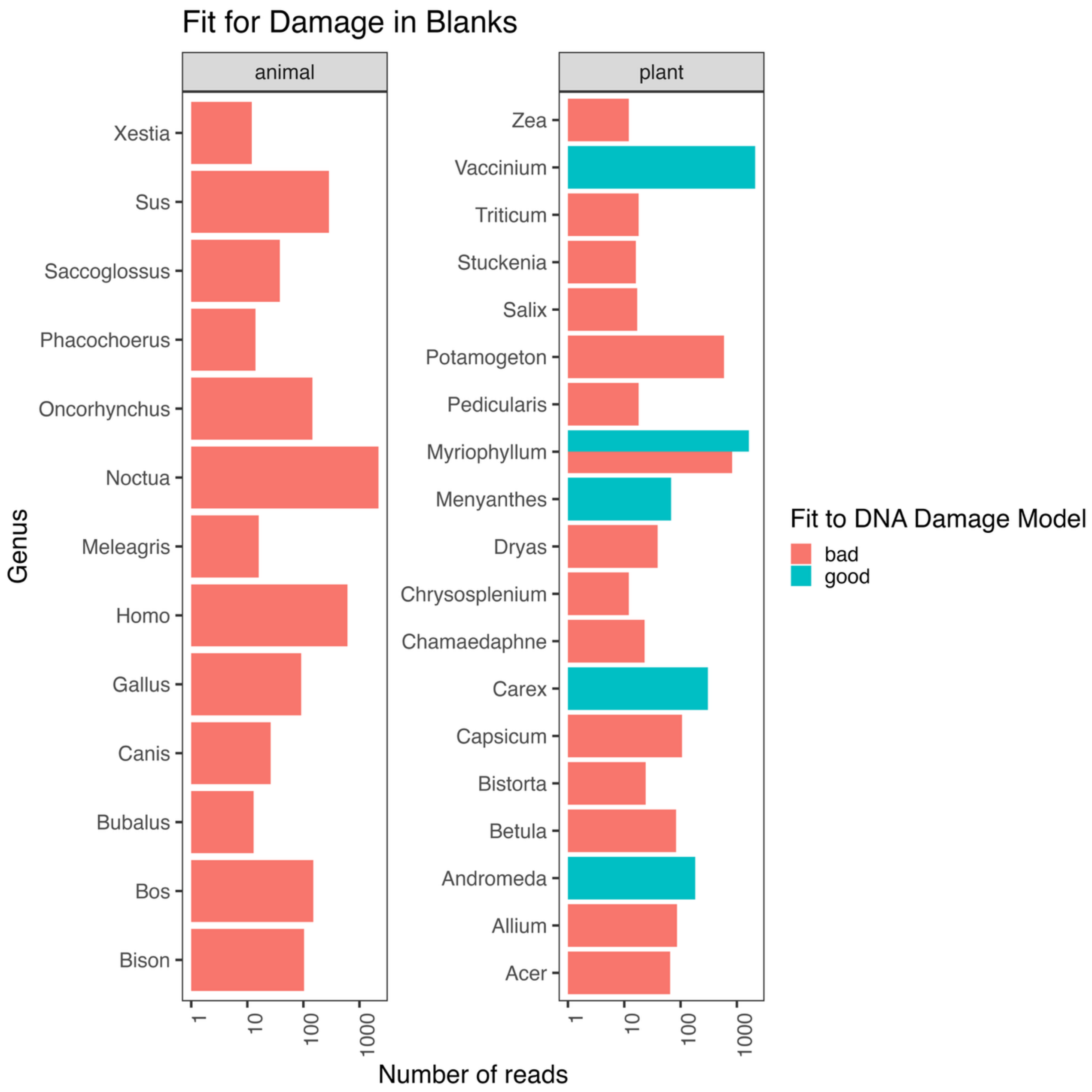

**Figure S32. Testing the fit of the DNA damage using the CCC method described above with all sequence data from controls merged.** Where good equals taxa with damage and bad are taxa that do not fit the expected DNA damage patterns.

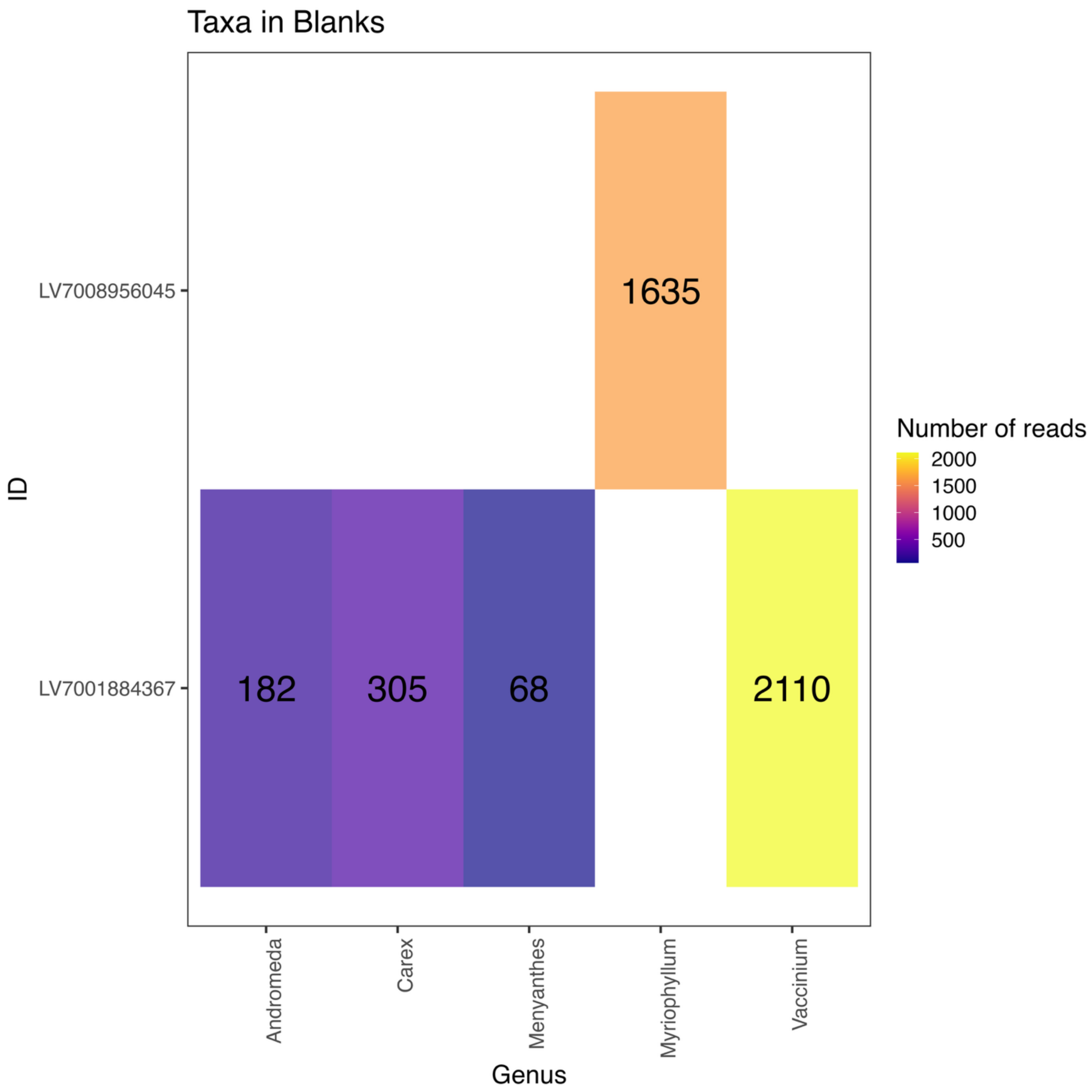

**Figure S33. Plotting out the distribution of taxa in their respective control samples from which the taxa have ancient DNA damage patterns.** Numbers are count of classified sequences to the respective taxa.

**Chloroplast DNA and TrnL barcodes**

In addition to the taxonomic profiles based on both the organelle and autosomal DNA references. We sought to profile the plants using a standard ancient plant DNA mini-barcode (trnL g-h) that generally are applied to ancient DNA.

We evaluated the specificity and sensitivity of two reference databases: a complete trnL gene database and a barcode-only database. Simulated ancient DNA datasets were generated from chloroplast genomes of two representative Arctic plant taxa to assess performance.

*Betula nana* (NCBI accession NC_033978.1)
*Dryas alaskensis* (NCBI accession NC_088483.1)

We used gargammel v1.1.4 to produce simulated FASTQ files with an average fragment length of ~45 bp and a deamination damage profile modelled after Günther and coworkers[37] as shown in Figure S34. Coverage ranged from 280× to 3×, corresponding to datasets of 1M, 500k, 250k, 100k, 50k, and 10k reads, each simulated in 10 bootstrap replicates.

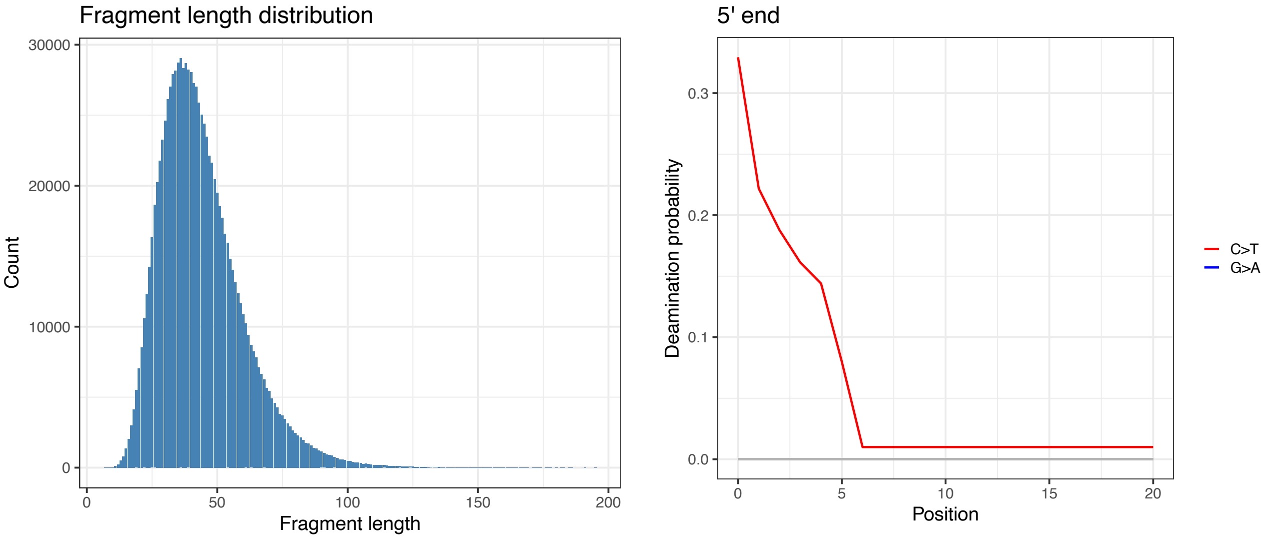

**Figure S34. Example of the degree of DNA damage and the fragment length distribution from the simulated data.**

We next constructed a comprehensive database of chloroplast sequences from the Viridiplantae clade by downloading all available full chloroplast genomes from NCBI RefSeq on July 11th, 2025, using the query:

("Viridiplantae"[Organism] OR Viridiplantae[All Fields]) AND (refseq[filter] AND chloroplast[filter]AND("100000"[SLEN]:"500000"[SLEN])).

This search yielded **12,816 complete chloroplast references**. From this collection, we extracted the **trnL gene sequences** when annotated in the GenBank entry, producing a trnL-specific database of **11,791 references** (differences arise from missing or inconsistent GenBank annotations, not from absence of the gene itself). To refine this into the shorter **trnL barcode region**, we applied the program **rCRUX** (https://github.com/CalCOFI/rCRUX; Curd et al., 2024) with standard parameters, in combination with custom R scripts (https://github.com/nicolaavogel/barcodeMiner/tree/main/scripts). We used the primer pair described in Taberlet et al. (2006): forward GGGCAATCCTGAGCCAA and reverse CCATTGAGTCTCTGCACCTATC. This produced a **barcode-level database comprising 10,851 sequences**.

Overall, we identified **9,967 references** containing both a full trnL gene and its barcode region. In addition, **692 references** contained a barcode region but lacked trnL annotation in NCBI, while **1,823 references** had annotated trnL sequences but no extractable barcode region (Fig. S35). These discrepancies likely reflect annotation differences or too low similarity between primer and reference rather than true absence of the trnL gene. For transparency, accession numbers and taxonomic details of the missing entries are provided in the supplementary datasets trnL missing data and barcode missing data.

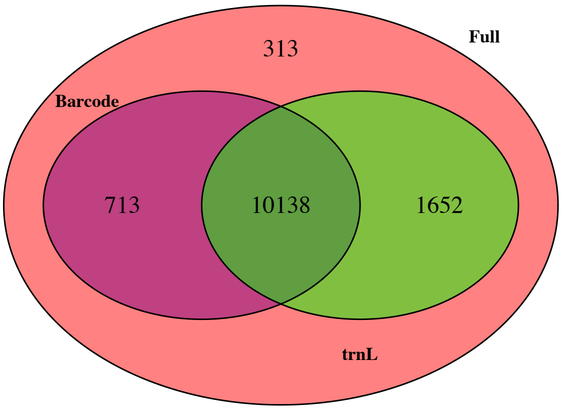

**Figure S35. Venn diagram showing the overlap between the three Viridiplantae chloroplast databases.** The red circle represents the full chloroplast genome database (1), including 313 sequences lacking both trnL gene annotation in NCBI and identifiable barcode regions from the primer sequences. The green circle represents references with annotated trnL gene information, while the purple circle represents references for which the barcode region could be successfully

We first mapped all simulated reads against the barcode. And calculated the probability of having 1, 10, 50, or 100 reads assigned to any taxonomic rank (Figure S36). We find that the probability of detecting even a few reads (10 reads) aligning to the trnL (g-h) barcode requires to have a relatively high depth of coverage to the chloroplast in both species (>50X in both cases).

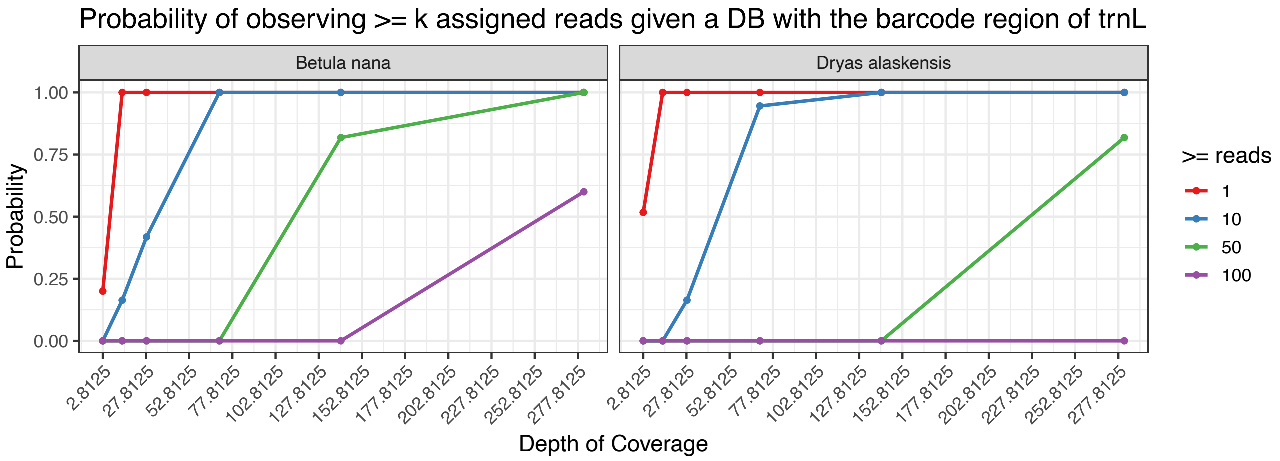

**Figure S36. Probability plot for the simulated data mapped to the trnL barcode.**

We then mapped all simulated reads against the trnL gene database. Again, calculating the probability for taxonomically classifying 1, 10, 50 and 100 reads at any rank (Figure S37). We find that the probability of detecting even reads (10 reads) aligning to the trnL gene substantially increases when aligning to the full gene sequence. Despite the increased sensitivity depth of coverage still needs to be relatively high, with a minimum (>3X in both cases) to classify 50 reads.

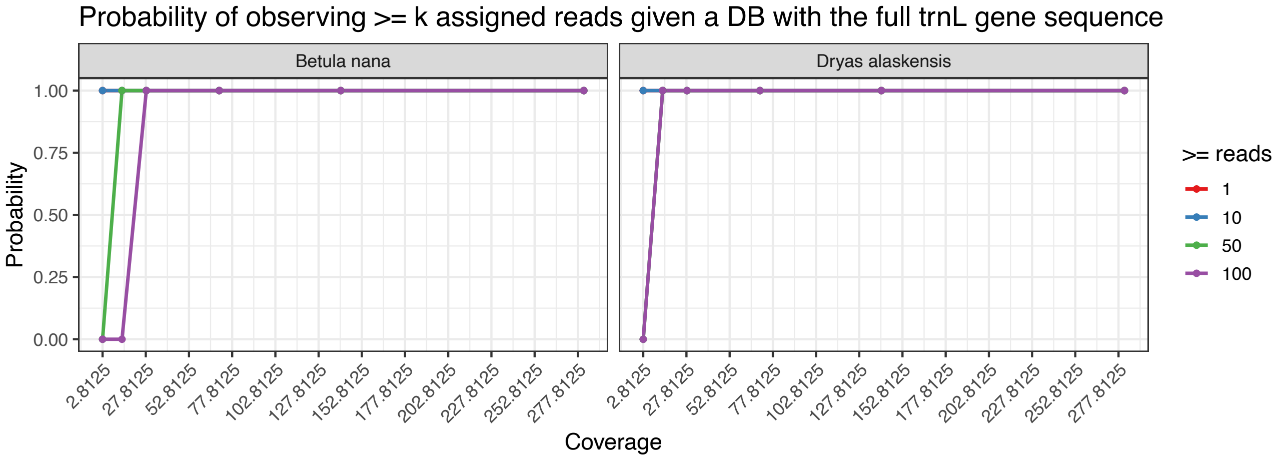

**Figure S37. Probability plot for the simulated data mapped to the full trnL gene.**

Lastly, we performed taxonomic profiling using both the barcode and the gene databases on 10 selected samples. Trimmed and quality-controlled reads were mapped using bowtie2 using the same settings as for the taxonomic profiling using the entire genome reference database and lastly parsed through metaDMG lca for taxonomic classification.

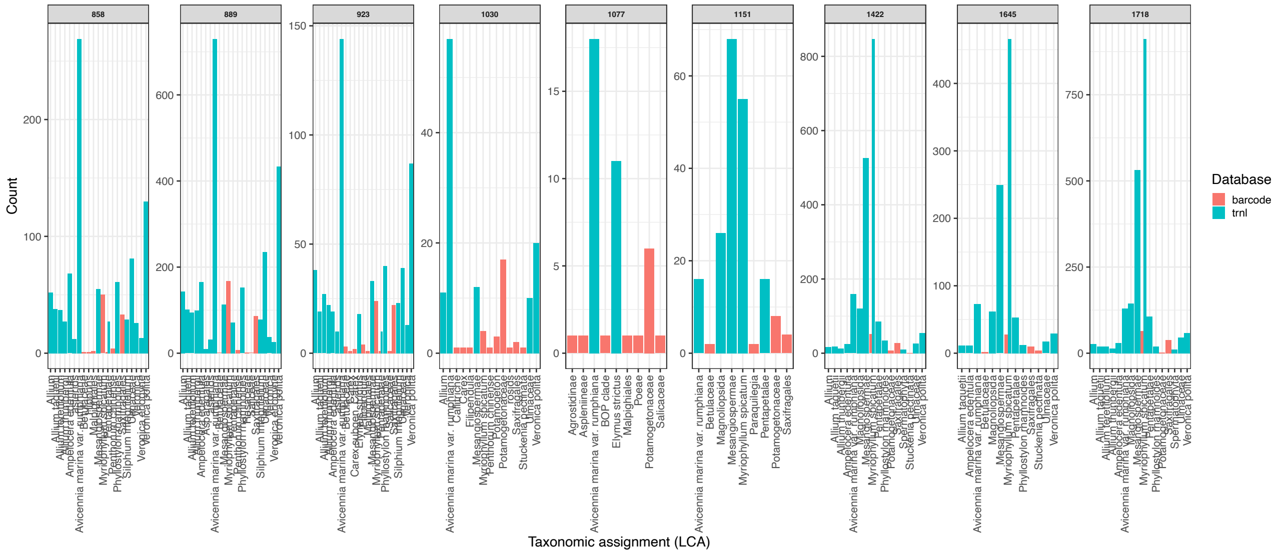

**Figure S38. Unfiltered number of classified reads assigned at all taxonomic levels for the barcode database, assignments for the tnrL database were filtered with a minimum assignment of 10 reads to minimise the noise in the samples.**

Our analysis shows that *Myriophyllum spicatum* is the dominant taxon, consistent with patterns observed in the broader empirical dataset (Figure S38). This species was also one of the few to surpass the minimum threshold of 100 reads set for inclusion in the analysis. In addition, *Veronica polita* (Gray Field-Speedwell) was detected at lower abundance, both genera are members of the Icelandic flora. The remaining two identified species *Avicennia marina* (grey mangrove) and *Phyllostylon rhamnoides* are subtropical or tropical. These latter identifications likely reflect limitations inherent in competitive mapping with short reads against databases restricted to a subset of plant genes.

**Description and Interpretation of the Eukaryote Metagenomic Records**

Environmental DNA enables the detection of many referenced organisms in sediment metagenomes, including those without preserved or identified macro or micro sub-fossils. Changes in the relative abundance of matched DNA sequences through the record can be interpreted as reflecting variation in the amount of biomass deposited at the site over time, much as sub-fossil remains (e.g. pollen, diatoms, algae, foraminifera) are used in paleoecology. While this provides valuable insights into temporal dynamics, translating eDNA signals into actual species counts or precise measures of abundance across the broader landscape remains challenging. As with other paleoecological proxies, a range of taphonomic and environmental factors constrain the representativeness of eDNA, but these uncertainties are inherent to all such approaches.

Extended Data Figs. 2–4 display the percentages of reads matched to metazoans, aquatic angiosperms and algae and all terrestrial viridiplantae, calculated from the read sums of each respective category. In these plots, variation in the type and proportion of matched reads is shown stratigraphically, analogous to a pollen diagram.

For each diagram, the total number of matched reads is represented as histograms with widths scaled to the deposition interval of the sample. Each dataset has been divided into five biozones: A-1 to A-5 for aquatic plants, M-1 to M-5 for metazoans, and T-1 to T-5 for terrestrial Viridiplantae. Biozones were defined using a stratigraphically constrained cluster analysis with the information statistic as the distance measure (CONISS). The resulting zones, together with the cladogram derived from dispersion distances, are shown in the final column of each figure, alongside a rate-of-change statistic and, in some cases, the Shannon diversity index.

The zonation in the diagrams do not correspond precisely because the aquatic and terrestrial biomes are not fully in phase. However, each begins with a marine zone at the base of the sequence (Zone 1), followed by a distinct post-marine or pre-settlement zone (Zone 2). The settlement zone (Zone 3) begins at approximately 900 CE in all diagrams and persists for two to four centuries. This is followed by a retraction zone (Zone 4) and, finally, Zone 5, which corresponds to 20th-century urbanization and eutrophication. Each diagram also includes the episodes used to describe the sequence in the main text, plotted adjacent to the age scale. Unlike the zones, these episodes are not statistically derived but represent a best-fit interpretation that integrates all compiled proxies presented in this study, together with historical and archaeological data, to partition the sequence. The correspondence between these episodes and the aquatic, metazoan, and terrestrial viridiplantae biozones can be evaluated directly by comparing the episodes with the zones in each diagram.

Extended Data Figure 5 presents a pollen diagram showing the proportions of pollen and spores identified from the same sediment sequence and depths as the metagenomic analyses of aquatic and terrestrial plants, allowing direct comparison. For this study the percentages were calculated by dividing individual taxa counts by the total number of pollen and spores in the sample regardless of habitat (i.e. terrestrial and aquatic taxa were included in the sum). The modelled pollen accumulation for *Betula*, total pollen and total charcoal particle accumulation are included in the diagram for comparison with the percentages. Palynologists recognize that pollen percentages do not correspond directly to the proportions of plants in the vegetation, owing to differences in pollen production and dispersal potential. Similarly, although some correspondence between biomass and DNA sequence abundance is evident, relative read frequencies, like pollen percentages, do not directly reflect plant proportions in the biome (see Supplementary Information: *Betula* comparison). A range of taphonomic factors also influence the biotic representativeness of eDNA, including relative genome size, depositional pathway, extraction protocols, and reference database quality among others.

The pollen sequence was subdivided using the same stratigraphically constrained cluster analysis as applied to the metagenomic data. However, the scarcity of pollen in the marine layers limited the analysis to a depth of ~230 cm, corresponding to ca. 600 CE. Pollen accumulation rates range from 11,949 grains $cm^{-2}$ $yr^{-1}$ at the top of the sequence to 74 at the base (mean = 1,903). Pollen accumulation rates are low compared with forested boreal sites at the same latitude but typical for Iceland, where cool summers restrict pollen production. The first pollen zone corresponds to Zone T-2 of the terrestrial Viridiplantae diagram, with P-2 corresponding to the settlement period zone (T-3), P-3 and P-4 to the retraction period, and P-5 to the 20th century. These diagrams are referenced and compared in the sections that follow.

**Metagenomic record of aquatic plants**

The lake's vascular flora, directly observed through macro- and microfossils, aligns well with the aquatic angiosperms identified in the metagenomic analysis. This agreement is weaker for algae and bryophytes. For instance, oogonia of both *Chara* and *Nitella* were found in the sediments, yet neither of these macroalga was detected in the metagenomic data. A similar discrepancy was observed for *Drepanocladus*, an aquatic bryophyte. While the failure to detect these genera could stem from limitation in the reference database, where algal and bryophyte genomes are underrepresented relative to those of vascular plants, that is not the case here because complete chloroplast genomes for these genera were represented in the database. This suggests that differential preservation and other taphonomic factors limit their detection. Substantial disparities were also evident in the abundance of reads assigned to different taxonomic groups, reflecting habitat differences. On average, reads linked to aquatic angiosperms were roughly ten times more abundant than those linked to terrestrial plants, and about three orders of magnitude more numerous than those matched to algae or metazoans. Read counts also varied considerably between samples, making the sample variance and standard error of the aquatic vascular plants anomalously large. All aquatic taxa were highly skewed and kurtotic relative to the terrestrial. This pattern is unlikely to be explained solely by nucleotide preservation, as reads from vascular plants, animals, and algae exhibited distinct stratigraphic distributions. Terrestrial and aquatic angiosperms were strongly correlated (Pearson's $r = 0.938$, $n = 62$, $P = 0.001$), whereas metazoans and algae showed only weak negative correlations with vascular plants and no significant correlation with each other.

**Extended Data Fig. 2** shows the aquatic viridiplantae, including fully aquatic vascular plants and chlorophytes. Several taxa that grow in shallow water or on permanently saturated substrates are also included in this diagram, plotted to the left of the aquatic angiosperms for comparison. These same taxa are also represented in the terrestrial viridiplantae diagram (**Extended Data Fig. 4**), where their percentages were calculated relative to the total terrestrial read sum, rather than the much larger aquatic read sum used for aquatic angiosperms. Only aquatic plants and algae were included in the calculation of biozonation and diversity statistics in **Extended Data Fig. 3**, whereas the shallow-water taxa were included in both the plotting and statistical analyses in **Extended Data Fig. 4**.

Despite the high abundance of reads from aquatic angiosperms, only ten genera were identified: *Lemna*, *Potamogeton*, *Stuckenia*, *Zannichellia*, and *Zostera* (Alismatales); *Ceratophyllum*; *Elatine*; *Myriophyllum*; *Callitriche*; and *Utricularia*. On average, 96.4% of the reads matched *Myriophyllum*, despite its relatively small genome size (~244 Mb[38]) Samples with lower proportions of Myriophyllum consistently contained markedly fewer reads from aquatic angiosperms overall. This suggests that, for most of the record, this hydrophyte dominated the aquatic community and contributed the majority of detrital biomass.

The *Myriophyllum* species native to Iceland and still present in the lake today is *M. alterniflorum*, a submerged hydrophyte intolerant of brackish water, rather than an emergent form. It dominates most of the sequence, except in Zone A-1 at the base, Zone A-3 (900–1100 CE), and Zone A-5 at the top (**Extended Data Fig. 3**). Shifts in the dominance of *Myriophyllum* relative to its competitors represent the principal ecological dynamic in the lake.

Algal reads are relatively abundant at the base of the sequence, dominated by the marine genera *Ulvaria*, *Ulva*, and *Pycnococcus*. In contrast, reads matching aquatic angiosperms are rare in this zone and are largely represented by *Zostera*, a marine seagrass. The first major ecological shift began around 300 CE, when algal reads from these marine taxa declined while *Myriophyllum* increased at the expense of *Zostera*. However, the overall aquatic angiosperm read count did not rise until after 600 CE, likely reflecting the accumulation of *Myriophyllum* biomass in the lake. By 660 CE, all marine taxa had disappeared—consistent with isolation from the sea—and were replaced in Zone A-2 by a freshwater hydrophyte community strongly dominated by *Myriophyllum*, with little evidence of planktonic algae (**Extended Data Fig. 3**). The presence of *Myriophyllum* reads in Zone A-1, alongside marine taxa, indicates that it was already well established in the upstream freshwater catchment and rapidly colonized the lake's photic zone as the basin freshened.

The dominant alga in Zone A-2 is *Spirogyra*, a freshwater macroalga that thrives under eutrophic conditions. Its maximum frequency occurs just after 800 CE, indicating that the lake was eutrophic in its freshwater phase prior to the Landnám. The transition to Zone A-3 at 900 CE therefore falls shortly after the onset of Landnám. Zone A-3 is characterized by a century-long rise in the frequencies of *Potamogeton* and *Stuckenia* at the expense of *Myriophyllum*, which began to decline around 960 CE. By 1080 CE, *Myriophyllum* accounted for only 20% of hydrophyte reads but had fully recovered by 1180 CE. However, the proportion of Myriophyllum reads rose more rapidly than total read abundance, suggesting that biomass recovery may have taken over a century longer.

Because of its finely divided leaves and submerged growth form, *Myriophyllum* is highly sensitive to turbidity in the photic zone, which restricts it to shallower water. In contrast, emergent

hydrophytes with floating leaves or mat-forming growth have a competitive advantage under turbid conditions, as they dominate the surface layer and shade deeper-growing taxa. Lakes in southwestern Iceland today host several *Potamogeton* species. Some are submerged (*P. perfoliatus*, *P. berchtoldii*, *P. alpinus*), while others (*P. gramineus, P. natans*) bear floating leaves. The most common *Stuckenia* species, *S. filiformis*, also produces floating leaves and typically dominates the upper photic zone. These traits would have given Potamogeton and Stuckenia a competitive edge over *Myriophyllum* if turbidity increased during Zone A-3 (**Extended Data Fig. 3**).
Zone A-3 begins shortly after Landnám, so anthropogenic disturbance of the catchment (e.g., cultivation and grazing) could have increased both nutrient and mineral sediment influx, raising turbidity. However, enhanced nutrient input from livestock and effluvium may not have been a major factor throughout the period, as planktonic algal reads are scarce in this zone and *Spirogyra* declines, consistent with less eutrophic conditions. Of the planktonic genera detected *Auxenochlorella* and *Chlorella* dominate. Their percentage distributions form jagged alternating peaks suggestive of intermittent monospecific blooms possibly indicative of sporadic nutrient influx. Turbidity in the photic zone could have been due to these intermittent algal blooms and suspended eroded sediment.
Multiple proxies—including titanium influx, density, magnetic susceptibility, and LOI—all point to increased minerogenic influx during this period (Main Fig. 1, 5; XRF section). Furthermore, episodes of accelerated sediment deposition would have favoured strongly rooted hydrophytes such as *Potamogeton* and *Stuckenia*, which obtain nutrients directly from sediments, while simultaneously impeding *Myriophyllum* germination and recruitment[39,40,41].
Although soil erosion in the lake's catchment may have contributed to the decline of *Myriophyllum* in Zone A-3, not all features of the record align with this explanation. Like *Myriophyllum*, the freshwater macroalgae *Chara* and *Nitella* are sensitive to a darkened photic zone. Yet their distinctive oospores persist in the sequence up to 140 cm depth, corresponding to ca. 1230 CE (the base of Zone A-4; **Fig. S2**). This suggests that, at least periodically, the photic zone remained clear enough to support these submerged hydrophytes.

Shifts in the presence and frequency of reads from shallow-water rooted hydrophytes and telmatic plants point to an additional dynamic. In the pre-settlement period, *Caltha* (marsh-marigold) was the only genus adapted to saturated substrates represented in the record. In Zone A-3, however, *Glyceria*, *Eleocharis*, *Juncus*, and *Luzula* all appear and increase. Given Tjörnin's shallow bathymetry, even a modest rise in lake level would have expanded habitat for rooted hydrophytes such as *Potamogeton* and *Stuckenia*, as well as the telmatic zone and surrounding wet meadow. Intentional modification—such as a partial impoundment of the Lækurinn outlet—may be the most parsimonious explanation for these settlement-period rise in lake-level.

A distinguishing feature of Zone A-4 is the persistence of telmatic taxa and hydrophytes that were rare or absent in Zones A-1 and A-2. Notably, *Isoëtes* read percentages peak in early Zone A-4, following the recovery of *Myriophyllum*. The genus is also recorded in the pollen study, where *I. echinospora* microspores comprise up to 70% of pollen and spores in Pollen Zone P-3 (**Extended Data Fig. 3, 5**). This indicates that *Isoëtes* was relatively common in the lake at the time. As a small rooted, submerged hydrophyte, *Isoëtes* would have benefited from new habitat created by the rising lake-level in Zone A-3 but remained limited to shallow water under turbid conditions, avoiding high-sedimentation areas. Its abundance in Zone A-4 suggests that minerogenic influx diminished, consistent with reduced cultivation and grazing in the catchment. These conditions facilitated both the spread of *I. echinospora* and the recovery of *Myriophyllum* in the Retraction Episode.

Discrepancies between the Lake Tjörnin pollen and metagenomic records provide further insight into *Myriophyllum*. Between 570 and 960 CE—when it likely constituted the bulk of detrital biomass—*Myriophyllum* pollen accounted for 2–44% of total pollen (mean = 15.7%). Values collapsed below 2% around 960 CE and remained low until ca. 1260 CE, when a gradual recovery began. From 1645 to 1900 CE, pollen percentages were robust (mean = 27.5%) before falling below 3% in the 20$^{th}$ century. While both pollen and read counts declined sharply around 960 CE, *Myriophyllum* reads recovered more quickly than pollen, suggesting that clonal reproduction may have played a role in biomass recovery.

Algal reads are particularly scarce in sediments from Zones A-3 and A-4 (900–1900 CE; **Extended Data Fig. 3**). The gyttja in Zone A-4 is detrital, composed mainly of decomposed aquatic macrophytes, chiefly *Myriophyllum* (**Fig. S2**). Both algal read counts and taxonomic richness increase in the fine, gelatinous gyttja deposited during the 20$^{th}$ century. In Zone A-5, several planktonic genera appear simultaneously for the first time in the record: *Pseudopediastrum*, *Tetradesmus*, *Desmodesmus*, *Balticola*, and *Stephanosphaera*. These taxa emerge at or just after the second decline of *Myriophyllum* (ca. 1850 CE), coinciding with the expansion of floating and mat-forming hydrophytes such as *Lemna* and *Zannichellia*. *Ceratophyllum* and *Elatine* also appear largely in this period, though at low frequencies, consistent with recent introductions. *Stuckenia* and *Potamogeton* frequencies also rise in Zone A-5, but unlike in Zone A-3, their dynamics are asynchronous, suggesting a more complex community structure.

Taken together, these data converge on a single explanation for the disappearance of *Isoëtes* and the second decline of *Myriophyllum*: effluent from the expanding city of Reykjavík fertilized planktonic algae, which darkened the photic zone and favoured surface-dominant hydrophytes. Ironically, the suppression of *Myriophyllum* under increasing eutrophication resulted in a more diverse hydrophyte community.

**Metagenomic record of terrestrial plants**

The metagenomic analysis identified 275 terrestrial and telmatic vascular plant genera after filtering (see Methods), of which 241 (88%) occur in the modern Iceland flora either as native species or historic introductions (Extended Data Fig. 4). Of these genera, 18 include at least one introduced species, either a cultivar or anthropochore. The remainder are plants that either cannot grow in Iceland or were never introduced. Most occur at trace frequencies and may be misidentified due to the issues with reference databases described above. However, some of these exotic genera including *Camellia* (tea), *Theobroma* (cacao), and *Acer* (maple) may be correctly matched as they

are confined to the top of the sequence. Reads matching *Acer* may come from trees planted as ornamentals in the city, while nucleic acids from tea and chocolate could come into the sediment from effluvium.
Of the eleven trees identified in the metagenomic analysis, only four were present in the pre-Landnám landscape: *Salix* (willow), *Betula* (birch), *Sorbus* (rowan) and *Populus* (poplar). Arboreal percentages were generally low and of these only *Salix* had values comparable to some of the grasses and forbes, ranging from 24 to 0.2% (Mean = 5%). The abundance of reads matched to non-native trees *Alnus* (alder), *Carpinus* (hornbeam), *Corylus* (hazel) and *Juglans* (walnut) was lower still. Percentages of *Corylus* increase in Zone T-3 which could be construed as evidence for a settlement period introduction that later failed, but this is not supported by pollen or macrofossils (Ext. Fig. 4). Many sequences mapped to non-native trees may simply be mis-mapped from related native taxa. The read percentages of non-native members of the Betulaceae, *Corylus*, *Alnus* and *Carpinus* are strongly correlated with *Betula* (Pearson's r = .982, .961 and .982 respectively, n= 62, P=.001). Therefore, reads assigned to allochthonous taxa that are strongly correlated with closely related native genera should be ignored in this study. This is particularly relevant when they are represented by so few reads. Some of these genera are also registered in the pollen study at low frequencies. Most of these are wind dispersed taxa, so the presence of these exotic taxa in the pollen record is probably due to long-distance transport.

Several closely related Rosaceous genera were detected including *Sorbus*, the genus that includes the native tree *S. aucuparia* (rowan), along with *Hedlundia* a hybrid genus derived from *S. aucuparia* and *Aria* (whitebeam) along with the cultivated genera *Malus* (apple) and *Pyrus* (pear). These very closely related genera form a tribe within the Amygdaloideae sub-family of the Rosaceae, known as the Maleae. However, these genera were not nearly as strongly correlated with each other in the sequence as the related genera in the Betulaceae described above. Therefore, we have less confidence that these are *Sorbus* nucleotides overmatched to other genera in the Maleae. To directly test whether these reads were overmatched to these exotic taxa from the native genus, *Sorbus* we conducted additional tests on limitations in the reference genomes in our database for the family Rosaceae. We extracted all reads assigned to the species within *Malus* and *Pyrus* from the Rosaceae family and then remapped against all available Rosaceae genomes from RefSeq with a minimum reference length of 100.000 bp (NCBI search string:
"("Rosaceae"[Organism] OR Rosaceae [All Fields]) AND (refseq[filter] AND ("100000"[SLEN] : "1000000000000"[SLEN]))")).
We collected a total of 2487 reference genomes from RefSeq release 229 from the 03.03.2025. Remapping against the custom Rosaceae database was done using bowtie2, and taxonomic assignment was made using metaDMG with the same parameters used in the metagenomic mapping (see Methods).
Because the NCBI repository grows constantly, new genomes could represent a broader spectrum of the diversity within families than our original database. Comparing the newly assigned LCA ranks using our custom Rosaceae database compared to the LCA assignments of the original, we found, that nearly all reads now fell at a higher taxonomic level, primarily within the tribe Maleae for Rosaceae (see Fig. S39). Some of the overmatching in our original mapping can be explained by the

lack of references for the native Iceland flora as well as reads mapping to conserved regions within the Rosaceae. Given this, there is no strong reason to suggest that the increase in reads mapping to *Malus* in the Settlement Episode is evidence of a failed introduction of this genus. However, the highest percentages of reads mapping to members of the Maleae other than *Sorbus* occur at the top of the record dating to the late 20th century, when at least whitebeam if not the others, could have been planted for urban landscaping. This, coupled with the fact that variance in the percentages of *Sorbus* and the other Maleae were not strongly correlated through the record justified collapsing the reads from genera in the Maleae tribe other than *Sorbus* (i.e. *Hedlundia*, *Malus*, and *Pyrus*) into a single category plotted alongside *Sorbus* in the diagram, rather than simply adding them to the *Sorbus* category on the assumption that they were all overmatched.

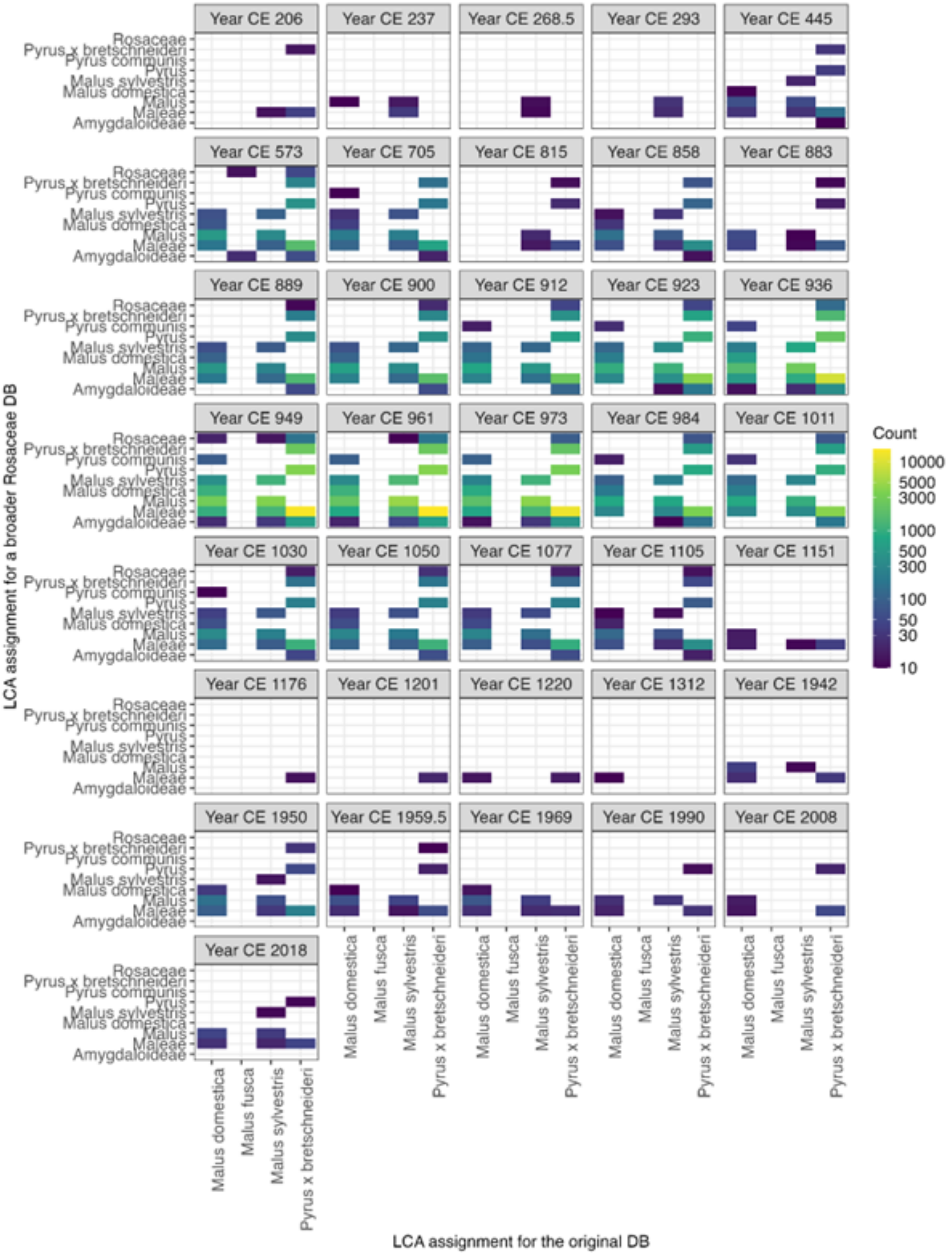

**Figure S39. LCA assignment comparison for all reads classified to the genera Malus and Pyrus.** The x-axis represents the original LCA assignment, while the y-axis shows new LCA assignments when competitive mapping against a broader database consisting of all references from RefSeq for the family Rosaceae. The count describes how many reads have been switched to an alternative LCA assignment.

The non-native genus *Prunus* which includes plumbs and cherries was also detected. Although *Prunus* shares the same Rosaceae sub-family the Amygdaloideae, with the Maleae tribe, it belongs to a separate tribe the Amygdaleae. Although it is certainly possible that reads matched to *Prunus* could also have been overmatched from *Sorbus*, this is somewhat less likely given its taxonomy. A *P. domestica* pyrene was recovered at the Lækjargata 10–12 site on the north bank of the Tjörnin dating to the settlement period[22] and several members of the genus were utilized elsewhere in Scandinavia during the early medieval period, including *P. spinosa* (Sloe), *P. domestica* (also subsp. *Insititia*) (plums and damsons) as well as *P. avium* (cherries)[42]. Therefore, members of this genus were very familiar to the Norse and were both imported to and utilized at the site leading to speculation that limited *prunus* cultivation was attempted in the Tjörnin catchment during the settlement period. This is probably unwarranted as reads matched to *Prunus* are very rare in the record and are not concentrated in layers dating to the settlement. Therefore, there is no incontrovertible evidence in the eDNA that exotic rosaceous trees were introduced as cultivars in the settlement period.

*Picea* (spruce) is the only arboreal taxa detected that is successfully naturalized today but its reads occur at only trace proportions throughout the record and there is no increase corresponding to its late introduction.

The 275 genera plotted in Extended Data Fig. 5 are ordered by habitat except pteridophytes and non-native plants. The category comprised of ruderal and cultivated plants typical of hay meadows in Iceland now, is the largest group with 47 genera (17%). Of these, ten genera were classified as introduced cultivars because the genera are not native to Iceland and have cultivated species. A second category of grasses included genera that may have occurred in the pre-Landnám flora but which also have introduced species. *Hordeum* (barley) has its highest frequencies during the settlement period (Zone T-3) where it comprises as much as 17% of the total reads. There is historical and archaeological evidence for barley cultivation in Tjörnin catchment[43]. The relatively high frequencies during Zone T-3 suggest that it was cultivated from 900 to 1200 CE followed by a sharp decline in the 13$^{th}$ century. In contrast, reads from *Avena* (oats) and *Triticum* (wheat) do not have peak percentages in this zone. *Triticum* is a diverse genus with both tetraploid and hexaploid species with multiple references of various quality, increasing the chance of overmatching from other grasses. *Triticum* reads matched at the bottom of the sequence attest to this problem. There is no historic evidence for *Triticum* or *Avena* cultivation in the settlement period or afterwards, but macrobotanical evidence suggests that oats may have occurred as a weed in barley fields[43].

In addition to these introduced cultivars, seven other non-native Poaceae genera were also detected: *Aegilops*, *Brachypodium*, *Bromus*, *Helictochloa*, *Hordelymus*, *Koeleria*, and *Phalaris*. None of these are naturalized and most have never been reported in Iceland (Table S4). To test whether these identifications were also due to overmatching from related native taxa we repeated our overmatching analysis from the Rosaceae family with the Poaceae to investigate if some of the potentially misassignments could have been due to a lack of native reference genomes in our original database. The newly collected database for the Poaceae consisted of 4330 reference genomes, collected from the RefSeq release 229, 03.03.2025 with the search string:

“("Poaceae"[Organism] OR Poaceae[All Fields]) AND (refseq[filter] AND ("100000"[SLEN] : "1000000000000"[SLEN]))”.

All mapping and taxonomic assignment parameters remained consistent with those described in the method section. The reads from our metagenomic mapping for the genera *Aegilops*, *Avena*, *Brachypodium*, *Bromus*, *Helictochloa*, *Hordelymus*, *Koeleria*, *Phalaris*, and *Triticum* were extracted and remapped. We were able to observe again a higher proportion of reads were placed at a higher taxonomic level with the update reference database (Fig. S40).

Other anthropochors or cultivars including *Brassica* (possibly kale/cabbage), *Trifolium* (clover) and *Cota/Anthemis* (possibly dyer's chamomile) have distributions consistent with an introduction at or shortly after the Landnám. Reads matched to Solanum are probably derived from the potatoes, which became a staple of the Icelandic diet in the 18$^{th}$ century. These only appear near the top of the sequence.

Supplementary Data Table S4 lists genera in the Poaceae registered in the metagenomic analysis along with their habitats and enumerates the species in each genus. Non-native species are listed and the date of introduction estimated. An increase in reads from these genera could be driven by their introduction and naturalization of the latter or the niche expansion of native species as conditions changed.

**Table S4. Lists genera in the Poaceae that may have occurred in the pre-Landnám flora, but which also have introduced species (based mainly on Kristinsson and colleagues[44]).**

| Poaceae Genera | Habit (Habitat) | Comments |
|---|---|---|
| *Hordeum* | Cultivar | 1 cultivated species. |
| *Avena* | Cultivar | Possibly cultivated historically in Iceland but poorly attested. |
| *Dactylis* | Cultivated | 1 sp. *D. glomerata* was introduced before 1810, prob. for haymaking. Naturalized. |
| *Phleum* | Cultivated(hay)/meadow/ruderal | *P. alpinum* is native, *P. prantense* was probably introduced for haymaking before the mid-18th century. |
| *Festuca* | | 3 species, *F. rubra* introduced ca mid-19th century for haymaking. |
| *Poa* | | 7 species *P. pratensis* ssp. *pratensis* is probably introduced for hay. All are now naturalized. |
| *Deschampsia* | | 3 species *D. beringensis* was introduce ca 1970 for haymaking. *D. glomerata* was introduced before 1810, prob. for haymaking. Both are now naturalized. |
| *Alopecurus* | Cultivated/Pond edge/shallow water/wet meadow/meadow/ruderal | 3 species *A. aequalis* (aquatic) is probably native. *A. geniculatus* (wet meadow/meadow/ruderal) is prob. introduced before the mid-18th century. *A. pratensis* was introduced ca 1900 for haymaking. All are now naturalized. |
| *Lolium* | Meadow/Ruderal | 1 species. Introduced ca 1900. Naturalized. |
| *Elymus* | | 2 species. *E. repens* is considered imported early in the post-settlement period. |
| *Agrostis* | | All 5 species are probably native. |
| *Avenella* | | 1 species, *A. flexuosa* is native. |
| *Anthoxanthum* | | 1 species, widly distributed. |
| *Calamagrostis* | Wet Meadow | 1 species |
| *Glyceria* | Pond edge/shallow water | Fine. 1 species (known to be harvested for its seeds). Very restricted distribution in Iceland (not in RVK). Possibly introduced. |
| *Catabrosa* | Pond edge/shallow water | 1 species |
| *Trisetum/Koeleria* | Rocky substrates and meadows | 2 species |
| *Arrhenatherum* | Rocky substrates | 1 species, endangered, grows in only two place in Iceland. |
| *Phippsia* | Wet meadow/rocky substrates. | 1 species, *P. algida* grows in wet, sparsely vegetated and gravelly substrates, mainly in the highlands. |
| *Puccinellia* | Sea Coast | 2 species |
| *Leymus* | | 1 species |
| *Milium* | Survives in remnant woodland and places providing natural protection | 1 species, *Milium effusum*. Probably native. |
| *Aegilops* | Non-Native/Not naturalized | |
| *Brachypodium* | | |
| *Helictochloa* | | |
| *Hordelymus* | | |
| *Phalaris* | | Not native, but *P. canariensis* was found in Reykjavík, Húsavík and Sauðárkrókur between 1893 and 1966. |
| *Bromus* | | Four species of *Bromus* have been recorded, none have naturalized. |

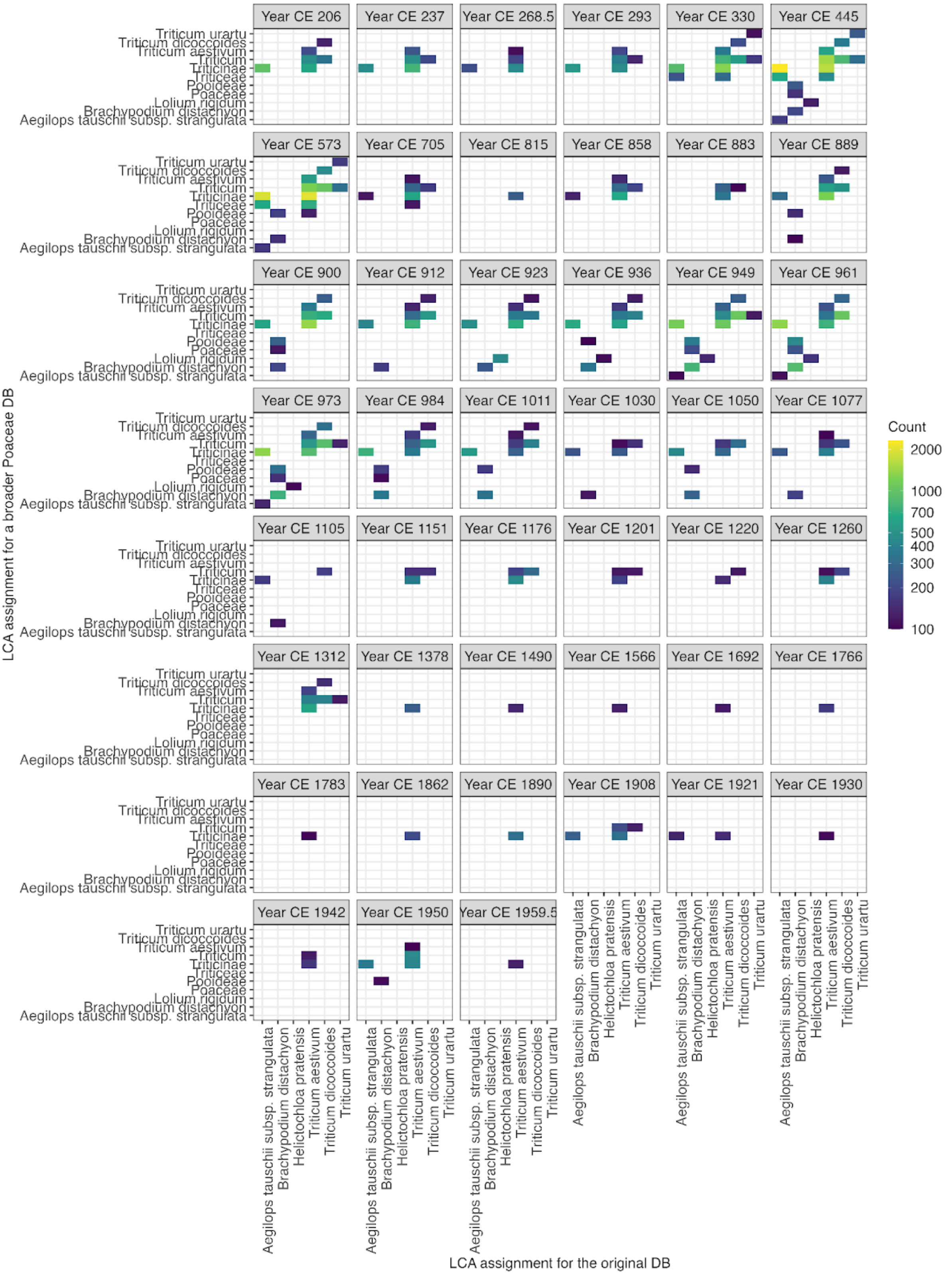

**Figure S40. LCA assignment comparison for all reads classified to the genera *Aegilops, Avena, Brachypodium, Bromus, Helictochloa, Hordelymus, Koeleria, Phalaris, Triticum*.** The x-axis represents the original LCA assignment, while the y-axis shows new LCA assignments when competitive mapping against a broader database consisting of all references from RefSeq for the family Poaceae. The count describes how many reads have been switched to an alternative LCA assignment. All assignments with fewer than 100 counts are filtered out for clarification.

Though most genera comprise only a small percentage of reads throughout the sequence, several are very well represented. The coast adapted grasses *Puccinellia* and *Leymus* dominate the base of the sequence (Zone T-1) along with *Honckenya* (sea sandwort). These all decline sharply and then disappear in Zone T-2 as the Tjörnin is isolated. The dominance of *Carex* (sedges) throughout the record is also expected, as some species in this genus comprise much of the biomass in all low elevation meadows in Iceland. In contrast, the high frequencies of *Phegopteris* (beech ferns) are difficult to explain. In Iceland, ferns in this genus grow in wet meadows and along rock outcrops, cave mouths and cliffs, but is not a common plant. While the number of reads assigned to the *Carex* may be commensurate with its proportion of the vegetation, this cannot be true of *Phegopteris* or *Drosera* (sundews). The last is a small carnivorous plant growing on histic soils, typically raised bogs, habitats that do not dominate the Tjörnin catchment. Therefore, some factor(s) other than biomass proportion must affect sequence abundance. These could include differences in the relative size of genomes between all plant references used as well as genomic coverage in the databases. *Phegopteris* increases in frequency from 0 to 2% in zone T-1 to well over 40% in T-2 the pre-settlement. Its proportions drop to well under 10% in the settlement phase, only to recover to over 40% in the Retraction Episode. *Carex* shows a similar pattern, rising from low proportions at the base of the record to over 40% in samples in Zone T-2 only to decline during Zone T-3. Like *Phegopteris* it recovers in T-4, albeit not as sharply. Both of these taxa declined as willow, birch and range of meadow grasses and anthropochores increased along with barley. *Carex* and *Phegopteris* recovered as managed hay meadows contracted, and cultivation ceased in the catchment, consistent with the Episode III retraction of cultivation. Both plants declined again as the catchment was paved in the 20$^{th}$ century.

**Comparison of palynological and metagenomic records**

The pollen analysis identified a total of 71 taxa to different taxonomic levels (Extended Data Fig 5). Sixteen of the 62 genera could be provisionally identified to species, twelve to family, one to tribe, one to class and two to subfamilies. 18 (29%) of the genera identified in the pollen analysis were not detected in the metagenomic analysis at the level of filtering used. These included: *Juniperus* (juniper), a common arboreal plant in the Tjörnin catchment, *Lupinus* (lupins) a recent introduction from North America, *Artemisia* (wormwood), and *Urtica* (nettles) important ruderal and invasive plants. Pollen from the telematic plant *Sparganium* (bur-reed) along with spores from the native pteridophytes, *Selaginella*, *Polypodium* and *Sphagnum* along with *Calluna* (heather) was also only identified in the pollen study. However, the related genera, *Vaccinium* (whortleberry/bilberry) and *Empetrum* (crowberry) were detected in the metagenomic analysis. Most of these taxa are present in the PhyloNorway database[33] so they may have been detected at levels below the filtering threshold. The Cyperaceae family, which includes *Carex* was the most abundant pollen type followed by the Poaceae which unlike the Cyperaceae, are represented by a range of genera in the metagenomic analysis, these include many from meadow and ruderal habitats as well as the coastal grasses *Puccinellia* and *Leymus* and some others. Because the various grass genera subsumed in the

Poaceae family are adapted to different habitats this pollen category lacks the ecological resolution of the metagenomic analysis.

Though these same broad taxonomic groups dominated both records, variance in neither Poaceae nor Cyperaceae pollen corresponds to their metagenomic analogues. The reads from several grass genera including *Poa*, *Alopecurus*, *Deschampsia*, *Agrostis* and the cultivar *Hordeum* rise in frequency during the settlement period and decline during the Retraction Episode. In contrast, the percentage of grass pollen increases gradually throughout the record and does not decline at that time. *Carex* in the metagenomic analysis reaches its highest percentages in Zone T-4 during the Retraction Episode, after declining during the settlement period as grasses and anthropochores increased. In contrast, Cyperaceae pollen is well represented throughout the microfossil record including the settlement period. It does decline slightly in biozones P-4 and P-5 corresponding to the Urbanization Episode (IV). The *Carex* decline in the metagenomic data only occurs in Zone T-5 at the top of the sequence (Extended Data Fig. 5).

Pteridophytes comprise the other major taxonomic category in both records, and this is also completely out of phase in the two studies. Polypodiopsida spores comprise between 40 and 50% of all micro sub-fossil in pollen zone P-2 which corresponds to the settlement period (Episode II). In contrast, few pteridophytes reads are detected during this period in the metagenomic analysis. This emphasizes the different taphonomic processes governing the incorporation of microfossils and nucleotide s from the same plants.

We suspect that relatively little nucleic acid is derived from the pollen and spores identified in the sediments because cytoplasm encapsulated in the sporopollenin of intact microfossils is resistant to DNA extraction. Therefore, most of the analysed sequences are from extracellular DNA adhering to minerals or degraded organic matter which either forms in or is washed into the basin, rather than microfossils. The lack of correspondence between nucleotides and microfossils in the most common taxa supports this conjecture.

If this generalization is accurate, it follows that pteridophytes comprised less of the vegetation near the lake during the Settlement Episode despite their relatively high spore frequency which, is difficult to explain. The reduction of nucleotides from ferns during the period is consistent with cultivation and to a lesser extent grazing which would have reduced the proportion of ferns in the vegetation directly adjacent to the lake while, ferns further from the lake in uncultivated or generally less impacted areas could have continued to produce copious wind dispersed spores. However, this does not explain why Polypodiopsida spores are so abundant in the settlement Zone P-2.

Total pollen accumulation is higher in the Settlement Episode when minerogenic influx also increases probably due to increased erosion. If the Urbanization Episode (which has anomalously high pollen influx) is excluded from the correlation analysis, spores of Pterospida and *Sphagnum* are correlated with pollen accumulation through the sequence (Pearson's $r = 0.58$, $n = 53$, $P = 0.001$). This suggests that spores of various pteridophytes and bryophytes may be mobilized from the surface and redeposited in the lake with eroded sediments.

In contrast, microspores of *Isoëtes echinospora* and *Myriophyllum* pollen comprise larger percentages of samples accumulated when total pollen influx was low. At least two factors could contribute to this: microfossils from submerged aquatic plants are produced in the lake and are not subject to variation in the factors controlling influx like surface erosion, and the fact that neither of these aquatic plants thrive when sediment influx is high.

*I. echinospora* is the most abundant spore in pollen zone P-3 and its correspondence with *Isoëtes* read abundance in the metagenomic record has already been noted. This correspondence supports the interpretation of an expanded telmatic habitat (possible due to a lake-level rise flooding the lake shore) and reduced sedimentation during the Retraction Episode (III). *Myriophyllum* pollen and read abundance correspond to a lesser degree. The *Myriophyllum* declines in the metagenomic record are confined to short periods in the settlement period (Episode II) and during the $20^{th}$ century. *M. alterniflorum* pollen declines sharply in the Settlement Episode (II) Zone P-2 but recovers only gradually during the Retraction Episode. The highest percentages of *M. alterniflorum* pollen fall in pollen biozone P-4 corresponding to the early Urbanization Episode (IV). *M. alterniflorum* pollen disappears in the $20^{th}$ century along with *Myriophyllum* nucleotides. Here, discrepancies between the eDNA and pollen records show that *Myriophyllum* probably comprised much of the lakes biomass even when it produced relatively little pollen, suggesting clonal rather than sexual reproduction during long periods. This is particularly evident between the $11^{th}$ to the $17^{th}$ centuries.

Both proxies also concur regarding the native arboreal taxa *Betula* and *Salix*. Pollen influx from both is very low for most of the record except for late Episode IV, where it is probably derived from tended trees planted in yards and public parks. The slight *Betula* pollen percentage decline recorded at the top of Zone P-1 corresponding to the Settlement Episode is due to constraint. Actual *Betula* pollen accumulation increases slightly at this time but was always very low prior to urbanization. Suggesting that these trees and/or shrubs in the natural flora produced very little pollen, relying on clonal reproduction, and/or never comprised a large proportion of the vegetation relative to the grasses, sedges and ferns, even during the Pre-human Episode. The four to tenfold increase in nucleotide reads matched to *Betula* and *Salix* respectively, persisted through the first three centuries of the Settlement Episode, when grazing pressure was most intense. Though browsing livestock may have moved biomass from these plants into the lake via their dung, both the pollen and metagenomic records are consistent with an expansion of birch and willow however slight, rather than extirpation or reduction. Not surprisingly, reads from *Geranium*, in Iceland a plant associated with the birch understory, also increase during the settlement period. It seems unlikely that this habitat could persist along with livestock if not protected, perhaps in fenced off coppices of birch and willow.

Finally, the microcharcoal influx recorded in the pollen record indicates that natural fire was very rare in the pre-settlement period. Charcoal influx rises in the Settlement Episode. It then falls in pollen zone P-3 corresponding to the Retraction Episode III but there are two sharp spikes centred on the mid-$19^{th}$ and $20^{th}$ centuries respectively (Episode IV). The reduction in charcoal influx in the Retraction Episode further confirms the contraction or reduction of settlement during this period in the catchment. Its gradual resumption after 1700 (biozone P-4) is consistent with the growth of the urban settlement and later, industries (Extended Data Fig. 5).

**Metazoan metagenomic record**

Extended Data Figure 2 shows the relative proportions of all filtered reads (mean read length ca. 45 base pairs) matched to metazoan genera in the Tjörnin sequence, plotted against the modelled calendar-year age. Genera are ordered primarily by habitat and secondarily by taxonomy. The sequence is divided into five biozones, defined by stratigraphically constrained cluster analysis. Supplementary Data Table S5, lists the class, order, and common English names of each metazoan genus, together with the total mapped reads from all samples.

Although metazoan reads are roughly two orders of magnitude less abundant than those of terrestrial plants, reads assigned to *Anas* (dabbling ducks), *Ovis* (sheep), and *Bos* (cattle) are about an order of magnitude more common than those from other terrestrial animals—likely because dung from waterfowl and livestock feeding on or near the lake was incorporated directly into the sediment. Marine reads are dominated by fish, *Halocynthia* (sea squirts), and *Mytilus* (mussels), while freshwater reads are dominated by two ostracod genera. The highest read concentrations occur at the base of the sequence and again in the 10th and 20th centuries (Extended Data Fig. 2). As noted above, the distribution of metazoan reads shows no correlation with aquatic or terrestrial plant reads, consistent with their distinctive sources and taphonomic pathways.

The PhyloNorway genome skim database used in this study includes many vascular plants from Norway and adjacent polar regions, with complete chloroplast and nuclear ribosomal assemblies[33]. In contrast, the metazoan reference set is far less comprehensive and of more uneven quality. Many invertebrate genera likely escape detection because no reference exists, and some taxa are represented only by mitochondrial or chloroplast sequences. Others are included through nuclear genome skims of variable coverage. Consequently, the range of eukaryotic taxa detected reflects, to some extent, the limitations of the reference database, with metazoans more strongly affected than viridiplantae in this study.

**Table S5. Lists the class, order and common English names of each metazoan genus along with their total reads from all samples.** Exotic genera never recorded in Iceland's flora are printed in red while introduced domestic animals are shown in blue.

**Aquatic Genera**

| Habitat | Genus | Read count | Class | Order/Suborder | CommonName |
|---|---|---|---|---|---|
| Fully Marine | *Anguilla* | 3238 | Actinopterygii | Anguilliformes | Eels |
| | *Clupea* | 8961 | Actinopterygii | Clupeiformes | Herrings |
| | *Gadus* | 28325 | Actinopterygii | Gadiformes | Cod |
| | *Sebastes* | 7329 | Actinopterygii | Scorpaeniformes | Rockfish |
| | *Lineus* | 717 | Anopla (underthephylum Nemertea) | Heteronemertea | Ribbon worms |
| | *Halocynthia* | 17306 | Ascidiacea | Stolidobranchia | Sea squirts |
| | *Asterias* | 608 | Asteroidea | Forcipulatida | Starfish |
| | *Aulacomya* | 417 | Bivalvia | Mytilida | Ribbed mussel |
| | *Limnopema* | 235 | Bivalvia | Mytilida | Golden mussel |
| | *Mytilus* | 10879 | Bivalvia | Mytilida | Mussels |
| | *Octopus* | 2131 | Cephalopoda | Octopoda | Octopuses |
| | *Strongylocentrotus* | 1590 | Echinoidea | Echinoida | Sea urchins |
| | *Saccoglossus* | 3746 | Enteropneusta | Enteropneusta | Acorn Worms |
| | *Biomphalaria* | 807 | Gastropoda | Hygrophila | Aquatic gastropod |
| | *Patella* | 1134 | Gastropoda | Patellogastropoda | Limpets |
| | *Membranipora* | 1550 | Gymnolaemata | Cheilostomatida | Bryozoans (encrusting bryozoans) |
| | *Penaeus* | 2048 | Malacostraca | Decapoda | Shrimp |
| | *Portunus* | 4027 | Malacostraca | Decapoda | Swimming crabs |
| | *Bradleycypris* | 396 | Ostracoda | Podocopida | Seed shrimp |
| | *Cyprideis* | 13035 | Ostracoda | Podocopida | Seed Shrimp |
| | *Priapulus* | 4799 | Priapulida | Priapulida | Priapulid Worms |
| Freshwater-Brackish | *Gasterosteus* | 18469 | Actinopterygii | Gasterosteiformes | Sticklebacks |
| | *Brachionus* | 5914 | Rotifera | Ploima | Rotifers |
| | *Keratella* | 1171 | Rotifera | Ploima | Rotifers |
| | *Otomesostoma* | 2170 | Platyhelminthes | Proseriata | Free-living flatworm |
| Primarily Freshwater | *Chironomus* | 24724 | Insecta | Diptera | Non-biting midges |
| | *Gieysztoria* | 11585 | Rhabditophora | Rhabdocoela | Flat worms |
| | *Chrissia* | 21381 | Branchiopoda | Diplostraca | Water Fleas |
| | *Eurycercus* | 418 | Branchiopoda | Diplostraca | Water Fleas |
| | *Plumatella* | 279 | Phylactolaemata | Plumatellida | Bryozoan (freshwater) |
| | *Cyprinotus* | 557 | Ostracoda | Podocopida | Seed Shrimp |
| | *Darwinula* | 10616 | Ostracoda | Podocopida | Seed Shrimp |
| | *Eucypris* | 3583 | Ostracoda | Podocopida | Seed Shrimp |
| | *Heterocypris* | 66612 | Ostracoda | Podocopida | Seed Shrimp |
| | *Notodromas* | 12706 | Ostracoda | Podocopida | Seed Shrimp |
| | *Sarscypridopsis* | 71627 | Ostracoda | Podocopida | Seed Shrimp |
| | *Hydra* | 6228 | Hydrozoa | Anthoathecata | Hydra |

**Terrestrial Genera**

| Habitat | Genus | Read count | Class | Order/Suborder | Common Name |
|---|---|---|---|---|---|
| Terrestrial | *Anas* | 146653 | Aves | Anseriformes | Ducks |
| | *Anser* | 8046 | Aves | Anseriformes | Geese |
| | *Aythya* | 7692 | Aves | Anseriformes | Diving ducks |
| | *Bos* | 24625 | Mammalia | Artiodactyla | Cattle |
| | *Ovis* | 69504 | Mammalia | Artiodactyla | Sheep |
| | *Sus* | 14097 | Mammalia | Artiodactyla | Pigs |
| | *Equus* | 4799 | Mammalia | Perissodactyla | Horses |
| | *Homo* | 17518 | Mammalia | Primates | Humans |
| | *Cricetulus* | 7037 | Mammalia | Rodentia | Hamsters |
| | *Dromiciops* | 416 | Mammalia | Microbiotheria | Colocolo opossum |
| | *Contarinia* | 5146 | Insecta | Diptera | Gall Midges |
| | *Drosophila* | 1226 | Insecta | Diptera | Fruit flies |
| | *Timema* | 1204 | Insecta | Phasmatodea | Stick insects (or walking sticks) |
| | *Lumbricus* | 5651 | Clitellata | Opisthopora | Earth worms |

Several genera detected in the metagenome are also preserved as macrofossils. DNA from chironomids is present throughout the sequence, as are their head capsules (see Supplementary Section: Chironomid Analysis). Although oribatid mites are not recorded, DNA from several genera of Branchiopoda and one genus of Phylactolaemata (Plumatella) is detected, consistent with macrofossil evidence in the sediment log.

Six of the 37 aquatic genera identified do not occur in Iceland. Some of these marine invertebrates likely represent overmatched sequences to reference genomes and instead derive from related allochthonous genera. Normally, sequences shared between genera from conserved regions of the genome would only resolve to higher taxonomic levels (e.g., family or order). However, overmatching can occur when reference genomes from non-local taxa are of higher quality or more numerous than those of native taxa. For example, most bivalve reads match the local genus *Mytilus*, but others align with *Aulacomya* and *Limnoperna*, Pacific genera within Mytilida. A similar pattern is observed in gastropods: the native tortoise limpet *Testudinalia testudinalis* was not detected, but reads were matched to the related non-native genus *Patella*. Marine arthropods may also be overmatched when the reference database lacks local genera, leading to assignments to Pacific taxa in the same orders (e.g., Decapoda and Podocopida).

By contrast, some taxa cannot be explained by overmatching to close relatives. Reads assigned to the insect genus *Timema* (stick insects) are unlikely to be correct, as no members of this order (Phasmatodea) are native to Iceland. Their presence may reflect a poorly sequenced or contaminated genome. The same explanation may apply to the few reads matched to *Dromiciops* (the Colocolo opossum), a genus only distantly related to any mammal known from Iceland, in the past or present.

Although taxa with demonstrably contaminated genomes were removed, all other genera meeting the filtering criteria are retained and plotted, including exotic taxa unlikely to be mapped correctly. Retaining these assignments is preferable to arbitrary removal and highlights the importance of genome quality and the limitations of the metagenomic reference protocol, which remains under continuous refinement.

The only other exotic mammal present in the record at this level of filtering is *Cricetulus* (hamsters). Reads assigned to this genus are more numerous than those of *Dromiciops* or *Homo* and increase after the settlement period. However, they are unlikely to represent overmatching to another rodent genus. No rodents are native to Iceland, and *Mus musculus* (the Eurasian house mouse)—introduced at Landnám—is present in the reference database. It is therefore improbable that the *Cricetulus* reference genome would attract reads from *M. musculus* or similar species, nor is it likely that any rodent biomass would yield so many reads. A more plausible explanation is contamination of the *Cricetulus* genome with incorporated microbial sequences, which are erroneously mapping to bacterial DNA present in the sediments. This phenomenon has been reported previously, but robust and scalable computational methods for the detection and decontamination of large reference genomes are still lacking[45].

A similar issue almost certainly underlies the abundant reads mapped to the genus *Phasianus*. These account for more than 20% of metazoan reads in some samples and occur throughout the sequence. Because *Phasianus* (pheasants) are not native to Iceland and no breeding populations have ever been established through introduction, it is impossible that these reads derive from true pheasants. Overmatching from related genera is also implausible. *Lagopus muta* (rock ptarmigan) is the only native member of the Phasianidae in Iceland. *Gallus gallus domesticus* (the domestic chicken) is first attested archaeologically in Reykjavík in the mid-17$^{th}$ century (see Supplementary section: Historical, Archaeological, and Demographic Context). Both species have high-quality reference genomes available, yet neither is detected in this analysis. Instead, the *Phasianus* reference genome is suspected—although not known—to be heavily contaminated with microbial sequences. Accordingly, reads mapping to *Phasianus*, along with those mapping to other contaminated genomes (e.g., *Oncorhynchus* and *Lipotes*), were excluded from the analysis[45].

Reads matched to Homo (humans) also occur at trace frequencies throughout the sequence, except for a single sample from ca. 1700 CE. More human nucleotides might be expected given the volume of effluvium known to have been released into the basin during the Urbanization Episode. The metazoan bio-zones correspond closely with those generated from the aquatic plants. Reads from marine invertebrates dominate the base of the Tjörnin sequence from ca. 200 to 650 CE (Metazoan Zone M-1, Extended Data Fig. 2). The same segment is dominated by marine algae and Zostera (Zone A-1, Ext. Fig 2). *Halocynthia*, *Mytilus*, *Portunus* (crabs) and *Octopus* dominate the base of the sequence. Reads matching to *Contarinia*, a terrestrial gall midge would be derived from the terrestrial component of the mixed detritus that accumulated when the Tjörnin was a lagoon (Fig. S2). The ostracod *Notodromas* succeeds *Cyprideis* as the lagoon shifts from marine to brackish. The appearance followed by an increase in the read frequency of non-biting midges (Chironomus) after ca. 450 CE suggests that the benthic zone had begun to freshen. In short, the metazoan succession in Zone M1 shows the same transition from a marine to a brackish and then finally freshwater system seen in the aquatic plant record (Zone A-1, Extended Data Fig. 3).

Metazoan biozone M-2 (Extended Data Fig. 2) also corresponds to biozone A-2 calculated from aquatic plant reads (Extended Data Fig. 3). This zone features the establishment of chronomids and several freshwater ostracod genera. Reads from the fully-marine taxa common in Zone M-1 either disappear or are reduced to trace frequents, except for *Saccoglossus* (acorn worms) and aquatic gastropods which may have recolonized the lake periodically via marine surges. The brackish adapted *Gasterosteus* (stickleback) and the anadromous *Anguilla* (common eel) are also detected throughout the sequence.

Biozone A-2 is differentiated by the presence of the macroalgae Spirogyra and an increase in *Myriophyllum* biomass, indicative of a freshening and increasingly eutrophic lake. Zone M-2, the corresponding biozone in the metazoan record is distinguished by abundant reads matching to the genus *Anas* (dabbling ducks). Most of these reads are probably from *A. platyrhynchos* (Mallards), but *A. acuta* (Northern Pintail) and *A. strepera* (Gadwall) may also have visited the lake. Reads from *Aythya* (diving ducks) were not as common, but both *A. fuligula* (the Tufted Duck) and *A. marila* (the Scaup) may have foraged in the lake during this period. The Tjörnin was clearly a productive duck habitat between ca. 650 and 900 CE and this must have impacted the aquatic macrophyte community[46]. In contrast, reads from *Anser* (grey geese) were only detected at trace values in this zone. Several species of *Anser* including *A. brachyrhynchus* (Pink-footed Goose) and *A. anser* (the Greylag Goose) breed in Iceland today[47], but they were relatively rare in the lake at this time. The genus *Branta* (black geese) was not detected in the metagenomic record at all. This suggests that in the pre-Landnám period ducks dominated the lake, while geese were relatively rare.

The transition from Zone M-2 to M-3 is generated by a rapid shift in the terrestrial vertebrate community. The proportion of *Anas* declines sharply as those of introduced mammals including: *Bos*, *Ovis*, *Sus* (pigs), *Equus* (horse) rise rapidly to dominate the assemblage. The earliest reads from sheep are detected at about 900 CE shortly after the traditional date for the beginning of the Landnám at 877 CE, but the read counts for all these mammals increases dramatically by 940 CE, suggesting that the Norse settlement at the lake had grown by this time.

In contrast, *Anas* read frequency sharply declined after human arrival. The average *Anas* read count in Zone M-2 is ca. three times that of Zone M-3, indicating that fewer ducks visited the lake after the settlement was established. On the other hand, reads from *Anser* increase by a hundredfold in Zone M-3. Given the preponderance of reads from domestic taxa in Zone M-3 and the scarcity of *Anser* reads in Zone M-2, it is possible that most of the *Anser* reads detected in Zone M-3 come from *A. anser domesticus* (domesticated geese) kept by Norse settlers. Domestic geese were introduced to Scandinavia in the Roman Iron Age and were common animals on Norse farmsteads[48–50]. Domestic ducks did not become common in Europe until much later in the medieval period[51]. Thus, the shift in the avifauna from a large population of wild ducks to presumable fewer possibly domestic geese in Zone M-3 is an anthropogenic one. Zone M-3 lasted about 400 years, from just after the Landnám to ca. 1350 CE, and includes the entire period of the Icelandic Commonwealth.

Copious reads from pigs and horses were registered along with those from sheep and cattle in this zone. The last three were important to the Icelandic economy throughout its history, but this not the case for pigs. Archaeological data from multiple sites attests to the presence of swine at several early settlements in Iceland, but they were never numerous[52–54] Medieval swine in Iceland could not forage on pannage as they did in much of Europe, so they were probably fed food refuse close to the farmsteads while rooting in home-fields and middens. With time, their numbers dwindled, and they were increasingly confined to wealthier farms, typically ecclesiastical estates. Sometime in the early 17th century, swine disappeared completely from the archaeological record[52]. The concurrent extinction of the Scottish grice is probably not coincidental. In both regions, declining cereal cultivation and a shift to intensive sheep herding for fabric production made pigs less economical in the post medieval and early modern periods[52,55].
At about 1350 CE, the beginning of Zone M-4, the proportion of eDNA reads from swine drop to trace levels. Those from horses also declined sharply compared to those of sheep and cattle. However, the read counts of all domestic mammals dropped by tenfold in this zone consistent with a significant drawdown of the mammalian biomass in the Tjörnin catchment at this time. This implies a shift in the intensity of cultivation and grazing as well as the type of settlement.

Other distinguishing features of Zone M-4 include an increase in the *Chironomus* (non-biting midge) reads suggesting that these benthic insects recovered from a slight decline during the previous zone. This may be linked to reduced minerogenic influx at this time or nutrient influx (see description for Aquatic Zone A-4, above and supplementary Data section on *Chironimids*).
The increase in reads matched to *Lumbricus* in this zone are more difficult to explain. Erosion from cultivation and grazing would be expected to mobilize sediment containing these terrestrial annelids

and their DNA, but *Lumbricus* only increases in Zone A-4 when anthropogenic disturbance is reduced. Another possibility is that the increase tracks the subsequent establishment and growth of a population of a *Lumbricus* species first introduced in the previous zone, A-3. Today on Iceland, *L. terrestris* and *L. castaneus* along with species in the related genera *Aporrectodea* and *Octolasion* are distributed in anthropogenic habitats, such as hay-fields, pastures, and gardens, first created in the Tjörnin catchment during Zone A-3. While it is considered likely that the presence of lumbricid earthworms in Iceland is a consequence of the Landnám, they might have been introduced or established somewhat later in the Tjörnin catchment[56]. Reads from *Lumbricus* decrease during the 20$^{th}$ century as the catchment is paved over.

Another distinctive feature of Zone M-4 is the rise of two fish genera, *Gasterosteus* (sticklebacks) followed by *Gadus* (cod). *Gasterosteus* are a marine genus very tolerant of brackish conditions. They were well represented at the base of the sequence but contributed few reads in Zone M-3. However, *Gasterosteus* comprises as much as 25% in the period 1450 and 1500 AD at the base of Zone M-4. They may have flourished in the lake at that time due to increased marine influx (Fig. S). The presence of Gasterosteus is noted in old stories about the Tjörnin recorded in the 18$^{th}$ century, in which other fish are mentioned as well including *Salmo salar* (Atlantic salmon) and *Salmo trutta* (brown trout). These anadromous genera were not detected in the metagenomic analysis. In contrast, *Gadus* is a marine fish that could not have lived in Lake Tjörnin during its brackish/freshwater phase. Copious sequences matched to Gadus began to accumulate at ca. 1450 and were particularly abundant in the late 19th century. These were probably derived from fish waste dumped into the lake by Reykjavik´s fish mongers and are thus indicative of catchment urbanization.

Zone M-5 corresponds to the 20$^{th}$ century. The increased eutrophication at that time is described in the previous section on the aquatic flora (Aquatic Zone A-5). At least three species of ostracod, *Darwinula*, *Heterocypris*, *Sarscypridopsis* all flourish in this zone. *Heterocypris*, and *Sarscypridopsis* appear in the record for the first time, so they may have been introduced to the lake at the turn of the last century.

The proportion of reads matched to *Anas* increase only slightly in this zone even though the Tjörnin is now known as the "world's largest bread soup" because citizens and tourists alike feed bread to ducks, drawing large flocks to the lake[57]. However, the average number of *Anas* reads was ca. tenfold higher in Zone M-2. This suggests that either the concentration of ducks feeding in the lake was much higher in the pre-settlement period, or there are taphonomic factors affecting the preservation of these reads at the top of the sequence.

Human sequences are also conspicuous by their absence in Zone M-5. Effluvium from the city was allowed to drain into the lake until the 1960s, which would have been a source of human nucleic acid[1]. The lipid analysis is also consistent an influx of human faeces at this time, but the associated DNA either did not survive or was not detected in these samples. Though this zone yielded many reads relative to most of the lake sequence (See Metazoan Sum, Extended Data Fig. 2), these nutrient rich sediments harboured relatively few vertebrate reads.

Though reads matched to *Clupea* (herring) never occur at high frequencies, their distribution has historical implications. The commercial herring fishery in Iceland began in the late 19$^{th}$ century when Norwegian fishermen introduced drift netting from steam ships. Towns like Siglufjörður on Iceland's north coast, were the first to develop into herring processing centres. The industry grew and came increasingly under Icelandic control in the early 20$^{th}$ century. By the 1930s and 1940s, herring had become Iceland's most valuable export, accounting for up to 40% of the nation's foreign exchange earnings. However, in 1969, stocks collapsed due to overfishing[58].This chronology corresponds closely to the distribution of reads matched to *Clupea* in the record. The majority of these are registered in samples dated from 1908 to 1969, with the maximum value falling in 1921. This suggests that these reads were incorporated as herring was processed or transhipped through Reykjavik during the boom years of the herring fishery. The sample dated to 1969 is the last to have *Clupea*. A few reads are also registered in zones M-3 after the Landnám and in M-1, the marine layer at the base of the sequence.

**Metazoan eDNA phylogeny**

**Horse:** modern horses can be classified into 18 major mitochondrial haplogroups[59]. A recent study on Icelandic horses identified six haplogroups (B, D, L, M, N, Q) currently present in Iceland and the Reykjanes peninsula[59,60]. However, only three of the six (Q, M, and N) are supported by our eDNA data (Extended Data Fig. 10), and almost all eDNA supports are found in the period of the earliest Viking settlement at Reykjavík. For haplogroups N and Q, eDNA data is insufficient to place the identifications to specific breeds, and hence only provides support at ancestral nodes of the haplogroups. Haplogroup Q contains both modern Icelandic horse and the Lewitzer ponies. The latter is originated in northeastern Germany and was developed through crosses among Shetland ponies, Arabians, Hutsuls, and occasionally Trakehners. Haplogroup M is the only lineage for which eDNA supports were placed to a close-to-tip branch, where eDNA reads was placed to a node leading to two individuals of modern Icelandic horse and the Vyatka house. The later is a hardy, medium-sized indigenous Russian breed historically valued for its strength, endurance, and adaptability, and traditionally employed in farming, transport, and postal troikas[61].

**Sheep:** many sheep eDNA reads were assigned to mitochondrial haplogroup B1, which predominantly comprises sheep from Northern Europe[62], including all modern Icelandic sheep. One SNP supported a placement on the ancestral node to four Icelandic landraces, three of which are recognized as the Icelandic Leader sheep. As a traditional landrace of Iceland, the Icelandic Leader sheep is renowned for their resilience and capacities in guiding sheep flocks under harsh weather conditions[63,64]. Owing to its importance, the Icelandic Leader sheep was mentioned in the Icelandic law book Jónsbók to as early as 1281 CE, referred as metfé ("priceless animals")[65] Their introduction as early as 936 CE, suggested by our eDNA data, could have contributed to the early settlement and the establishment of husbandry under Iceland's challenging environment (Extended Data Fig. 10).

**Cattle:** eDNA reads were dominantly assigned to mitochondrial haplogroup T (including subgroups T1, T2, T3, and T4) for cattle[66]. However, despite the abundance of mitochondrial reads, further resolution within and among these subgroups is limited, as haplogroup T lacks sufficient internal phylogenetically informative variations. Nonetheless, many reads covering SNPs that represent private variants were placed to specific individuals. For instance, between 912 and 984 CE, we identified reads matching private SNPs from the Rendena breed, primarily found in northern Italy and associated with haplogroup T2. Similarly, reads matching to private SNPs from the rare Katerini breed, originating from northern Greece, were detected in the layer dated to 1766 CE. In addition, several reads were also aligned to individuals outside haplogroup T, including those to haplogroups Q, P, and I. While these ending tips on the tree may have captured eDNA reads through specific private variants, no reads were assigned to their respective backbone branches (i.e., internal nodes along the tree linking the root to these tips), despite these nodes containing sufficient diagnostic mutations. These tip-level placements therefore likely resulted from shared private variants rather than phylogenies. This suggests that the real populations contributing to the Icelandic cattle eDNA signals are likely not sampled in the reference genome panel (Extended Data Fig. 10).

**Pigs:** from the layer dated to 949 CE, we detected eDNA corresponding to the Norwegian Landrace[67], yet further inferences require a more in-depth understanding of the genetic background of this breed (Extended Data Fig. 10).

**Impact of the Landnám on *Betula* populations**

Conventional wisdom holds that the Norse Landnám (settlement) in the late 9th century was a turning point in Iceland's ecological history, leading to the near-total deforestation of birch woodlands (*Betula pubescens*). Paleoecological data, particularly pollen and sediment analyses, demonstrate that Iceland's lowlands were once dominated by birch forests, which rapidly declined after human arrival. The longstanding paradigm holds that deforestation was primarily anthropogenic. Pollen records consistently show a sharp drop in *Betula* pollen around the Landnám tephra horizon (877 CE), replaced by indicators of open landscapes, such as grass and heath pollen. The widespread presence of charcoal layers and introduced plant species further support the role of human activity in transforming the landscape. Livestock grazing is also recognized as the key factor preventing birch regeneration. Sheep and other domesticated animals continuously browsed saplings, ensuring that once forests were cleared, they could not recover. The infield/outfield agricultural system reinforced this pressure, leading to long-term vegetation shifts and soil degradation. Thus, the legacy of Landnám is still evident. Modern studies estimate that around 1% of Iceland's original birch cover remains, with extensive soil erosion persisting across the island. While reforestation efforts have seen limited success in recent decades, the transformation of Iceland's ecosystems due to Norse settlement remains one of the most striking examples of human-driven environmental change in the North Atlantic.

At the core of the narrative that humans were responsible for clear-cutting *Betula* forests are data from pollen percentage diagrams. Such diagrams show the relative abundance of different plant taxa in a sample, facilitating the reconstruction of past vegetation composition[68]. However, this does not account for absolute pollen input, overrepresentation or underrepresentation that may occur due to differential pollen production among plant species, or the zero-sum situation in which one taxon may appear to decline only because others increased. To avoid such problems some studies, use pollen concentrations, which indicate how much pollen is preserved in a sediment layer. This is useful for assessing sedimentation processes and pollen preservation but fails to consider sedimentation rate variations over time. Others use pollen accumulation rates (PAR) to evaluate vegetation changes (Table S6, Figure S41), which provide an absolute measure of past pollen influx and makes it possible to infer past vegetation productivity, climate change, and biomass in the catchment over time. However, PAR numbers from lakes can be difficult to interpret due to the possible influx of redeposited *Betula* (and other) pollen from eroded soil. Among the twelve sites across Iceland where PAR studies were conducted, two sites show a post-Landnám *Betula* increase, four sites show a pre-Landnám decrease, three sites have records outside the time range relevant for Landnám, while two sites have insufficient age constraints to interpret any transitional changes. Only one site, Reykholtsdalur, shows a rapid *Betula* decline after Landnám. Thus, the PAR values do not provide a consistent signal for a rapid *Betula* decline after Landnám across Iceland, casting doubt about the equilibrium–Landnám disturbance–new equilibrium ecological model, which has been part of the prevailing mid-to-late 20th-century narrative.

That we found little or no evidence for birch woodland within the Tjörnin catchment compares well with other coastal sites such as Ketilsstaðir[69] and Reykjanes[70]. These were environments where pioneers could settle and farm without clearing birch[71], but which provided unwooded wetlands for grazing and hay meadows. This is corroborated by many place names within Reykjavík that bear witness to wetland habitats that are now lost.

**Table S6. Pollen studies from Iceland with calculated PAR values and our inference of their results[68,72–79].**

| Reference | Site name | Latitude | Longitude | Age range of pollen study | PAR inference | PAR range |
|---|---|---|---|---|---|---|
| **Erlendsson PhD thesis (2007), Erlendsson & Edwards 2009** | Ketilsstaðir | 63.443889 | -19.168056 | AD 500–1500 | Pre-CE 870 betula decline | 100-800 |
| **Erlendsson PhD thesis (2007), Erlendsson & Edwards 2009** | Reykholtsdalur | 64.662047 | -21.305906 | c. CE 500-1000 | Betula decline at landnam | 300-1750 |
| **Erlendsson PhD thesis (2007), Erlendsson & Edwards 2009** | Stóra-Mörk | 63.661464 | -19.905161 | c. CE 500-1000 | Pre-CE 870 betula decline | 0-400 |
| **Eddudottir et al. 2020** | Galtaból, NW Iceland | 65.265083 | -19.7266 | c. 4200-0 cal. yr BP | Decline in Betula around 1000 CE (low resolution) | 150-600 |
| **Eddudottir et al. 2018** | Kagaðarhóll ake, NW Iceland | 65.587778 | -20.132778 | c. 10,100–7000 cal. yr BP | Not relevant to Landnam transition | 100-2500 |
| **Eddudottir et al. 2016** | Barðalækjartjörn, NW Iceland | 65.42 | -19.873056 | c. 10,300–200 cal. yr BP | Decline in Betula around 1000 CE (low resolution) | <1000 |
| **Eddudottir et al. 2015** | Kagaðarhóll, NW Iceland | 65.587778 | -20.132778 | Early Holocene to c. 3000 cal. yr BP | Not relevant to Landnam transition | 0-2000 |
| **Eddudottir et al. 2017** | Kagaðarhóll, Barðalækjartjörn, Northwest Iceland | 65.587778 | -20.132778 | Hekla 4 tephra impact (c. 4200 cal. BP) | Not relevant to Landnam transition | >500 |
| **Wastl et al. 2001** | Vesturardalur-Skíðadalur, Tröllaskagi, Northern Iceland | 65.75 | -18.75 | c. 9200 BP – Present | Insignificant (very low resolution) | 0-2750 |
| **Hiles et al. 2021** | Kalmanstjörn, Mývatnssveit, NE Iceland | 65.657667 | -17.233833 | Last 3000 years | Post-landnam betula increase | <10 |
| **Lawson et al. 2007** | Helluvaðstjörn, Mývatnssveit, Northern Iceland | 65.572869 | -17.180797 | Last 3500 years | Post-landnam betula increase | 200-1600 |
| **Dixon, 1997** | Háls, southwestern Iceland | 64.933086 | -23.317196 | Poor chronology | NA | NA |

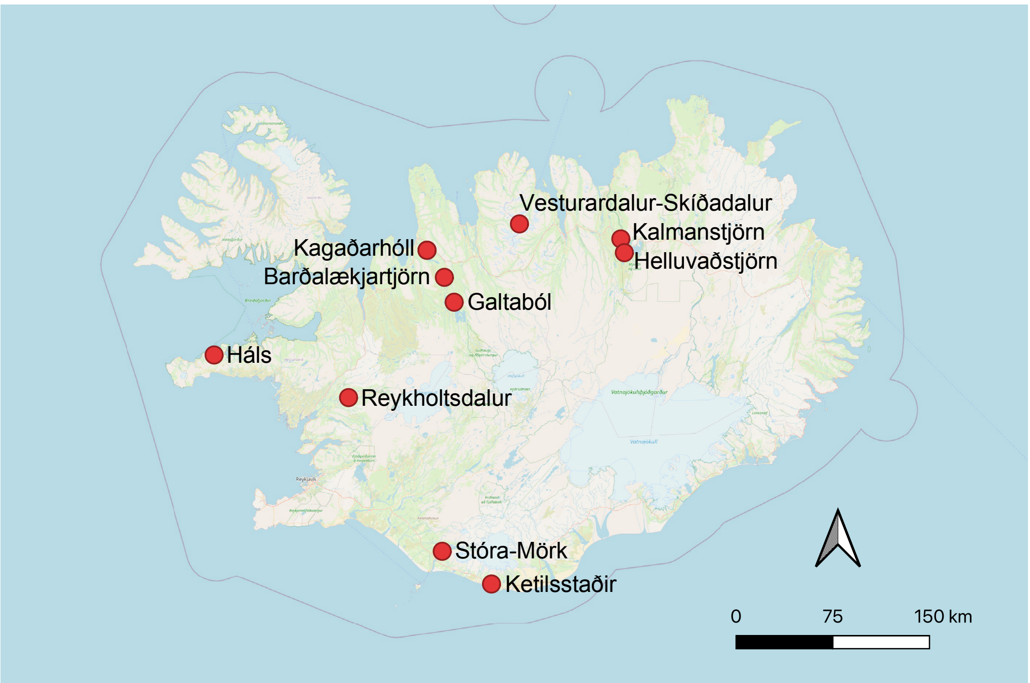

**Figure S41. Map of pollen records from Iceland with PAR calculations.**

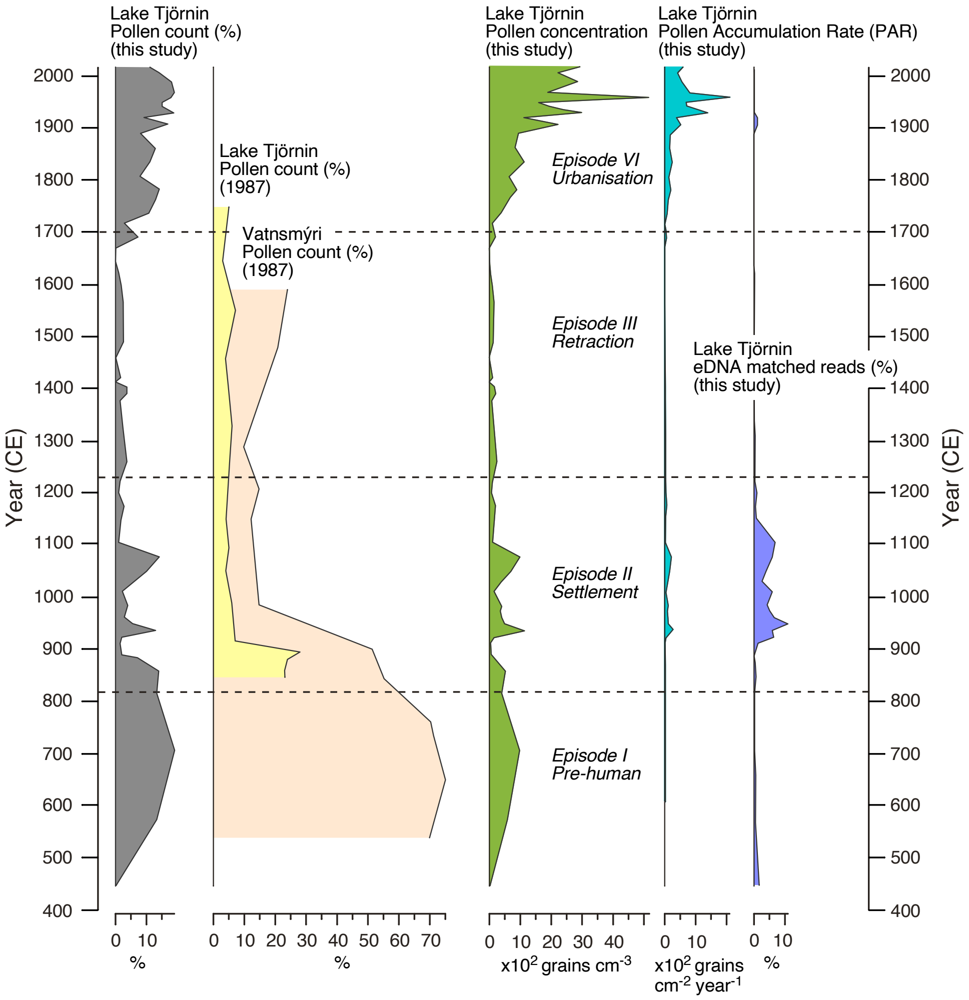

**Figure S42. Comparison of pollen records from 1987[80] by Hallsdottir and this study (2025) with eDNA results from the key taxa of Betula.**

Figure S42 compares previous pollen study of *Betula* within the Tjörnin catchment expressed as percentages[80], with a new pollen record (concentration and PAR; see supplementary information section: Comparison of palynological and metagenomic records and Extended Data Fig. 5) as well as eDNA results from this study. Hallsdóttir[80] found a significant decline in *Betula* at the time of settlement, most notably in the peat succession from Vatnsmýri, where *Betula* percentages dropped from 70% to less than 15%, and from 25% to 5% in the Tjörnin lake record. The discrepancy in the size of the *Betula* percentages between studies is due to the exclusion of sedges from the base sum in the Vatnsmýri study, but not at Tjörnin.

Environmental DNA does not show this trend; instead, it indicates an increase in *Betula* after Landnám, lasting until 1150 CE, which is mirrored by the PAR pattern, although it begins to decline around 1100 CE. We interpret this as evidence of increased biomass and human activity in the catchment. Tree biomass and pollen accumulation rates (PAR) are closely related, as modern PAR values from lake sediments have been shown to be robust and precise estimators of contemporary aboveground biomass for major tree taxa, such as *Pseudotsuga*, *Pinus*, *Notholithocarpus*, and the TCT pollen group (i.e., Taxodiaceae, Cupressaceae, and Taxaceae, which include coniferous trees such as redwoods, cypresses, junipers, and yews). By calibrating PAR-biomass models using spatially explicit vegetation data, historical changes can be inferred in forest biomass, with greater precision achieved when using assemblage-level relevant source areas of pollen compared to taxon-specific estimates[81,82].

**The ancestry of Icelandic Viking Age barley**

Initially, joint SNP-calling was performed using a global reference panel of 116 diverse domesticated barley accessions. This approach clustered the ancient barley eDNA reads from Tjörnin with domesticated six-rowed barley of European origin (Fig. S43B)[83]. In a second approach, individual mapping of ancient eDNA reads was performed against wild and landrace accessions within the extended barley pangenome[84], including two newly assembled genomes of historically relevant six-rowed barley landraces Bere (Scottish Isles, Northern Europe's oldest known barley landrace[85]) and Asplund (selection from a Swedish landrace). Among the 55 diverse genomes tested, the ancient samples exhibited the greatest mapping breadth when aligned with Bere and Asplund genomes (Figs. S43C). In a third approach we estimated the relative proportions of reference genomes from the extended barley pangenome[84] and the Bere and Asplund assemblies represented in the ancient short reads identified across the Tjörnin core sample. We show that when barley reads are found in the data, starting at ca. 912 CE (Fig. S43A), the dominant variety is Scottish Bere barley with a smaller proportion of HOR 2180 (Landrace from former Czechoslovakia). As these proportions are relatively constant in time, we interpret this as evidence for a variety most closely related to Bere, but not 100% identical to Bere as the best statistical model is a mixture between Bere and a few other varieties with minor contributions. Finally, we called ancient SNPs from the most deeply sequenced core sample (CGG3016635, dated 984 CE) and subsequently filtered them against unimputed SNP variants from the Core1000 panel[84] to retain exclusively SNP positions represented in contemporary barley diversity. Using the resulting 4,901 ancient SNP positions distributed across barley chromosomes 1H to 7H, we constructed a phylogenetic tree, which grouped the ancient Icelandic barley with European or European-derived six-row barley varieties. Notably, three out of seven of these barley lines displayed a distinctive and rare MKK3 haplotype linked explicitly to Nordic landraces such as Bere and Asplund (Fig. S44; Table S7;[86]). In summary, this analysis suggests that the ancient barley from the Tjörnin sample is closely related to Bere barley and landraces carrying a distinctive and rare MKK3 haplotype linked to Nordic landraces.

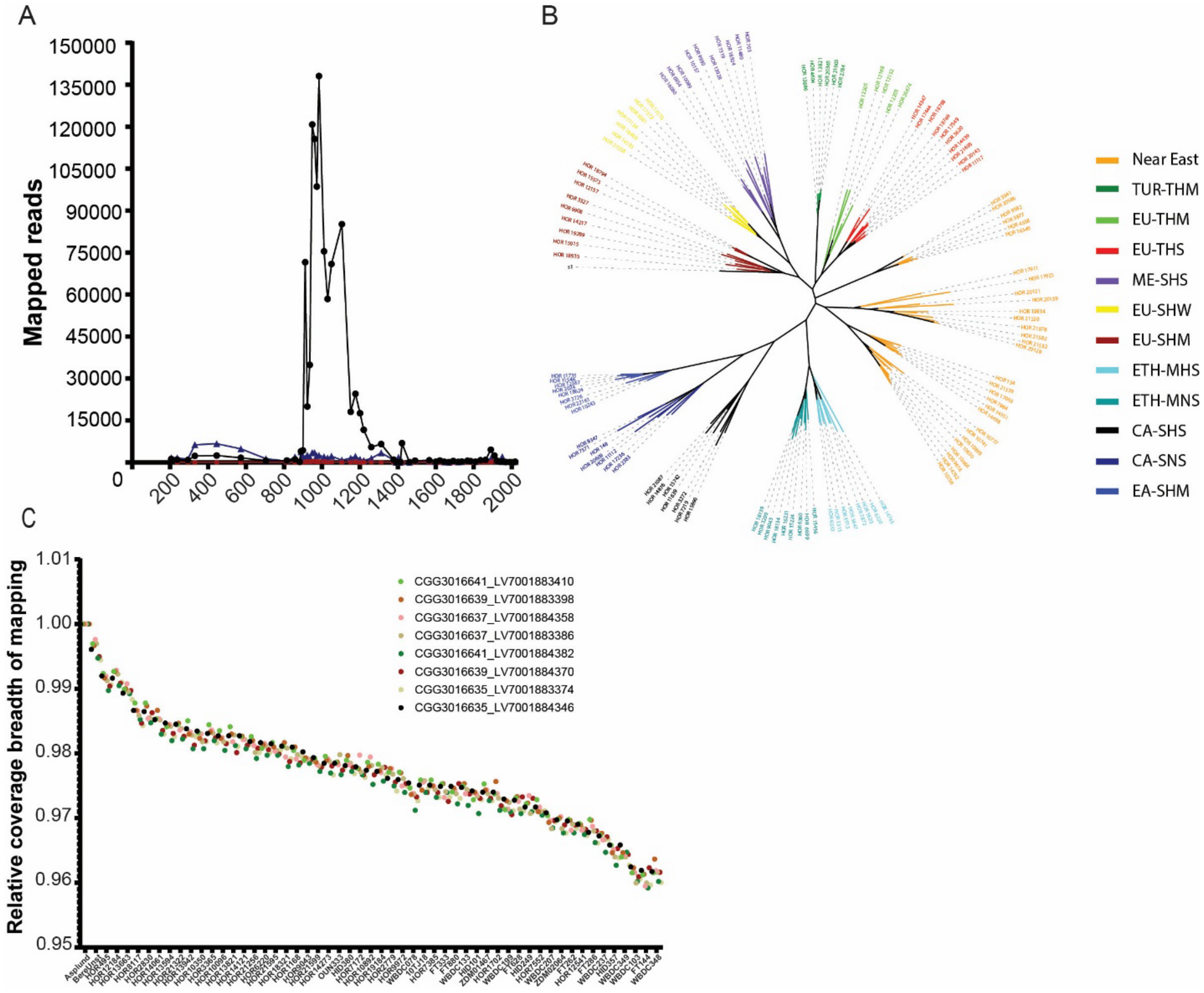

**Figure S43. The ancestry of Icelandic Viking Age barley.** A) eDNA reads competitively mapped to *Hordeum vulgare* (black circles), *Avena* (red squares) and *Triticum* (dark blue triangles) identified across time in the Lake Tjörnin sediment core. B) Unrooted neighbour-joining tree showing the relationships between the ancient environmental DNA sample from Island and ecogeographically diverse modern domesticated barley accessions from Guo and coworkers[83]. C) Relative coverage breadth of ancient barley reads mapped to 53 wild and land race barley assemblies from the extended pan genome and the Bere and Asplund assemblies.

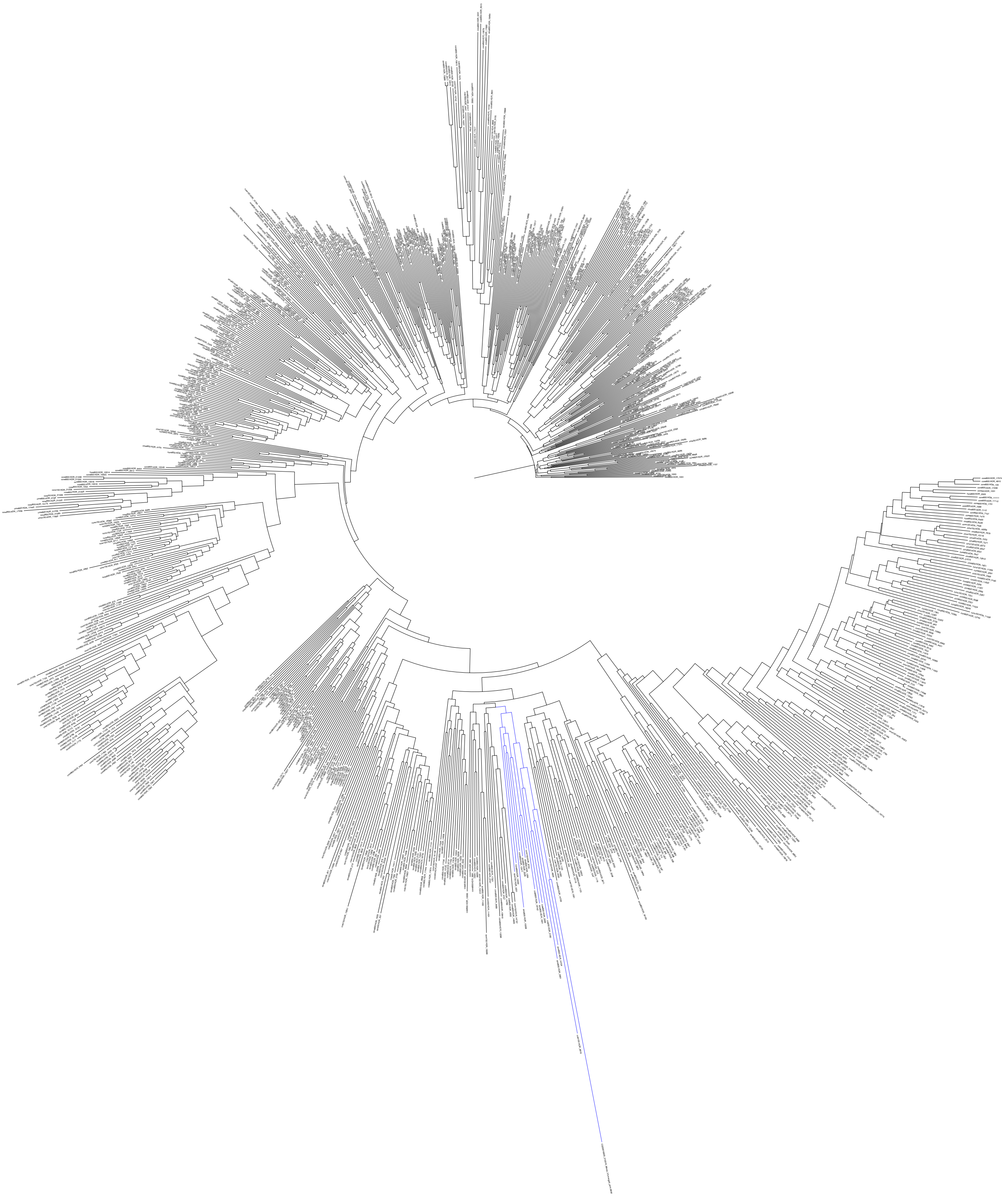

**Figure S44. Phylogenetic tree showing the ancient Icelandic barley samples phylogenetic clustering in the CORE1000.**

#### Table S7. Subclade clustering phylogenetically with ancient barley sample

| Type | Name | Row type | Accession | *MKK3* CN | $MKK3^{E165Q}$ |
|---|---|---|---|---|---|
| Traditional cultivar/landrace | Orkisz | convar. vulgare/convar. hexastichon | HOR_8819 | 3 | 0 |
| Advanced/improved cultivar | CARLE | convar. vulgare/convar. hexastichon | HOR_14704 | 2 | 1 |
| Advanced/improved cultivar | Ochnella | convar. vulgare/convar. hexastichon | HOR_2961 | 3 | 1 |
| Advanced/improved cultivar | | convar. vulgare/convar. hexastichon | HOR_2180 | 1 | 0 |
| Advanced/improved cultivar | Podhorecky | convar. vulgare/convar. hexastichon | HOR_2099 | 1 | 0 |
| Advanced/improved cultivar | Tiroler Pfauengerste | convar. distichon | HOR_1601 | 2 | 1 |
| Traditional cultivar/landrace | LANDSORTE NR.750 | convar. vulgare/convar. hexastichon | HOR_19742 | 3 | 0 |

#### EM Method

We here describe the method used to identify the origin of the barley variety based on the short read data. This method is based on a slight modification of the method described in earlier studies by Pipes and colleagues [87]. This provides maximum likelihood estimates of the proportions of reference genomes represented in a pool of short reads:

##### Notation

For each read $i = 1, \ldots, N$ and reference genome $j = 1, \ldots, K$:

- $n_{ij}$: number of sites compared (read length)
- $d_{ij}$: number of mismatches observed
- $\pi_j$: true proportion of reads from genome $j$ (to be estimated)

The goal is to estimate the mixture proportions $\boldsymbol{\pi} = (\pi_1, \ldots, \pi_K)$ subject to $\Sigma_{j=1}^{K} \pi_j = 1$ given the observation of alignments of reads to all genomes.

##### Likelihood Model

This method assumes a single error rate $\varepsilon$ for all comparisons and symmetric errors among all nucleotides. For read $i$ and genome $j$, the conditional likelihood is then:

$$Q_{ij} = P(\text{read } i \mid \text{ read } i \text{ is from genome } j) = (\epsilon/3)^{d_{ij}}(1-\epsilon)^{n_{ij}-d_{ij}}$$

The full log likelihood is obtained as

$$\prod_{i=1}^{N}\sum_{j=1}^{K} Q_{ij}\pi_j$$

We optimize this likelihood function using an EM algorithm with SQUAREM optimization.

##### EM Algorithm

**E-step:** Calculate posterior probabilities

$$w_{ij} = \frac{Q_{ij}\pi_j}{\sum_{k=1}^{K} Q_{ik}\pi_k}$$

**M-step:** Update proportions

$$\pi_j^{(\text{new})} = \frac{1}{N}\sum_{i=1}^{N} w_{ij}$$

The error rate is similarly estimated using the EM algorithm using the following M-step:

$$\epsilon^{(\text{new})} = \frac{\sum_{j=1}^{K}\sum_{k\neq j}\sum_{i=1}^{N} w_{ij} d_{ik}}{\sum_{j=1}^{K}\sum_{k\neq j}\sum_{i=1}^{N} w_{ij} n_{ik}}$$

To apply this method, we used the reads mapped to all 78 reference genomes as described in the methods section. We included only reads that aligned to all genomes as differences in assembly might otherwise bias the analyses. The number of such reads per sample, along with the parameters of the EM can be found in source data (EM). The full results of proportion per genome per sample can be found in Source data (proportions). There are a constant low rate of barley reads that do not get assigned a high proportion to any one genome. This likely represents a low level of contamination or species misassignments. However, we notice that at the time where a large proportion of barley reads are found in the data, starting at ca. 912 CE, the dominant variety is Bere with a smaller proportion of HOR_2180 (Fig S45). As these proportions are relatively constant in time, we interpret this as evidence for a variety most closely related to Bere, but not identical to Bere as the best statistical model is a mixture between Bere and a few other varieties with minor contributions. In more recent times, when barley reads again become scarce in the sample, the likely artefactual observations of trace amounts of barley with no likely source genome re-appears.

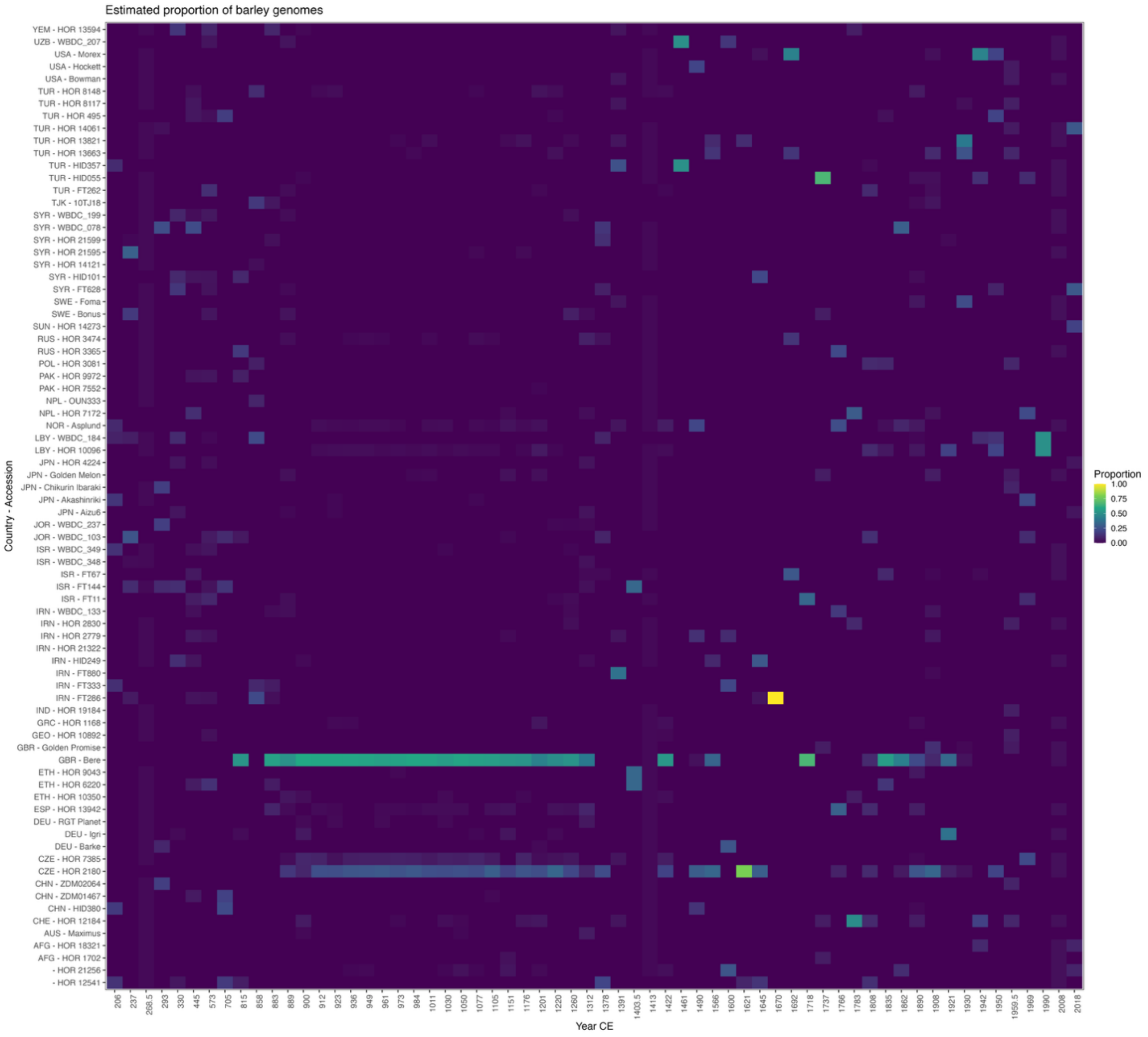

**Figure S45. Proportions of barley reads assigned to reference genomes over time.** Heatmap showing the estimated proportion of reads assigned to each of 78 barley reference genomes across samples ordered by date. Proportions were inferred using an EM algorithm under a uniform error model. A clear signal appears around 912 CE, dominated by Bere with minor contributions from HOR 2180, suggesting a mixed barley variety related to Bere. Earlier and later periods show low, diffuse signals consistent with contamination or misassignment.

**Comparative analysis of historical herbarium barley**

To further evaluate the identity of the Icelandic Viking Age barley, we analysed historical herbarium samples collected in Iceland. The aim was to test whether the barley identified in the Lake Tjörnin sediment core corresponds to the same variety as historically cultivated barley in Iceland, or whether it represents a distinct type.

**Samples**

We analysed two herbarium grain samples housed at the Natural History Museum of Denmark (Fig S46). The first specimen (ID:  D_1934) was collected in Djúpivogur, Iceland, in 1934). The second specimen (ID: M_1897) was collected in 1897 in Mödruvellir, Iceland, and was noted to be infected with *Ustilago hordei.*

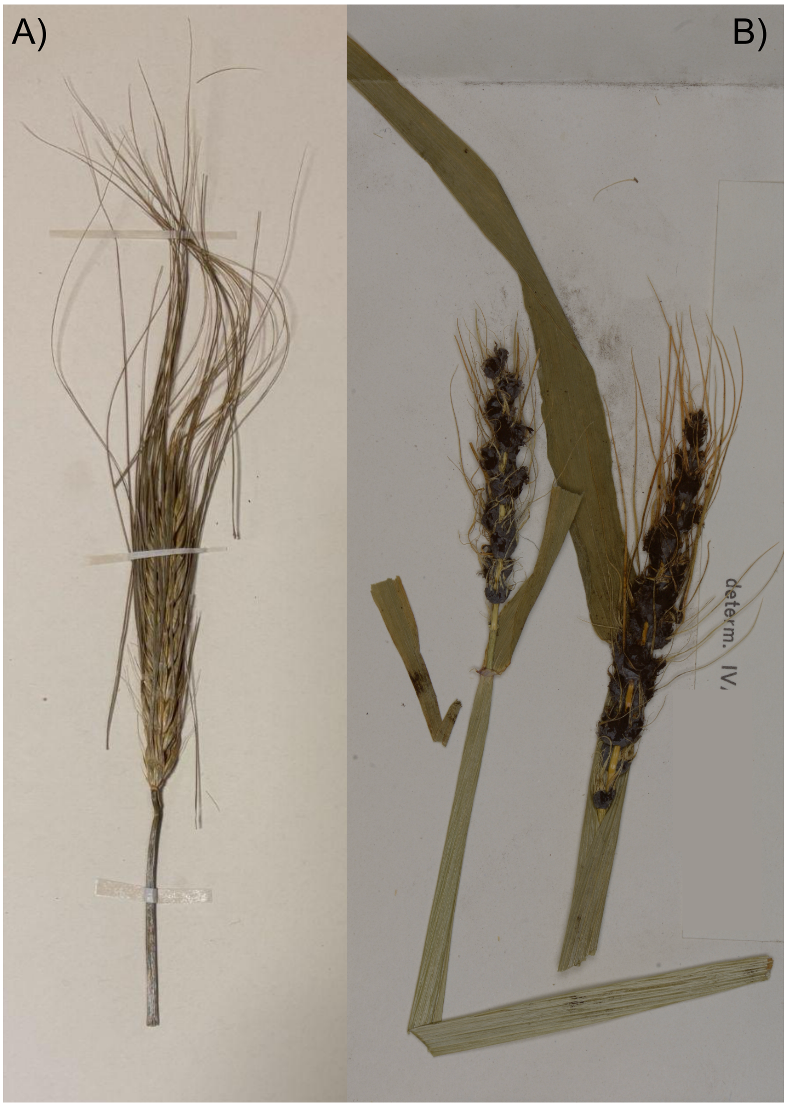

**Figure S46. Herbarium barley specimens from Djúpivogur (A, D_1934) and Mödruvellir (B, M_1897), Natural History Museum of Denmark.**

**DNA recovery**

Two historic herbarium barley specimens curated at the Natural History Museum of Denmark were selected as comparators for the barley detected in the Tjörnin sediment core: Djúpivogur, a seed with attached awn (Figure S46A; D_1934), and Mödruvellir, a flat-pressed stem (Figure S46B; M_1897). A no-template control (NTC) was included throughout. Sampling, pre-treatment, DNA extraction and library preparation were performed in dedicated clean rooms at the Globe Institute, University of Copenhagen, Denmark.

Because the specimens were likely treated with mercuric chloride as part of museum preservation efforts, surfaces were washed with EDTA to chelate $Hg^{2+}$ and avoid interference with dithiothreitol (DTT) in the lysis buffer.

The specimens were further broken down with a scalpel to reduce particle size, then washed for 10 min at room temperature with 50 mM EDTA (Invitrogen), followed by two rinses with molecular-grade H2O (BioNordika). Washed material was transferred to SAFE 2D-barcoded 2 mL tubes (LVL technologies) preloaded with 200 µL 0.1 mm ceramic beads and 200 µL 0.5 mm ceramic beads (Omni), and 800 µL N-phenacylthiazolium bromide (PTB)-based extraction buffer[88]. The PTB buffer contained 0.4 mg/mL Proteinase K (Roche), 2.5 mM PTB (Fluorochem), 50 mM DTT (Roche), 1% SDS (Sigma-Aldrich), 10 mM Tris (Thermo Scientific), 10 mM EDTA (Invitrogen), and 5 mM NaCl (Invitrogen) in molecular-grade H2O. Samples and the NTC were bead-beaten on a FastPrep-96 (MP Biomedicals) for two rounds at 1600 RPM for 30 s each, separated by 30 s. Tubes were then incubated with shaking overnight at 37° C. After incubation a final 30 s bead-beating at 1600 RPM was performed, followed by a brief centrifugation to pellet insoluble material. A 500 µL supernatant aliquot was transferred to a new tube. A 240 µL portion of lysate was combined with 950 µL QSB1 (Qiagen) and 10 µL paramagnetic beads (G-Biosciences) and rotated for 15 min. The supernatant was discarded, and the beads were washed once with MW1 buffer (Qiagen) and twice with 80% ethanol (Sigma-Aldrich). Residual ethanol was removed and beads were air-dried for 5 min at room temperature, then DNA was eluted in 50 µL EB buffer (Qiagen). DNA yield and fragment size were assessed with the Qubit 2.0 dsDNA HS assay (Invitrogen) and the Agilent Fragment Analyzer system. Extracts were converted to unique dual-indexed single-stranded version 2.0 libraries following Gansauge and coworkers[89] and sequenced on a NovaSeq 6000 S4 flow cell with 2x100 bp paired-end reads.

**Basic bioinformatics**

Adapter sequences were trimmed and overlapping reads were collapsed conservatively using AdapterRemoval (2.3.2[90]). Reads below 30 base pairs were discarded. Collapsed reads were mapped to the *Hordeum vulgare* Morex reference genome using bwa (0.7.17,[91]) with seeding disabled (-l 512). Reads in lane were merged, and duplicates were marked using Picard MarkDuplicates (2.25.0, http://picard.sourceforge.net) with a pixel distance of 12000.

Both specimens yielded well-preserved endogenous barley DNA. D_1934 contained 89% endogenous DNA with an average coverage of 2.38x, while M_1897 contained 91% endogenous DNA at 3.77x coverage, confirming suitability for comparative genomic analysis.

**Ancestry estimation**

Reads from each herbarium specimen were downsampled into ~40M-read chunks and processed through the same competitive mapping eDNA pipeline as for the lake sediments. Reads assigned to the genus *Hordeum* were aligned against the panel of 78 barley reference genomes, filtering for primary alignments only. Reads aligning across all references were extracted, merged, and analysed using the EM mixture estimation framework (see below).

Both herbarium barley specimens showed highly similar ancestry profiles, closely resembling the sedimentary eDNA signal (Fig. S47). As in the core data, the dominant component was Bere, accompanied by a consistent contribution from HOR 2180. In addition, both herbarium specimens revealed a clear Asplund component not evident in the sedimentary eDNA, suggesting that Icelandic barley cultivated in the late 19th–early 20th centuries represented a Bere-related variety with a new Asplund component representing import from Scandinavia.

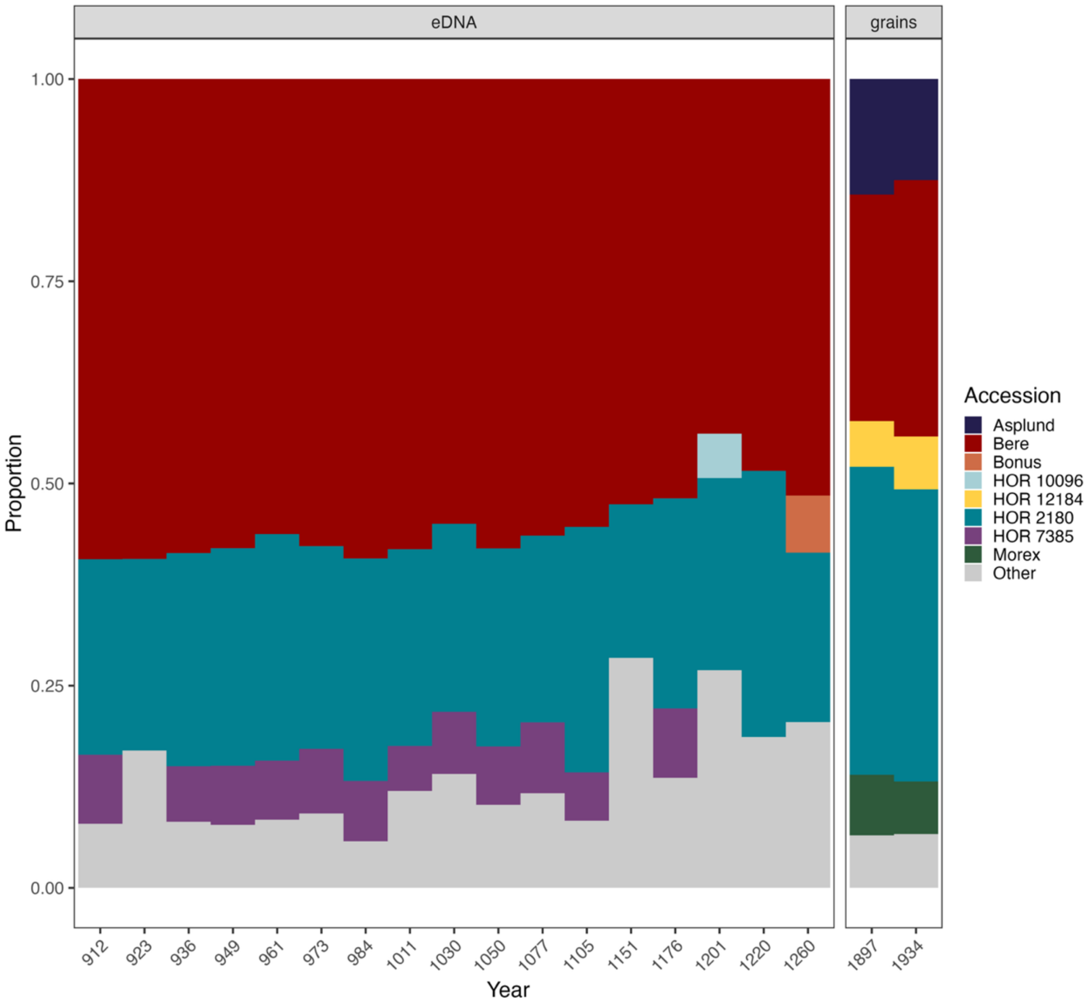

**Figure S47. Genetic ancestry of Icelandic Viking Age barley from sedimentary eDNA and historical herbarium grains.** Estimated ancestry proportions for barley accessions recovered from the Tjörnin sediment core (left panel, 912–1260 CE) and two herbarium specimens collected in 1897 and 1934 (right panel). Proportions were inferred using an expectation–maximization algorithm applied to short reads competitively mapped to 78 reference barley genomes. The eDNA signal is dominated by Bere (red) with consistent contributions from HOR 2180 (blue) and minor fractions of other accessions. In contrast, the herbarium grains exhibit a similar Bere-dominated profile but additionally contain a distinct Asplund component (purple), supporting historical import of Scandinavian barley varieties.

### Prokaryotic Components of the Metagenome

**Microbial taxonomic profiling**

We reconstructed the microbial taxonomic profiles following the ancient metagenomic analysis workflow described in Fernandez-Guerra and colleagues [92](https://github.com/GeoGenetics/2025-

kapk-microbial). Briefly, after an initial quality control reads were extended with the kmer-based assembler Tadpole in BBTool[93] with strict parameters, and then dereplicated using VSEARCH –fastx-uniques[94]. Information from the dereplicated extended reads was used to filter the original quality-controlled reads, which were then used for downstream analyses. Filtered reads were mapped against a curated, de-replicated, custom reference database using Bowtie2[95]. For the reference database construction, we sourced data from NCBI[32], GTDB v207[96], PhyloNorway pastid sequences[13] and IMG/VR v4 genomic data[97]. Additionally, we incorporate metagenome-assembled genomes from TARA Oceans[98], Genomes from Earth's Microbiomes[99], the glacier microbiome catalogue[100] and genomes recovered from a permafrost study[101]. Mapping duplicates were then removed with MarkDuplicates (Picard Toolkit). Multi-mapping reads were resolved using the "reassign" function in filterBAM via an Expectation-Maximisation approach with the parameters -i 0 -a 92 -n 3. The resulting BAM file was subsequently filtered using the "filter" function in filterBAM with the following parameters: -N -e 0.6 -n 3 -A 94 -a 92 --min-breadth 0.01 --min-expected-breadth-ratio 0.75. The "lca" function in filterBAM was used to estimate Lowest Common Ancestor (LCA) genome-normalized taxonomic abundances. These abundances are based on the 80% truncated average depth (TAD80). The number of reads in this region are estimated with the formula N = (G*C)/L, where G represent the length of the TAD80 region, C is the coverage in this region and L is the average length of reads mapping to the reference. Then, taxonomic abundances are estimated by normalizing the number of reads (N) by the length of the TAD80 region (G) and scaled by 1 million. Post-mortem DNA damage was estimated with metaDMG[102] in LCA mode (Figure S48).

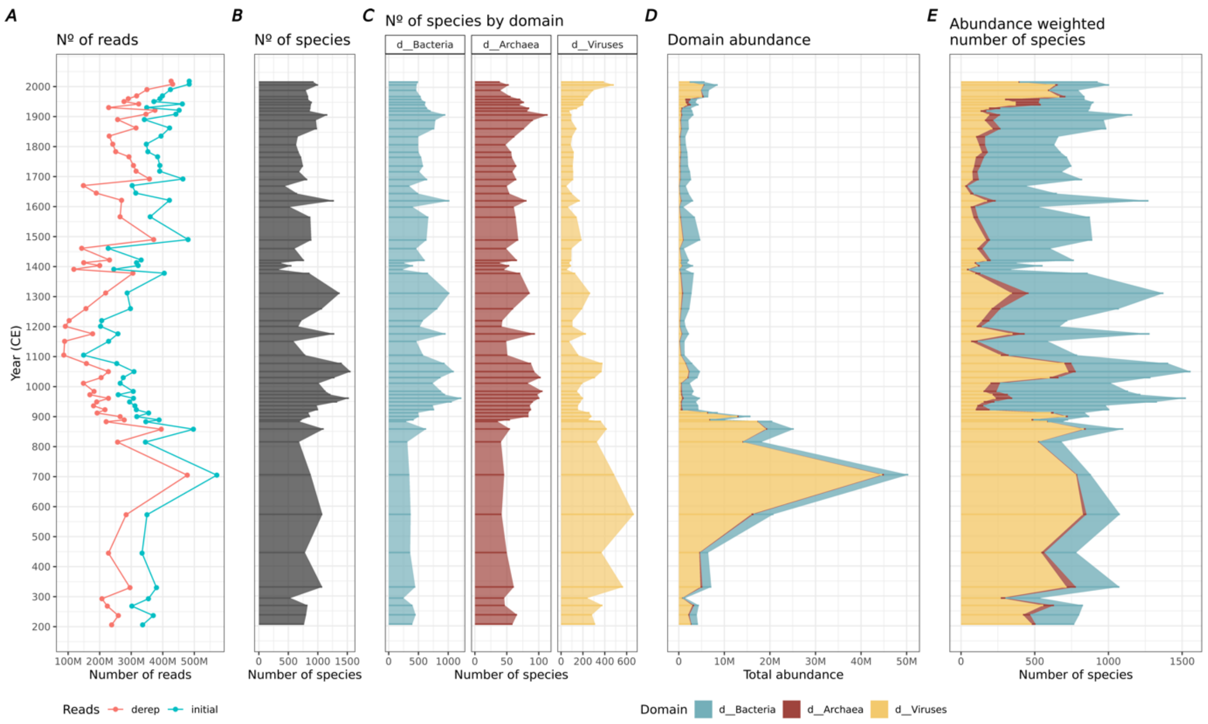

**Figure S48. A) Number of initial and dereplicated number of reads per sample.** B) Total number of observed species per sample and C) number of observed species by microbial domain.

D) Domain abundance per sample and E) abundance-weighted contribution of each domain to the total number of observed species.

**Dataset filtering**

For ordination-based analyses and reference-based metabolic reconstruction only the archaeal and bacterial fractions were considered. After the viral fraction removal, the remaining dataset was filtered to exclude potentially modern communities by removing (i) widespread taxa detected in more than 50% of the samples and with little or no damage ($A\_b < 0.05$) in any of the observations together with (ii) taxa with low (<1.5) abundance variance coefficient. Species with very low abundance (mean abundance < 1e-4) and occurrence (detected in less than 4 samples) were also excluded.

**Ordination-based analyses**

Bray-Curtis dissimilarity was calculated for each pair of samples after Hellinger standardization[103] and the resulting distance matrix was used for the non-metric multidimensional scaling (NMDS) ordination. Additionally, Bray-Curtis dissimilarities were square-root-transformed to perform hierarchical clustering with Ward's (ward.D2) method. The optimal number of clusters was identified by maximizing the average silhouette width after sequentially test over increasing number of clusters. Indicator species[104] were identified for each cluster of samples, not allowing multiple groups combinations and selecting species with high statistical significance (p-value < 0.01) and indicator value (stat>0.5).

The unsupervised clustering of samples based on microbial community composition revealed ten distinct clusters that accurately recapitulate the major transitions in lake history (Figure S49). Indicator species analysis identified idiosyncratic microbial assemblages associated with each cluster, representing variable proportions of the overall community (Figure S50). Some taxa served as strong indicators of past conditions and environmental transitions in the region. The first cluster resembled a rich marine-specific microbiome, with bacterial genera adapted to cold marine environments, including *Polaribacter*, *Psychrobacter*, and *Psychromonas*, alongside other marine-associated taxa such as *Poseidonibacter* and *Methyloceanibacter*. The transition to freshwater conditions is evident in the second cluster, which contains freshwater and brackish representatives of the cyanobacterial genus *Dolichospermum*. Human arrival and settlement are reflected in the increased abundance of taxa adapted to detritus-rich environments, such as representatives of the order Sedimentisphaerales (phylum Planctomycetota), suggesting a higher input of organic matter likely driven by anthropogenic activity in the catchment area. Additionally, specific host-associated archaeal and bacterial species were detected during this phase (Fig. S51), most notably *Methanobrevibacter*, a methanogenic archaeon with representatives isolated from the gut microbiomes of cattle, goats, sheep, and other domesticated animals. This approach also resolved fine-scale temporal patterns during the most recent phase of urbanization and industrialization. The first cluster within this episode suggests a period of elevated primary productivity, potentially linked to a high nutrient influx, followed by a second cluster characterized by taxa associated with anaerobic processes. Finally, a distinct event is marked by a sharp increase in wastewater-associated taxa, particularly representatives of the genera *Rubrivivax* and *Accumulibacter*.

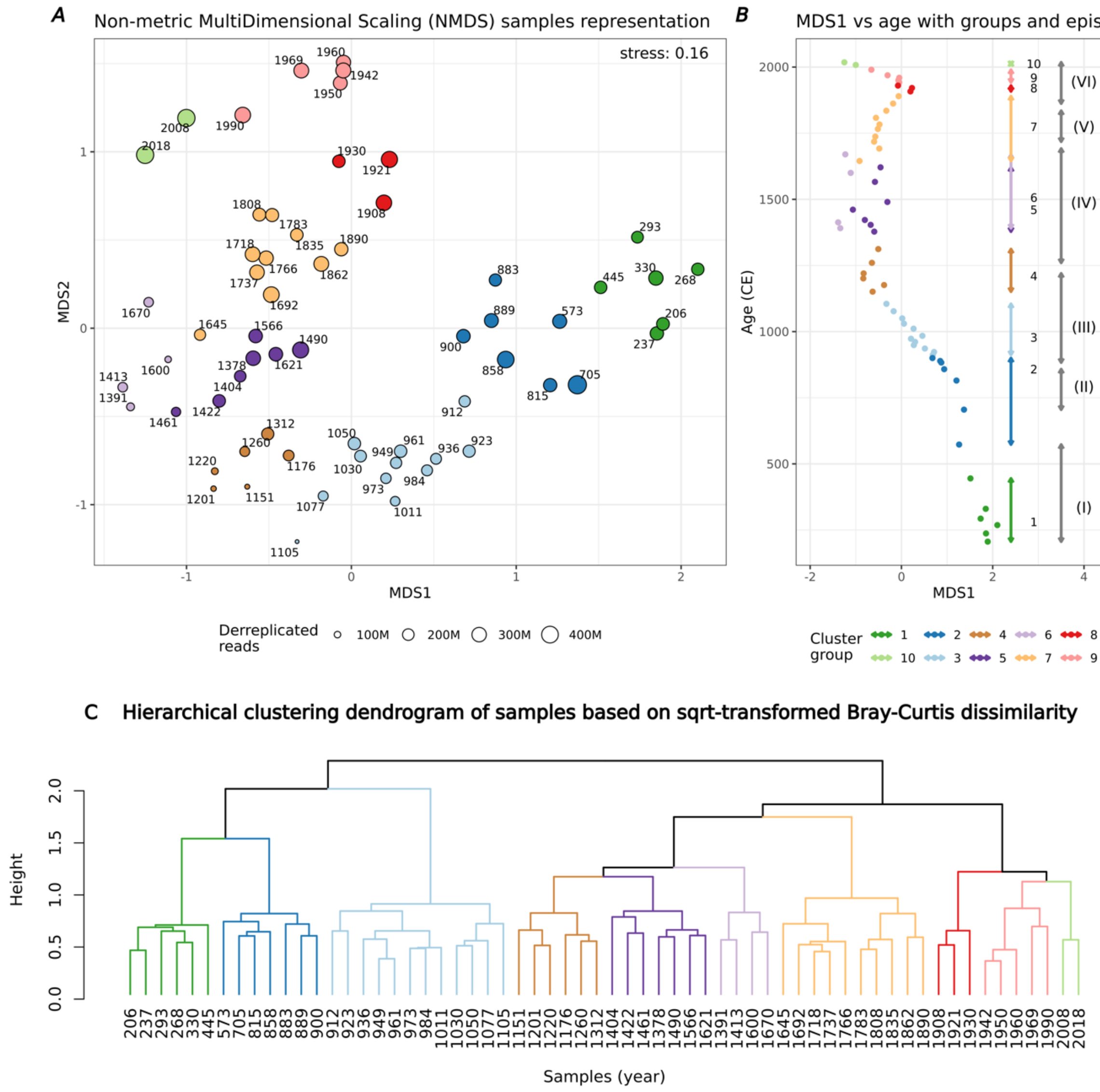

**Figure S49. A. Non-metric multidimensional scaling (NMDS) samples, ordination based on pairwise Bray-Curtis dissimilarities.** B. MDS1 scores plotted against sample age. Coloured vertical lines indicate the temporal span of each of the 10 identified sample clusters, while grey lines indicate the span of previously described episodes. C. Hierarchical clustering dendrogram based on square-root-transformed Bray-Curtis dissimilarities.

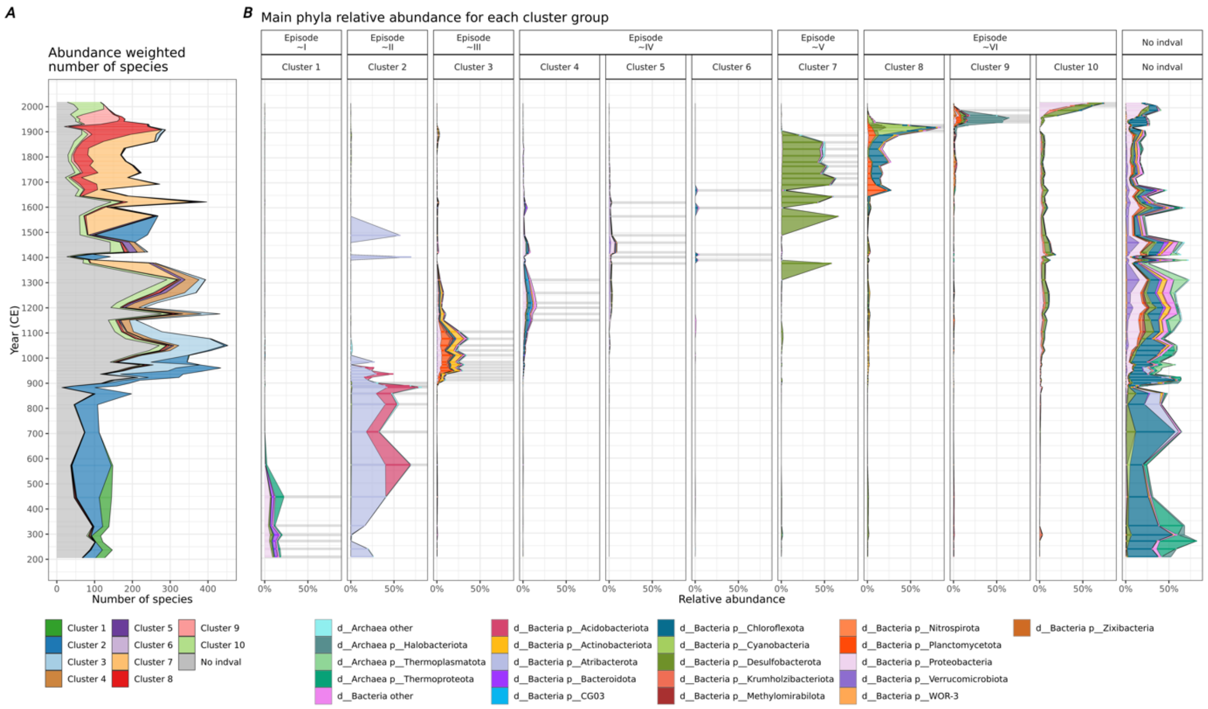

**Figure S50. Contribution of indicator species from each sample cluster over time.** A. the abundance-weighted proportion of the total number of observed species, and B. their relative abundance. Horizontal grey lines indicate samples assigned to each cluster of samples.

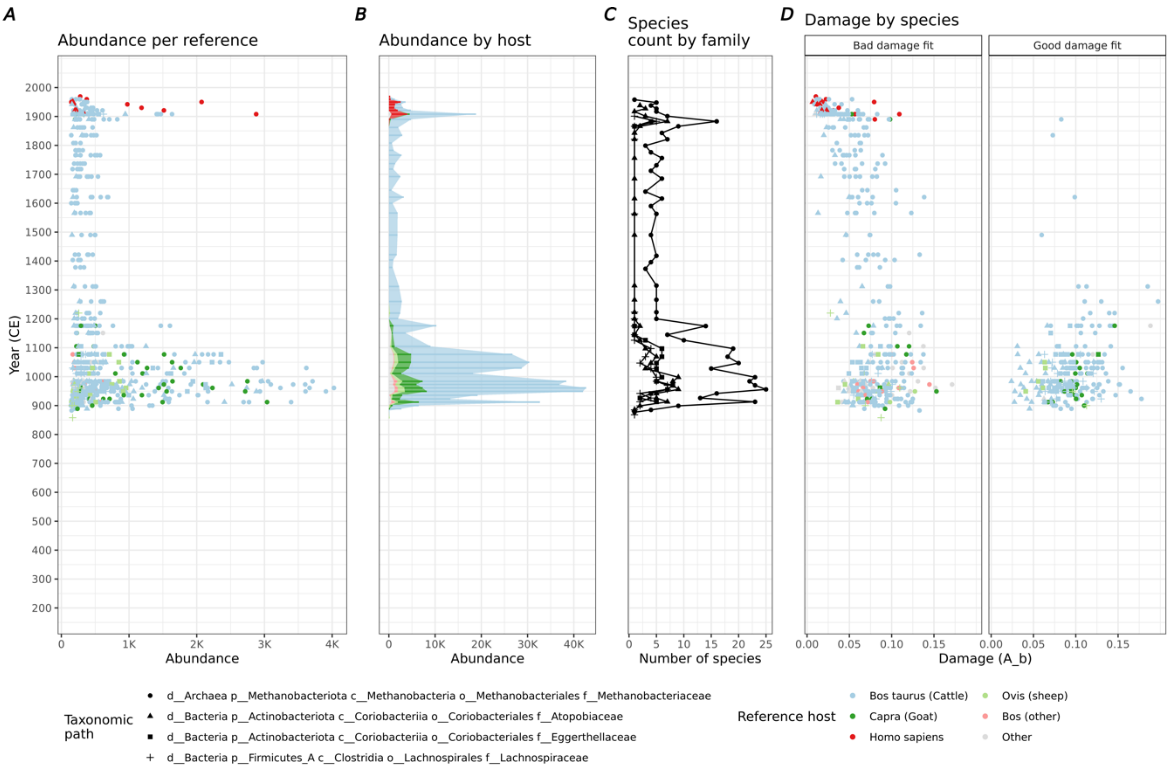

**Figure S51. Detection of potential gut microbiome taxa in selected mammalian hosts.** Potential hosts were assigned based on the isolation source of the reference genomes. A) Relative abundance of observed subspecies (reference genomes). B) Total abundance grouped by host species. C) Number of observed species per host family. D) DNA damage estimates for each observed subspecies.

#### Reference based metabolic reconstruction

We estimated the metabolic potential of the archaeal and bacterial reference genomes using the Anvi'o tool Anvi-estimate-metabolism[105,106], run with default parameters. We targeted metabolic pathways involved in the nitrogen (Fig. S52) and sulphur (Fig. S53) cycles. Nitrogen-related pathways included assimilatory nitrate reduction, nitrification, nitrogen fixation, denitrification, and dissimilatory nitrate reduction. Complete nitrification and anammox were also targeted but not detected. Sulphur-related pathways included dissimilatory sulphate reduction, assimilatory sulphate reduction, sulphate–sulphur assimilation, and thiosulfate oxidation via the SOX complex.

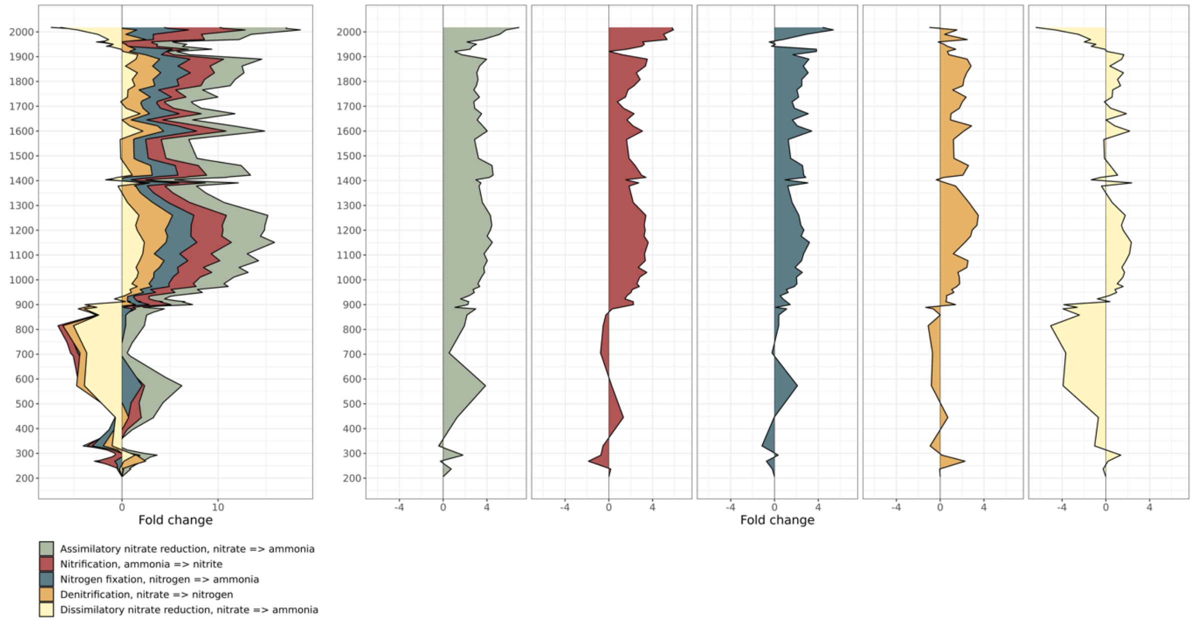

**Figure S52. Log2 fold change in relative abundance of taxa exhibiting Nitrogen-related metabolisms according to the reference-based metabolic reconstruction.** Targeted KEGG modules are Nitrogen fixation (M00175), Nitrification (M00528), Denitrification (M00529), Dissimilatory nitrate reduction (M00530) and Assimilatory nitrate reduction (M00531). Modules for Anammox (M00973) and Complete nitrification (M00804) were also targeted but not detected.

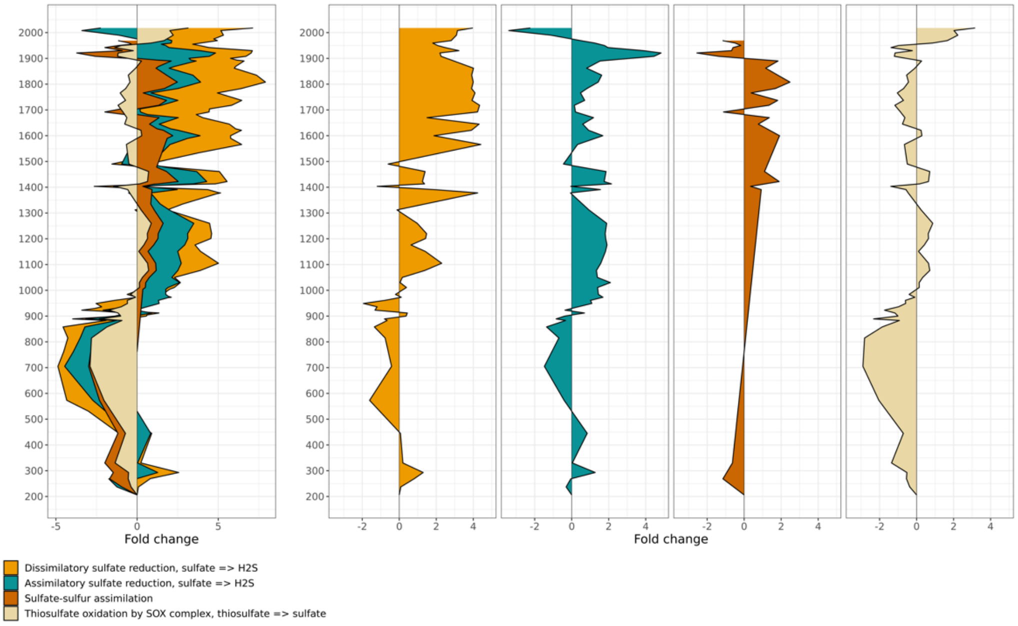

**Figure S53. Log2 fold change in relative abundance of taxa exhibiting Sulphur-related metabolisms according to the reference-based metabolic reconstruction.** Targeted KEGG modules are Dissimilatory sulphate reduction (M0596), Assimilatory sulphate reduction (M00176), Sulphate-sulphur assimilation (M00616) and Thiosulfate oxidation by SOX complex (M00595).

**Microbial source tracking**

We used getBiomes (https://github.com/aMG-tk/get-biomes) to retrieve biomes metadata and associated raw sequence data from MGnify and the European Nucleotide Archive (ENA) to generate the source dataset for sourcetracking analysis. The tool was run with the following parameters: --ena-filter {library_layout: PAIRED, library_strategy: WGS, library_source: METAGENOMIC, library_selection: RANDOM, read_count: 10000000} --combine --exclude-terms human,16S. The retrieved raw sequences were processed using the same pipeline applied to ancient data, with settings optimized for modern metagenomic data. This included quality filtering with fastp, selection of the forward read for unmerged pairs, no reads extension, and taxonomic annotation with a 95% ANI threshold. Post-mapping steps were identical to those used for the ancient datasets, with the exception that post-mortem damage was not assessed. The same database and pipeline were used to annotate, filter, and estimate taxonomic abundances for both the sources and sinks. The resulting sources and the filtered dataset analyzed in this study were used as a input for meta-Sourcetracker[107] with the following parameters: --sink_rarefaction_depth 0 --source_rarefaction_depth 0 --per_sink_feature_assignments --restarts 100 --draws_per_restart 5 –diagnostics.

The obtained results reproduced several previously observed patterns (Fig. 4 B). The marine signal was clearly recovered during the earliest phase, while the high input of organic matter associated with human arrival and settlement was captured by an increased contribution of soil-associated taxa. Likewise, the signal of mammal-associated taxa was also detected during the settlement phase and the initial occupation of the catchment area.

**The ancient microbial record**

The analyses undertaken for the recovery and identification of the ancient microbial record are described in Methods. Extended Data Fig. 9 shows by symbol size the number of reads aligned passing all filters, with read ANI (Average Nucleotide Identity) indicated by symbol fill colour. The taxa are arranged by their occurrence over time in the core, with Extended Data Fig. 9 displaying them from left to right by types that occur only in the centuries prior to human arrival, those that occur only in the centuries after human arrival, and taxa that are present throughout the record.

Thus, in the lowest levels of the Tjörnin core when this was still a marine embayment, display a rich marine-specific microbiome. We detected bacterial genera adapted to cold marine environments, including *Polaribacter*, *Psychrobacter* and *Psychromonas*, alongside other marine-associated taxa such as *Poseidonibacter* and *Methyloceanibacter*. Source-tracking approaches further support the marine origin of both prokaryotic and viral assemblages (Fig. 4). Among the 61 microbial species detected using the pathogen- and fungal-specific workflow, seven (11%) were exclusively discovered before the formation of Tjörnin (i.e., before 600 CE), these include pathogenic bacteria that target marine fish and animals, e.g., *Vibrio splendidus* (Atlantic salmon, cod and other fish, crustaceans, and shellfish) and *Vibrio cyclithrophicus* (shellfish larvae) (Fig. 3A; Extended Data Fig. 9).

As the Tjörnin basin was gradually isolated from the sea and steadily freshened, we see a corresponding change in the microbial species record. The increase in freshwater along with an increase in temperature combined to support the presence and persistence of *Mitosporidium daphnia* dating from approximately 700 CE until about 1250 CE. This parasite infects the planktonic crustacean *Daphnia*, which thrives in freshwater at temperatures higher than 11°C, although some strains can tolerate brackish water (Extended Data Figs. 2, 9).

One alga (*Aureococcus anophageefferens*) and 23 microbial species (39% of all identified species using the pathogen- and fungus-specific workflow) were not significantly affected by changes in salinity and existed both before and after human arrival. However, five of these (5/23, 22%) were affected by temperature changes, disappearing at the start of or during the first half of the warm period. Among these, three were pathogenic fungi capable of infecting various types of grasses (*Alternaria arborencens*, *Pyrenophora tritici-repentis*, and *Parastagonora nodosum*) possibly reflecting the above-described changes in grass taxa, or larch and poplars (*Melampsora larici-populina).* The fourth was a lichen-associated bacterium first identified in Antarctica (*Burkholderia* sp. PAMC 28687). Among the remaining eighteen microbial species, some were non-pathogenic bacteria or fungi associated with marine, plant, and soil environments. However, we also identified pathogenic species, including the occasional presence of the bacterium *Vibrio anguillarum*, which can cause disease in fish and shellfish, as well as gastroenteritis in humans – though we note this taxon occurs only rarely after Landnám (Extended Data Figs. 3, 9).

With the arrival of Norse on Iceland, the microbial species analysis offers direct, unequivocal, and continuous evidence of human presence from approximately 900 CE and onwards. This is demonstrated through the detection of the yeast *Debaryomyces hansenii*, which is a component of the human gut microbiome, and *Staphylococcus sp. AntiMn-1*, which is associated with the nose and skin microbiome (Extended Data Fig. 9). Although these taxa can be found in many habitats and not just human gut – including sea water – neither of those taxa are present in the pre-600 or 600-900 sections of the core, as they ought to be if they were marine in origin. Nor are they in the 'before and after human arrival column' if they were from humans and seawater (Extended Data Fig. 9). We also continuously detect *Bacillus sp. 1s-1* from about 900 CE; this bacterium could originate from the gastrointestinal tract of humans, animals (primarily ruminants), and/or marine sponges, suggesting that imported human and animal microbial species might affect animals residing within the lake.

After Landnám, the detection rate of certain plant-associated microbial species significantly increased. For example, *Botrytis cinerea* is rare before human arrival, but this necrotrophic fungus which can infect wild and domesticated plants including berries and vegetables, occurs almost continuously after human settlement. This pattern may be linked to the increased cultivation of susceptible host plants. Likewise, plant cultivation is a potential cause for the abrupt increase in the detection of *Pseudomonas syringae* i.e. the cause of canker. A similar rise in detection frequency was observed for *Pseudomonas aeruginosa*, as well as for ten other *Pseudomonas* species that were present before human arrival, and *Clostridium perfringens*. Both *Pseudomonas aeruginosa* and *Clostridium perfringens* are significant pathogenic bacteria that can lead to various infections in animals and humans, including gastroenteritis and life-threatening sepsis (Extended Data Fig. 9). Similarly, human arrival coincided with the sustained abundance of plant symbionts, notably, of the arbuscular mycorrhizal (AM) fungus *Rhizophagus irregularis* (phylum Glomeromycota, Ext. Data Fig 9). *R. irregularis* presence predates human arrival (c. 800 CE) as Tjörnin transitioned into a landscape with increasing abundance of terrestrial plants. Its continued abundance until 1200s might be linked to farming, with the increased cultivation of cereal crops such as wheat and barley and other host species (e.g. fruit trees) for AM fungi. Engaging in AM symbiosis provides mineral nutrients to the plant host and is also known to improve plant stress tolerance and resistance to other microbial pathogens[108]. Notably, *R. irregularis* is a generalist, cosmopolitan AM fungi species considered to thrive in disturbed, agricultural soils. This altogether supports an emergence coincident with the increase in agriculture in Tjörnin.

Moreover, we identified 28 additional host-associated microbial species (Fig. 4; Extended Data Fig. 4), including the fungi *Phialocephala scopiformis*, an endophyte of conifer needles, and *Hyaloscypha bicolor* (part of the *Rhizoscyphus ericae* aggregate), which primarily forms ericoid mycorrhizal symbiosis with Ericaceae plants like *Calluna* (heather) and blueberries. This reflects the importance of ericaceous pioneer plants and their associated mycorrhizal symbionts in the soil types of Iceland.

The microbial record shows a sudden and continuous presence of several Aspergillus species following human settlement, particularly *Aspergillus fumigatus*, a fungus found in soil and compost heaps, and *Aspergillus nidulans*, associated with compost and stored seeds (Extended Data Fig. 9). Moreover, we detected *Penicillium rubens*, a soil-dwelling fungus that often grows on vegetables and decaying vegetation; this signal is particularly strong during the warm period, perhaps reflecting temperature-induced farming choices and practices over time. All these fungi thrive in damp environments and can cause indoor mould. While *Aspergillus fumigatus* is the most common cause of the disease Aspergillosis (a respiratory infection), *Aspergillus nidulans* and *Penicillium rubens* are opportunistic pathogens in humans. The sudden appearance of these and other new microbial species suggests that the settlers' subsistence lifestyle had a significant impact on the pathogenic challenges they faced and on local microbial diversity.

As climatic conditions changed over time, accompanying the decline of barley, and perhaps more broadly of agriculture, there is a corresponding decline in detection of the root fungal symbiont *R. irregularis* (Ext Fig 9) which forms a symbiotic relationship with barley and other plants, and serves to improve nutrient uptake. It drops below and never recovers to levels seen prior to 1200 CE, providing strong correlation between its detection and human agriculture in that period. As AM fungi are obligate biotrophs that require lipids provided by plant hosts to complete their lifecycle, their co-occurring decline befits the shifts in human activities in that period.

**Viral source ecosystems**

To assign an ecosystem of origin to the viral genomes included in our reference database, we used the ecological metadata available in IMG/VR v4[97]. This database integrates metadata from the Genomes OnLine Database (GOLD) to provide standardized ecosystem classifications for uncultivated viral genomes (UViGs) derived from globally distributed metagenomic datasets. This approach also demonstrated that viral communities reproduced the previously reported patterns with even greater clarity and resolution (Fig. 4 A). The transition from marine to freshwater conditions was distinctly recovered, as was the increased contribution of terrestrial taxa following human arrival. Finally, the fine-scale transition observed during the most recent episode—already identified through clustering analysis—was also captured by the viral source-tracking approach.

**Viral community diversity**

To test if virus and host community diversity were changing across the different episodes, we performed NMDS analyses based on Bray-Curtis distance with PERMANOVA using the R package vegan, followed by a pairwise comparison between groups using the R package rvaidememoire. In both cases 999 permutations were considered, and for the pairwise comparison Pillai test with False Discovery Rate for *p*-value correction was used. Next, to identify the individual viruses significantly contributing to the changes across episodes, an indicator species analysis was performed using the multipatt function from the R package indicspecies with 999 permutations (Fig. S54).

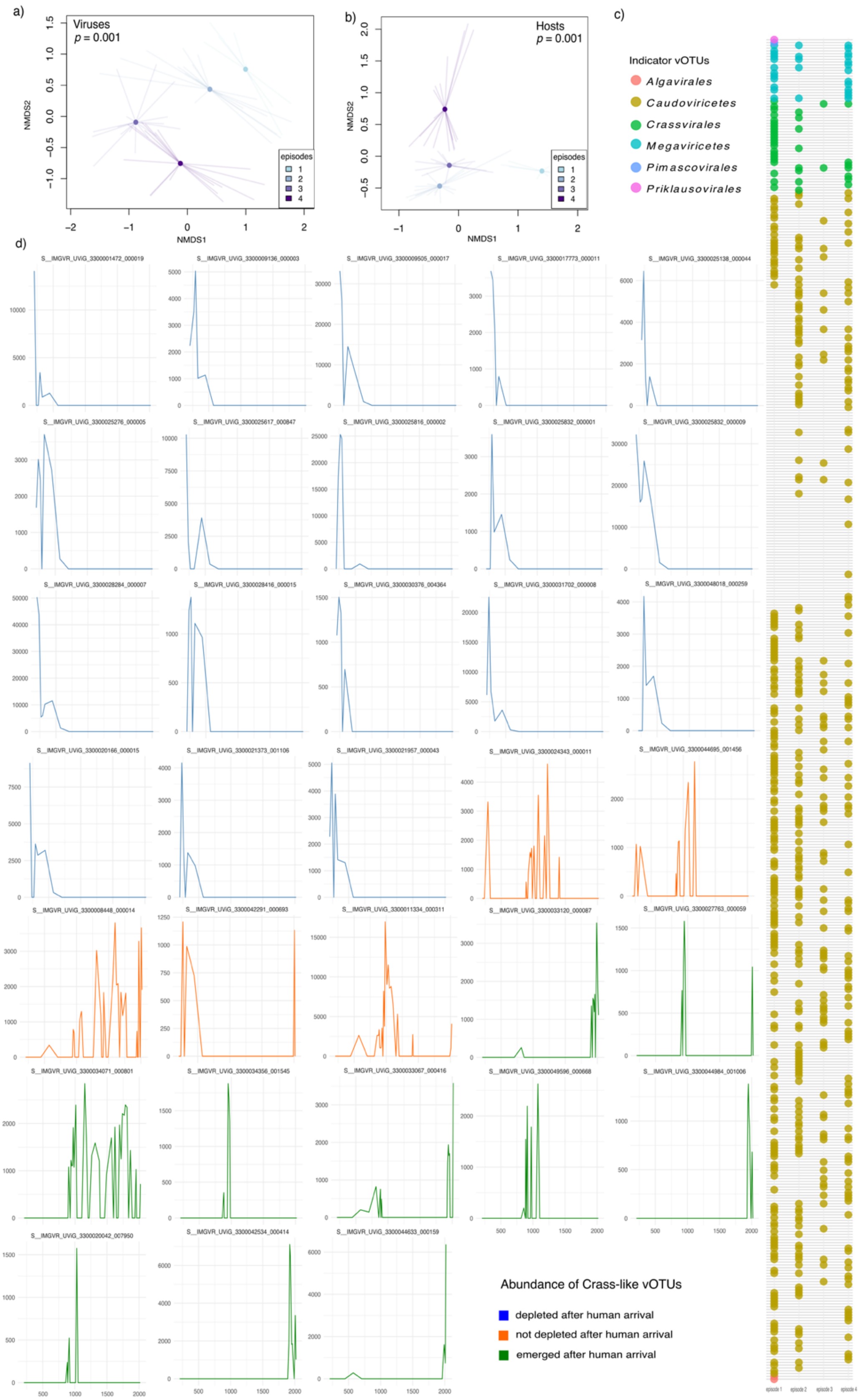

**Figure S54. Shifts in virus community diversity.** In total, 1,084 viral operational taxonomic unities (vOTUs) were detected in the data set. They were classified within the phyla Uroviricota

(95.29%), Nucleocytoviricota (4.33%) and Preplasmiviricota (0.33%). The vast majority were viruses of Bacteria and Archaea belonging to the class Caudoviricetes (95.29%), with 4.88% of these classified within the order Crassvirales. The remaining 4.71% were classified as members of the class Megaviricetes (4.33%), including the orders Algavirales (0.46%) and Pimascovirales (0.09%) that infect single-celled eukaryotes and animals, and the class Maveriviricetes (0.37%) that co-infect hosts with Megaviricetes. Non-metric multidimensional scaling (NMDS) analyses with PERMANOVA and pairwise comparison showed that both a) virus and b) microbial host communities were significantly different across the four episodes ($p < 0.01$). In the NMDS analyses points represent centroids that are connected to the individual observations by the lines. c) Indicator species analysis detected 482 vOTUs that were significantly associated to the environmental changes that marked the different episodes ($p \leq 0.05$). Each line represents an individual vOTU that was detected as indicator and the points, coloured by taxonomy, indicate the episode these vOTUs were associated with. This analysis detected 33 vOTUs that were classified as Crassvirales. Given their relevance as key members of the human gut microbiome, d) the abundance (y-axis) of each of these Crassvirales clades were represented across time (x-axis). The species name of the reference crAssvirus clade detected is indicated at the top of each plot.

#### Historical, Archaeological, and Demographic Context

**Estimate of the Icelandic population**

The earliest firm indication for the pre-modern population of Iceland is the 1703 census, which counted 50,140 people[109]. The census was taken following a prolonged period of demographic decline and age-models of the 1703 population suggest that it had been reduced by as much as 10% since the late 17th century[110]. Counts of farm units in c. 1100 (4560), 1311 (3812) and 1703 (4024) give a sense of long-term stability but may mask fluctuations in the numbers and sizes of households. There are two main views of the development of the pre-1703 population[111]. One postulates population growth until the 14th century, cut short by the 15th century epidemics and a subsequent inability to reach the pre-1400 maximum. The other sees general stability characterised by intermittent fluctuations on the scale of +/- 10% and a major downturn in the 15th century, followed by full recovery to pre-plague levels. The latter view is represented in the graph. There are also differences of opinion about population growth at the beginning of settlement. A traditional view[112] sees a small founding population with incremental growth only reaching full carrying capacity in the 12th century. A contrasting view[113] argues for a very large founding population and sharp initial population growth reaching the 50.000 level already in the 10th century. This latter view is represented in Figure 5E. Data from after 1700 CE were retrieved from https://statice.is/statistics/population/inhabitants/overview/.

**Summary of the archaeological evidence from Reykjavik area**

Despite the celebrated sagas of Iceland that recount its settlement (Landnám) in the 9th century, the earliest of the these was not written until several centuries after Landnám[114]. Hence, those several centuries are part of Iceland's prehistory and understanding this period relies upon archaeological and palaeoecological material. Fortunately, there have been numerous archaeological excavations in the central downtown area (Miðbær) of Reykjavík over the last fifty years (Fig. 1). Material pertaining to archaeological excavations is drawn from both published work and the grey literature (e.g. contract excavation reports). In some instances, animal and plant assemblages have yet to be fully analysed and published, and it is likely that this review will not be comprehensive. Furthermore, no consideration is given here to the abundance of the taxa identified in the assemblages, nor is there any attempt to interpret hunting or agricultural equipment, or items used to process foodstuffs e.g. querns. Similarly, while palynological data might be referred to in relation to Tjörnin and Vatnsmýri[5,80], it is not considered in depth.

In addition to the archaeological evidence, there are two historical texts that are important to this review process: The Icelandic land register known as *Jarðabók*, compiled between 1702 and 1714 CE[115], and the *Diplomatarium Islandicum* (DI, I–XVI)[116], a sixteen-volume collection of letters, receipts, rulings, reports, inventories and ledgers etc. relating directly to Icelandic affairs from the 10th century until the 16th century. Other sources of material will be utilized to supplement the information gleaned from Jarðabók and DI. These include descriptions of Iceland written by Icelandic and foreign geographers/travellers in the 18th and early 19th centuries, such as Ólafsson, Hooker and MacKenzie[117–119].

**The archaeology of Reykjavík**

It is estimated that there were five longhouses on the north bank of Tjörnin following Landnám[120]. Four of these households were concentrated around the nexus of modern Aðalstræti, Tjarnagata and Suðurgata, as reviewed by Sverrisdóttir[121]. There was also an associated area of workshops, smithies and kilns closer to Tjörnin[122]. The discovery of an early household at Lækjargata challenges the conventional understanding of human settlement in Reykjavík, expanding the area under occupation, perhaps even implying a proto-urban context[120].

Mammals and birds recovered in zooarchaeological assemblages from Reykjavík are tallied in Appendix 1 & 2. One of the most striking features of the zooarchaeological assemblages is the disappearance of the walrus (*Odobenus rosmarus*) and great auk (*Pinguinus impennis*) from archaeological faunas by 1200 CE. Walrus disappearance has been attributed to over-exploitation by human settlers[123]. In fact, its presence in Iceland and its potential as a source of ivory is thought to have been a reason for the colonisation of Iceland in the first place. Regarding the auk, there were no suitable nesting cliffs for this species in the Reykjavík area, and in comparison with butchering patterns of birds at other sites in Iceland, Amorosi suggests that its presence (as well as that of other Alcids) in the faunal assemblages of Reykjavík is probably a consequence of exploratory/hunting trips off the coast of southwest Iceland and thus part of a more general utilisation of seabirds[124,125] which survive in Iceland to the present[126]. Species that are preyed upon include *Cepphus grylle*, *Uria aalge*, *Alca torda*, *Fratercula arctica*, *Phalacrocorax carbo*, known collectively as *svartfuglar*. Other edible species include *Fulmaris glacialis*, and potentially, *Morus bassana*, which is still eaten in the Outer Hebrides.

The only seal species that breed on Icelandic shores are the harbour seal (*Phoca vitulina*) and the grey seal (*Halichoerus grypus*), with the former being most common[127,128]. The harp seal (*Pagophilus groenlandicus*) has been observed near Reykjavík in Faxaflói[129], while the ringed seal (*Phoca hispida hispida*), bearded seal (*Erignathus barbatus*) and hooded seal (*Crystophora cristata*) have all been recorded in the seas around Iceland[129]. All belong to the family Phocidae. Of these taxa, *P. vitulina* and *E. barbatus* have been identified in Reykjavík faunal assemblages, the remainder are simply identified as Phocid. Seals were hunted in a variety of ways in the past in Iceland i.e. clubbed on sea ice, clubbed or trapped on the shore, caught in nets, speared from boats, and later, shot with guns[130].

For the Cetacea, in only one instance is there an attempt to attribute species i.e. *Balaenoptera borealis* (with reservations). Cetacea bones, especially those of large whales, are probably derived from whale wrecks. It is possible that pods of small whales were driven into shallow bays, while individual small whales might have been speared from a boat (Kristjánsson 1980). The lack of whale bone in post 17th century Icelandic sites may be due to the collapse of North Atlantic whale populations driven by industrial whaling in Europe and America in the 18th and 19th centuries. The presence of arctic fox (*Vulpes lagopus*) in the assemblages seems slightly late (post-1226 CE), given that it is native to Iceland. Its presence in faunal assemblages might suggest the utilisation of the animal for fur.

Overall, there is a sense that early settlers in Iceland relied heavily upon wild animals and birds for subsistence while their domestic herds became established and even continued to do so for some time afterwards[131].

Of the domestic animals, cattle (*Bos taurus*) and sheep (*Ovis/Capra* and/or *Ovis aries*) are ubiquitous from Landnám down to 1900 CE, horses (*Equus caballus*) only slightly less so. It has been argued that pigs (*sus scrofa*) were a feature of Landnám and quickly disappeared from the Icelandic landscape in subsequent years[132]. However, in Reykjavík, the record is quite consistent into the modern period, and recent research from northern Iceland suggests that the story is more complex[133].

The Landnám hen, as its name suggests, is thought to have been introduced to Iceland at the time of settlement. This may be a modern colloquialism, as no bones of *Gallus gallus* have been found in early faunal assemblages, and it does not appear in Reykjavík until the mid-17$^{th}$ century. Of cats and dogs (*Felis domesticus* and *Canis familiaris*), only the latter are present from Landnám, with cats first appearing in the medieval period. Small dogs are often assumed to have been lap dogs, belonging to high status people[134]. Some of them may have been present to control rodents[135]. Rats (*Rattus sp.*) do not appear in the Icelandic faunal assemblage until after 1650 CE. As this is long after the two outbreaks of Plague in Iceland in the 15$^{th}$ century it has been the subject of much discussion in Iceland concerning the actual nature of the disease, vectors of circulation, and impact[136,137].

Analysis of fishbone is very limited, and most is from archaeological contexts that post-date 1500 CE. In the earliest assemblages, identification is limited to Gadid or *Gadus morhua*[138,139]. The later assemblages are far more diverse, and along with *G. morhua*, include other Gadids: *Pollachius virens* and *Melanogramus aeglfinus*, as well as *Molva molva*, *Brosme brosme*, *Hippoglossus hippoglossus* and *Anarchichas lupus* [135,140]. The difference in diversity between the assemblages is likely due to effort regarding analysis rather than any change in fishing habits. For example, it is important to note that an entire layer of fishbone at Suðurgata 3–5 (870–1226 CE) was never analysed[124,141].

Although many excavation reports for Reykjavík cite the presence of molluscs, only a limited selection have been identified to genus or species level, including *Littorina littorae*, *Balanus sp. Mytilus edulis* and *Mya sp*., dated to between 1650 and 1900 CE [135,140]. Only one excavation has analysed insect remains to genus and species level[142]. While all these taxa were present, the scarcity of detailed data reflects a lack of specialist studies.

Concerning plant remains, there are surprisingly few reports regarding excavations in Reykjavík[120,138,141,143,144]. Particular attention has been given to wood remains to understand where timber for construction and crafts was being sourced, whether imported, driftwood or native woodland[138,144,145].

Other aspects of the archaeobotanical assemblages are related to farming activities[120,141,144]. It is argued that the presence of *Hordeum vulgare* suggests that cereal cultivation was attempted during the early stages of Iceland's settlement[120,141]. Indeed, based upon historical and place name studies, cereal cultivation is thought to have persisted in southwest Iceland into the 16th century (Ólsen 1910). Both interpretations are borne out by palynological evidence[4,80,146,147]. The presence of *Avena sativa* and *Linum usitatissimum* is attributed to the seeds of these plants being accidentally introduced (archaeophytes), mixed in with the seed of *H. vulgare*, along with the seeds of other plants associated with cultivated ground e.g. *Matricharia inodora*, *Stellaria media*, and *Spergula arvense*.

In contrast, native plants are not as well represented, probably due to the archaeological context, i.e. from within structures. However, the influence of Tjörnin and Vatnsmýri is present in the form of *Potamogeton sp.*, *Menyanthes trifoliata*, and Cyperaceae. That *Empetrum nigrum* and *Vaccinium uliginosum* occur might suggest active exploitation of these native fruit bearing species. *M. trifoliata* and *Ascophyllum nodosum* may also have been used for food[120,148]. Imported fruit is represented by the find of a stone from a *Prunus sp.* at Lækjargata[120].

While no attempt is made here to synthesize the sequencing of architectural remains from the various excavations of Reykjavík, in relation to natural events that might negatively impact upon the residents living there, it is worth noting findings from Aðalstræti 14:

"… *after a certain amount of soil building a new settlement occurred during the late Viking Age or Middle Ages. This settlement was also covered by sand and gravel layer which at this time washed over the entire plot*." [141]

Here, archaeology infers two instances in the past when this site was possibly flooded, with the subsequent deposition of sand and gravel. Abandonment of the site is not permanent, and nor does it mean that the entirety of the Reykjavík area was evacuated at this time.

**From Vík to Reykjavík, 1200–1700 CE**

It is convention that Reykjavík was formerly known as Vík[149]. The earliest historical reference to Vík is from a cartulary for 1200 CE which identifies a church there (DI-XII, p. 9). Archaeologists believe that this church was located at the corner of what is now Aðalstræti and Kirkjustræti, where a physical church presence persisted into the modern period[150]. Archaeologists and historians also think that the church sat immediately north of the farm of Vík[150,151]. This places the farm on the northern shore of Lake Tjörnin, on the gravel ridge that separates the lake from the sea[5]. It is the only medieval farmstead known to have been located directly upon the shore of Lake Tjörnin according to historical sources.

In 1367 CE, the Church of St John the Apostle was described as located on the land of the farm of Vík (DI-III, p. 220). This entry states that the church had a right to all the Vík lands that extended westwards to Sel. The church also owned twelve cows. The location and extent of the land associated with the Church of St John the Apostle at Vík are reiterated in two subsequent inventories from 1379 and 1397 CE (DI-III, pp. 339–340; DI-IV, p. 109). Amid detailed descriptions of church valuables e.g. altar cloths, candlesticks, censers and the like, details of livestock remained consistent with the inventory of 1367 CE, although by 1397 CE, the church had also acquired twelve ewes.

The 1379 CE inventory provides some details on the natural assets attached to the Vík church (DI-III, pp. 339–340). There were rights to arable land on offshore islands and driftwood along much of the shore of the peninsula of Seltjarnarnes. The importance of driftwood as a resource in Iceland cannot be underestimated, and the church at Vík was securing a significant right[145]. On the landholding of Vík itself, the church asserted rights to woodland (*skog*) and a shieling (*sel*) on the Hill of Vík (*vikvrholt*). It is possible that this inventory is formulaic, that there was in fact no woodland present, here simply asserting a right should there be woodland. However, as the 1379 CE inventory is far more detailed than that of 1367 CE, it appears an effort was being made to more accurately record the resources and rights belonging to the Church of St John the Apostle at Vík. In short, it is likely that there was indeed a woodland on the landholding of Vík in the 14th century.

In terms of a specific location, *Vikvrholt* literally means Hill of Vík and it would be reasonable to assume that the woodland was on the hill closest to the eponymous farm i.e. the hill now known as Landakotshæð (Fig. S55). This hill also lies west of the Vík farmstead, in keeping with the church portion of the farm asserted by the 14th century ledgers. However, there is a prevailing view that the actual hill concerned is Öskjuhlið[150], southeast of Reykjavík, and an important landmark within the context of the Seltjarnarnes peninsula (Fig. S55). This identification involves the conflation of the shieling mentioned in 1379 CE, with a shieling named *Víkursel* (lit: shieling of Vík) cited as belonging to Vík in the 18th century Jarðabók[115]. It is by no means certain that these two shielings are one and the same, and the premise has been questioned[152]. It is perhaps best therefore, to consider the 14th century account in isolation, that it is simply referring to a shieling (and woodland) on a hill near Vík. About the character of the woodland, note that downy birch (*Betula pubescens*) is the only native tree species thought to have formed a woodland habitat-type in Iceland prior to the 19th century[44].

References to Vík in the late 15th and early 16th centuries are primarily concerned with the sale or exchange of property. The documentation details only the location and something of the extent of the landholdings, and nothing of the livestock or natural resources available to the landowners (DI-VI, p. 127 & pp. 610–611; DI-VII, p. 458 & p. 754). The latter point also applies to another church inventory from 1548 CE (DI-XII, p. 129), but in this instance, rental payments in butter and whey (to preserve foodstuffs) point to livestock, while a further payment in fish implies active pelagic fishing for the first time at Vík. Other items are listed that also suggest fishing activity e.g. fishing line, hooks, a container for hooks, and leather and presumably waterproof clothing. Arguably, there is a sense of a shift from the land to the sea in terms of economic activity, reiterated further by a ledger account from 1552 CE detailing a payment of five hundred fish to the Danish governor at Bessastaðir (DI-XII, p. 418). The species of fish are never named in these accounts, but payments most likely comprised of stockfish (*skreið*) i.e. unsalted fish, dried in cold air, primarily cod (*G. morhua*), but also other Gadid species[109].

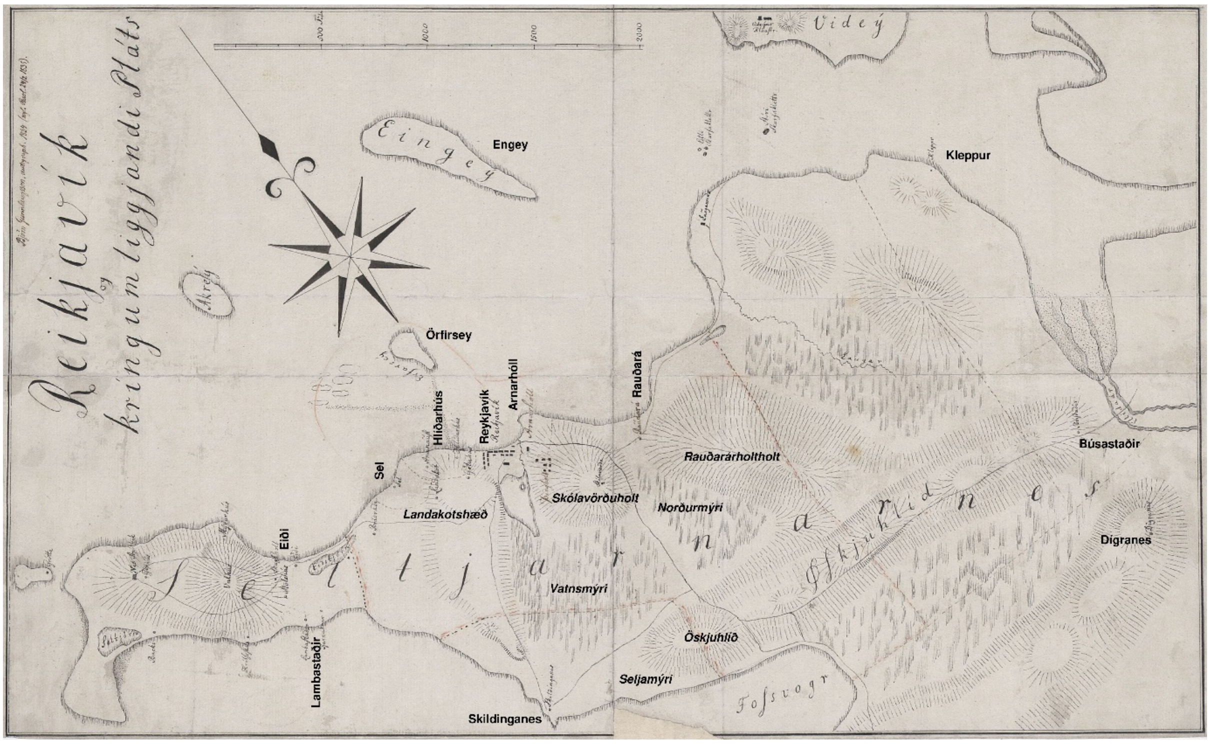

**Figure S48. Map of the District of Seltjarnarnes[153] amended to emphasize the names of the principal farms in the district and give names to the landmarks discussed in the text i.e. hills and wetlands (all spellings are in modern Icelandic).**

In 1570 CE, pastoral concerns were reasserted regarding the rental payment for ten cows (*kúgildi*). Ambiguity arises however, as a single cow can also be equivalent to six ewes[112]. It is also important to bear in mind that the inventories cited here do not represent the entirety of the farm of Vík, only the portion allocated to the church. It is not until the compilation of a land register 1686 that the *kúgildi* for the whole farm of Reykjavík can be seen[112]. At this time, the farm was divided between the Danish Crown and a private owner, with the value of fifteen cows (or equivalent) due to each.

It is also perhaps worth noting that the same land register identifies a *kúgildi* of six cows for Hlíðarhús and six cows for Skildinganes, farms that either neighbour Reykjavík or hold land overlooking Vatnsmýri[112]. Skildinganes is also mentioned in two documents from 1553 CE. One, a property grant, listed three cows (or equivalent) at Skildinganes (DI-XII, p. 524). The other, a ledger, states that the rent was paid in woollen cloth (*vaðmál*), which could imply the presence of sheep at Skildinganes rather than cattle (DI-XII, p. 577).

Overall, prior to 1700 CE, the only livestock identified at Vík are cattle and sheep, with an emphasis on cattle. Later accounts do state that the region of Gullbringussýsla generally had few sheep[154]. In terms of wider resources, there is a single instance of a woodland, while fisheries are only alluded to via payments, or through the presence of fishing equipment. Nothing that might be considered detrimental in terms of environmental impact is recorded in association with the farm of Vík. This includes an eruption known to have occurred on the Reykjanes peninsula c. 1226 CE (sometimes known as the Medieval eruption) and the Katla 1500 CE eruption[155,156]. The ash fall of both events is recorded in archaeological contexts and in the sediments of Tjörnin[5,122].

**Reykjavík and Jarðabók, 1703 CE**

Reference to Jarðabók provides the first insight into the actual complexity of the Vík landholding. By the time it was compiled in the early 18$^{th}$ century, the farm was known as Reykjavík, comprising of the home farm and nine sub-tenancies; Landakot, Götuhús, Gríota (Grjóta), Melshús, an unnamed unit, Stöðlakot, Skalholltskot (Skálholtskot), Hólakot, and a house with no land (for convenience referred to as Tómthús, as it is described in Jarðabók[115].

In relation to Lake Tjörnin, Reykjavík was on the gravel ridge to the north, Landakot on the summit of the hill to the west (Landakotshæð), Hólakot and Melshús were lower down on the western shore, and Stöðlakot, Skálholtskot were on the eastern bank (Fig. S56). Some consider the unnamed tenancy to have been a leasehold known as Suðurbær[151]. If so, it too was located on the shore of Lake Tjörnin, just south of Reykjavík farm. The remaining sub-tenancies of Götuhús and Grjóta were situated on the northern, seaward, side of the gravel ridge that separated Lake Tjörnin from the sea (Fig. S56). The specific location of Tómthús is unknown but it was probably seaward, as the residents were employed in the fisheries.

Hlijdarhús (Hlíðarhús) and the sub-tenancy of Ananaust (Ánanaust) neighboured Reykjavík to the northwest (Fig. S56). Both were once part of the Reykjavík estate. Árnarhóll and the sub-tenancy of Litla Arnarhóll lay to the northeast[115]. Neither Hliðarhús or Arnarhóll nor their subsidiaries are likely to have had a direct influence on Tjörnin. Nevertheless, their inclusion in the analysis does provide further information on the natural resources available near Reykjavík as well as environmental factors that might have had some bearing on land use practices. To the south of Reykjavík (Fig. S55), Skildinganes was comprised of a home farm and four sub-tenancies, households largely dependent on fisheries, as well as the sub-tenancy of Litla Skildinganes[115].

Details of livestock, shore rights, access to fuel, and engagement in fisheries and so on for each of these farms and tenancies are given in Table 1. It is obvious from Table 1 that cattle, sheep and horses were a daily feature of life in Reykjavík in the early 18$^{th}$ century, as a source of dairy produce and as beasts of burden. It is also clear that virtually every tenant and sub-tenant had access to, or served on a boat and was engaged in the pelagic fishery. This is also apparent from rental payments for land and livestock being met with fish in most instances. No fish species are mentioned, but as discussed, the primary product would have been stockfish[109]. Most tenants and sub-tenants in the area depended upon peat for fuel, but there is some reference to brushwood (*hrís*) i.e. Landakot, Arnarhóll and nearby Rauðará[115], all situated on, or with access, to high ground (Fig. S55). Brushwood might pertain to *Betula nana* and/or a range of ericaceous plant species.

Shore rights were the exclusive right of the home farms, with those of Skildinganes the most extensive. It is interesting to note that there were complaints that the rights of Reykjavík in relation to shellfish and lumpfish (*Cyclopterus lumpus*) were not always respected. In this case, ignoring these rights was seen to have led to the overexploitation of the species concerned[115]. That the *C. lumpus* fishery was considered a shore right, rather than part of the wider fishery, may be due to *C. lumpus* breeding and laying eggs in shallow, coastal waters[157]. Females are targeted for the roe, while males are harvested for their meat. Unfortunately, no shellfish taxa are mentioned specifically in relation to any home farm.

A small selection of edible shore plants is named in Jarðabók (Table 1). The identification of shore plants here is based upon the translation of the terms used in Jarðabók into modern Icelandic i.e. *fjaragras*, *sölvafjara* and *marálmur* are *Chondrus crispus*, *Palmaria palmate* and *Zostera angustifolia* respectively. *Murukjarna* (*Alaria esculenta*) and *biollur* (possibly *A. nodosum*), were also found in the wider vicinity[115].

Driftwood was potentially available to all, though the amount varied according to farm, and most complained of not having enough. It is worth bearing in mind that the Icelandic term *rekavon*, can include whale wrecks (Cetacea) as well as driftwood. Rights to hunting seals in Skerjafjörður were held only by Skildinganes and Reykjavík. No species are mentioned and nor is the method of hunting.

A particular feature of Reykjavík and its sub-tenancies was an obligation to assist with the movement of captured falcons (*Falco rusticola*) from the mews at Bessastaðir to the trading port known as Hólmskaupstaður near Reykjavík. It is possible, that on occasion, falcons were held at Reykjavík for short periods of time. The birds were destined for the royal mews in Copenhagen[158].

There are only two references to adverse environmental conditions in the Reykjavík area during this time including, flooding at the home farm of Reykjavík, possibly due to high rainfall or snowmelt coincident with a high tide and damage inflicted by the sea to the home field at Skildinganes, possibly due to flooding or rock debris cast ashore by the sea during storms (salt might be a further factor)[159].

The occurrence of sea ice was another problem that often beset the island, usually in the north and east, but occasionally along the south coast[160]. However, the year 1695 CE was particularly exceptional, with sea ice extending along the south coast, around Reykjanes and across Faxaflói to Borgarfjörður; thereby enclosing Reykjavík completely. This would have impacted fisheries to some degree (the sources in Jarðabók say that fishing was year-round when conditions allowed), and hampered shipping[160]. However, no reference is made to this event in the Jarðabók entries for Reykjavík and its neighbours, even though it occurred less than ten years before the land register was being compiled.

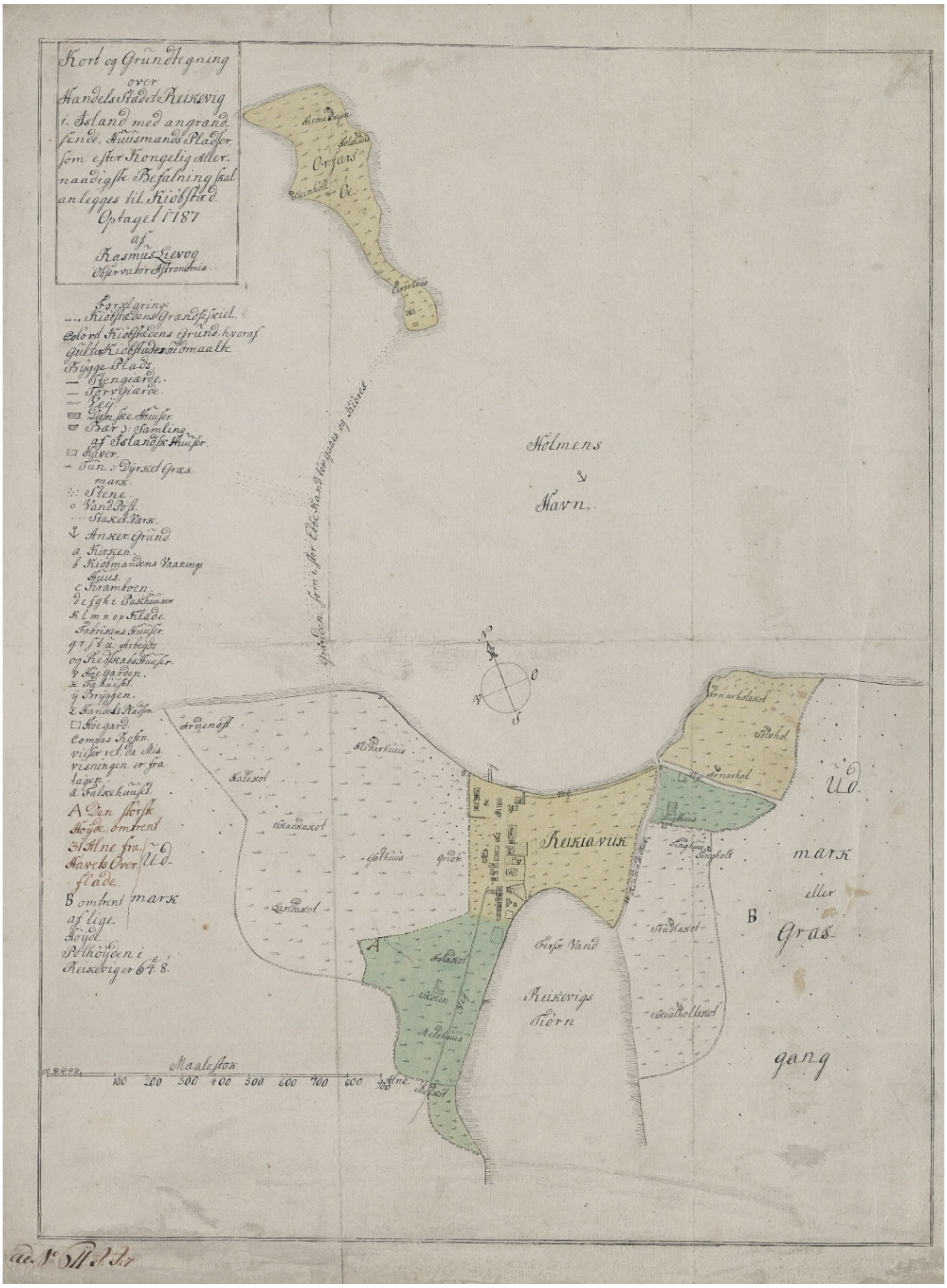

**Figure S49. Map of late 19th century Reykjavík and the locations of sub-tenancies and neighbouring farms[161].** Note that the divisions do not denote the boundaries of landholdings.

**Table S8. Livestock, shore rights, access to fisheries and fuel for the farms and sub-tenancies in the Reykjavík locale[159] (Magnússon & Vídalín 1926, Vol. III, pp. 230–270)**. About shore rights: a) *Chondrus crispus*; b) *Palmaria palmate*; c) *Zostera angustifolia*; d) Mollusca; e) *Cyclopterus lumpus*; f) Phocidae; g) driftwood. Regarding fuel: P) peat; B) brushwood.

| Farm or sub-tenancy | Cattle | Sheep | Horses | Shore rights | Fishing boat | Fuel | Falcons |
|---|---|---|---|---|---|---|---|
| Hlíðarhús | ✓ | ✓ | ✓ | | ✓ | P | ✓ |
| Ánanaust | ✓ | | | | ? | P | ✓ |
| Reykjavík | ✓ | ✓ | ✓ | b, d, e, f, g | ✓ | P | ✓ |
| Götuhús | ✓ | | ✓ | | ✓ | P | ✓ |
| Grjóta | ✓ | | ✓ | | ✓ | P | |
| “Tómthús” & sub-sub-tenant | ✓ | | | | ✓ | ? | ✓ |
| “Suðurbær” | ✓ | ✓ | ✓ | | ✓ | P | ✓ |
| Landakot | ✓ | ✓ | ✓ | | ✓ | B | ✓ |
| Hólakot | ✓ | | | | ✓ | P | |
| Melshús | ✓ | | ✓ | | ✓ | P | ✓ |
| Stöðlakot | ✓ | | | | ✓ | P | ✓ |
| Skálholtskot | ✓ | | | | ✓ | P | |
| Arnarhóll | ✓ | ✓ | ✓ | a, g | ✓ | ? | |
| Litla Arnarhóll | ✓ | ✓ | ✓ | | ✓ | B | |
| Skildinganes | ✓ | ✓ | ✓ | a, b, c, d, e, f, g | ✓ | P | |
| Litla Skildinganes | ✓ | ✓ | ✓ | | ✓ | P | |

**Reykjavík, 1750–1800 CE**

In the mid-18th century Eggert Ólafsson and Bjarni Pálsson were tasked by Danish authorities with collating a report on the nature of Iceland. The results of this research were published in the *Ferðabók Eggerts Ólafssonar og Bjarna Pálssonar 1752–1757*[117], essentially a geography of Iceland. At this time, Reykjavik farm had become the base of the Innréttingar, a development company established by Icelanders with the support of the Danish Crown[111]. The Innréttingar had two large barrels for fish (or fish oil?) stationed at the farmstead[162], and in 1752 CE Aðalstræti – a street including a row of houses – was constructed between the farmstead and the harbour area [111]. The houses incorporated workshops involved in rope-making, the tanning of hides, and the spinning and weaving of wool. A woolen mill was established in Elliðaárdalur, c. 5 km east of Reykjavik (Elliðaárdalur), and salt-panning was initiated in 1755 CE[162]; neither enterprise survived over the longer term. Reykjavik was formerly declared a town by Danish royal decree in 1786 CE[111].

According to Olafur and Pálsson[162], fish species that were actively sought by fishermen in sea off Reykjavík included cod (*G. morhua*), halibut (*H. hippoglossus*), skate (*Dipturus batis*), and plaice (*Pleuronectes platessa*). It was also known as a specific area of Iceland where basking sharks (*Cetorhinus maximus*) were hunted[162]. With regard to the southland generally, and the hot pools of Laugarnes specifically (c. 3 km east of Reykjavík), two types of eel (named by the author in Latin),*Muraena unicolor* (possibly a type of Moray eel) and *Anguilla auctorum* (European eel), were found although they were not eaten by Icelanders[162]. Harbour seals (*P. vitulina*) were said to be common in the waters around Reykjavík, and were often shot[162]. On the shore, mussels (*M. edulis*) were harvested, and it is perhaps worth noting that bird eggs were gathered on the nearby island of Viðey, as well as the down of the eider duck (*Somateria mollissima*)[162].

Along with the urban and industrial developments in Reykjavík, in 1752 CE experiments in arable cultivation and arboriculture were also initiated at the behest of the Innréttingar[162]. Conifers were introduced, but which species are unclear. Terms such as fir and spruce are used interchangeably in the narrative. The experiments with hemp and flax may have been intended for the rope-makers on Aðalstræti[136]. Potatoes (*Solanum tuberosum*) are discussed by Ólafsson and Pálsson more generally in relation to the south of Iceland, but given the context of the discussion, it may be fair to assume that they were also growing in Reykjavík at this time. Generally, Ólafsson and Pálsson felt that these experiments were successful, probably conscious of their sponsor, Frederik V, King of Denmark[162]. In truth, the cereal crops were not threshed and were fed to livestock. Experiments in tree planting failed completely, attributed to limited expertise and harsh weather.

More disastrous, and with far reaching repercussions for the entirety of the island, was the introduction of English rams to the Reykjavík area by the Innréttingar in 1750[111]. These sheep were infected with scab mite (*Psoroptes ovis*) and up to 60% of the Icelandic herd had to be culled to contain the parasite[163]

**Reykjavík, ca. 1810 CE**

The publication of the *Ferðabók* [162] drew other travellers to Iceland's shores from across Europe. Here, the accounts of Iceland by the botanist Willam J. Hooker (1811) and the geologist George S. MacKenzie (1811) are important owing to the details of their descriptions of Reykjavík. Hooker provided the first real taxonomic consideration of plants for Reykjavík[118,119].

Hooker described the countryside surrounding Reykjavík as barren, and although the land to the east was vegetated, it was strewn with boulders. He described Lake Tjörnin as surrounded by wetland, except to the north (i.e. Reykjavík), and devoid of trees[118] (Hooker 1811, p. 29). MacKenzie[119] noted that the area of Reykjavík on the banks of the stream that drained Lake Tjörnin was subject to flooding during high tides, including the vegetable gardens of the town's residents. Hooker provides some detail regarding the cultivation of garden vegetables and herbs by the locals, and MacKenzie gives details of cultivation experiments conducted by his travelling companions [118,119]. These accounts of plants and cultivation are generally consistent with the descriptions of Ólafsson and Pálsson, but also include a few additions[162]. Hooker also comments that a species of *Draba* (Brassicaceae) was growing abundantly on the roof of one of the houses on Aðalstræti, and that "… scurvy-grass (*Rumex acetosa* and *digynus*)" i.e. common and mountain sorrel (*R. acetosa* and *Oxyria digyna*), formed part of his diet while in Reykjavík[118].

Domestic animals in the town included cow, sheep, horse, and dog. The latter are identified by Hooker as *Canis islandicus* (Icelandic sheepdog) but there were also some mongrels thought to originate from Denmark[118,119]. MacKenzie states that in 1810 the horses of Reykjavík were poorly fed (possibly because it was late winter/early spring), and that in the hinterlands horses were suffering from an unspecified disease[119]. During one outing Hooker saw an Arctic fox (*Canis* *lagopus* i.e. *V. lagopus)*[118], the skins of which were sold in Reykjavík [118,119]. The only sea mammals observed were seals (*Phoca sp.*)[118].

Hooker considered the Raven (*Corvus corax*) common in Reykjavík[118]. He was also gifted the eggs of the Arctic tern (*Sterna hirundo* i.e. *Sterna paradisaea*) to eat while in Reykjavík, which were presumably acquired from somewhere close by[118]. Hooker noted in his description of the town that the harbour area and beach were used for drying stockfish. Molluscs could be found on the shores around Reykjavík included *Balanius sp.* (Balanidae), *Mya truncata*, *Venus islandica* i.e. *Arctica islandica*, *Lepas sp.* (Lepadidae), *Bulla sp.* (Bullidae) and *Turbines sp.* (Turbinidae)[118].

MacKenzie's descriptions of Reykjavík tend to be more general, but as a trading centre, there were a number of plant and animal derivatives to be found that were for sale on the streets. In terms of export there was oil (fish?), fish (stockfish?), tallow (animal fat), wool, butter, fox and swan skins (*V. lagopus* and *C. Cygnus* respectively), and horses[119]. These were exchanged for tobacco (*Nicotiana sp.*), spirits, grain (specifically rye; *Secale cereal*), linen (*L. usitatissimum*) and cotton (Malvaceae) goods. In the households of Reykjavík, Viðey and Nes, he noted the presence of bread (imported cereals), wine (potentially of various fruits), sugar (*Saccharum sp.*), sago (*Metroxylon sp.*), tea (*Camilla sinensis*), coffee (*Coffea sp.*), chocolate (*Theobroma cacao*), curry (probably representative of an array of spices), beef and ham, alongside the usual Icelandic staples of mutton, cheese and butter[119]. A particularly interesting feature is reference to a "*vaccine virus*" to inoculate children against smallpox (*Variola virus*). MacKenzie states this was brought with him from Britain[119]. Smallpox outbreaks occurred frequently in Iceland e.g. 1670, 1707 and 1785 with that of 1707–1709 particularly virulent and responsible for the death of over a quarter of the population of Iceland[111].

**Tables listing taxa reported from archaeological excavations and historical sources**

The tables in this section list the taxa identified in multiple zooarchaeological (Tables S9 – 11) and macrobotanical analyses (Tables S12) as well as the angiosperms (Tables S13-14), lichens and mosses (Table S15) and marine macroalgae (Table S16) mentioned in the historical accounts reviewed above.

**Table S9.** Mammal bone from Reykjavík, 870–1800 CE

| Taxa/Site | a & i) Tjarnagata 4 (late 9th century) | b) Aðalstræti 14-16 (late 9th century) | c) Aðalstræti 18 (870-1226 CE) | c) Suðurgata 3-5 (870-1226 CE) | d) Alþingisreitnum (870-1226 CE) | e) Lækjargata 10-12 (870-1226 CE) | d) Alþingisreitnum (1226-1500 CE) | f) Tjarnagata 3C (post 1500 CE) | d) Alþingisreitnum (1500-1800 CE) | g) Aðalstræti 10 (ca. 1650-1750 CE) | h) Aðalstræti 10-16 (pre-1764 CE) | h) Aðalstræti 10-16 (post-1764 CE) | i) Aðalstræti 8 (1700-1900 CE) | h) Aðalstræti 10-16 (1800-1900 CE) | d) Alþingisreitnum (post-1800 CE) | h) Aðalstræti 10-16 (post-1900 CE) |
|---|---|---|---|---|---|---|---|---|---|---|---|---|---|---|---|---|

| | | | | | | | | | | | | | | | | |
|---|---|---|---|---|---|---|---|---|---|---|---|---|---|---|---|---|
| **Cervidae** | | | | | | | | | | | | | | | | |
| *Cervus sp.* | | | | | | | | | ✓ | | | | | | | |
| **Bovidae** | | | | | | | | | | | | | | | | |
| *Bos Taurus* | ✓ | ✓ | ✓ | | ✓ | ✓ | ✓ | ✓ | ✓ | ✓ | ✓ | ✓ | ✓ | ✓ | ✓ | ✓ |
| *Ovis/capra* | ✓ | ✓ | ✓ | ✓ | ✓ | ✓ | ✓ | ✓ | ✓ | | | | ✓ | | ✓ | |
| *Capra hircus* | | | | | | | | | | ✓ | | | | | | |
| *Ovis aries* | ✓ | | | | ✓ | | | ✓ | ✓ | ✓ | ✓ | ✓ | | ✓ | ✓ | ✓ |
| **Equidae** | | | | | | | | | | | | | | | | |
| *Equus caballus* | ✓ | ✓ | | | ✓ | ✓ | ✓ | ✓ | ✓ | ✓ | | ✓ | ✓ | | ✓ | |
| **Suidae** | | | | | | | | | | | | | | | | |
| *Sus scrofa* | ✓ | ✓ | | | ✓ | ✓ | ✓ | ✓ | ✓ | ✓ | | ✓ | | | ✓ | ✓ |
| **Canidae** | | | | | | | | | | | | | | | | |
| *Vulpes lagopus* | | | | | | | ✓ | ✓ | ✓ | | | | | | | |
| *Canis familiaris* | | | | | ✓ | | ✓ | ✓ | ✓ | ✓ | | | ✓ | | ✓ | |
| **Felidae** | | | | | | | | | | | | | | | | |
| *Felis domesticus* | | | | | | | ✓ | | ✓ | | | | | | ✓ | |
| **Rodentia** | | | | | | | | | | | | | | | | |
| *Rattus sp.* | | | | | | | | | | | | | | | ✓ | ✓ |
| *Rattus norvegicus* | | | | | | | | | | ✓ | | | | | | |
| **Phocidae** | | | | | | | | | | | | | | | | |
| *Phoca sp.* | ✓ | | | | ✓ | ✓ | ✓ | ✓ | ✓ | ✓ | | | ✓ | | ✓ | |
| *Phoca vitulina* | ✓ | | | | ✓ | | ✓ | ✓ | ✓ | | | | | | ✓ | |
| *Erignathus barbatus* | | | | | | | ✓ | | | | | | | | | |
| **Odobenidae** | | | | | | | | | | | | | | | | |
| *Odobenus rosmarus* | ✓ | ✓ | | | ✓ | ✓ | | | | | | | | | | |
| **Cetacea** | | | | | | | | | | | | | | | | |
| *Cetacean sp.* | ✓ | | | ✓ | ✓ | | ✓ | ✓ | | | | | | | ✓ | |
| *Balaeoptera sp. (borealis?)* | ✓ | | | | | | | | | | | | | | | |

a) Grímsson & Einarsson 1970; b) Tinsley & McGovern 2001; c) Nordahl 1988; d) Pálsdóttir 2010; e) Hicks 2016; f) Perdikaris et al. 2002; g) Harrison et al. 2008; h) Tinsley & McGovern 2002; i) Amorosi 1996

#### Table S10. Bird bone from Reykjavík, 870–1800 CE[1]

| Taxa/Site | **Tjarnargata 4 (late 9th century)**<br>Grímsson & Einarsson 1970 & Amorosi 1996 | **Lækjargata 10-12 (870-1226 CE)**<br>Hicks 2016 | **Alþingisreitnum (870-1226 CE)**<br>Pálsdóttir 2010 | **Alþingisreitnum (1226-1500 CE)**<br>Pálsdóttir 2010 | **Tjarnagata 3C (post-1500 CE)**<br>Perdikaris et al. 2002 | **Aðalstræti 10 (ca. 1650-1750 CE)**<br>Harrison et al. 2008 | **Alþingisreitnum (1500-1800 CE)**<br>Pálsdóttir 2010 | **Alþingisreitnum (post-1800 CE)**<br>Pálsdóttir 2010 |
|---|---|---|---|---|---|---|---|---|
| **Podicipedidae** | | | | | | | | |
| *Podiceps auritus* | | | | √ | | | | |
| **Phalacrocoracidae** | | | | | | | | |
| *Phalacrocorax aristotelis* | | | | √ | | | | |
| *Phalacrocorax carbo* | | | | | | √ | | √ |
| **Anatidae** | | | | | | | | |
| *Anser sp.* | | | √ | √ | | √ | √ | |
| *Cygnus cygnus* | | | √ | | | √ | √ | |
| *Somateria mollissima* | | | | √ | | | √ | |
| *Mergus merganser* | | | | | | √ | | |
| *Melanitta nigra* | | | | | | √ | | |
| *Anas platyrynchus* | | | | | | √ | | |
| **Charadriidae** | | | | | | | | |
| *Pluvialis sp.* | | | | | | √ | | |
| *Pluvialis apricaria* | | | | | √ | | | |
| **Procellariidae** | | | | | | | | |
| *Fulmaris glacialis* | | | | | √ | | | |
| **Laridae** | | | | | | | | |
| *Larus sp.* | √ | | √ | √ | √ | √ | √ | |
| *Larus canus* | | | | | √ | | | |
| *Larus argentatus* | | | | | √ | | | |
| *Larus glaucesens* | √ | | | | | | | |
| *Larus marinus* | | | | | | | √ | |
| *Rissa tridactyla* | | | √ | √ | | √ | | |
| ***Alcidae*** | | | | | | | | |
| *Alcid sp.* | | | | | | √ | | |
| *Uria sp.* | √ | | √ | √ | | √ | √ | √ |
| *Uria aalge* | √ | | √ | | | | | |
| *Pinguinus impennis* | √ | √ | √ | | | | | |
| *Alca torda* | | | | | | √ | | |
| *Fratercula arctica* | √ | | √ | √ | √ | | √ | |
| **Sulidae** | | | | | | | | |
| *Morus bassanus* | | | | √ | | | √ | |
| **Accipitridae** | | | | | | | | |
| *Haliaeetus albicilla* | | | √ | | | | | |
| **Falconidae** | | | | | | | | |
| *Falco rusticolus* | | | √ | | | | | |
| **Phasianidae** | | | | | | | | |
| *Lagopus mutus* | | | | | | √ | √ | √ |
| *Gallus gallus* | | | | | | √ | | √ |
| ***Corvidae*** | | | | | | | | |
| *Corvus corax* | | | | | | | √ | |

| | |
|---|---|
| ***Troglodytidae*** | |
| *Troglodytes spp.* | ✓ |

#### Table S11. Insect remains from Alþingisreitnum, 870-1226 CE[1]

| **Taxa** (Konráðsdóttir 2010) | **Alþingisreitnum** (870-1226 CE) |
|---|---|
| **Pediculidae** | |
| *Pediculus humanus* | ✓ |
| **Trichodectidae** | |
| *Damalinia ovis* | ✓ |
| **Carabidae** | |
| *Bembidion bipunctatum* | ✓ |
| *Bembidion grapii* | ✓ |
| *Patrobus septentrionis* | ✓ |
| *Patrobus atrorufus* | ✓ |
| *Pterostichus adstrictus* | ✓ |
| *Calathus melanocephalus* | ✓ |
| **Hydrophilidae** | |
| *Cercyon sp.* | ✓ |
| *Cercyon littoralis* | ✓ |
| **Staphylinidae** | |
| *Omalium laeviusculum* | ✓ |
| *Omalium septentrionis* | ✓ |
| *Xylodromus concinnus* | ✓ |
| *Stenus sp.* | ✓ |
| *Lathrobium brunnipes* | ✓ |
| *Philonthus sp.* | ✓ |
| *Quedius sp.* | ✓ |
| *Quedis mesomelinus* | ✓ |
| *Atheta sp.* | ✓ |
| *Oxypoda sp.* | ✓ |
| **Elateridae** | |
| *Hypnoidus riparius* | ✓ |
| **Cryptophagidae** | |
| *Cryptophagus spp.* | ✓ |
| *Atomaria sp.* | ✓ |
| **Latridiidae** | |
| *Latridius sp.* | ✓ |
| *Latridius pseudominutus* | ✓ |
| *Corticaria sp.* | ✓ |
| **Ptinidae** | |
| *Tipnus unicolor* | ✓ |
| *Ptinus fur* | ✓ |
| **Scarabidae** | |
| *Aphodius lapponum* | ✓ |
| **Curculionidae** | |
| *Otiorhynchus arcticus* | ✓ |
| Otiorhynchus nodosus | ✓ |

#### Table S12. Macro-botanical remains from Reykjavík, 870-1800 CE[1]

| Taxa/Site | Tjarnagata 4: late 9th century Grímsson & Einarsson 1970 | Aðalstræti 14: 870-1226 CE Nordahl 1988 | Suðurgata 3-5: 870-1226 CE Nordahl 1988 | Lækjargata 10-12: 870-1226 CE Mooney & Guðmundsdóttir 2020 | Alþingisreitnum: 870-1226 CE Guðmundsdóttir 2010 | Alþingisreitnum: 1226-1500 CE Guðmundsdóttir 2010 | Alþingisreitnum: 1226-1500 CE Martin 2010* | Alþingisreitnum: 1500-1800 CE Guðmundsdóttir 2010 |
|---|---|---|---|---|---|---|---|---|
| Betulaceae | | | | | | | | |
| *Betula sp.* | | √ | | | √ | √ | | √ |
| *Betula pubescens* | √ | | | | | | | |
| Fagaceae | | | | | | | | |
| *Quercus sp.* | | | √ | | √ | √ | | √ |
| *Fagus sp.* | | | | | √ | | | |
| Oleaceae | | | | | | | | |
| *Fraxinus sp.* | | | | | √ | | | |
| *Fraxinus excelsior* | | | | | | √ | | √ |
| Salicaceae | | | | | | | | |
| *Salix sp.* | | | | | √ | √ | | √ |
| Rosaceae | | | | | | | | |
| *Sorbus sp.* | | | | | √ | √ | | √ |
| *Prunus sp.* | | | | √ | | | | |
| *Geum sp.* | | | √ | | | | | |
| Pinaceae | | | | | | | | |
| *Larix sp.* | | | | | √ | √ | | √ |
| *Larix decidua* | √ | | | | | | | √ |
| *Picea sp.* | | | | | √ | √ | | √ |
| *Pinus sp.* | | | | | √ | √ | | √ |
| *Pinus sylvestris* | | | | | √ | √ | | √ |
| Cupressaceae | | | | | | | | |
| *Juniperus communis* | | | | | | √ | | |
| Taxaceae | | | | | | | | |
| *Taxus baccata* | | | | | √ | | | |
| Cyperaceae | | | | | | | √ | |
| Poaceae | | | | | | | √ | |
| *Hordeum vulgare* | | | √ | √ | | | √ | |
| *Festuca sp.* | | | √ | | | | | |
| *Phleum pratense* | | | √ | | | | | |
| *Avena sp.* | | | | √ | | | | |
| Caryophyllaceae | | | | | | | | |
| *Stellaria media* | | | √ | √ | | | √ | |
| *Spergula arvensis* | | | √ | √ | | | | |

[1] Further data is available, but it is not clear which phase of the excavation that the sample and context numbers for these plant remains emanated from.

| | | | | | | | |
|---|---|---|---|---|---|---|---|
| Polygonaceae | | | | | | | |
| *Polygonum sp.* | | | ✓ | | | | |
| *Rumex sp.* | | | | | | | ✓ |

| Taxa/Site (continued) | Tjarnagata 4: late 9th century Grímsson & Einarsson 1970 | Aðalstræti 14: 870-1226 CE Nordahl 1988 | Suðurgata 3-5: 870-1226 CE Nordahl 1988 | Lækjargata 10-12: 870-1226 CE Mooney & Guðmundsdóttir 2020 | Alþingisreitnum: 870-1226 CE Guðmundsdóttir 2010 | Alþingisreitnum: 1226-1500 CE Guðmundsdóttir 2010 | Alþingisreitnum: 1226-1500 CE Martin 2010* | Alþingisreitnum: 1500-1800 CE Guðmundsdóttir 2010 |
|---|---|---|---|---|---|---|---|---|
| Lamiaceae | | | | | | | | |
| *Stachys sp.* | | | ✓ | | | | | |
| *Galeopsis sp.* | | | ✓ | ✓ | | | | |
| Asteraceae | | | | | | | | |
| *Matricaria inodora* | | | ✓ | | | | | |
| Violoideae | | | | | | | | |
| *Viola sp.* | | | | | | | ✓ | |
| Brassicaceae | | | | | | | | |
| *Thlaspi arvense* | | | ✓ | | | | | |
| Ericaceae | | | | | | | | |
| *Empetrum nigrum* | | | ✓ | ✓ | | | | |
| *Vaccinium uliginosum* | | | | ✓ | | | | |
| Fabaceae | | | | | | | | |
| *Medicago lupulina* | | | ✓ | | | | | |
| Linaceae | | | | | | | | |
| *Linum usitatissimum* | | | | ✓ | | | | |
| Menyanthaceae | | | | | | | | |
| *Menyanthes trifoliata* | | | | ✓ | | | | |
| Potamogetonaceae | | | | | | | | |
| *Potamogeton sp.* | | | | ✓ | | | | |
| Fucaceae | | | | | | | | |
| *Ascophyllum nodosum* | | | | ✓ | | | | |

#### Table S13. Plants cultivated in Reykjavík in the mid-18th and early 19th centuries

| *Latin | Common name | Ólafsson 1752 (1975) | Hooker 1809 (1811) | MacKenzie 1810 (1811) |
|---|---|---|---|---|
| *Secale cereale* | Rye | ✓ | | |
| *Hordeum vulgare* | Barley | ✓ | | |
| *Avena sativa* | Oats | ✓ | | |
| *Poaceae sp.* | Hybrid-grains | ✓ | | |
| *Linum usitatissimum* | Flax | ✓ | ✓ | |
| *Cannabis sativa* | Hemp | ✓ | ✓ | |
| *Lathyrus sp.* | Peas | ✓ | | ✓ |
| *Brassicaceae* | Mustard | | ✓ | ✓ |
| *Brassicaceae* | Cabbage | | ✓ | ✓ |
| *Brassicaceae* | Rutabaga | | ✓ | ✓ |
| *Brassicaceae* | White cabbage | ✓ | | |
| *Brassicaceae* | Kohlrabi | | | ✓ |
| *Brassicaceae* | Cauliflower | ✓ | | |
| *Brassicaceae* | Turnip | | | ✓ |
| *Brassicaceae* | Turnip-radishes | | ✓ | |
| *Raphanus sp.* | Radish | ✓ | ✓ | ✓ |
| *Daucus carota* | Carrots | | ✓ | |
| *Solanum tuberosum* | Potatoes | ? | ✓ | ✓ |
| *Nasturtium officinale* | Watercress | ✓ | ✓ | ✓ |
| *Thymus sp.* | Garden thyme | ✓ | | |
| *Salix sp.* | Willow | ✓ | | |
| *Sambucus sp.* | Elder | ✓ | | |
| Generic term | conifer | ✓ | | |

*The Latin nomenclature is inferred for the most part as the authors did not always use it to identify the plants they were discussing.

#### Table S14. Vascular plants observed by Hooker (1811)

| Page no. | *Latin | **Habitat | **Location | Comment |
|---|---|---|---|---|
| 48 | *Polypodium arvonicum* | rocks | Vatnsmýri | *Woodsia alpine* according to Kristinsson et al. 2018 |
| 20, 39 | *Lychnis alpina* | coast, upland | Arnarhóll, Skólavörðuholt | |
| 39 | *Saxifraga sp.* | upland | Skólavörðuholt | Hooker suggests *S. triscuspidata.* |
| 20 | *Saxifraga tricuspidata* | coast | Arnarhóll | |
| 20 | *Cardamine petraea* | coast | Arnarhóll | |
| 20 | *Draba incana* | coast | Arnarhóll | |
| 20 | *Draba contorta* | coast | Arnarhóll | |
| 20 | *Stellaria sp.* | coast | Arnarhóll | Hooker suggests *S. groenlandica.* |
| 20 | *Silene acaulis* | coast | Arnarhóll | |
| 20 | *Cerastium alpinum* | coast | Arnarhóll | |
| 39 | *Dryas octopetala* | upland | Skólavörðuholt | |
| 48 | *Rubus saxitalis* | rocks | Vatnsmýri | |
| 39 | *Vaccinium uliginosum* | upland | Skólavörðuholt | Used for tea |
| 39 | *Salix sp.* | upland | Skólavörðuholt | Two or three species. |
| 39 | *Salix lanata* | upland | Skólavörðuholt | |
| 48 | *Carex spp.* | wetland | Vatnsmýri | Two new species according to Hooker. |
| 20 | *Juncus trifidus* | coast | Arnarhóll | |
| 20 | *Juncus biglumis* | coast | Arnarhóll | |
| 21 | *Festuca vivipara* | coast | Arnarhóll | |

*The Latin nomenclature may not correspond with modern taxonomic categorisation.
**The locations and habitats are inferred from the descriptions of Hookers excursions (1811).

#### Table S15. Lichens and mosses observed by Hooker (1811)

| Page no. | *Latin | **Habitat | **Location | Comment |
|---|---|---|---|---|
| 19, 40 | *Trichostomum canescens* | coast, upland | Arnarhóll, Skólavörðuholt | |
| 48 | *Trichostomum ellipticum* | rocks | Vatnsmýri | |
| 20 | *Endocarpon tephroides* | coast | Arnarhóll | |
| 20 | *Lecidea sp.* | coast | Arnarhóll | A species unknown to Hooker. |
| 20 | *Lecidea geographica* | coast | Arnarhóll | |
| 20 | *Lecidea fusco-lutea* | coast | Arnarhóll | |
| 20 | *Cetraria islandica* | coast | Arnarhóll | |
| 20 | *Cetraria nivalis* | coast | Arnarhóll | |
| 20 | *Parmelia scrobiculata* | coast | Arnarhóll | |
| 20 | *Parmelia brunnea* | coast | Arnarhóll | |
| 20 | *Stereocaulon globiferum* | coast | Arnarhóll | |
| 20 | *Baeomyces endivifolius* | coast | Arnarhóll | |
| 20 | *Baeomyces vermicularis* | coast | Arnarhóll | |
| 40 | *Cornicularia sp.* | upland | Skólavörðuholt | Hooker says similar to *C. bicolor*, but grey and much larger. |
| 20 | *Encalypta lanceolata* | coast | Arnarhóll | |
| 20 | *Dicranum latifolium* | coast | Arnarhóll | |
| 20 | *Buxbaumia foliosa* | coast | Arnarhóll | |
| 20 | *Polytrichum hercynicum* | coast | Arnarhóll | |
| 39 | *Splachnum vasculosum* | upland | Skólavörðuholt | |

| | | | |
|---|---|---|---|
| 39 | *Splachnum mnioides* | upland | Skólavörðuholt |
| 48 | *Meesia dealbata* | wetland | Vatnsmýri |
| 48 | *Hypnum filamentosum* | rocks | Vatnsmýri |

*The Latin nomenclature may not correspond with modern taxonomic categorisation.
**The locations and habitats are inferred from the descriptions of Hookers excursions (1811).

#### Table S16. Marine Macroalgae observed by Hooker (1811)

| Page no. | *Latin | Comment | **Habitat | **Location |
|---|---|---|---|---|
| 31 | *Fucus Palmatus* | | | |
| 31 | *Fucus esculentus* | | | |
| 31 | *Fucus digitatus* | | | |
| 31 | *Fucus ciliatus* | | | |
| 31 | *Fucus dentatus* | | | |
| 31 | *Fucus purpurascens* | | | |
| 31, 65 | *Fucus saccharinus* | A variety of . | shore | Between Reykjavík & Laugarás. |
| 31 | *Fucus plumosus* | | | |
| 31 | *Fucus flagelliformis* | | | |
| 31 | *Fucus rubens* | | | |
| 31 | *Fucus ramentaceus* | Only found in Iceland. | | |
| 31 | *Conferva faeniculacea* | Of Hudson. | | |

*The Latin nomenclature may not correspond with modern taxonomic categorisation.
**The locations and habitats are inferred from the descriptions of Hookers excursions (1811).

#### Table S17. Birds observed by Hooker (1811)

| Page no. | Common name (Hooker 1811) | *Latin (Hooker 1811) | **Latin | Comment |
|---|---|---|---|---|
| 72 | Whooper swan | | *Cygnus cygnus* | |
| 72 | Eider duck | | *Somateria mollissima* | |
| 72 | Red-throated merganser | *Mergus serrator* | *Mergus serrator* | |
| 72 | Cormorant | | *Phalacrocorax carbo* | |
| 72 | Puffin | | *Fratercula arctica* | |
| 72 | Black guillemot | *Colymbus troile* | *Cepphus grylle* | |
| 66 | Tern sp. | *Sterna hirundo* | *Sterna paradisaea* | |
| 31 | Snipe | | *Gallinago gallinago* | |
| 273 | Phalarope sp. | *Phalaropus glacialis* | *Phalaropus sp.* | Red-necked phalarope (*Phalaropus lobatus*) |
| 24, 31 | Raven | | *Corvus corax* | |
| 31 | Common wagtail | | *Motacilla sp.* | White wagtail (*Motacilla alba alba*) |
| 31 | Snow bumting | | *Plectrophenax nivalis* | |

*These are Latin names given by Hooker.
** Latin nomenclature is inferred as the author did not always use it to identify the birds observed.

153. Björn Gunnlaugsson. Reikjavík og kríngum liggjandi Pláts. *Íslandskort.is; National Library of Iceland* https://islandskort.is/map/1122#?c=0&m=0&s=0&cv=0&r=0&xywh=-4150%2C0%2C12490%2C6728.

154. Horrebow, N. *The Natural History of Iceland.* (London, 1758).

155. Larsen, G. (2010) 3 Katla Tephrochronology and eruption history. In Schomacker, A., Krüger, J. and Kjar, K.H., Eds., Developments in Quaternary Sciences, 13, Elsevier, Amsterdam,

23-49. - References - Scientific Research Publishing. https://www.scirp.org/(S(czeh4tfqyw2orz553k1w0r45))/reference/referencespapers?referenceid=1102686.

156. Sæmundsson, K., Sigurgeirsson, M. Á. & Friðleifsson, G. Ó. Geology and structure of the Reykjanes volcanic system, Iceland. *J. Volcanol. Geotherm. Res.* 391, (2020).

157. Kennedy, J., Jónsson, S. Þ., Kasper, J. M. & Ólafsson, H. G. Movements of female lumpfish (Cyclopterus lumpus) around Iceland. *ICES J. Mar. Sci.* 72, 880–889 (2015).

158. Mehler, N., Kűchelmann, H. C. & Holterman, B. The export of gyrfalcons from Iceland during the 16th century; a boundless business in a proto-globalised world. in *Raptor and human – falconry and bird symbolism throughout the millennia on a global scale AU2 - Gersmann, K.H. AU2 - Grimm, O.* 995–1020 (Wachholtz Verlag – Murmann Publishers, Kiel/Hamburg, 2018).

159. Magnússon, Á. & Vídalín, P. *Jarðabók (Vol. III)*. (Hin Íslensk Fræðafelagið í Kaupmannahöfn & Möller SL, Kaupmannahöfn, 1926).

160. Vilmundarson, Þ. Heimildir um hafís á siðari öldum. in *Hafísnn AU2 - Einarsson, M. Á.* 313–332 (Almenna Bókafélagið, Reykjavík, 1969).

161. Lievog, R., L. Í.-. Kort og Grundtegning over Handel Stædet Reikevig i Island med angrændsende Huusmænds Pladser, som efter Kongelig Allernaadigste Befalning skal anlegges til Kiöbstæd. *Íslandskort.is; National Library of Iceland* https://islandskort.is/map/1123#?c=0&m=0&s=0&cv=0&r=0&xywh=-3969%2C0%2C13149%2C7083 (1787).

162. Ólafsson, E. *Ferðabók Eggerts Ólafsson Og Bjarna Pálsonnar Um Ferðir Þeirra á Íslandi Árin 1752-1757, II*. (Örn og Örlygur, Reykjavík, 1975).

163. Jóhannesson, T. *Agriculture in Iceland: Conditions and Characteristics*. (Agricultural University of Iceland, Iceland, 2010).
